# Trial-to-trial variability of spiking delay activity in prefrontal cortex constrains burst-coding models of working memory

**DOI:** 10.1101/2021.01.30.428962

**Authors:** Daming Li, Christos Constantinidis, John D. Murray

## Abstract

A hallmark neuronal correlate of working memory (WM) is stimulus-selective spiking activity of neurons in prefrontal cortex (PFC) during mnemonic delays. These observations have motivated an influential computational modeling framework in which WM is supported by persistent activity. Recently this framework has been challenged by arguments that observed persistent activity may be an artifact of trial-averaging, which potentially masks high variability of delay activity at the single-trial level. In an alternative scenario, WM delay activity could be encoded in bursts of selective neuronal firing which occur intermittently across trials. However, this alternative proposal has not been tested on single-neuron spike-train data. Here, we developed a framework for addressing this issue by characterizing the trial-to-trial variability of neuronal spiking quantified by Fano factor (FF). By building a doubly stochastic Poisson spiking model, we first demonstrated that the burst-coding proposal implies a significant increase in FF positively correlated with firing rate, and thus loss of stability across trials during the delay. Simulation of spiking cortical circuit WM models further confirmed that FF is a sensitive measure that can well dissociate distinct WM mechanisms. We then tested these predictions on datasets of single-neuron recordings from macaque prefrontal cortex during three WM tasks. In sharp contrast to the burst-coding model predictions, we only found a small fraction of neurons showing increased WM-dependent burstiness, and stability across trials during delay was strengthened in empirical data. Therefore, reduced trial-to-trial variability during delay provides strong constraints on the contribution of single-neuron intermittent bursting to WM maintenance.

**Significance Statement:** There are diverging classes of theoretical models explaining how information is maintained in working memory by cortical circuits. In an influential model class, neurons fire exhibit persistent elevated memorandum-selective firing, whereas a recently developed class of burst-coding models suggests that persistent activity is an artifact of trial-averaging, and spiking is sparse in each single trial, subserved by brief intermittent bursts. However, this alternative picture has not been characterized or tested on empirical spike-train data. Here we combine mathematical analysis, computational model simulation and experimental data analysis to test empirically theses two classes of models and show that the trial-to-trial variability of empirical spike trains is not consistent with burst coding. These findings provide constraints for theoretical models of working memory.

## Introduction

Working memory (WM) is a core cognitive function for temporary internal maintenance, and manipulation, of information across seconds-long delays. A key neuronal correlate of WM found by many studies is memorandum-selective spiking activity during mnemonic delays in neurons recorded from prefrontal cortex (PFC) of monkeys performing WM tasks (Funahashi et al., 1989; Miller et al., 1996; Romo et al., 1999; Leavitt et al., 2017). These common observations have motivated an influential class of theoretical models proposing neurobiological mechanisms for how PFC circuits could produce selective spiking activity patterns that can persist over seconds to support WM (Amit and Brunel, 1997; Wang, 1999, 2001; Lim and Goldman, 2013).

Recently, alternative proposals have challenged the role of persistent activity in WM on the basis that stable activity patterns may be inadequate to describe the dynamics of neural activity on single trials. In particular, Lundqvist et al. (2016a) contended that “this seemingly continuous delay activity may, however, reflect averaging across trials and/or neurons.” Their analysis of local field potential (LFP) recordings from PFC found intermittent bursts of gamma-band power on single trials, and argued that these findings provide evidence in favor of models of WM maintenance based on intermittent bursts of neuronal activity (Lundqvist et al., 2016a). Theoretical modeling studies have proposed potential mechanisms for how WM traces can be maintained without persistent spiking activity or stable activations across trials. In particular, short-term facilitation of synapses can maintain WM information, with neuronal spiking exhibiting sparse bursting through synaptic reactivation and adaptation (Mongillo et al., 2008; Lundqvist et al., 2011; Fiebig and Lansner, 2017; Mi et al., 2017).

These divergent proposals for mechanistic theories of WM function have contributed to to debate on the nature of WM delay activity in PFC (Constantinidis et al., 2018; Lundqvist et al., 2018). Prior studies characterizing the temporal stationarity of WM delay spiking activity were mostly based on trial-averaged firing rates, and therefore did not specifically examine the stability of those activations across trials (Murray et al., 2017; Spaak et al., 2017; Cavanagh et al., 2018). The burst-coding proposal, whose support comes across-trial instability of LFP power (Lundqvist et al., 2016a), has not been directly tested at the level of single-neuron spiking activity. It is therefore necessary to create a testing framework addressing this issue. Here we propose that stability across trials is a key feature differentiating these two classes of models, and characterize the spike count trial-to-trial variability measured by Fano factor (FF). FF has been shown to be associated with the engagement of neurons in behavioral tasks (Churchland et al., 2010; Hussar and Pasternak, 2010; Chang et al., 2012; Purcell et al., 2012a), and was shown in modeling studies to be higher in a clustered network with neurons transitioning between high and low activity states (Litwin-Kumar and Doiron, 2012; Deco and Hugues, 2012).

In this study, we analyzed a doubly stochastic Poisson model which allows different levels of spiking burstiness, and demonstrate that FF is a sensitive measure that can well dissociate bursting and non-bursting regimes. We show that if selective WM delay spiking activity were mediated by intermittent bursting, then testable implications include significant increase in FF and thus loss of stability across trials during the WM delay. Next, we simulate biophysically grounded spiking neural circuit models of WM maintenance, which can generate non-bursting persistent activity or intermittent bursting, to further confirm these results. Finally, we test the theoretical implications using single-neuron spike-train data recorded in macaque later PFC during three WM tasks of different modalities. In clear contrast to burst-coding model predictions, a population-level decrease in FF was observed in all the tasks, implying that across-trial stability is strengthened during delay. The magnitudes of FF were also smaller than burst-coding model predictions. The incompatibility between theoretical predictions and results from empirical spike-train data suggests that the extent to which intermittent burst spiking could contribute to WM delay activity is strongly limited.

## Materials and Methods

### Fano factor

Single-neuron spike-train data consists of multiple trials of recordings of spiking times under each task condition. Spiking activity is analyzed from the perspectives of intensity and variability across trials during each task epoch, leading to the concepts of mean firing rate and Fano factor (FF). These measures are both calculated using spike trains from multiple trials, and can be calculated for a specific time bin during the trial locked to a task event such as stimulus onset. We divide the time axis into bins of size Δ, and count the number of spikes that fall into each time bin in each single trial. Let *N* (*t*, Δ) be the spike count in the time window (*t, t* + Δ) in a trial. The mean firing rate *r*_mean_ at a given time bin under a stimulus condition is given by:

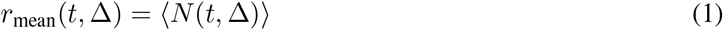

where 〈…〉 denotes the average calculated across trials. FF at a given time bin under a stimulus condition is defined by:

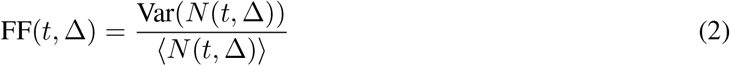

where the average and the variance are also both calculated across trials (Fano, 1947). FF is therefore a second-order statistic quantifying trial-to-trial variability. Low FF implies stability across trials, whereas high FF implies instability across trials. Note that the FF for a Poisson process is 1.

### Doubly stochastic Poisson model

Poisson spiking models are commonly used as statistical models of cortical spiking variability, and were motivated based on the observed proportional relationship between the mean and variance of neural responses (Tolhurst et al., 1981, 1983; Softky and Koch, 1993; Shadlen and Newsome, 1998; Oram et al., 1999). An inhomogeneous Poisson spiking process is a point process determined by a varying firing rate without random fluctuations, which has equal mean and variance of spike count. A doubly stochastic Poisson process is an extension to this with the underlying firing rate following another random process (Cox, 1955), which can model the spike count over-dispersion in empirical data (Churchland and Abbott, 2012). This paradigm is suitable for the purpose of this study, because it allows incorporation of the quiescent and bursty state transitions directly into the firing rate (Miller, 2006). In our study, the underlying firing rate follows a two-level telegraph process, as illustrated in Fig. 1a.

**Figure 1:**
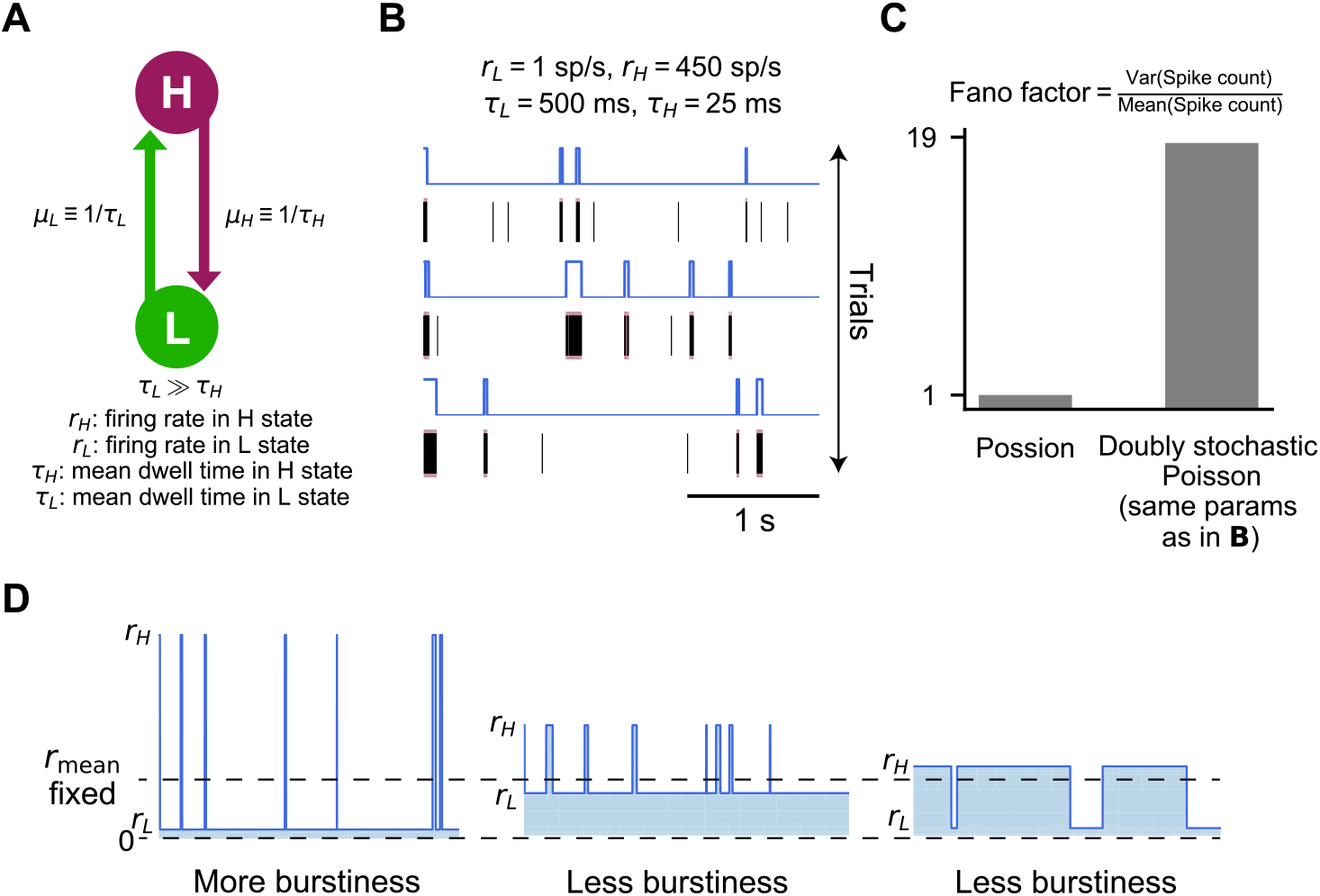
Two-state telegraph process as a doubly stochastic Poisson model of neuronal burst spiking. **(a)** Neuronal spiking is modeled as a doubly stochastic Poisson process with the underlying firing rate following a random two-level telegraph process. *H* and *L* denote the high and low firing states, with respective firing rates *r*_*H*_ and *r*_*L*_, transition rates leaving the states *μ*_*H*_ and *μ*_*L*_, *τ*_*H*_ and *τ*_*L*_ are the mean dwell time in each state. **(b)** Three sample trials of model-simulated telegraph processes with the corresponding generated spike trains. The model parameters used here are in a biophysically plausible range: *r*_*L*_ = 1 sp/s, *r*_*H*_ = 450 sp/s, *τ*_*L*_ = 500 ms, *τ*_*H*_ = 25 ms. Colored bursts typically occur in the high firing state. **(c)** Any homogeneous Poisson process with constant firing rate has Fano factor (FF) equal to 1, whereas the example doubly stochastic Poisson process, defined in **(b)**, has FF equal to 18.6. **(d)** The model in general can generate different levels of burstiness at fixed mean firing rate *r*_mean_ by tuning the four parameters. The intermittent bursting regime is characterized by low *r*_*L*_, high *r*_*H*_, and low 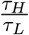. Less burstiness can be achieved by raising τ_*L*_ the low firing rate *r*_*L*_, or by increasing the ratio 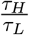.

In the two-level telegraph process model, the two hidden states, denoted *L* (for low) and *H* (for high), have firing rates *r*_*L*_ and *r*_*H*_, respectively. The transition rates leaving the respective states are denoted *μ*_*H*_ and *μ*_*L*_. Equivalently, we introduce the mean dwell time of each state: *τ*_*H*_ and *τ*_*L*_, which are related to the transition rates by 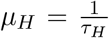 and 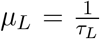. The dwell time in each state follows exponential distribution, with mean equal to *τ*_*H*_ and *τ*_*L*_, respectively. The four parameters {*r*_*L*_*, r*_*H*_, *τ*_*L*_, *τ*_*H*_} fully define the spiking process, and are same across trials under one stimulus condition.

A key advantage of this simple model is that it can be solved analytically. With the model parameters defined above, one have immediately the mean firing rate:

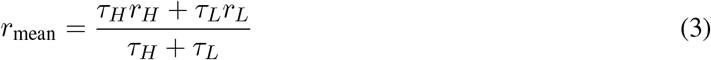

If we call the time-dependent firing rate *λ*(*t*), and denote *N* (*t*, Δ) the spike count in the time window (*t, t* + Δ), then we have:

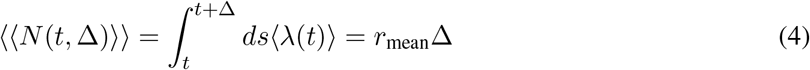

where the inner average is with respect to the fluctuations of the firing rate, and the outer average is with respect to the Poisson process. By the properties of Poisson process, the second moment of the spike count is given by:

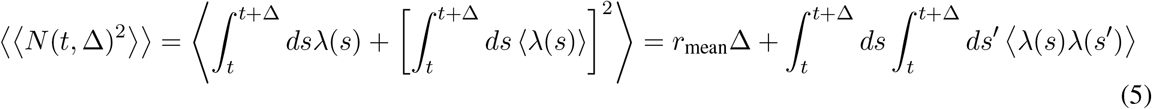

Hence the variance of the spike count is given by:

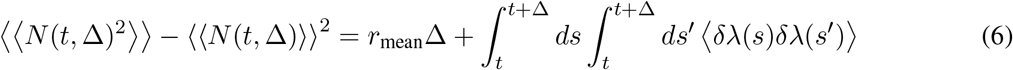

where *δλ*(*t*) = *λ*(*t*) − 〈*λ*(*t*)〉.

The autocovariance of the underlying telegraph firing rate process has the form:

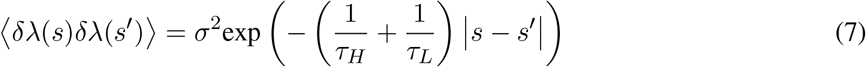

where 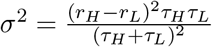. Therefore the characteristic time of the exponential decay is 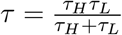. Direct calculation of the integral gives:

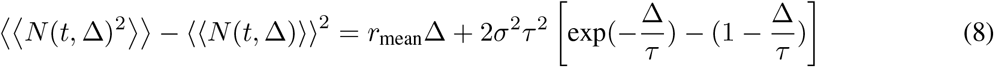

Therefore, FF is given by:

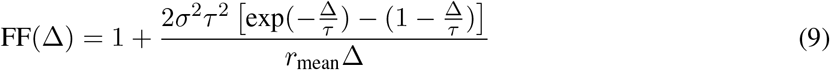

FF therefore depends on the four parameters {*r*_*L*_, *r*_*H*_, *τ*_*L*_, *τ*_*H*_} and on the bin size Δ.

### Circuit model simulations

We simulated three biophysically-based cortical circuit models to test that FF is a sensitive measure that can differentiate distinct WM mechanisms. Compte et al. (2000) built a recurrent circuit model with synaptic mechanisms for stable activity in spatial WM processes, in which leaky integrate-and-fire neurons were arranged on a ring with the angular location corresponding to the preferred cue position. Non-bursting persistent activity was maintained by strong recurrent connections. Here, we adopt a spatially uniform discrete node attractor network model based on similar mechanism applied to a visual motion discrimination decision making task (Wang, 2002) (Model 1 below), first introduced by Brunel and Wang (2001). Non-bursting persistent activity realized by stable attractor networks typically fails to reproduce irregular spiking activity (Barbieri and Brunel, 2008; Durstewitz and Gabriel, 2007; Renart et al., 2007). To address this issue, we also included an attractor model with balanced slow excitation and fast inhibition implementing a negative derivative feedback mechanism, introduced by Lim and Goldman (2013), that generates persistent irregular firing patterns during WM delays (Model 2 below). For the generation of bursting in selective WM activity, Lundqvist et al. (2011) proposed that bursting can be realized through the competition between cellular adaptation and synaptic augmentation in a cyclic attractor network. Similarly, we added spike-frequency adaptation and short-term facilitation to Model 1 and obtained intermittent bursting (Model 3 below).

### Model 1: Stable attractor network

The model for generating non-bursting persistent activity is the same as described in Wang (2002) (see also, Brunel and Wang, 2001), with minor modification to the parameter values. There are three excitatory pyramidal cell pools with in total 1600 pyramidal cells, with two selective to the stimuli (240 neurons for each) and the third one nonselective (1120 neurons), as well as an inhibitory interneuron pool (400 neurons). Both pyramidal cells and interneurons are described by leaky integrate-and-fire neurons and are characterized by a resting potential *V*_L_ = −70 mV, a firing threshold *V*_th_ = −50 mV, a reset potential *V*_reset_ = −55 mV, a membrane capacitance *C*_m_ = 0.5 nF for pyramidal cells and 0.2 nF for interneurons, a membrane leak conductance *g*_L_ = 25 nS for pyramidal cells and 20 nS for interneurons, and a refractory period *τ*_ref_ = 2 ms for pyramidal cells and 1 ms for interneurons. The corresponding membrane time constants are *τ*_m_ = *C*_m_*/g*_L_ = 20 ms for excitatory cells and 10 ms for interneurons. Below the threshold, the membrane potential *V* (*t*) of each unit follows

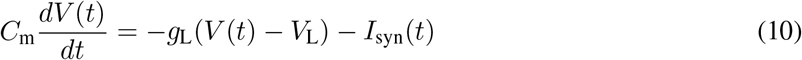

where *I*_syn_(*t*) represents the total synaptic current flowing into the cell.

The network consist of pyramid-to-pyramid, pyramid-to-interneuron, interneuron-to-pyramid, and interneuron-to-interneuron connections. Recurrent excitatory postsynaptic currents (EPSCs) have two components mediated by AMPA and NMDA receptors, respectively. External synaptic inputs send to the network all the information (stimuli) received from the outside world, as well as background noise due to spontaneous activity outside the local network. In simulations, external EPSCs were mediated exclusively by AMPA receptors. The total synaptic currents are given by

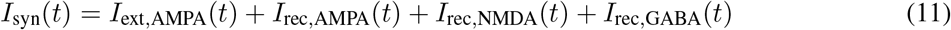

in which

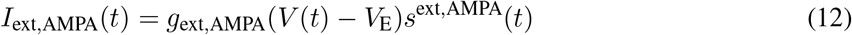

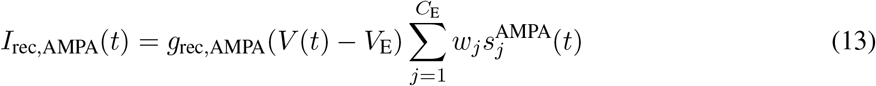

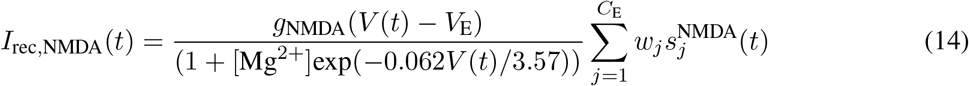

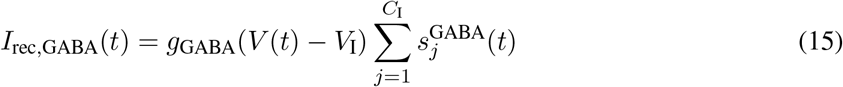

where *V*_E_ = 0 mV, *V*_I_ = −70 mV. The dimensionless weights *w*_*j*_ represent the structured excitatory recurrent connections (see below), the sum over *j* represents a sum over the synapses formed by presynaptic neurons *j*. NMDA currents have a voltage dependence that is controlled by extracellular magnesium concentration, [Mg^2+^] = 1 mM. The gating variables, or fraction of open channels *s*, are described as follows.

The AMPA (external and recurrent) channels are described by

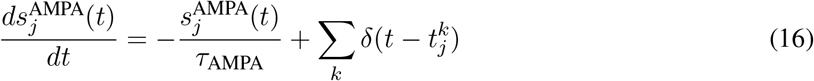

where the decay time of AMPA currents is taken to be *τ*_AMPA_ = 2 ms, and the sum over *k* represents a sum over spikes emitted by presynaptic neuron *j*. In the case of external AMPA currents, the spikes are emitted according to a Poisson process with rate *ν*_ext_ = 2.3 kHz independently from cell to cell. NMDA channels are described by

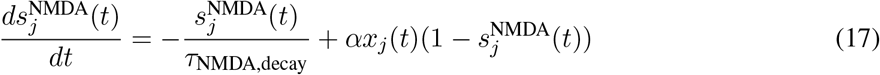

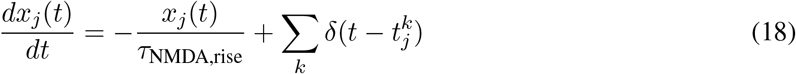

where the decay time of NMDA currents is taken to be *τ*_NMDA,decay_ = 100 ms, *α* = 0.5 ms^−1^, and *τ*_NMDA,rise_ = 2 ms. Last, the GABA synaptic variable obeys

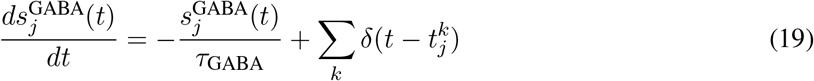

where the decay time constant of GABA currents is taken to be *τ*_GABA_ = 5 ms. Note that the very fast rise times (< 1 ms) of both AMPA and GABA currents are neglected. All synapses have a latency of 0.5 ms.

We used the following values for the recurrent synaptic conductances (in nS) in the *N* = 2000 neurons network: for pyramidal cells, *g*_ext,AMPA_ = 2.07, *g*_rec,AMPA_ = 0.05, *g*_NMDA_ = 0.165, and *g*_GABA_ = 1.3; for interneurons, *g*_ext,AMPA_ = 1.62, *g*_rec,AMPA_ = 0.04, *g*_NMDA_ = 0.13, and *g*_GABA_ = 1.0.

The coupling strength between a pair of neurons is prescribed according to a “Hebbian” rule: the synapse is strong (weak) if in the past the two cells tended to be active in a correlated (anticorrelated) manner. Hence, inside a selective pool, *w*_*j*_ = *w*_+_, where *w*_+_ > 1 is a dimensionless parameter that is equal to the relative strength of potentiated synapses with respect to the baseline. We used *w*_+_ = 1.84. Between two different selective pools, and from the nonselective pool to selective ones, *w*_j_ = *w*_−_, where *w*_−_ < 1 measures the strength of synaptic “depression.” We used *w*_−_ = 0.852. Other connections have *w*_j_ = 1. We chose the stimulus strength to be *μ* = 38 Hz and the task coherence to be 50%. We simulated 10 trials with the same timing of the task epochs as in the experiments (a foreperiod of 1000 ms, a stimulus onset of 500 ms, and a delay of 3000 ms), and then performed the analysis in the excitatory pool that was stimulated by the preferred cue.

### Model 2: Stable attractor network with negative derivative feedback

We use the same spiking network as described in Lim and Goldman (2013), with 1600 pyramidal neurons, 400 interneurons, and 2000 neurons in the external pool (not shown). The random connection probability *p* is set to 0.5. The remaining parameters are the same as listed in Lim and Goldman (2013). This network implements a negative derivative feedback mechanism through balanced slow excitation and fast inhibition. A brief stimulus is given to the external pool, and the negative derivative feedback provides a corrective signal that maintained delay activity. Simulation was done by running the MATLAB code available at ModelDB (entry 181010l; https://senselab.med.yale.edu/ModelDB/ShowModel?model=181010). We simulated 10 trials with a foreperiod of 1000 ms, a stimulus onset of 100 ms, and a delay of 4000 ms.

### Model 3: Burst-firing cyclic attractor network (Model 1 with addition of short-term facilitation and spike-frequency adaptation)

To create WM delay activity maintained by intermittent bursts, we use a synaptic model allowing a cyclic attractor. Motivated by the model of Lundqvist et al. (2011), we add spike-frequency adaptation mediated by calcium-activated potassium channels and NMDA short-term facilitation to the pyramidal leaky integrate-and-fire neurons to Model 1. First, we add a spike-frequency adaptation current to Equation 10:

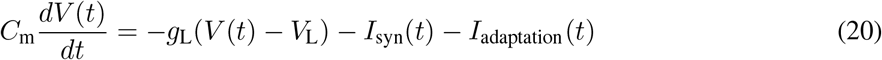

where

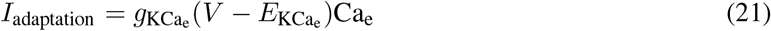

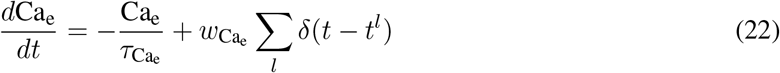

with *l* the index of spikes. The calcium ions decay exponentially at a fast timescale 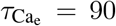 ms, and upon each firing there is an amount of calcium influx 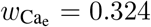. Next we add short-term facilitation to NMDA synapses between pyramidal neurons, so that Equations 17 and 18 now read:

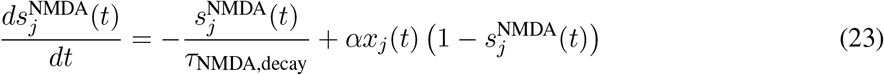

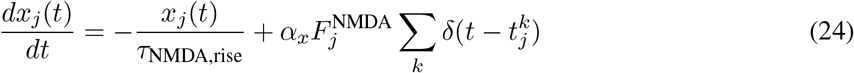

and

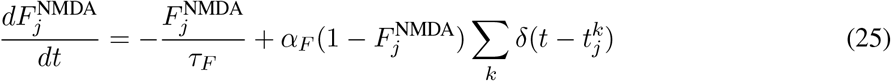

with *α*_x_ = 300, *α*_*F*_ = 0.008, and *τ*_*F*_ = 3000 ms, such that *F*^NMDA^ decays on a much longer timescale compared to adaptation. To obtain strong intermittent bursts, we modify the previously defined parameters such that *w*_+_ = 5.14, *w*_−_ = 0.5, *ν*_ext_ = 1.9 kHz, and *μ* = 152 Hz, with the rest being the same.

Again, we simulated 10 trials with the same timing of the task epochs as in the experiments (a foreperiod of 1000 ms, a stimulus onset of 500 ms, and a delay of 3000 ms), and then performed the analysis in the excitatory pool that was stimulated by the preferred cue. For each trial, we shifted the delay activity by a random amount between 0 and 2000 ms, to account for the fact that bursts of spiking may not be phase-locked with the stimulus onset in empirical data.

### Working memory tasks and datasets

We applied our analysis framework on previously collected single-neuron spike-train datasets recorded from monkey PFC during three WM tasks of different modalities: an oculomotor delayed response (ODR, visual) task (Constantinidis et al., 2001), a vibrotacticle delayed discrimination (VDD, somatosensory) task (Romo et al., 1999), and a spatial match/non-match (MNM, visual) task (Riley et al., 2018).

The ODR data was collected from two macaque monkeys (COD, MAR), recorded from the dorsolateral PFC (areas 8 and 46) of the left hemisphere from both monkeys. The ODR task (Fig. 4a) has eight stimuli for angular locations (0°, 45°, 90°, 135°, 180°, 225°, 270°, 315°). The subject fixates on a central point for a period of 1 s, and a visuospatial cue of one of these spatial angles is presented for 0.5 s, followed by a 3-s mnemonic delay. After the delay, the subject makes a saccadic eye movement to the remembered location.

The VDD data was collected from three macaque monkeys (R13, R14, R15), recorded from the inferior convexity of the PFC of the hemispheres on the contralateral side. The VDD task (Fig. 4b) data consists of two set of stimuli for vibrotactile frequencies: (10 Hz, 14 Hz, 18 Hz, 22 Hz, 26 Hz, 30 Hz, 34 Hz) for monkey R13, and (10 Hz, 14 Hz, 18 Hz, 24 Hz, 30 Hz, 34 Hz) for monkeys R14 and R15. The analysis results of these two sets of VDD stimuli were finally concatenated, as we were only interested in characterizing spiking trial-to-trial variability with respect to tuning, while the exact frequencies were not important. After a 1-s foreperiod, the subject receives a 0.5-s vibrotactile stimulus of variable mechanical frequency (cue 1, f1) to the finger, followed by a 3-s mnemonic delay. After the delay, a second stimulus (cue 2, f2) is presented and the subject reports, by level release, which stimulus had a higher frequency.

The MNM data was collected from three male rhesus monkeys (ADR, ELV, NIN), recorded from areas 8a, 8b, 9, 10, 9/46, 45, 46, and 47/12 of the lateral PFC of the right hemispheres of ADR and NIN, and from both hemishpheres of ELV. In the MNM task (Fig. 4c), a 2° square stimulus was presented in one of nine possible locations (labeled by angular locations similar to ODR, plus one at the center) arranged in a 3 × 3 grid with a 10° distance between adjacent stimuli. The subject fixates for a period of 1 s, followed by a stimulus onset of duration 0.5 s. After a mnemonic delay of 1.5 s, a second stimulus is shown for 0.5 s either matching the first in location or of a different location. After a second delay of 1.5 s, the subject makes a saccade to one of the two choice targets representing match and non-match shown respectively on an additional display.

In those experiments, typically 5-20 trials (correct and incorrect) for each stimulus were recorded from each neuron. During the recordings, neurons were not pre-selected for tuning properties.

### Data preprocessing

All the data manipulations and statistical tests were done using scripts written in Python. We followed a procedure similar to what was described in Barak et al. (2010) for filtering neurons for analysis. We only included correct trials in the analysis. For each dataset, neurons with mean firing rate lower than 1 sp/s (spike per second) in all task epochs (foreperiod, cue, delay) were removed. Trials were excluded if > 1% of the interspike intervals were < 1 ms. Neurons for which > 10% of the trials met this criterion were removed. The VDD dataset contains many unstable recordings that result in a drifting firing across trials. To exclude such neurons, stability of the recordings was assessed by performing a t-test on the spike counts of the first versus the second half of the trials during the forperiod. Neurons for which *P* < 0.05 were removed. The ODR and MNM datasets suffer much less from this issue, and only fewer than five such neurons were excluded by a sanity check with visual inspection when performing the analysis. In each dataset, neurons with at least 8 correct trials per stimulus condition were kept for analysis. This minimal filtering step was only based on data quality, but not related to tasks or tuning, and should not introduce sampling bias to the study. This procedure eventually yielded 642 neurons for the ODR task, 667 neurons for the VDD task, and 1009 neurons for the MNM task.

### Data analysis

When analyzing the experimental datasets, we divided the time axis into bins of Δ = 250 ms, which is relatively long compared to proposed burst durations. If there was no spike in a given time bin in any trial under a certain stimulus condition, then that time bin was not included in the analysis. In the analysis of delay activity, the first time bin immediately after the cuing period was not included to account for the potential non-zero relaxation time. ODR and MNM neurons with stimulus-selective delay activity were selected using one-way ANOVA (*P* < 0.05) over spike counts across stimulus conditions. VDD neurons with stimulus-selective delay activity were selected by testing a linear dependence between the mean firing rates during delay and the stimulus frequency strengths (*P* < 0.05). Neurons with at least consecutive 500 ms of selective activity during delay were selected in this procedure, which we also called neurons with well-tuned delay activity, and the analysis was done only on the time bins with stimulus-selective activity. Under each stimulus condition, *r*_mean_ and FF were computed during foreperiod and delay respectively, then were averaged across time bins to yield single scalars. For each single neuron, the stimulus under which *r*_mean_ was highest was called the preferred stimulus, while that under which *r*_mean_ was smallest was called the least preferred stimulus. We compared FF under these two stimulus conditions during delay to the baseline mean FF during the foreperiod. We defined neurons with increased FF under the preferred stimulus those whose FF was higher during delay under the preferred stimulus than both in the foreperiod and during delay under the least preferred stimulus. We also counted neurons with FF and *r*_mean_ significantly positively correlated across stimulus conditions (Pearson r > 0, *P* < 0.05), by concatenating all the time bins with well-tuned activity during delay.

To take into account of the temporal dynamics of coding and FF, we also compared FF in each single time bin during delay to the foreperiod baseline, in which case the preferred and the least preferred stimulus were defined according to the firing rate in each time bin.

When analyzing the simulated data of Model 1 and Model 3, we computed FF during foreperiod and delay, only for the neural pool that was activated by the stimulus, without discarding the time bin immediately after the cuing period. When analyzing the simulated data of Model 2, we computed FF during the last 500 ms of the foreperiod, and the last 3000 ms during the delay, for pyramidal neurons with non-zero firing rates during these periods. The time bin size used was still 250 ms.

## Results

### Types of stability of spiking activity during working memory

There are various candidate neural mechanisms for working memory (WM) delay activity (Barak and Tsodyks, 2014; Chaudhuri and Fiete, 2016; Murray et al., 2017). For instance, stable attractor networks implementing a positive feedback mechanism allow excitation to reverberate in a recurrent network, thereby storing information (Compte et al., 2000; Wang, 2002). Burst-coding models, on the other hand, rely on synaptic mechanisms such as synaptic facilitation with information encoded in calcium kinetics (Lundqvist et al., 2011). Attractor networks are not limited to discrete or one-dimensional attractors. For instance, chaotic attractor models can generate more complex patterns subserved by highly irregular dynamic attractors (Pereira and Brunel, 2018). There are also transient memory mechanisms such as the feedforward chains, where the signal decays rapidly at each unit but lifetime becomes longer when passed along the line (Goldman et al., 2008).

These models can generate memorandum-selective neural states that account for different types of stability of WM. We here propose that models of WM delay activity can be qualitatively assessed from two distinct properties (Table 1). One property is stationarity across time. This can be assessed by analysis of the trial-averaged peristimulus time histogram (PSTH), with stationarity defined such that the mean firing rate during delay keeps roughly at a constant level. The other property is stability across trials, which is assessed by the variability of spiking activity across trials. These two properties provide axes that allow us to classify existing WM models, and guide us choosing a measure to dissociate divergent mechanisms.

**Table 1:** Qualitative characterization of various working memory models in terms of stationarity across time and stability across trials.

|  | Stationarity<br>across time | Stability<br>across trials |
| --- | --- | --- |
| Stable attractor models<br>(e.g. Compte et al., 2000) | High | High |
| Feedforward chain models<br>(e.g. Goldman, 2009) | Low | High |
| Burst-coding models<br>(e.g. Lundqvist et al., 2011) | High | Low |
| Chaotic attractor models<br>(e.g. Pereira and Brunel, 2018) | High | Low |

For instance, the stable attractor models, the burst-coding models, and the chaotic attractor models of WM are all featured by stationarity across time (Compte et al., 2000; Lundqvist et al., 2011; Pereira and Brunel, 2018), whereas the feedforward chain model of WM exhibits non-stationarity through its temporal variations of trial-averaged firing rate (Goldman et al., 2008). On the other hand, the burst-firing and the chaotic attractor models are featured by the sparseness of spiking within single trials, which imply that the bursts of spikes are mostly not aligned across trials, resulting in low levels of across-trial stability. In contrast, the stable attractor model and the feedforward chain model exhibit higher levels of stability across trials, as within each single trial, the stable attractor is characterized by sustained activity, and the feedforward chain model is characterized by consistent transient dynamics following stimulus onset.

PSTH time courses with and without strong temporal variations have both been widely observed in empirical PFC recordings (Constantinidis et al., 2001; Brody et al., 2003; Tiganj et al., 2018), and these findings can test the stationarity of neuronal responses in relation to theoretical models (Murray et al., 2017). Because PSTH analyses rely on trial-averaging, they are not able to test the stability of neuronal firing across trials, which can provide important constraints to theoretical proposals for burst-firing. We therefore focus in this study on characterizing across-trial stability in PFC recordings and in computational models.

### Fano factor as a measure of trial-to-trial variability

To assess stability across trials, we focus on characterizing spike count trial-to-trial variability measured by Fano factor (FF) (Fano, 1947; Shadlen and Newsome, 1998; Miller, 2006), which has been shown to be related to neural engagement in cognitive tasks (Churchland et al., 2010; Hussar and Pasternak, 2010; Chang et al., 2012; Purcell et al., 2012b). Poisson processes are widely chosen spiking models based on the observed proportional relationship between the mean and variance of spiking responses in cortical neurons (Tolhurst et al., 1981, 1983; Softky and Koch, 1993; Shadlen and Newsome, 1998; Oram et al., 1999). A simple extension for neuronal spiking is an inhomogeneous Poisson model, determined by a time-varying firing rate without randomness, which has equal mean and variance of spike counts within the same time window, and thus FF equal to 1. In a stable attractor network, excitatory input stabilizes a specific attractor, so that spiking variability is quenched. This is likely to result in sub-Poisson statistics and FF smaller than 1 (Barbieri and Brunel, 2008; Durstewitz and Gabriel, 2007; Renart et al., 2007; Roudi and Latham, 2007). Burst-coding models, on the other hand, have two sources of variability: the variability of the firing rates and the variability of spikes (Miller, 2006; Churchland et al., 2011). Therefore, the extra variability is likely to result in super-Poisson statistics and FF larger than 1. This motivates us to build a doubly stochastic Poisson spiking model with the firing rate randomly transitioning between high and low activity states to capture the main characteristics of the alternative burst-coding proposal.

### A doubly stochastic Poisson spiking model

We first aimed to build a parsimonious model that can be parametrically tuned to reach either the stable or burst-spiking regimes. This allows us to make predictions on how FF behaves differently in the two distinct scenarios. A doubly stochastic Poisson process is an extension to the classic Poisson model, with the underlying firing rate following another random process, which can capture firing rate fluctuations in addition to spiking variability. This paradigm is suitable for the purpose of this study, because it allows incorporation of the low-activity quiescent and high-activity bursty state transitions directly into the firing rate. Burst-coding WM models hypothesize that each single trial consists of sparse sharp bursts of spiking, while during most of the time, neurons are in a baseline state of low-rate firing. According to this proposal, the memorandum selectivity of WM-related delay activity arises from modulation of these burst events (Lundqvist et al., 2016a, 2018). These proposals motivated us to translate these features into a doubly stochastic Poisson spiking model, where neuronal spiking is a Poisson process with the underlying firing rate following another two-level telegraph random process, as illustrated in Fig. 1a.

Here the two firing states *L* (for low) and *H* (for high or bursting) have firing rates *r*_*L*_ and *r*_*H*_, respectively. In the burst-coding scenario, *r*_*L*_ corresponds to the background firing that is homogeneous across all stimulus conditions, and should be relatively low. Bursts are most likely to occur when the neuron is in the high firing state *H*, with *r*_*H*_ ≫ *r*_*L*_. We also introduce the mean dwell time in each state: *τ*_*H*_ and *τ*_*L*_. Since the bursts are claimed to be sparse, we have *τ*_*L*_ ≫ *τ*_*H*_. The four parameters {*r*_*L*_, *r*_*H*_, *τ*_*L*_, *τ*_*H*_} fully define the spiking process, and are the same across trials under one stimulus condition. Sample trials of a two-level telegraph process with their generated spike trains are shown in Fig. 1b. Based on the aforementioned considerations, we ignored possible ramping of mean firing rate *r*_mean_ which occur at a slower timescale, and focused on characterizing the across-trial variability (FF) of spike counts, which does not depend on slow modulation of the mean firing rate as a function of time from delay period onset.

Note that this model can also produce a non-bursting regime at given mean firing rate by flexibly tuning the parameters, for instance by raising the low-state background firing rate *r*_*L*_, or by increasing the mean dwell-time ratio 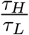 (Fig. 1d). *r*_mean_ and FF of this model can be analytically solved in the general form (with derivations shown in the Materials and Methods section).

### Theoretical implications of burst-coding models

If WM delay were modulated by sharp intermittent bursts, there would be the following important implications derived from the doubly stochastic Poisson model in the burst-coding regime, which can be tested in empirical WM neuronal recordings.

### Implication 1: Burst strength must be very strong if the burst coding assumptions were valid

If WM delay coding were instantiated as sparse intermittent bursts, we can estimate how strong the burst would need to be. For this purpose, we transformed Equation 3 and expressed the burst strength (firing rate in the high state) *r*_*H*_ as a function of the mean firing rate *r*_mean_, the mean dwell time ratio 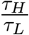 and the firing rate in the low state *r*_*L*_:

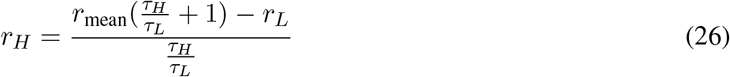

If WM were subserved by sharp intermittent bursts, we would have *r*_*H*_ ≫ *r*_*L*_ and 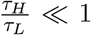. Fig. 2a is a visualization of the magnitudes of *r*_*H*_ by varying other parameters in a biophysically reasonable range consistent with burst-coding model assumptions. We fixed the firing rate in the low state *r*_*L*_ = 1 sp/s (spike per second), and varied *r*_mean_ between 10 and 100 sp/s and the mean dwell time ratio 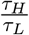 between 0.01 and 0.1 which imposes sparsity. It can be seen that the implied burst strength *r*_*H*_ tends to be very high, even exceeding the biophysical limit when both the observed *r*_mean_ is high and the bursts are sparse 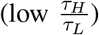. This provides a first strong constraint on the burst sparsity of delay activity: the bursts must be very strong and cannot be very sparse if the observed mean firing rate is high.

**Figure 2:**
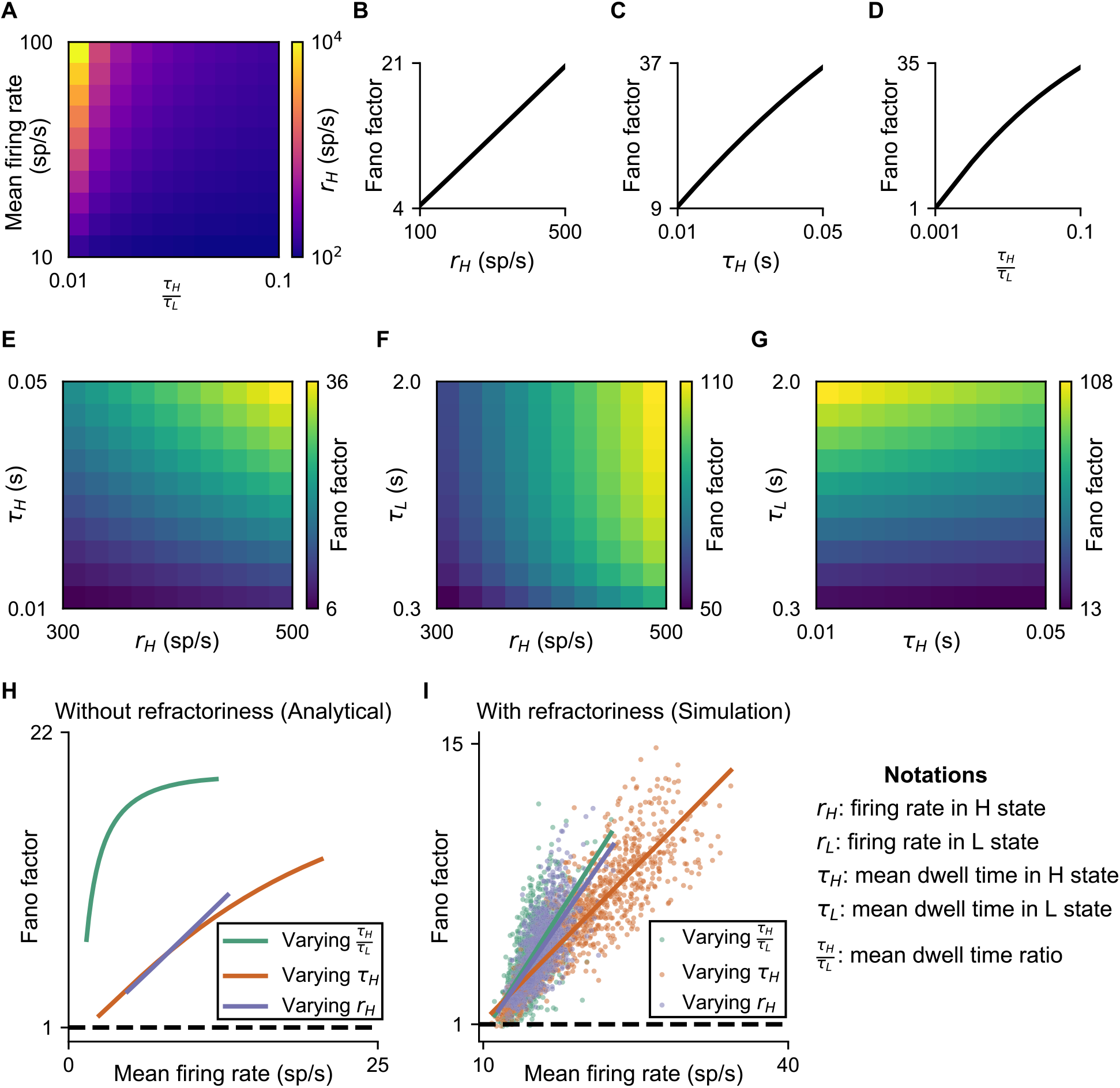
Theoretical predictions of the intermittent burst-coding model. **(a)** Firing rate in the high state should be very high under the sparse intermittent burst-coding scenario, given fixed firing rate in the low state. Heatmap of *r*_*H*_ is shown as a function of the mean firing rate *r*_mean_ and the dwelltime ratio 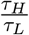 according to equation 26, with firing rate in the low state fixed. Parameters were chosen in a biophysically reasonable range: *r*_*L*_ = 1 sp/s is fixed; *r*_mean_ is varied between 10 and 100 sp/s as typically observed in experiments; 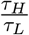 is varied between 0.01 and 0.1 as implied by sparsity. **(b)** FF grows with burst strength *r*_*H*_, with *r*_*L*_ = 1 sp/s, *τ*_*H*_ = 25 ms, *τ*_*L*_ = 500 ms, and time-bin width Δ = 250 ms fixed. **(c)** FF grows with burst duration *τ*_*H*_ in the sparse coding regime, with *r*_*L*_ = 1 sp/s, *r*_*H*_ = 500 sp/s, 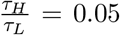 and Δ = 250 ms fixed. **(d)** FF grows with burst rate 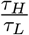 in the sparse coding regime, with *r*_*L*_ = 1 sp/s, *r*_*H*_ = 500 sp/s, *τ*_*L*_ = 500 ms, and Δ = 250 ms fixed. **(e–g)** Magnitudes of FF are large within a biophysical range with *r*_*L*_ = 1 sp/s, Δ = 250 ms, varying (*r*_*H*_, *τ*_*H*_), (*r*_*H*_, *τ*_*L*_), and (*τ*_*L*_, *τ*_*H*_) respectively, with the mean firing rate *r*_mean_ = 30 sp/s fixed. **(h)** FF increases monotonically with the mean firing rate. When varying 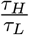 between 0.001 and 0.025, we fixed *r*_*L*_ = 1 sp/s, *r*_*H*_ = 450 sp/s, *τ*_*H*_ = 25 ms, and Δ = 250 ms. When varying *τ*_*H*_ between 5 ms and 75 ms, we fixed *r*_*L*_ = 1 sp/s, *r*_*H*_ = 200 sp/s, *τ*_*L*_ = 500 ms, and Δ = 250 ms. When varying *r*_*H*_ between 80 sp/s and 600 sp/s, we fixed *r*_*L*_ = 1 sp/s, *τ*_*H*_ = 25 ms, *τ*_*L*_ = 800 ms, Δ = 250 ms. **(i)** FF is still positively correlated with the mean firing rate after adding refractoriness into model simulation by allowing only the interspike intervals greater than 2 ms. Varying 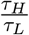: Pearson r = 0.64 (*P* < 10^−110^). Varying *τ*_*H*_: Pearson r = 0.87 (*P* < 10^−300^). Varying *r*_*H*_: Pearson r = 0.71 (*P* < 10^−150^). The parameters used in simulation are the same as in **(h)**.

### Implication 2: Fano factor should be very large if the burst-coding assumptions were valid

It can be shown that FF as a function of the model parameters in general is given by:

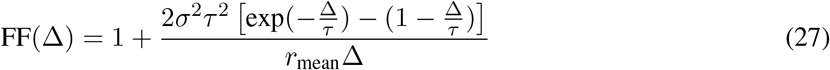

with Δ the time bin size, 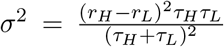 and 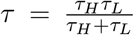 (see Materials and Methods section for derivation). Notice that here FF must be greater than 1 because of the extra variability due to the fluctuations of the underlying firing rate, compared to homogeneous Poisson model whose FF is equal to 1 (Fig. 1c).

Equation 27 has important neurophysiological implications. In the fast rate fluctuations limit (Δ ≫ *τ*) that we are interested in, FF saturates to 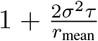. Exhibiting sharp intermittent bursts require *r*_*H*_ ≫ *r*_*L*_ and *τ*_*H*_ ≪ *τ*_*L*_, so that *τ* is close to *τ*_*H*_. We chose a time bin size of 250 ms for analysis, which is long enough compared to the mean burst duration. Fig. 2b–i show the properties of FF implied by equation 27, in a biophysical parameter range. If we fix *r*_*L*_ = 1 sp/s and *r*_mean_ to be what is usually observed in empirical data (30 sp/s for instance), burst-coding assumptions imply very large magnitudes of FF (Fig. 2e–g). These conclusions remain qualitatively the same under reasonable modification of parameters. These results suggest that if burst-coding assumptions applied, across-trial stability would be low as reflected in a high FF.

### Implication 3: Fano factor should be positively correlated with the mean firing rate if the burst-coding assumptions were valid

Equation 27 also implies that FF and *r*_mean_ are positively correlated under the burst-coding assumptions. With other model parameters fixed, FF grows with burst strength *r*_*H*_, burst duration *τ*_*H*_, and burst rate 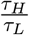 (Fig. 2b–d). We emphasize that the monotonic dependence on burst duration *τ*_*H*_ and burst rate 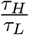 must be in the sparse burst-coding regime, such as 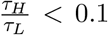. In that limit, FF is positively correlated with *r*_mean_ (Fig. 2h), which means that FF is larger for the neuron’s the preferred stimuli. We also tested the robustness of this result under the introduction of refractoriness following each spike, by running simulations of the model allowing only interspike intervals greater than 2 ms, and still found the significant positive correlation between FF and firing rate (Fig. 2i).

### Trial-to-trial variability measured by Fano factor dissociates distinct circuit mechanisms

The doubly stochastic Poisson model is based on the mathematical abstraction of the burst-coding assumptions. Although it captures the most important properties of the burst-coding mechanism, the simplicity of the model does not include underlying biophysical processes included in circuit models of WM maintenance. We therefore sought to validate that our FF measures can be used to distinguish different dynamic regimes of WM delay activity in influential circuit models. We simulated three spiking circuit models to generate both bursting and non-bursting WM delay activities, and confirm that FF is a sensitive measure that can well dissociate WM circuit models instantiating distinct mechanisms.

Attractor networks have been the most popular class of models for information storage (Wang, 2001). To generate non-bursting persistent activity, we used the spatially uniform stable attractor network model developed by Wang (2002) with minor modifications, focusing on the delay activity in the neural pool that was activated under the stimulus (Model 1, Fig. 3a, see Materials and Methods for details). Information is maintained by strong recurrent connections between neurons within the same selective neuronal pool. We adopted this discrete node model architecture because it allows generation of localized delay activity in both non-bursting and bursting scenarios (Model 3 below), facilitating direct comparison.

**Figure 3:**
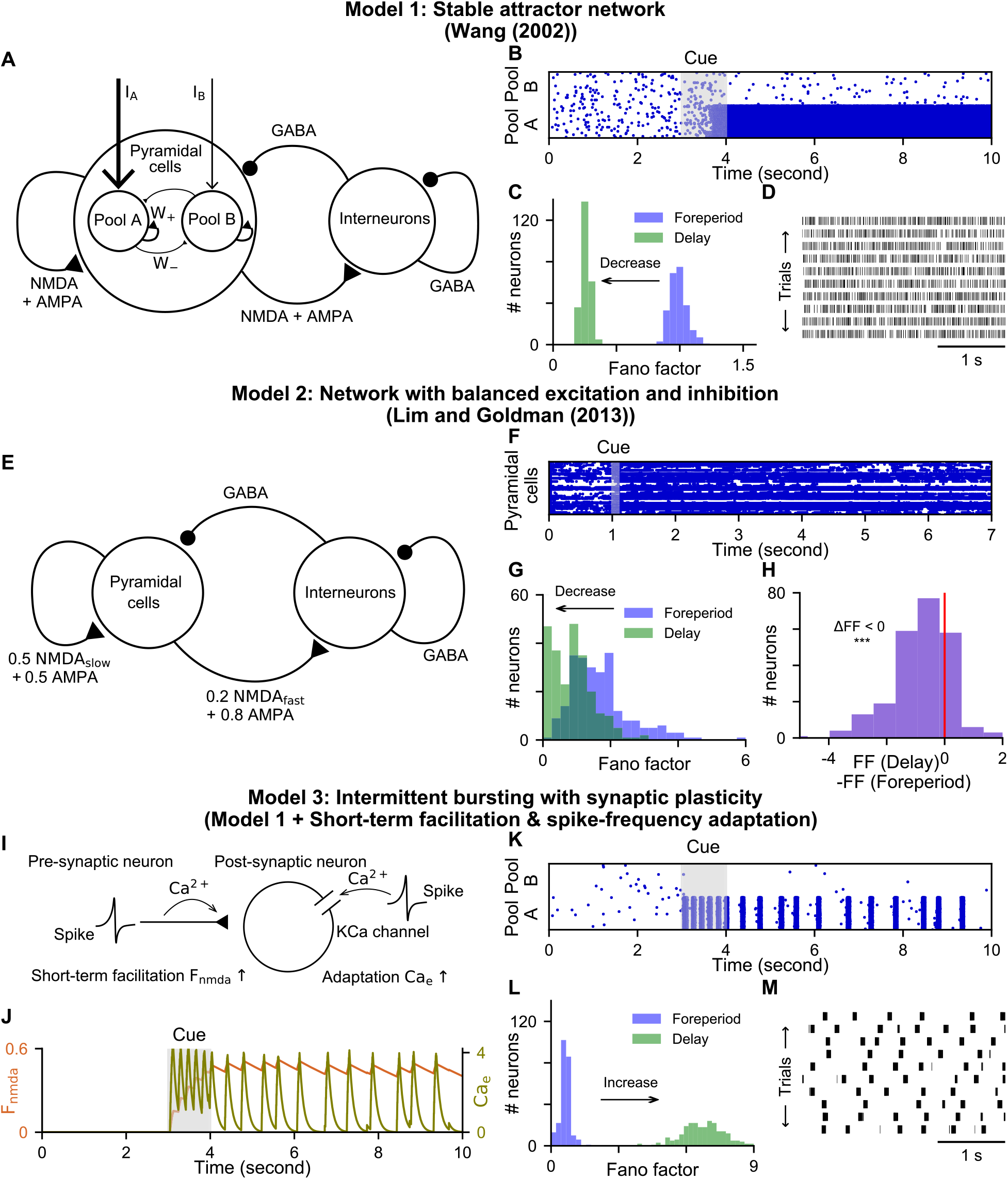
Numerical simulations confirm that FF is a sensitive measure which can dissociate distinct circuit mechanisms for WM delay activity. **(a)** The architecture of the spatially uniform discrete node spiking network model generating persistent activity (Wang, 2002). There are three pyramidal neural pools with two (A and B) selective to stimulus and one nonselective (not shown), as well as an interneuron pool. The two selective pools have strong recurrent excitation within themselves, mediated by NMDA and AMPA receptors. The two pools have lateral inhibition between them mediated by the GABAergic inhibitory population. Pool A is of interest in the study as it has elevated persistent activity under the stimulus. **(b)** Spike raster plot across a subset of neurons (40 from each pool) in a single trial with long-lasting persistent activity in Pool A during delay (0-3 s: foreperiod; 3-4 s: stimulus onset; 4-10 s: delay). **(c)** With non-bursting persistent activity, there is a global reduction in FF during delay compared to foreperiod. **(d)** Spike raster plot across trials during delay (only 3 seconds displayed) for a single neuron with non-bursting persistent activity. The mean firing rate with chosen simulation parameters is around 50 sp/s. **(e)** The architecture of the network implementing a negative derivative feedback mechanism (Lim and Goldman, 2013). **(f)** Spike raster plot across a subset of pyramidal neurons (80) in a single trial with long-lasting persistent activity realized by negative derivative feedback (0-1 s: foreperiod; 1-1.1 s: stimulus onset; 1.1-7.1 s: delay). **(g)** There is a global reduction in FF during delay compared to foreperiod. **(h)** Histogram of change in FF across neurons. FF during delay is significantly smaller than that during the foreperiod (*P* < 10^−26^). **(i)** Intermittent bursting is obtained by adding spike-frequency adaptation and short-term facilitation on top of the model in **(a)**. **(j)** Temporal evolution of the adaptation current and facilitation variables (0-3 s: foreperiod; 3-4 s: stimulus onset; 4-10 s: delay). **(k)** Spike raster plot across a subset of neurons (40 from each pool) in a single trial with long-lasting stable intermittent bursting in Pool A during delay. **(l)** With intermittent bursting, there is a global increase in FF from foreperiod to delay. **(m)** Spike raster plot across trials during delay (only 3 seconds displayed) for a single neuron with intermittent bursting. The mean firing rate with chosen simulation parameters is around 40 sp/s. Significance level: *: *P* < 0.05; **: *P* < 0.01; ***: *P* < 0.001

Stable attractor networks are known to produce a reduction of spiking variability to sub-Poisson levels in neurons participating in the persistent WM activity, because excitatory and inhibitory synaptic inputs can become unbalanced in a high-activity WM state (Barbieri and Brunel, 2008; Durstewitz and Gabriel, 2007; Renart et al., 2007; Roudi and Latham, 2007). For instance, overall excitatory input can stabilize one specific attractor state, resulting in reduced variability. To address this issue, we simulated a network which realizes stable persistent delay activity through balanced slow excitation and fast inhibition (Lim and Goldman, 2013) (Model 2, Fig. 3e, see Materials and Methods for details). Such a network implements a negative derivative feedback mechanism which generates spikes in a persistent way, while preserving Poisson-like irregular firing which is driven by fluctuations of subthreshold mean synaptic input when excitation and inhibition cancel each other out.

To generate transient bursts, on the other hand, we adopted a synaptic mechanism for maintaining WM traces over relatively long intervals. Mongillo et al. (2008) proposed that WM signals can be maintained through the mechanism of short-term synaptic facilitation, mediated by increased residual calcium levels at presynaptic terminals. In the model proposed by Lundqvist et al. (2011), neuronal spike-frequency adaptation reduces the lifetime of high-activity WM states, which generates short bursts of spiking, while short-term synaptic facilitation allows synapses to maintain a selective WM trace which causes reactivation of the associated pyramidal-neuron pool. These combination of faster spike-frequency adaptation and slower synaptic facilitation generates a WM code consisting of intermittent bursting in a selective pyramidal-neuron pool. To implement this proposed mechanism in our models, here we incorporated a spike-frequency adaptation current and short-term synaptic facilitation in the NMDA synapses in the excitatory neurons to Model 1 (Model 3, Fig. 3i, see Materials and Methods for details). In this scenario, WM information is maintained by spike-induced changes in synaptic weights in between bursts of selective spiking activity.

We compared simulated spike trains displaying non-bursting persistent activity and intermittent bursts in the neural population that is responsive to the stimulus. With the parameters chosen, all three models generated long-lasting WM delay activity (Fig. 3b,f,k). Sample single neuron raster plots were shown for Model 1 and Model 3, which are spatially uniform models (Fig. 3d, m). With Model 1, we found that there was a global reduction in FF during the delay compared to foreperiod, and neurons with strongest persistent activity fired with low variability with FF much smaller than 1 (Fig. 3c). With Model 2, there was also a shrink in FF during delay compared to foreperiod (Fig. 3g,h). For the bursting Model 3, Fig. 3j shows the temporal evolution of the facilitation and adaptation variables. In sharp contrast to Model 1 and 2, there was a dramatic increase in FF during they delay compared to foreperiod with (Fig. 3l). The magnitudes of FF in the biophysical Model 3 were significantly smaller than those in the doubly stochastic Poisson model. This is because in our mathematical model, the burst duration and the inter-burst intervals are completely random and follow exponential distribution, whereas synaptic biophysical models simulating real neuronal processes had much less flexibility and more constraints including refractoriness. Nonetheless, both mathematical and biophysical models yielded the same qualitative predictions. These simulation results from in a biophysically grounded circuit models further confirmed the conclusions in the previous section, thereby justifying that FF is a sensitive measure which can dissociate diverging WM mechanisms, and can be used to test on experimental data.

### Three spike-train datasets recorded in monkey lateral PFC during working memory tasks

We analyzed single neuron spike train data recorded from lateral PFC of macaque monkeys in three parametric WM tasks: an oculomotor delayed response (ODR) task (Constantinidis et al., 2001) (Fig. 4a), a vibrotacticle delayed discrimination (VDD) task (Romo et al., 1999) (Fig. 4b), and a spatial match/non-match (MNM) task (Riley et al., 2018) (Fig. 4c). The three tasks differ in several features, allowing us to test the generality of our findings. They differ in stimulus modality (visual for ODR and MNM vs. somatosensory for VDD), role of WM in guiding behavioral response (veridical report of location for ODR vs. binary discrimination for VDD vs. binary match/nonmatch report for MNM), and prototypical stimulus tuning curves of single PFC neurons (bell-shaped for ODR and MNM vs. monotonic for VDD). Delay activity in the ODR task can be attributed to response preparation in some neurons (Funahashi et al., 1993; Markowitz et al., 2015), whereas the VDD and MNM task delay activity are free of this premotor signal as the action is not specified until after the WM delay. The tasks have a 0.5-s cue epoch followed by a 3-s (ODR and VDD) or a 1.5-s (MNM) delay epoch, which are relatively long and allow characterization of time-varying WM representations.

**Figure 4:**
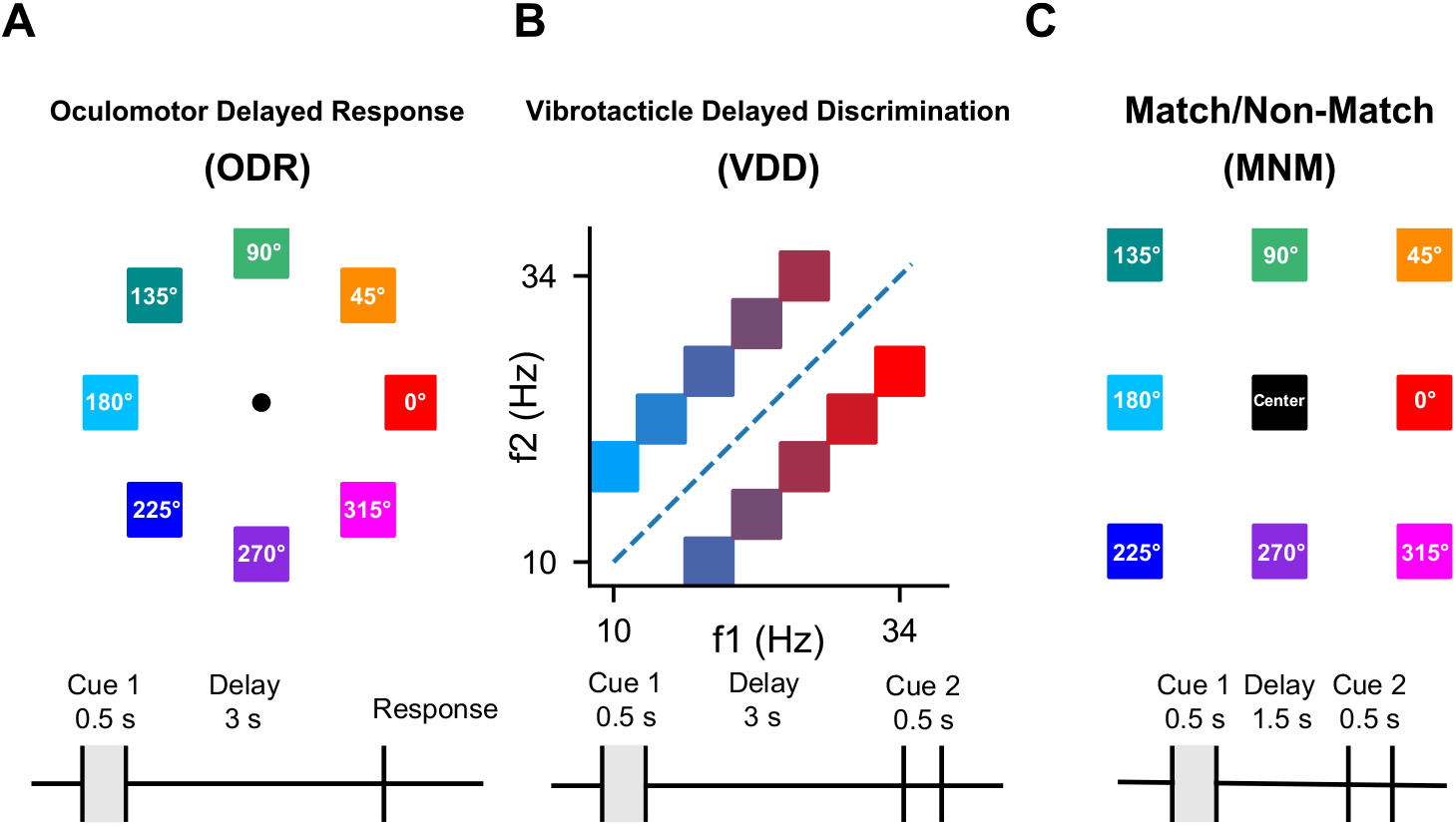
Working memory tasks. **(a)** In the oculmotor delayed response (ODR) task, the subject fixates on a central point for a period of 1 s, and a visuospatial cue of variable spatial angle is presented for 0.5 s, followed by a 3-s mnemonic delay. After the delay, the subject makes a saccadic eye movement to the remembered location (Constantinidis et al., 2001). **(b)** In the vibrotactile delayed discrimination (VDD) task, after a foreperiod of 1 s, the subject receives a 0.5-s vibrotactile stimulus of variable mechanical frequency (cue 1, f1) to the finger, followed by a 3-s mnemonic delay. After the delay, a second stimulus (cue 2, f2) is presented and the subject reports, by level release, which stimulus had a higher frequency (Romo et al., 1999). **(c)** In the spatial MNM task, the subject fixates for a period of 1 s, followed by a stimulus which is shown for 0.5 s. After a delay of 1.5 s, a second stimulus is shown for 0.5 s, either matching the first in location or of a different location. After a second delay of 1.5 s (not analyzed), the subject makes a saccade to one of the two choice targets representing match and non-match shown respectively on an additional display (not shown) (Riley et al., 2018).

### Fano factor observed in PFC inconsistent with burst-coding models

The key results we have established so far with statistical and biophysical modeling is that if WM delay activity were encoded as sharp intermittent bursts, there should be a dramatic increase in FF during delay compared to the foreperiod, and FF should be positively correlated with *r*_mean_, with FF highest under the preferred stimulus. To test if these predictions are true in empirical spiking data, we analyzed three single-neuron spike-train datasets recorded in lateral PFC during WM tasks described in the previous section, and characterized FF during different task epochs under each stimulus condition.

We selected neurons with at least consecutive 500 ms of stimulus-selective delay activity (details in Materials and Methods), and only analyzed time bins with stimulus-selective activity in correct trials. We divided task epochs into time bins of 250 ms, and computed *r*_mean_ and FF in each time bin under each stimulus condition. We defined the preferred and least-preferred stimulus conditions according to *r*_mean_ in the corresponding task epoch. A baseline level of FF was computed during the foreperiod by averaging across time for each neuron. Next, we measured FF averaged across trials during the delay under each stimulus condition, and compared to the baseline level. Correlation between *r*_mean_ and FF was computed as the Pearson correlation by concatenating all stimulus-selective delay-period time bins.

We sought to identify neurons exhibiting task-related burstiness, and therefore classified all selected neurons with criteria based on the theoretical implications of the burst-coding models examined in the sections above. Fig. 5 summarizes, for each task, the numbers of neurons with well-tuned delay activity that meet the following criteria: (i) higher FF for the preferred stimulus during delay than for the foreperiod and for the least-preferred stimulus during delay; (ii) FF > 3 (a threshold for the magnitude) for the preferred stimulus; (iii) positive correlations between *r*_mean_ and FF across stimulus conditions during delay (*P* < 0.05); and the intersections between these categories. Burt-coding models predict that neurons should be at the intersections of these categories. It can be seen from the Venn diagram, however, that numbers of neurons at the intersections are very small compared to the number of well-tuned neurons in each task (ODR: 8 of 179; VDD: 2 of 153; MNM: 0 of 75). Conversely, most of the well-tuned neurons are in none of these categories (ODR: 113 of 179; VDD: 106 of 153; MNM: 41 of 75).

**Figure 5:**
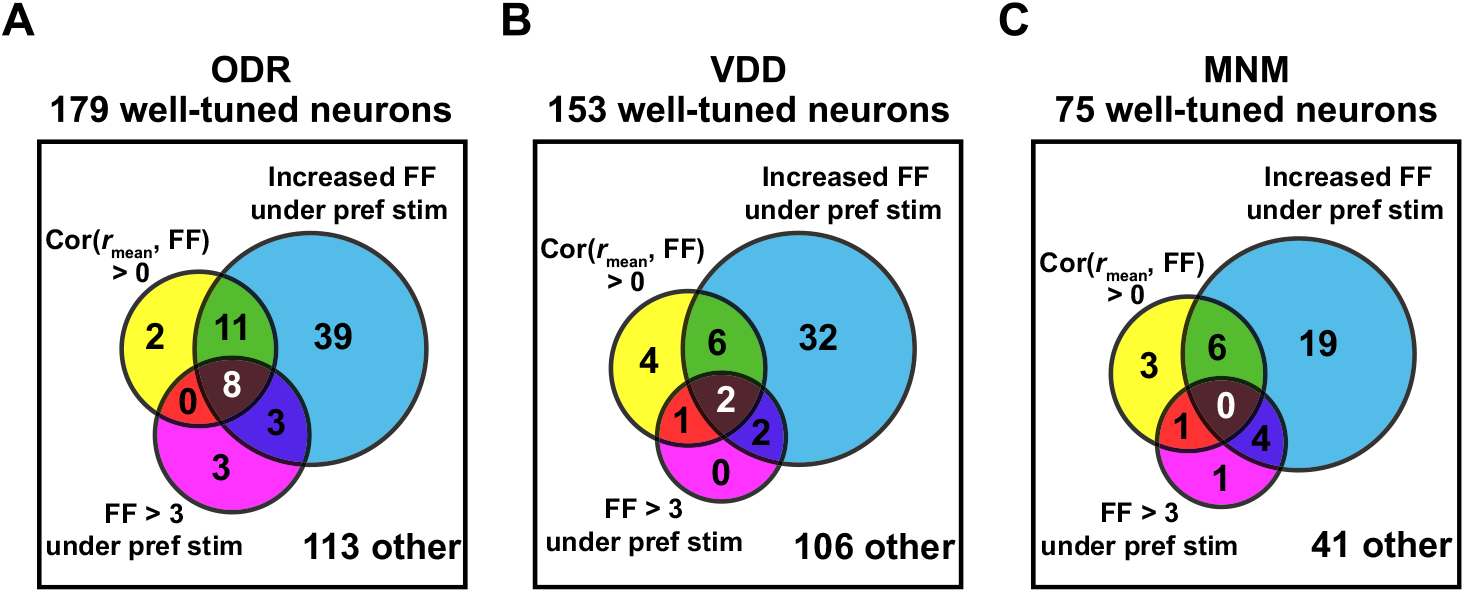
Venn diagram showing for each task (**a - c**) the number of neurons categorized as: displaying significant positive correlation between *r*_mean_ and FF, having FF > 3 under the preferred stimulus, having FF under the preferred stimulus higher than those in the foreperiod and under the least preferred stimulus (termed “Increased FF under pref stim” in the figure), and the combination of intersections between all three categories. The total number of well-tuned neurons and those that are in none of these categories are also shown. Numbers in each segment are exclusive.

Fig. 6 displays, for each task, plots for a single neuron with non-bursting persistent activity (Fig. 6a,c,e), and for the single neuron with maximal bursting behaviors chosen from neurons at the intersections of the three categories defined above (Fig. 6b,d,f). For completeness, we include such plots for all neurons with well-tuned delay activity in the Extended Data. It can be seen in general that stationarity across time is not a universal feature of WM delay activity: the mean firing rates can have trends such as ramping (Fig. 6a,c). We noticed that even among the neurons chosen from the intersections of the Venn diagram, firing patterns were still not as bursty as hypothesized by burst-coding models. For instance, the neuron in Fig. 6f has very low firing rates, and its sparsity was not due to burstiness but due to weak spiking activity. In Fig. 6d, the neuron has high FF during the foreperiod and under the least preferred stimulus during the delay. Therefore, it is an intrinsically bursty neuron, and the burstiness is not related to the task (Compte et al., 2003; Shinomoto et al., 2009). In addition, higher FF can be due to weak drift in overall levels of firing rate across trials.

**Figure 6:**
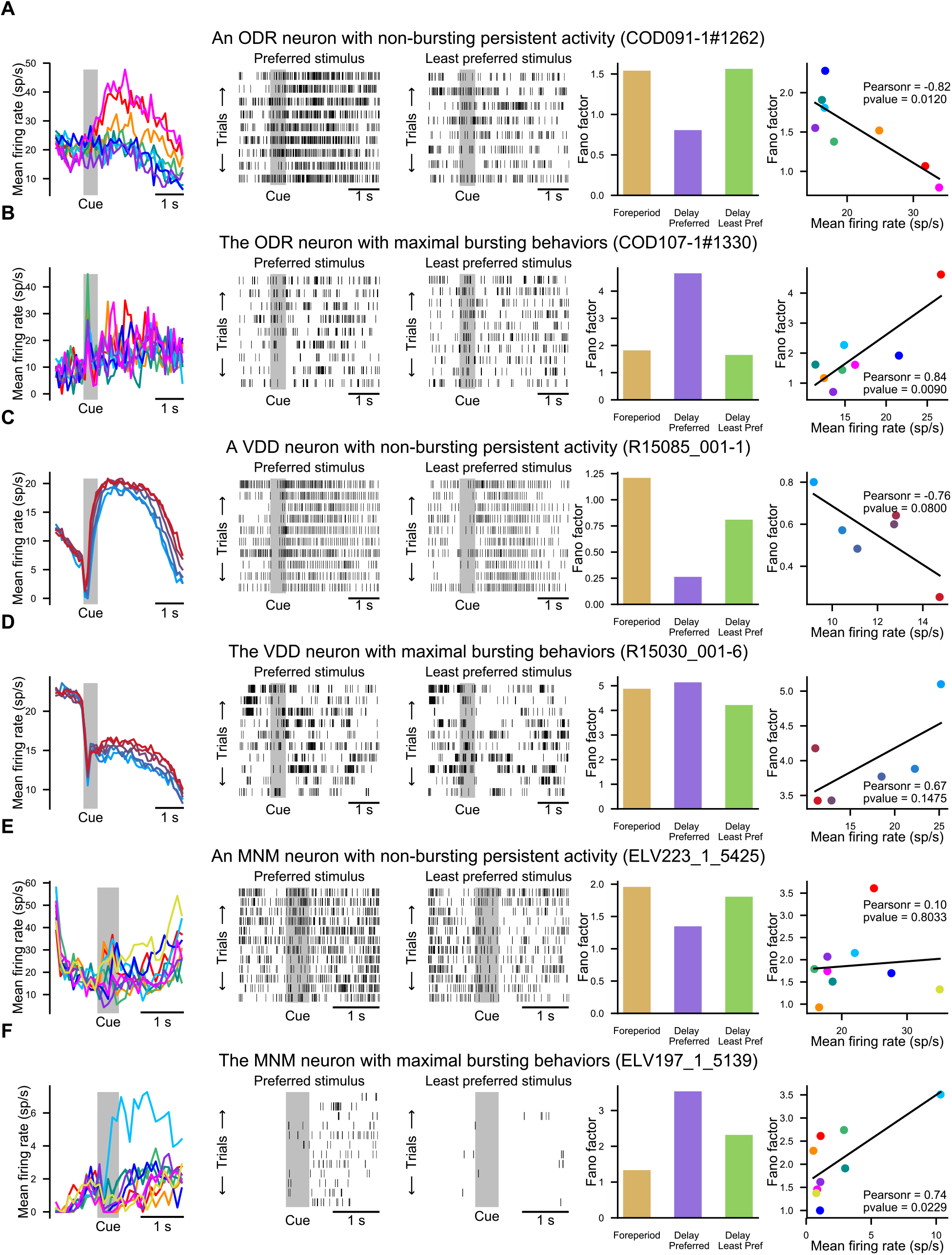
Sample neurons with their IDs in the datasets. The five panels from left to right are: (i) temporal evolutions of the mean firing rate under each stimulus condition, denoised by projecting to the first principal components capturing 80% of the variance of population activity of all well-tuned neurons in the same dataset (Murray et al., 2017); (ii) spike raster plot under the preferred stimulus; (iii) spike raster plot under the least preferred stimulus; (iv) bar plot of FF during foreperiod, under the preferred stimulus during delay, and under the least preferred stimulus during delay; and (v) scatterplot of FF vs. mean firing rate across stimulus conditions during delay. **(a)** A neuron with non-bursting stable activity in the ODR task. **(b)** The neuron with maximal bursting behaviors in the ODR task. **(c)** A neuron with non-bursting stable activity in the VDD task. **(d)** The neuron with maximal bursting behaviors in the VDD task. **(e)** A neuron with non-bursting stable activity in the MNM task. **(f)** The neuron with maximal bursting behaviors in the MNM task.

We next characterized these FF features at the population level. We plotted the histograms of FF during foreperiod, and under the preferred and the least preferred stimulus conditions during delay for each task. We found that the majority of neurons had FF lower than 3 in each one of the tasks (Fig. 7, top row), significantly lower than mathematical and circuit model predictions. We also created scatterplots of FF for preferred vs. least-preferred stimuli during delay, and the histograms of their differences (Fig. 7, bottom row). We found that in ODR and MNM datasets, but not in the VDD dataset, FF values for the preferred stimulus were larger than those for least preferred stimulus, at relatively small effect sizes with statistical significance (two-sided Wilcoxon signed-rank test, *P* = 0.003, *n* = 179, Cohen’s D = 0.24 for ODR; two-sided Wilcoxon signed-rank test, *P* = 0.10, *n* = 153, Cohen’s D = 0.19 for VDD; two-sided Wilcoxon signed-rank test, *P* = 0.002, *n* = 75, Cohen’s D = 0.35 for MNM).

**Figure 7:**
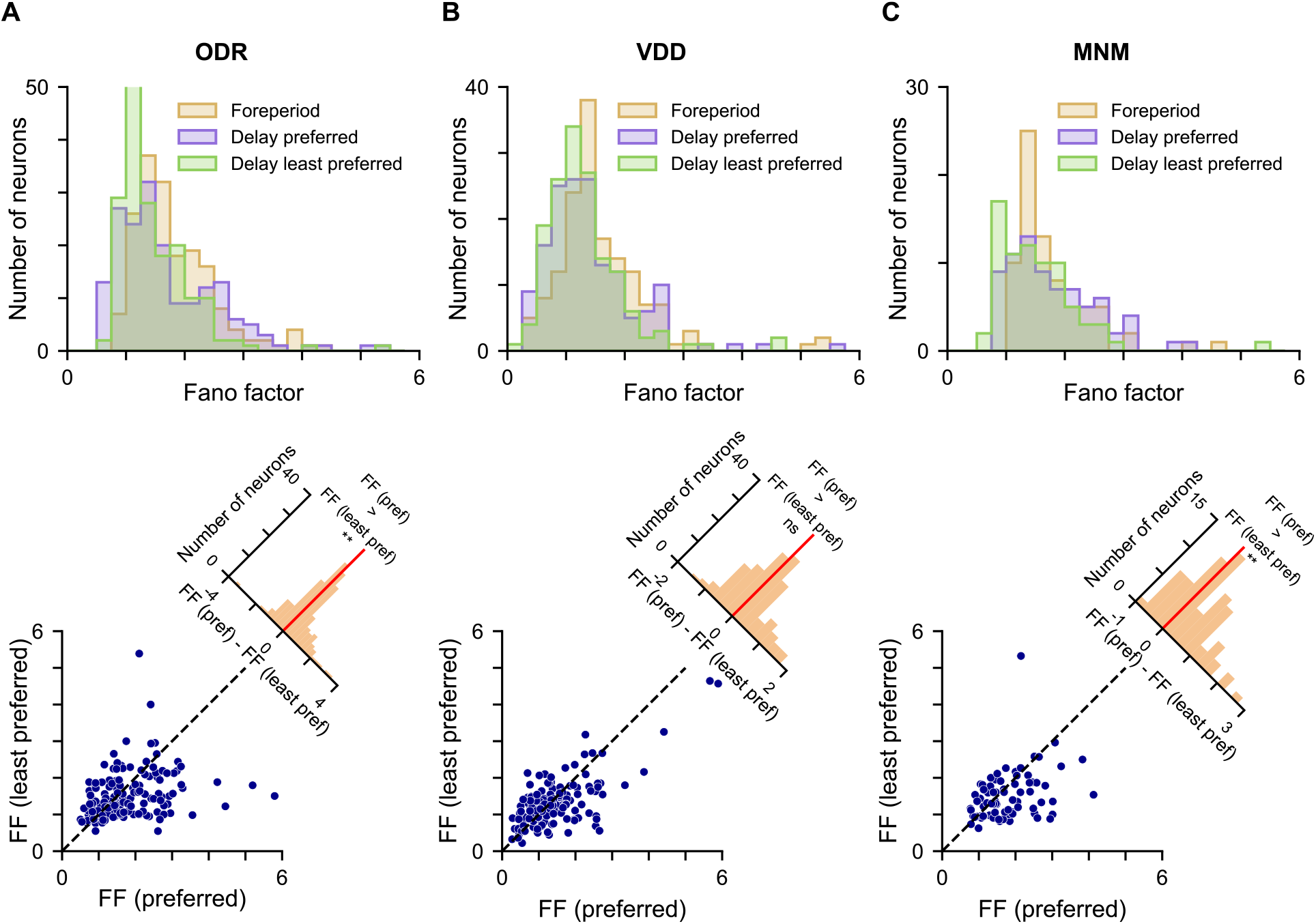
Population statistics of FF in each task. Top: Histograms of FF during the foreperiod, under the preferred stimulus during delay, and under the least preferred stimulus during delay, for all neurons with well-tuned delay activity. Bottom: Scatterplot of FF under the preferred stimulus versus under the least preferred stimulus, along with histogram of the difference in FF (preferred - the least preferred stimulus). **(a)** Results of the ODR task. FF under the preferred stimulus was slightly greater than that under the least preferred stimulus. **(b)** Results of the VDD task. FF under the preferred stimulus was not significantly different from that under the least preferred stimulus. **(c)** Results of the MNM task. FF under the preferred stimulus was greater than that under the least preferred stimulus. Significance level: *: *P* < 0.05; **: *P* < 0.01; ***: *P* < 0.001

We also characterized the distribution of the correlation between FF and mean firing rate (Corr(*r*_mean_,FF)) across stimulus of the entire population for each task. Similarly, in ODR and MNM datasets, the distributions are weakly biased to the positive side with large standard deviations, while in the VDD dataset, the distribution is centered around 0 (mean: 0.08, standard deviation: 0.26, two-sided Wilcoxon signed-rank test: *P* = 0.001 for ODR; mean: −0.04, standard deviation: 0.34, two-sided Wilcoxon signed-rank test: *P* = 0.30 for VDD; mean: 0.12, standard deviation: 0.29, two-sided Wilcoxon signed-rank test: *P* = 0.002 for MNM). This shows that in the empirical data overall there is no significant positive correlation between *r*_mean_ and FF as predicted by the burst-coding models.

As illustrated in the doubly stochastic model and the circuit models above, if WM delay activity were a result of transient bursts, there should be higher FF for the preferred stimulus during delay compared to the foreperiod baseline. We tested this through analysis of (neuron, bin) pairs during delay for each task, to control for any nonstationarity in FF across the delay. Here the preferred and least-preferred stimuli were defined based on the firing rates in each time bin, and the FF of this bin was compared to the baseline foreperiod FF. We found that in each task, FF values for both stimulus conditions during the delay were reduced compared to the foreperiod (Fig. 8a–c), at small effect sizes (ODR preferred stimulus: two-sided Wilcoxon signed-rank test, *P* = 1.4 × 10^−16^, *n* = 1151, Cohen’s D = 0.15; ODR least preferred stimulus: two-sided Wilcoxon signed-rank test, *P* = 3.8 × 10^−49^, *n* = 1053, Cohen’s D = 0.29; VDD preferred stimulus: two-sided Wilcoxon signed-rank test, *P* = 7.3 × 10^−25^, *n* = 794, Cohen’s D = 0.24; VDD least preferred stimulus: two-sided Wilcoxon signed-rank test, *P* = 2.2 × 10^−27^, *n* = 794, Cohen’s D = 0.34; MNM preferred stimulus: two-sided Wilcoxon signed-rank test, *P* = 0.004, *n* = 237, Cohen’s D = 0.07; MNM least preferred stimulus: two-sided Wilcoxon signed-rank test, *P* = 6.8 × 10^−6^, *n* = 237, Cohen’s D = 0.12). This suggests that WM delay activity is characterized by relatively high across-trial stability, which is in clear contradiction to burst-coding model predictions that are in the opposite direction.

**Figure 8:**
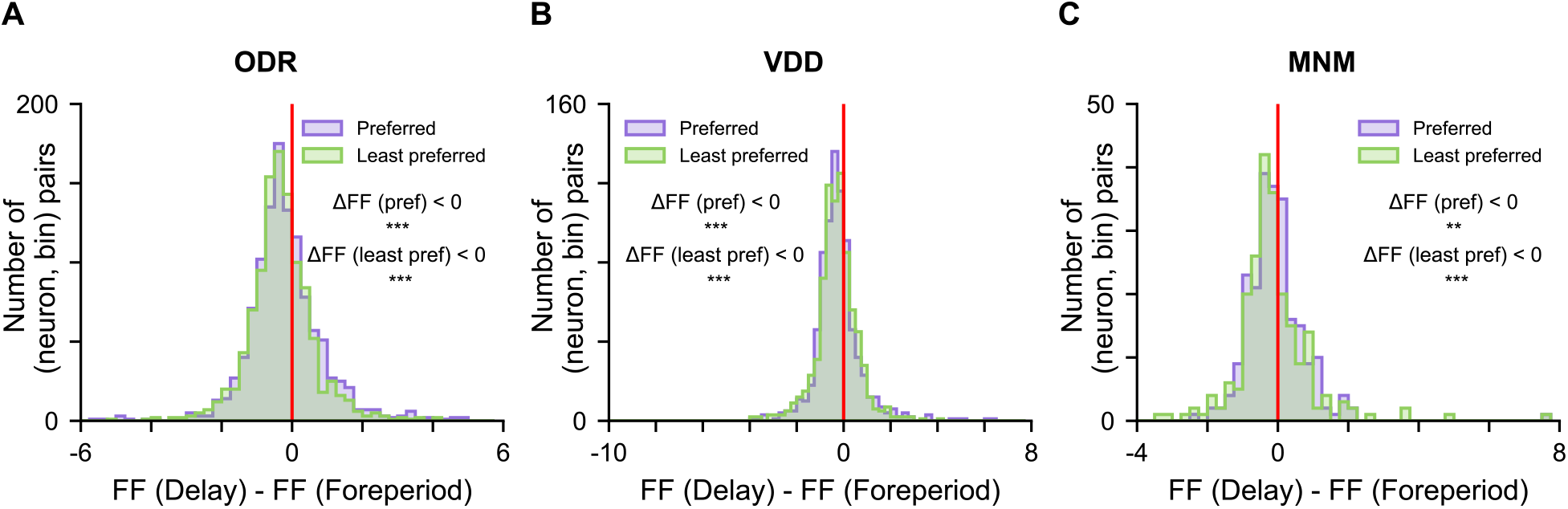
Statistics of FF of (neuron, bin) pairs in each task. Histograms of the change in FF (delay – foreperiod) under the preferred and least-preferred stimulus conditions, for all (neuron, bin) pairs with well-tuned delay activity. Empirical results contradict sharply the theoretical predictions of burst-coding models. **(a–c)** In all three datasets the change in FF was significantly less than zero for both preferred and least-preferred stimulus conditions. Significance level: *: *P* < 0.05; **: *P* < 0.01; ***: *P* < 0.001.

To summarize, we have the following findings in the three empirical spike-train datasets from WM tasks. First, the magnitudes of FF were much smaller than model predictions. Second, only a small fraction of neurons displayed increased FF for the preferred stimulus, and have positive correlation between FF and *r*_mean_ during delay. Third, FF values were overall reduced during WM delay compared to foreperiod baseline, although FF values for the preferred stimulus were reduced slightly less. Fourth, WM delay activity is characterized by stability across trials, but not necessarily stationarity across time. Reasoning by contradiction, we conclude that coding by intermittent bursts in single-neuron spiking cannot explain WM delay activity observed in PFC.

### Empirical results are better explained by the non-bursting picture

In the previous section, we observed small magnitudes of FF (mostly < 3) at the population level in each task, and very few neurons had FF increasing with the mean firing rate *r*_mean_ across stimulus conditions during delay. The doubly stochastic Poisson spiking model we built can provide insights on the interpretation of these results. The burst-coding model requires that the baseline firing should be weak, and therefore in terms of model parameters we should have *r*_*L*_ ≪ *r*_*H*_, with *r*_*L*_ small and homogeneous across stimulus conditions. It also requires that the bursts are sparse, namely 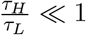. If these conditions were not met, for instance with *r*_*L*_ close to *r*_*H*_, or *r*_*L*_ tuned with stimulus, or 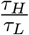 not low enough, FF would be lower at given observed mean firing rate level, according to Equation 27.

In Fig. 9a, as an illustrative example we depicted three sets of parameters all with the same *r*_mean_ ≈ 22 sp/s. Scenario (1) corresponds to the bursting regime with high FF. Scenario (2) corresponds to the a regime with a smaller difference between *r*_*L*_ and *r*_*H*_, and low FF. Scenario (3) corresponds to the a regime with burst duration twice as long as the low-state duration, and low FF. In Fig. 9b, we re-parametrized and plotted FF as a function of *r*_mean_ and the scaled difference of firing rates between the two states 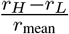. As shown mean there, the low FF (< 3) region, which is typically observed in empirical data, is characterized by a small difference between *r*_*L*_ and *r*_*H*_, and the parameter set of Scenario (2) lies in this region. This means that the default baseline state may not be as quiescent as expected, but is at a relatively high firing rate. In Fig. 9c, we re-parametrized and plotted FF as a function of *r*_mean_ and the mean dwell time ratio 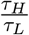. As shown there, low FF (< 3) region is characterized by very high 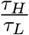, even far exceeding the sparse regime, and the parameter set of Scenario (3) lies in this region. This means that the actual burst duration is too long to be considered as a brief burst. Our mathematical formalism also provides an explanation for why most neurons had the same level of FF across stimulus conditions even though the firing rates were well-tuned. It is possible to fix FF, and solve for *r*_mean_ in terms of *r*_*L*_, 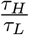, and *τ*_*H*_. As shown in Fig. 9d, we fixed FF (= 2 here, as typically observed in empirical data) and *τ*_*H*_ = 25 ms, and plotted *r*_mean_ against *r*_*L*_ and 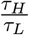. Similarly, higher *r*_mean_, which corresponds to preferred stimuli, can be obtained through stronger baseline firing *r*_*L*_ and/or longer burst durations (sacrifice of sparsity), without increasing FF.

**Figure 9:**
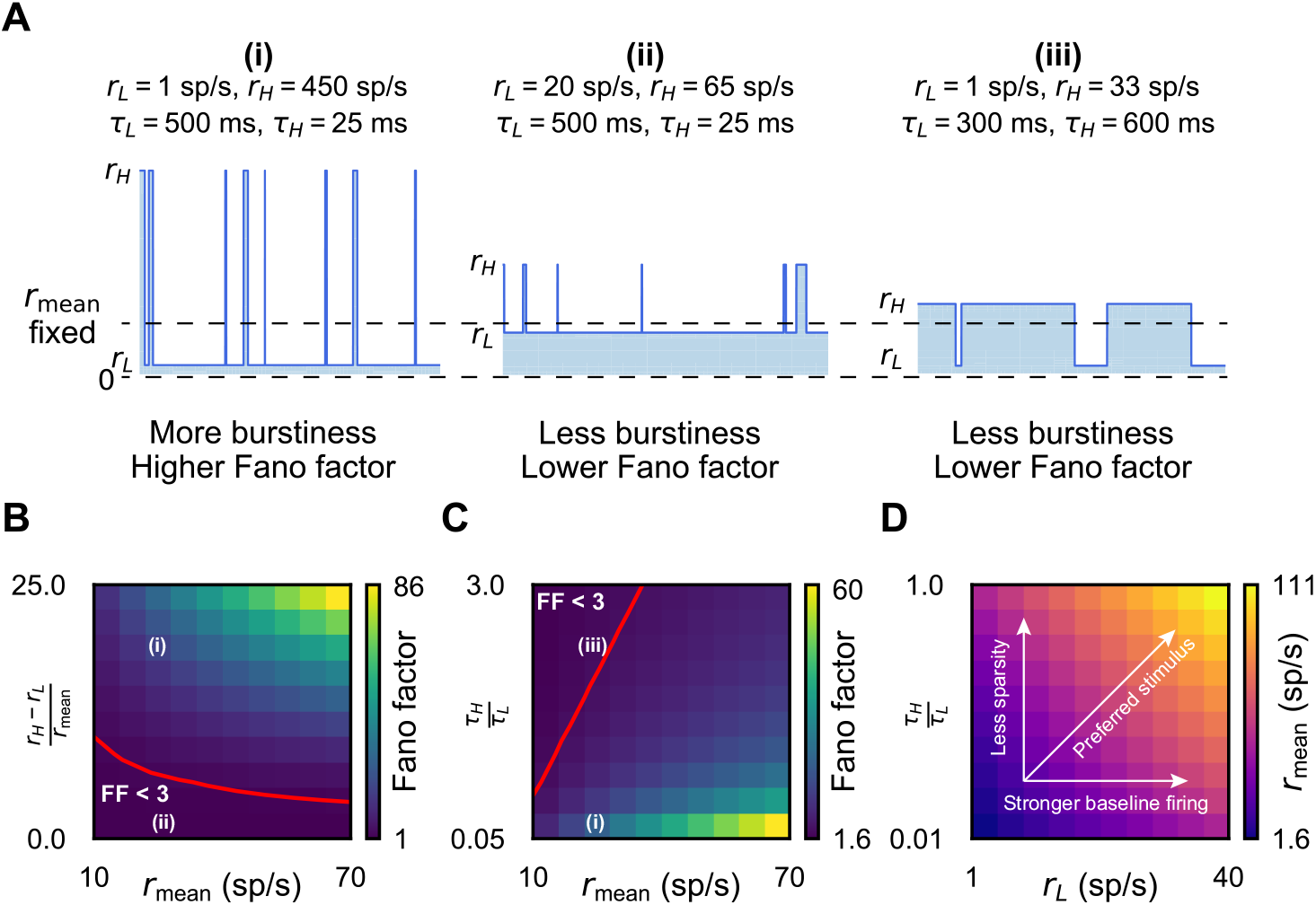
Interpretation of the empirical results in terms of the doubly stochastic model. **(a)** Given the same mean firing rate *r*_mean_ (≈ 22 sp/s in the schematics), less burstiness explains the observed low FF in empirical data (in correspondence to Fig. 1d). Scenario (1) with *r*_*L*_ = 1 sp/s, *r*_*H*_ = 450 sp/s, *τ*_*L*_ = 500 ms, and *τ*_*H*_ = 25 ms corresponds to the bursting regime with high FF. Low FF can be achieved either by smaller difference between *r*_*H*_ and *r*_*L*_ (scenario (2) with with *r*_*L*_ = 20 sp/s, *r*_*H*_ = 65 sp/s, *τ*_*L*_ = 500 ms, and *τ*_*H*_ = 25 ms), and/or higher mean dwell time ratio 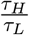 (scenario (3) with with *r*_*L*_ = 1 sp/s, *r*_*H*_ = 33 sp/s, *τ*_*L*_ = 300 ms, and *τ*_*H*_ = 600 ms). **(b)** Heatmap of FF as a function of mean firing rate and the scaled difference of firing rates in the two states 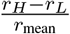, with *τ*_*L*_ = 500 ms, *τ*_*H*_ = 25 ms, and time bin width Δ = 250 ms. The red line is the contour line of FF = 3. Region where FF < 3, which was mostly observed in empirical data, is characterized by smaller difference in *r*_*H*_ and *r*_*L*_. Scenarios (1) and (2) in **(a)** are marked on the heatmap. **(c)** Heatmap of FF as a function of mean firing rate and mean dwell time ratio 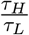, with *τ*_*L*_ = 300 ms, *r*_*L*_ = 1 sp/s, and time bin width Δ = 250 ms. The red line is the contour line of τL τ_*H*_ FF = 3. Region where FF < 3, which was mostly observed in empirical data, is characterized by high 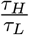. Scenarios (1) and (3) in **(a)** are marked on the heatmap. **(d)** Heatmap of *r*_mean_ as a function of *r*_*L*_ and 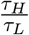 with FF = 2 fixed, which was typically observed in empirical data, and with *τ*_*H*_ = 25 ms, Δ = 250 ms. At fixed FF level, stronger mean firing rate under the preferred stimulus is achieved by elevating baseline firing and/or sacrificing spiking sparsity.

All these scenarios indicate that with empirical data, the corresponding doubly stochastic model corresponds to the baseline *L* state being higher and/or the high-rate state being much longer than proposed for burst-coding. In other words, modulation of delay activity across stimulus conditions is not well accounted for by transient bursts, and is well consistent with non-bursting theories.

## Discussion

In this study, we addressed an ongoing debate on the nature of WM delay activity between persistent activity and burst coding (Constantinidis et al., 2018; Lundqvist et al., 2018), by examining the implications of coding proposals on the stability across trials of neuronal spiking activity during WM delays. Statistical modeling using a doubly stochastic Poisson process and simulation of biophysically-based spiking circuit models demonstrated that FF is a sensitive measure that well dissociates these two divergent mechanisms, and that the burst-coding scenario would imply a significant increase in FF during the delay. In clear contrast, empirical spike-train data from PFC showed that FF is reduced and across-trial stability is strengthened during the WM delay. Very few neurons exhibited task-dependent burstiness. We conclude that the contribution of burstiness to WM delay spiking activity in PFC is strongly constrained.

### Relation to other empirical studies

We distinguish between across-trial stability and across-time stationarity as two important features to characterize WM delay activity, with across-trial stability as a key test of burst-coding models. We chose the commonly used FF as a measure for trial-to-trial variability, because there exist two distinct timescales under the burst-coding assumptions so that the time window size for analysis should be long enough relative to burst duration. There exist other measures of spiking variability, such as coefficients of variation, which are characterize the irregularity of spiking at fast timescale within single trials (Compte et al., 2003; Shinomoto et al., 2009). The transient bursts in the context of the debate we are addressing are meant to be sharp and sparse, which could be different from those defined elsewhere, such as intrinsic burstiness in neurons (Compte et al., 2003).

Observed reduction in FF at the population level during WM delays is consistent with other studies on single-neuron spiking. FF has been shown to be reduced after stimulus onset in multiple cortical regions (Churchland et al., 2010). Nawrot et al. (2008) applied a statistical model based on renewal processes and observed decreases in FF in monkey motor cortex during task engagement. In line with our findings, Chang et al. (2012) found that frontal eye field neurons had much broader spatial tuning of FF which was dissociated from behavioral engagement and distinct from firing rates. These properties can indeed be task-specific, however, such as the weak tuning of FF exhibited in the MNM dataset.

### Reevaluation of burst-coding proposals

Our analysis of the burst-coding scenario was based on proposals from empirical studies (Lundqvist et al., 2016b, 2018), as well as based on theoretical models for WM (Mongillo et al., 2008; Lundqvist et al., 2011; Mi et al., 2017). Our study built a testing framework for this, and can provide important insights into the various experimental results.

Before characterizing the FF measure, we showed that burst-coding proposals can require the firing rate within burst states to be very high, even potentially exceeding biophysical limits. The doubly stochastic Poisson model indicates that the observed small and reduced FF during WM delay conform to the non-bursting persistent activity scenario. Our conclusion that intermittent transient bursts are not plausible mechanism for the observed WM delay activity was established through reasoning by contradiction. However, the converse statement is not true: high FF does not necessarily imply bursting, as high FF can result from non-bursty over-dispersion of inter-spike intervals or from drifting excitability across trials.

Our findings are not inconsistent with previously observed intermittent bouts of elevated narrowband LFP power (Lundqvist et al., 2016a, 2018). LFP signals do not reflect individual spike contributions or single-neuron dynamics (Kajikawa and Schroeder, 2011). Therefore, LFP bursting is not equivalent to bursting of neuronal spiking, and is not incompatible with persistent activity. We also note that in the study of Lundqvist et al. (2016a), LFP bursts were mostly observed during the stimulus presentation, and much less so during the WM delay. It is thus unclear whether this LFP feature is directly related to WM maintenance, and whether those observations constitute evidence for burst-coding theoretical models.

### Implications for circuit models of working memory

We simulated and compared three biophysically detailed circuit mechanisms for generating WM delay activity. Attractor networks with recurrent connectivity have long been influential models of WM maintenance (Wang, 2001; Chaudhuri and Fiete, 2016). Wimmer et al. (2014) demonstrated that a diffusing bump attractor model captures key aspects of neuronal variability and correlations in spatial WM maintenance during the ODR task. There remain gaps between theoretical models and empirical recordings of neuronal spiking activity during WM. Empirical neuronal spiking data exhibits strong temporal variations in firing rates across delays, which fails to be captured by models with fixed-point attractors (Druckmann and Chklovskii, 2012; Barak et al., 2013; Murray et al., 2017; Cueva et al., 2020). Furthermore, attractor network models do not necessarily capture levels of spike-train irregularity observed empirically (Barbieri and Brunel, 2008; Renart et al., 2007; Roudi and Latham, 2007).

We showed that a fixed-point attractor network can reproduce reduced trial-to-trial variability during delay states, but with too low FF. Further modeling studies found that this issue could be ameliorated in a regime of excitation-inhibition balance (Lim and Goldman, 2013, 2014; Hansel and Mato, 2013; Renart et al., 2007). We verified this proposal by measuring FF in simulation of the model of Lim and Goldman (2013). At moderate firing rates, the model FF distribution resembled empirical data in magnitude, and reproduced the observed shrinkage of FF during delay compared to foreperiod. Burst-coding models, by contrast, operate in a different dynamic regime characterized by periodically occurring sharp bursts of spiking separated by longer periods of quiescence (Mongillo et al., 2008; Lundqvist et al., 2011; Fiebig and Lansner, 2017; Mi et al., 2017). We showed that a burst-coding model produces low across-trial stability with high FF, and found that empirical data better conforms to the across-trial stability of models with persistent activity.

It is also an important future direction to directly incorporate further neurobiological data into WM circuit models. For instance, the chaotic attractor network model by Pereira and Brunel (2018) was built with a learning rule inferred from data recorded in the inferior temporal cortex, which generated time-varying highly irregular activity within a WM attractor state. However, this model displayed strong firingrate fluctuations and likely exhibit incompatibly high FF values. Our study illustrates the utility of across-trial stability in empirical datasets to constrain and test WM circuit models.

### Limitations and future directions

One limitation of the datasets used here is that the number of trials for one stimulus condition is on the order of ten, which can yield large variance in FF estimation. Future experiments can collect larger number of trials of single-neuron recordings in WM tasks with a long enough delay period. However, our main aim was to characterize the qualitative relationships between FF and bursting, mean firing rate, stimulus conditions, and WM delay activity, which was not dependent on high precision in estimation of FF. Furthermore, we simulated same number of trials in the circuit models, which proved to be sufficient for this purpose. Therefore, this limitation should not affect the main conclusions of this study. Another limitation is that FF can be sensitive to drift and non-stationarity of overall firing rates across trials. Importantly, this issue would only inflate FF but not reduce it, and therefore our observations of small and reduced FF values are robust to this.

Although we demonstrated lack of specific support for the burst-coding proposal, trial-averaging in typical PSTH analyses could obscure more complex structures of spiking within single trials. PFC spike trains can exhibit high levels of irregularity during WM tasks (Compte et al., 2003; Shinomoto et al., 2009). However, spiking irregularity is not equivalent to across-trial sparseness as implied by intermittent bursting. It remains an open question how spike-count overdispersion relates to WM computations. Churchland et al. (2011) decomposed spike counts into two components and showed that across-trial variability distinguishes neural mechanisms in decision-making. More general statistical models have been proposed to explain spike-count overdispersion (Goris et al., 2014; Charles et al., 2018). Future analyses on richer datasets can further investigate relationships between spiking variability and cognitive function.

We focused on characterizing neurons with well-tuned firing rates during delay. In the scenario of population code, however, instead of relying on a small number of neurons with well-tuned activity to the stimulus, information can be distributed across neural groups with diverse tuning properties (Leavitt et al., 2017). Simultaneous recordings can inform the nature of shared neuronal variability (Cohen and Kohn, 2011), and potentially reveal complex neuronal interactions in spiking timing during cognitive functions (Crowe et al., 2010). The structure of neuronal correlations, and their modulation by task conditions, can constrain and inform models of cortical circuit dynamics and computation (Huang et al., 2019). Future studies can investigate neuronal correlations of simultaneous recordings during WM tasks (Leavitt et al., 2017), in relation to coding subspaces (Murray et al., 2017), which may provide crucial insights into the relationship between coordinated spiking dynamics and neural representations of cognitive states.

## Acknowledgements

We thank Norman Lam for useful discussions. This work was supported by National Institutes of Health grant R01 MH112746 (JDM), and by the National Eye Institute of the National Institutes of Health under award number R01 EY017077 (CC).

