## Supplementary Figures for "Trial-to-trial variability of spiking delay activity in prefrontal cortex constrains burst-coding models of working memory"

We include here plots of all the neurons with stimulus-selective delay activity in all the three tasks (179 for ODR, 153 for VDD, and 75 for MNM), similar to Figure 6 in the main text. Within each task, neurons are grouped according to Figure 5 in the main text by: those having either increased FF under the preferred stimulus during delay or having FF greater than 3 under the preferred stimulus or having positive correlation between mean firing rate and FF (group 1 below), and otherwise (group 2 below).

#### ODR NEURONS

##### Group 1

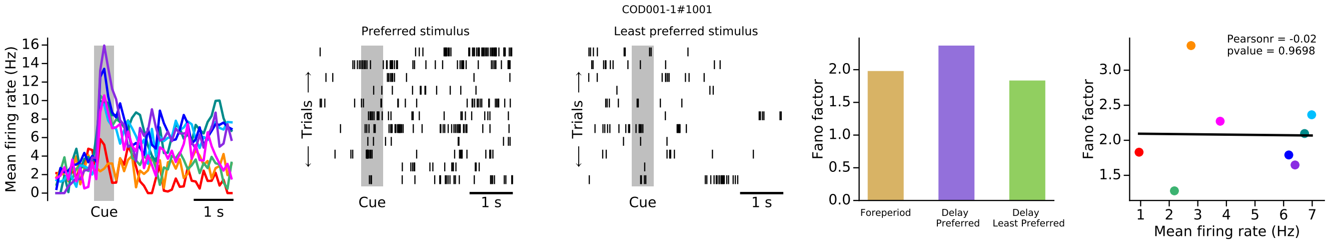


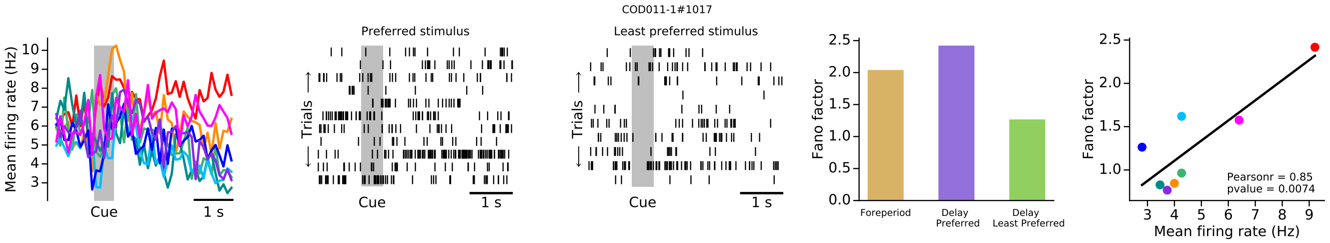


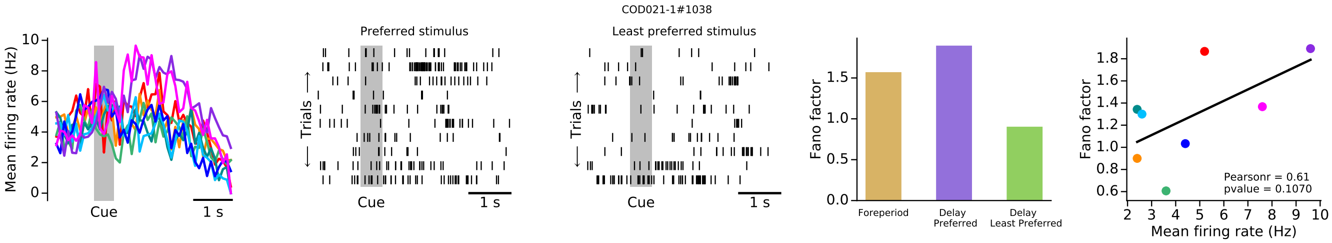


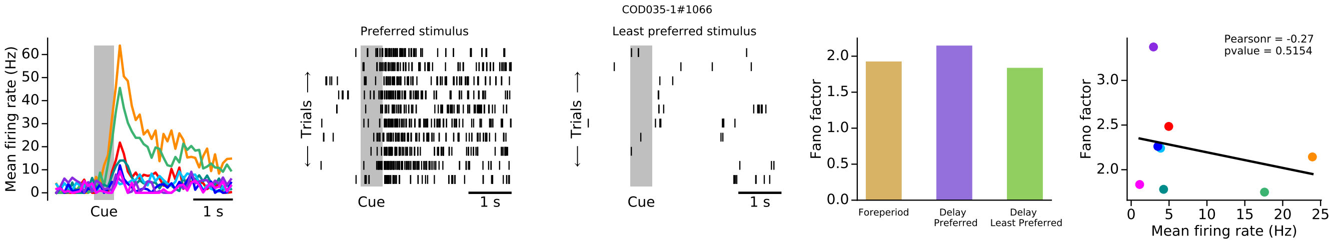

COD038-1#1073

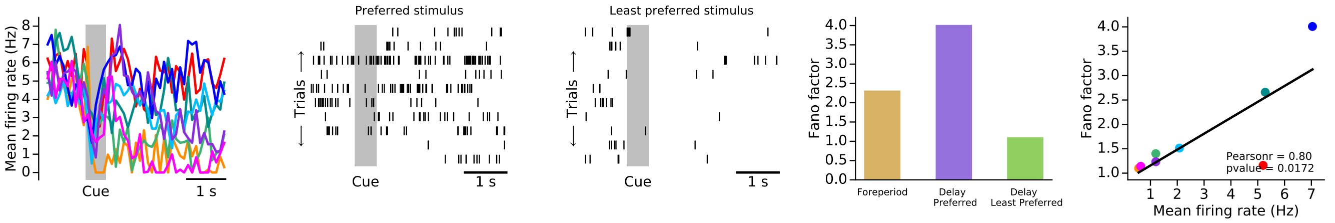


COD042-1#1085

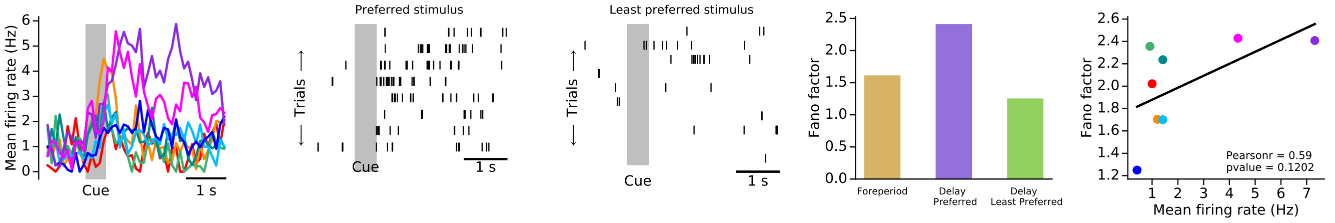


COD043-1#1087

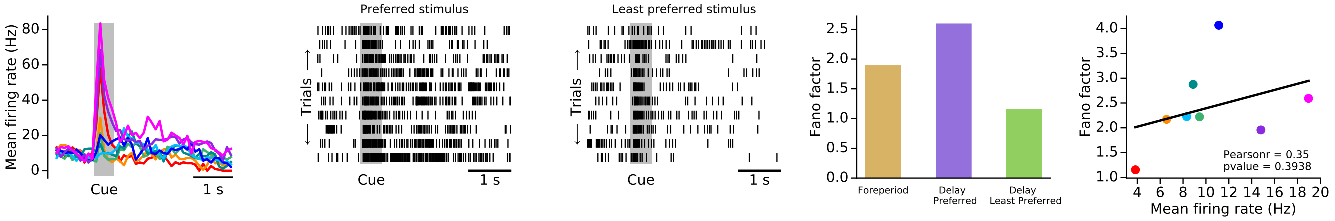


COD045-1#1091

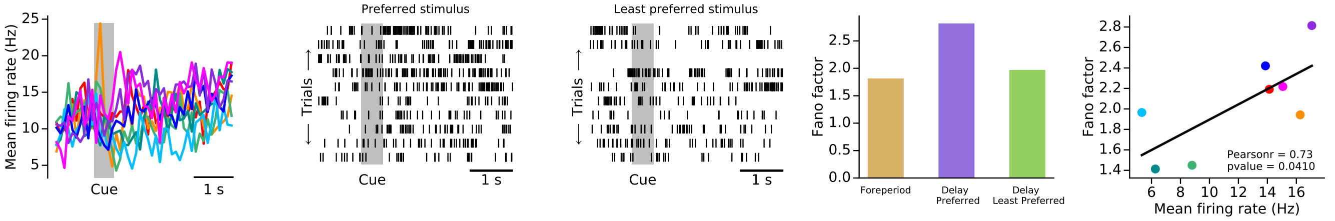


COD048-1#1100

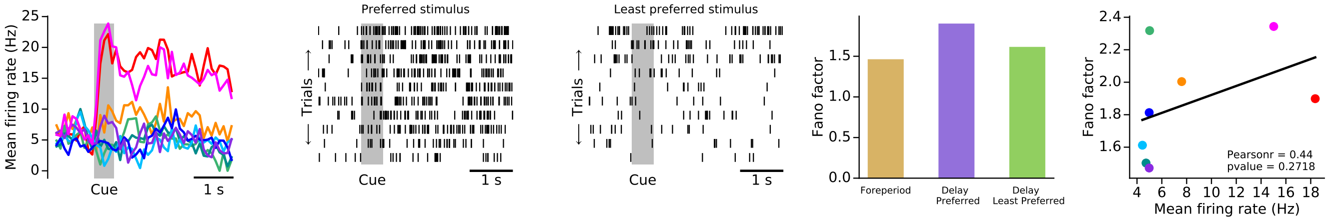

COD052-1#1114

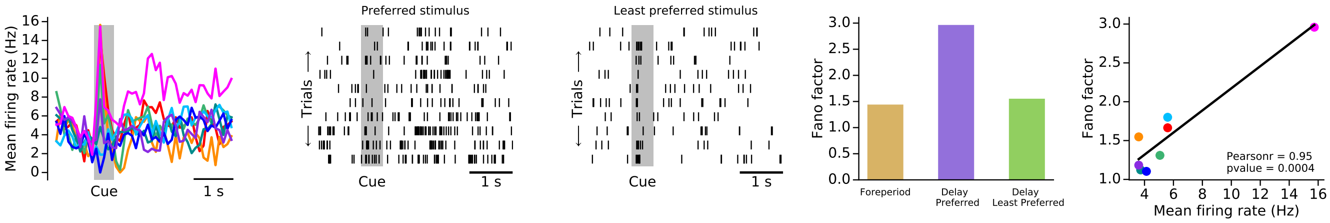


COD053-1#1115

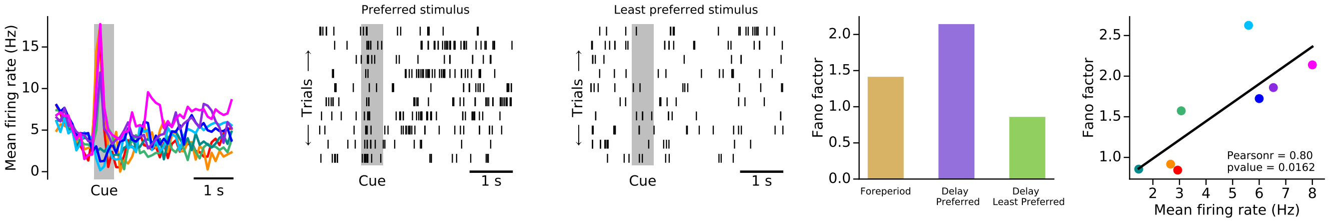


COD057-1#1130

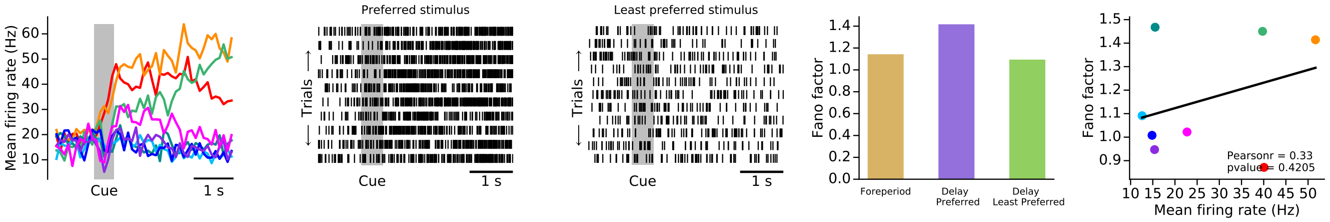


COD061-1#1143

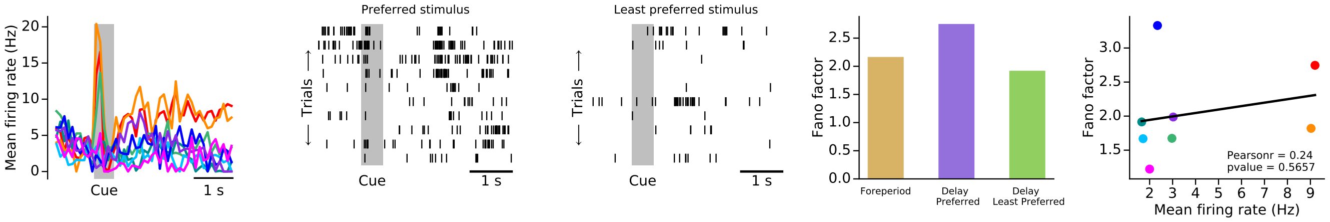


COD066-1#1157

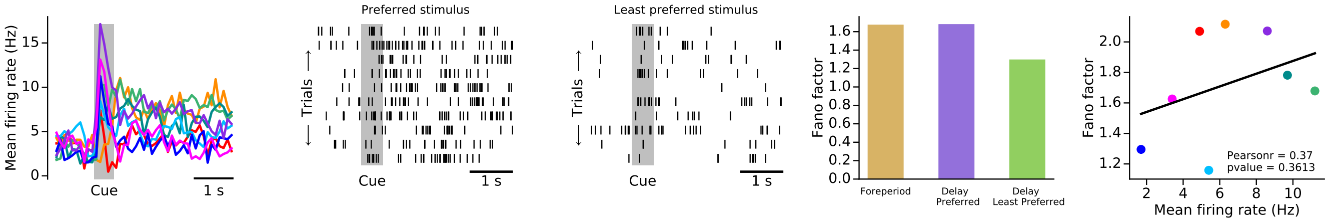

COD069-1#1168

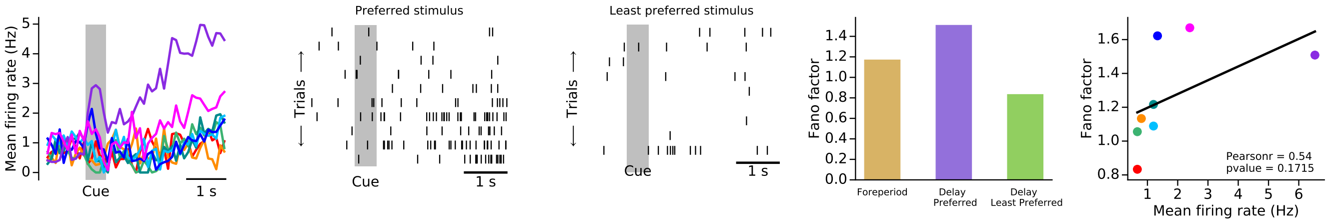


COD070-1#1174

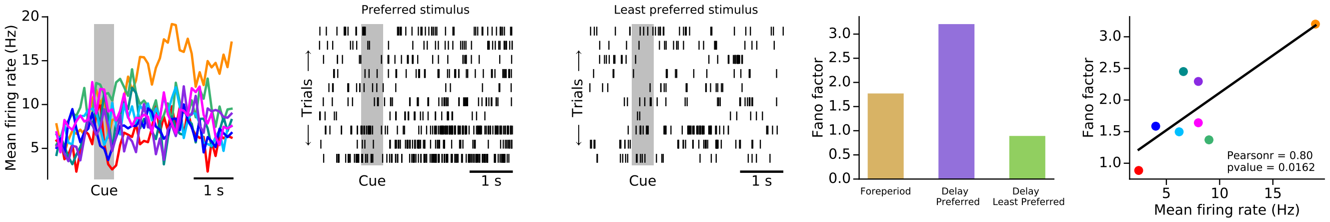


COD073-1#1182

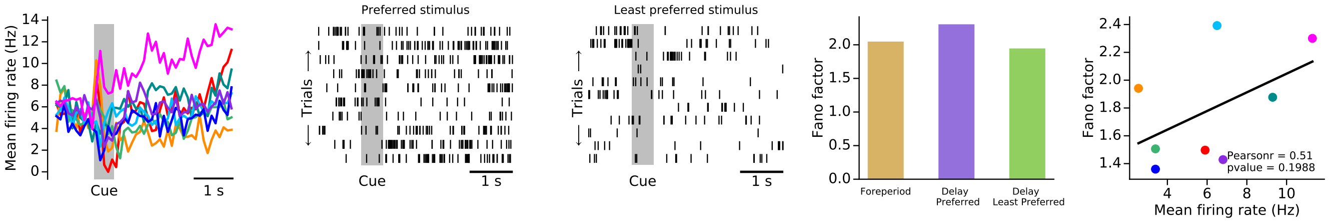


COD076-1#1188

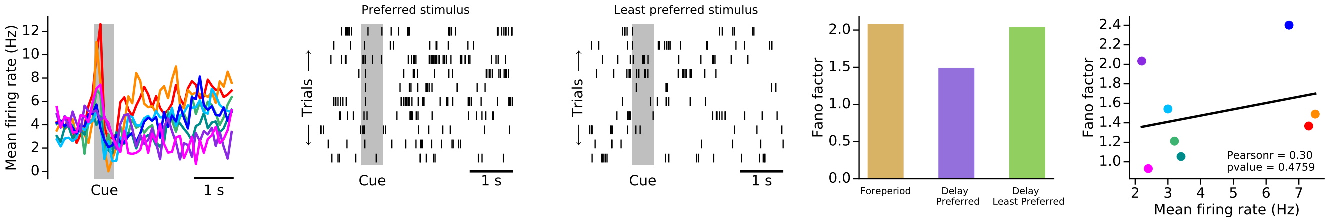


COD077-1#1193

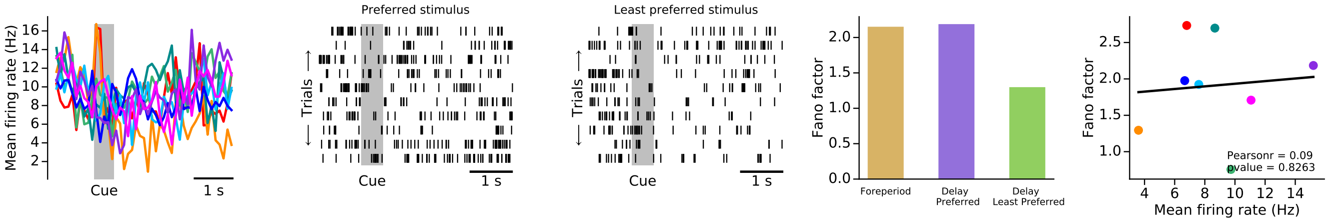

COD078-1#1195

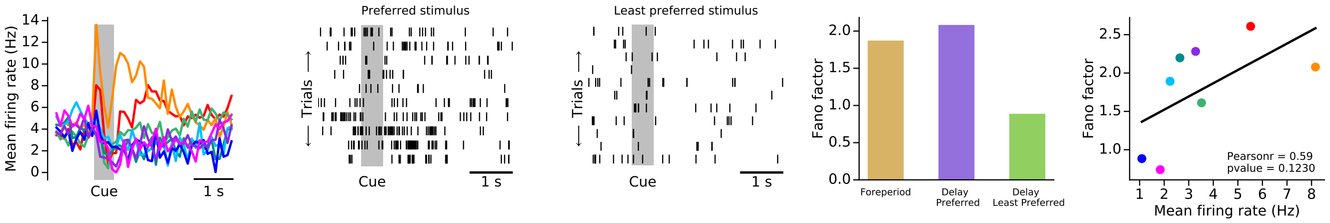


COD085-2#1234

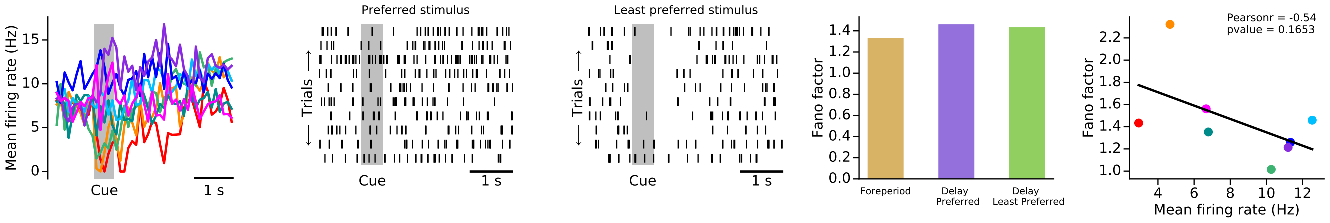


COD085-2#1237

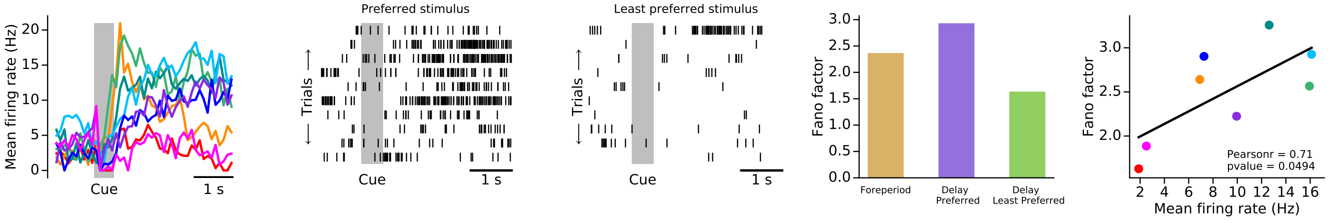


COD087-1#1245

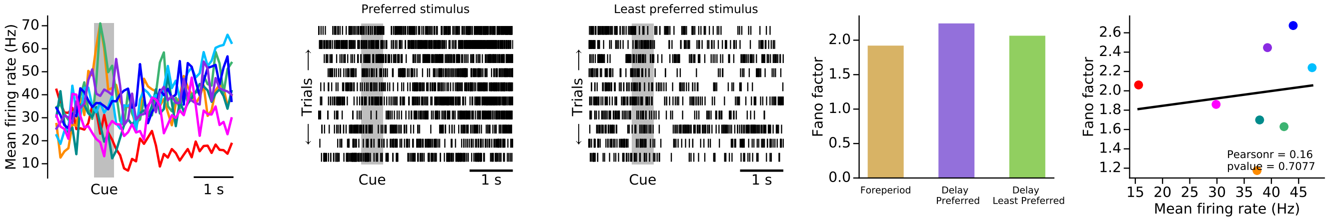


COD089-1#1252

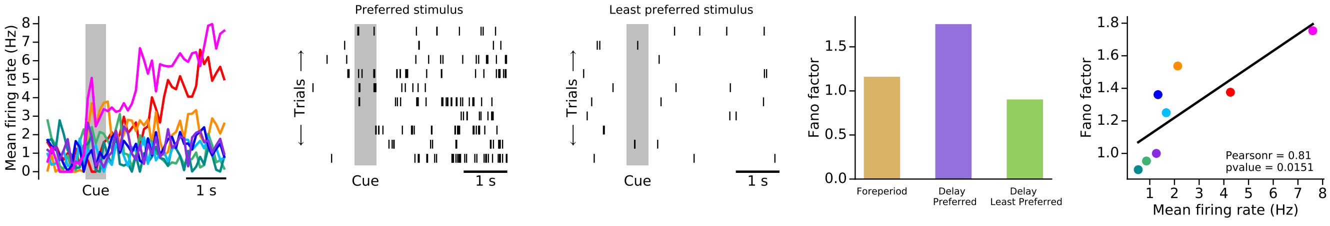

COD093-1#1267

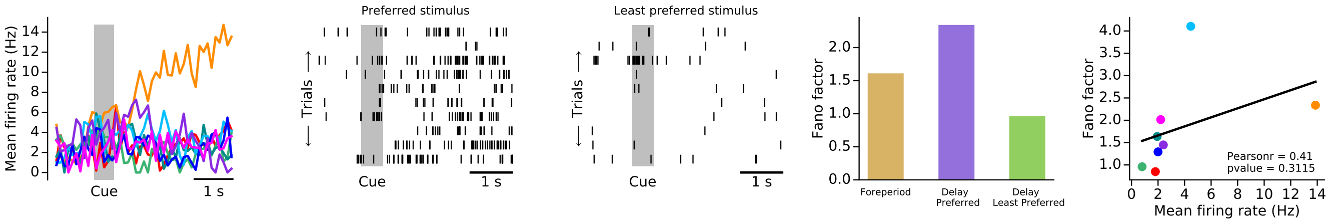


COD096-1#1274

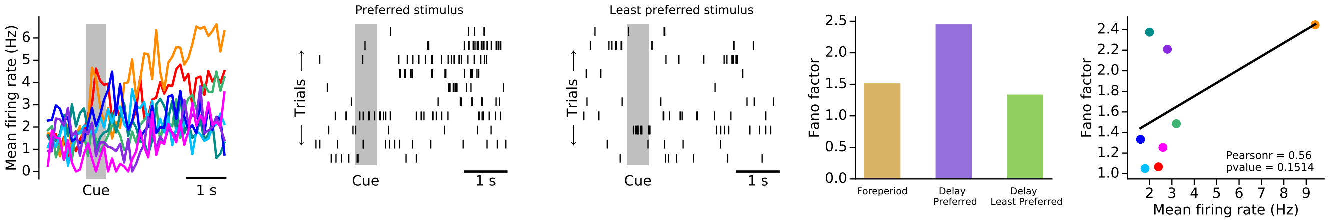


COD099-1#1291

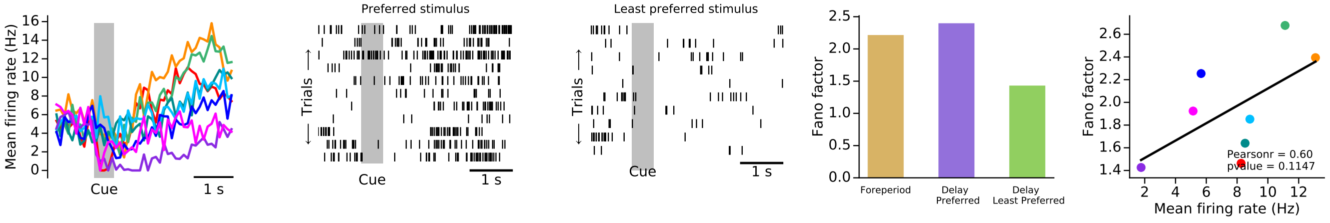


COD101-1#1299

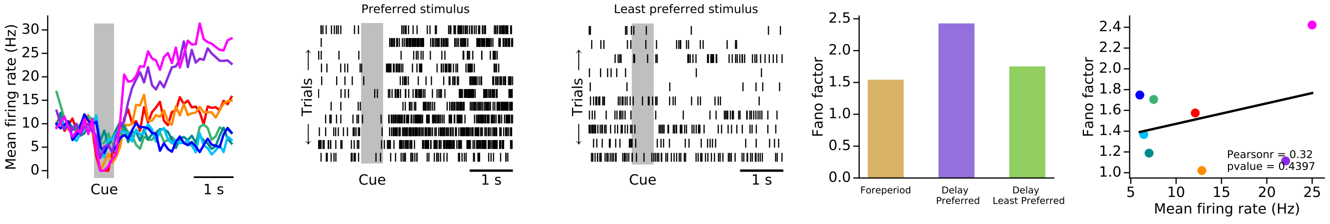


COD101-1#1300

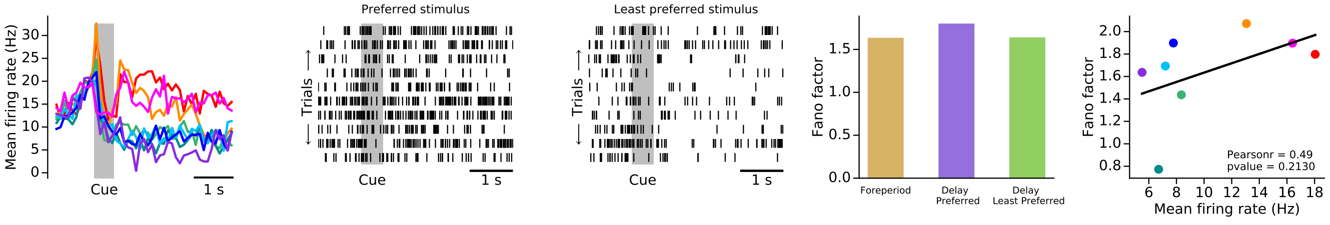

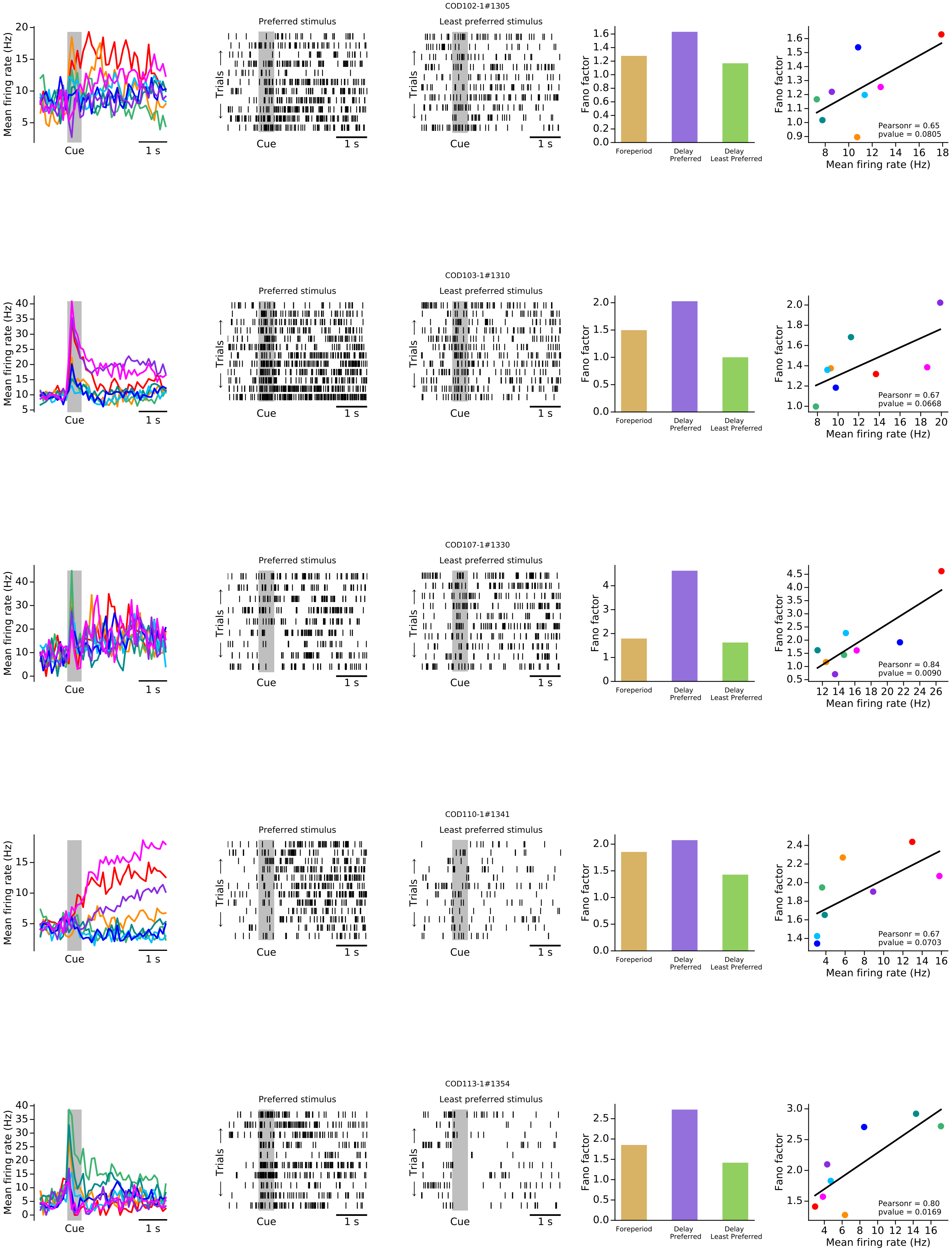
COD102-1#1305
Preferred stimulus
Least preferred stimulus
Mean firing rate (Hz)
Trials
Cue
1 s
Fano factor
Foreperiod
Delay Preferred
Delay Least Preferred
Mean firing rate (Hz)
Pearsonr = 0.65
pvalue = 0.0805
COD103-1#1310
Preferred stimulus
Least preferred stimulus
Pearsonr = 0.67
pvalue = 0.0668
COD107-1#1330
Preferred stimulus
Least preferred stimulus
Pearsonr = 0.84
pvalue = 0.0090
COD110-1#1341
Preferred stimulus
Least preferred stimulus
Pearsonr = 0.67
pvalue = 0.0703
COD113-1#1354
Preferred stimulus
Least preferred stimulus
Pearsonr = 0.80
pvalue = 0.0169

COD114-1#1358

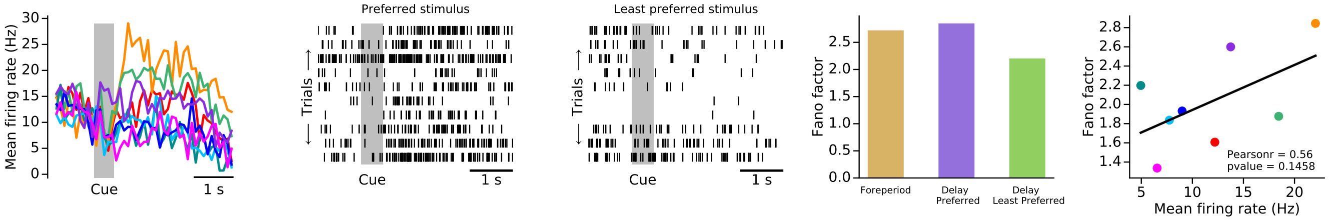


COD119-1#1382

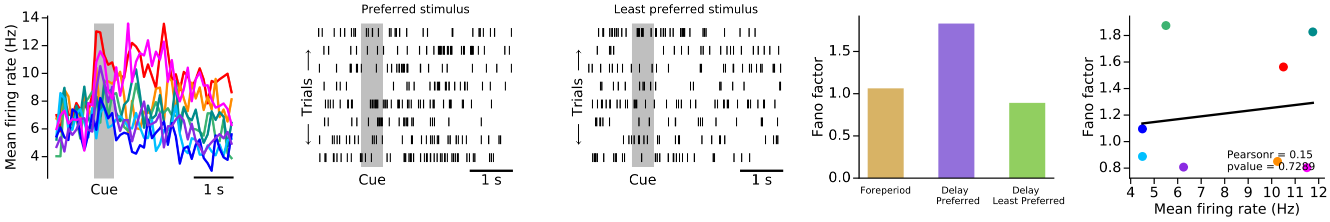


COD122-1#1395

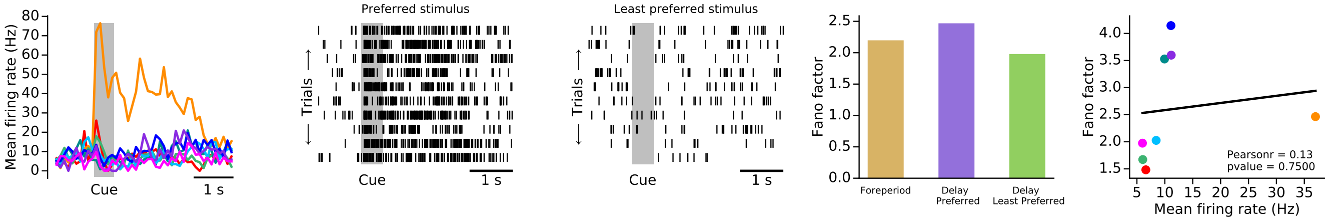


COD124-1#1406

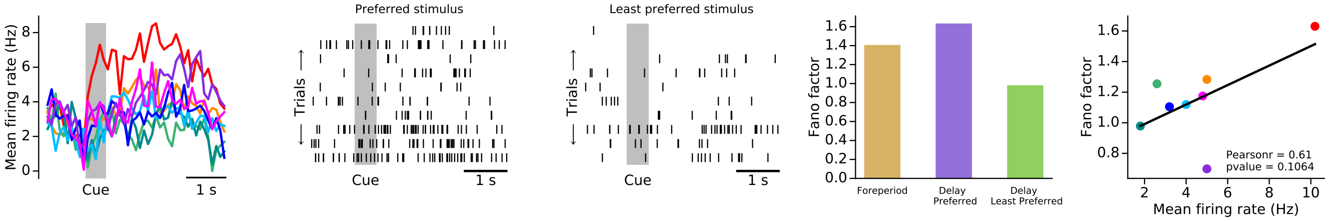


COD125-1#1409

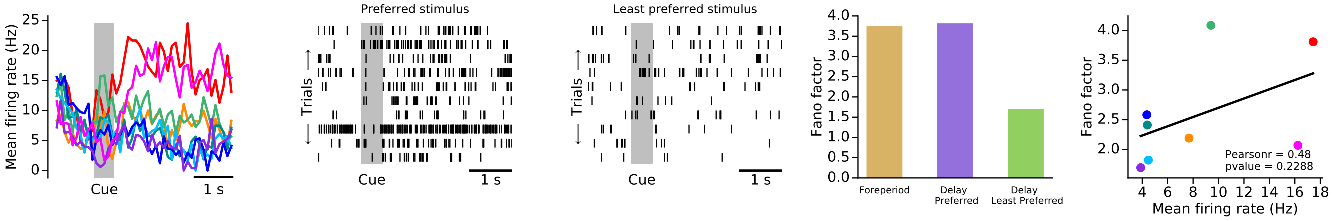

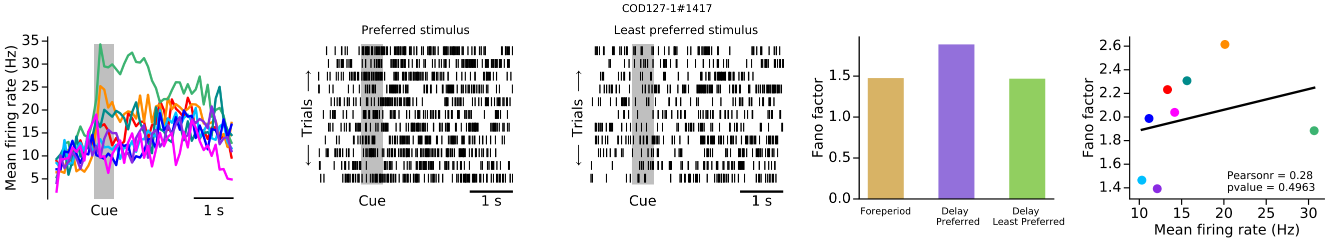
COD127-1#1417
Mean firing rate (Hz)
Cue
1 s
Preferred stimulus
Least preferred stimulus
Trials
Fano factor
Foreperiod
Delay Preferred
Delay Least Preferred
Pearsonr = 0.28
pvalue = 0.4963
Mean firing rate (Hz)

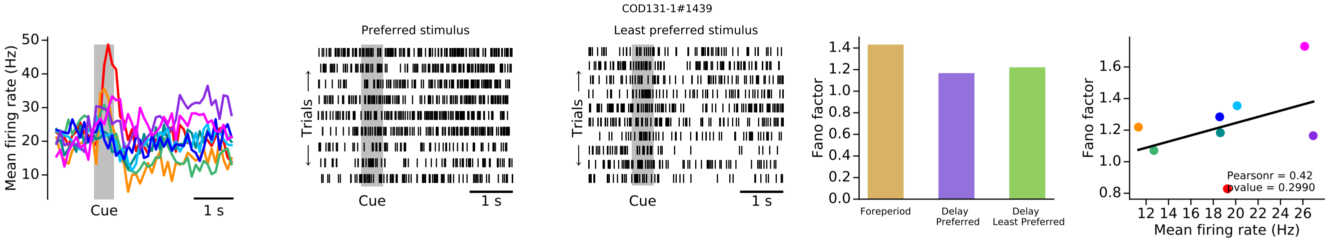
COD131-1#1439
Mean firing rate (Hz)
Cue
1 s
Preferred stimulus
Least preferred stimulus
Trials
Fano factor
Foreperiod
Delay Preferred
Delay Least Preferred
Pearsonr = 0.42
pvalue = 0.2990
Mean firing rate (Hz)

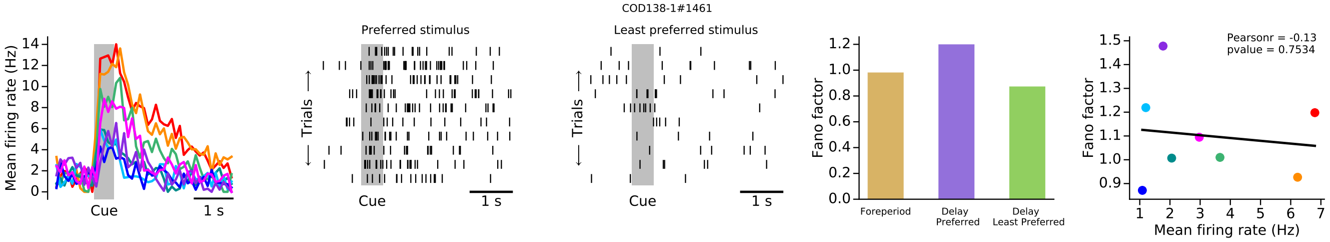
COD138-1#1461
Mean firing rate (Hz)
Cue
1 s
Preferred stimulus
Least preferred stimulus
Trials
Fano factor
Foreperiod
Delay Preferred
Delay Least Preferred
Pearsonr = -0.13
pvalue = 0.7534
Mean firing rate (Hz)

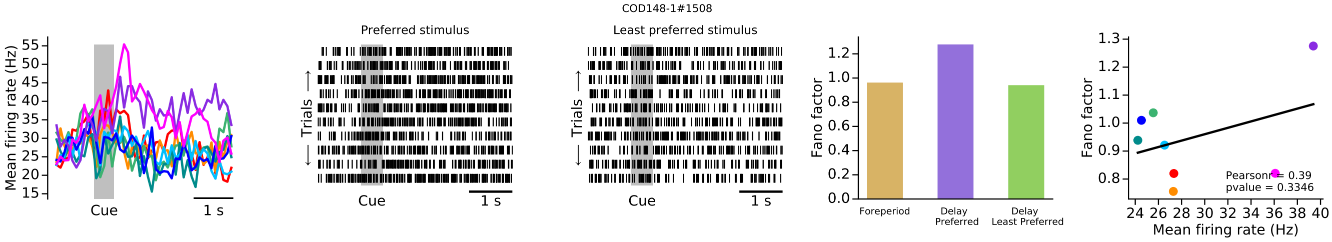
COD148-1#1508
Mean firing rate (Hz)
Cue
1 s
Preferred stimulus
Least preferred stimulus
Trials
Fano factor
Foreperiod
Delay Preferred
Delay Least Preferred
Pearsonr = 0.39
pvalue = 0.3346
Mean firing rate (Hz)

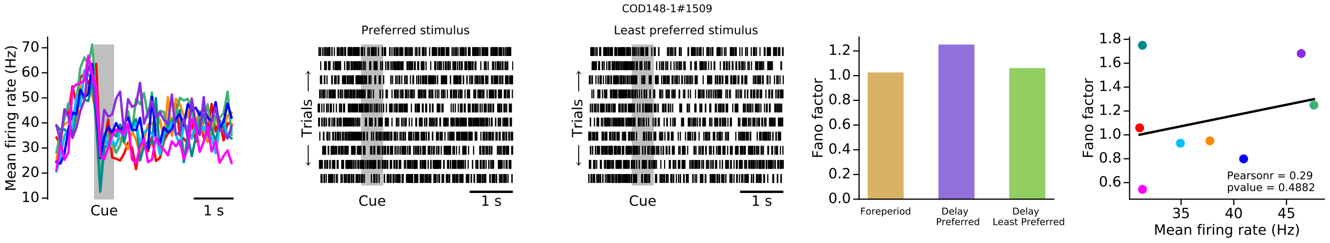
COD148-1#1509
Mean firing rate (Hz)
Cue
1 s
Preferred stimulus
Least preferred stimulus
Trials
Fano factor
Foreperiod
Delay Preferred
Delay Least Preferred
Pearsonr = 0.29
pvalue = 0.4882
Mean firing rate (Hz)

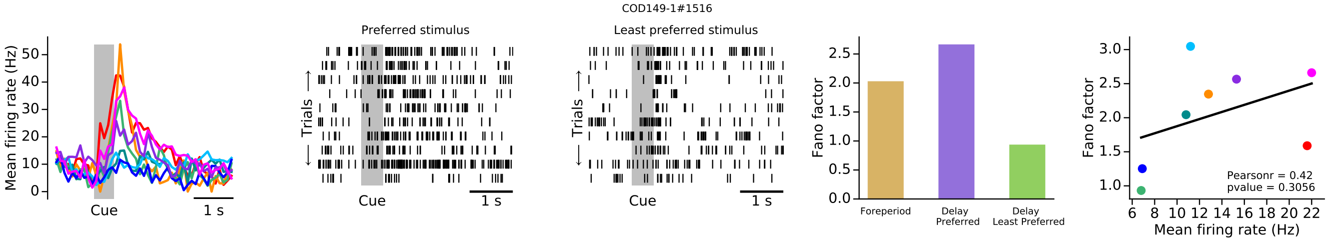
COD149-1#1516
Preferred stimulus
Least preferred stimulus
Mean firing rate (Hz)
Cue
1 s
Trials
Fano factor
Foreperiod
Delay Preferred
Delay Least Preferred
Pearsonr = 0.42
pvalue = 0.3056
Mean firing rate (Hz)

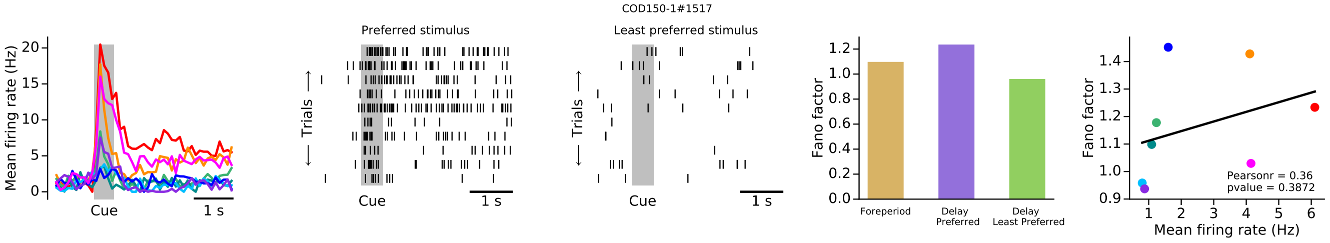
COD150-1#1517
Preferred stimulus
Least preferred stimulus
Mean firing rate (Hz)
Cue
1 s
Trials
Fano factor
Foreperiod
Delay Preferred
Delay Least Preferred
Pearsonr = 0.36
pvalue = 0.3872
Mean firing rate (Hz)

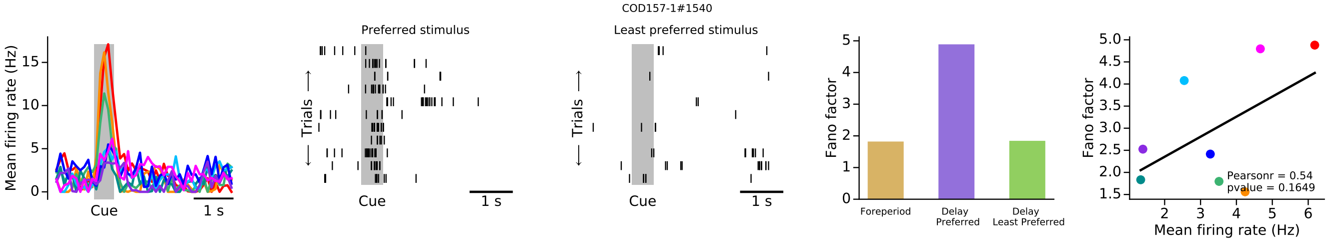
COD157-1#1540
Preferred stimulus
Least preferred stimulus
Mean firing rate (Hz)
Cue
1 s
Trials
Fano factor
Foreperiod
Delay Preferred
Delay Least Preferred
Pearsonr = 0.54
pvalue = 0.1649
Mean firing rate (Hz)

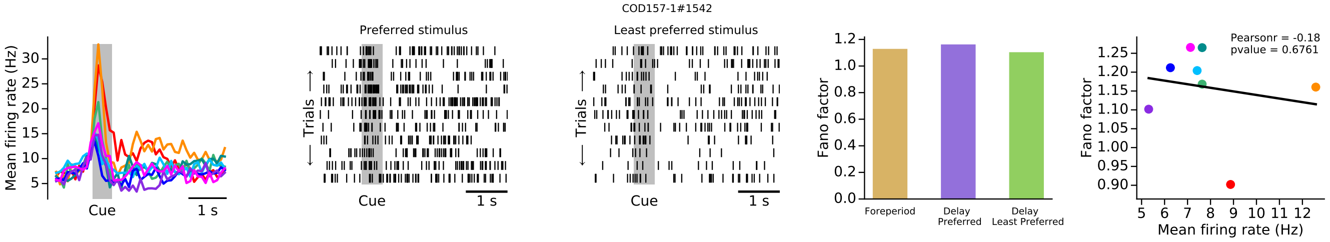
COD157-1#1542
Preferred stimulus
Least preferred stimulus
Mean firing rate (Hz)
Cue
1 s
Trials
Fano factor
Foreperiod
Delay Preferred
Delay Least Preferred
Pearsonr = -0.18
pvalue = 0.6761
Mean firing rate (Hz)

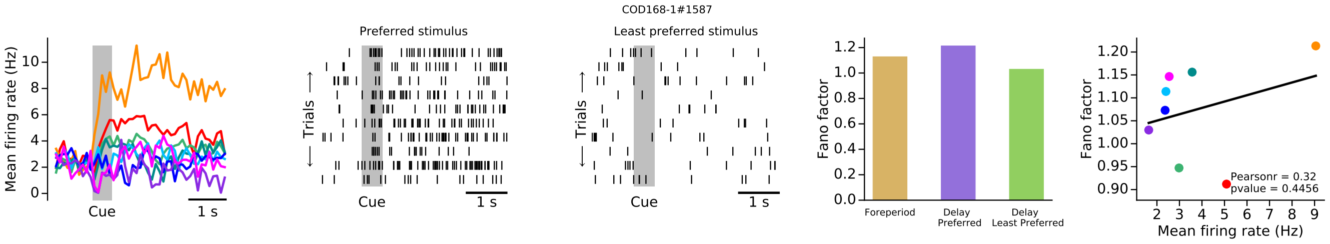
COD168-1#1587
Preferred stimulus
Least preferred stimulus
Mean firing rate (Hz)
Cue
1 s
Trials
Fano factor
Foreperiod
Delay Preferred
Delay Least Preferred
Pearsonr = 0.32
pvalue = 0.4456
Mean firing rate (Hz)

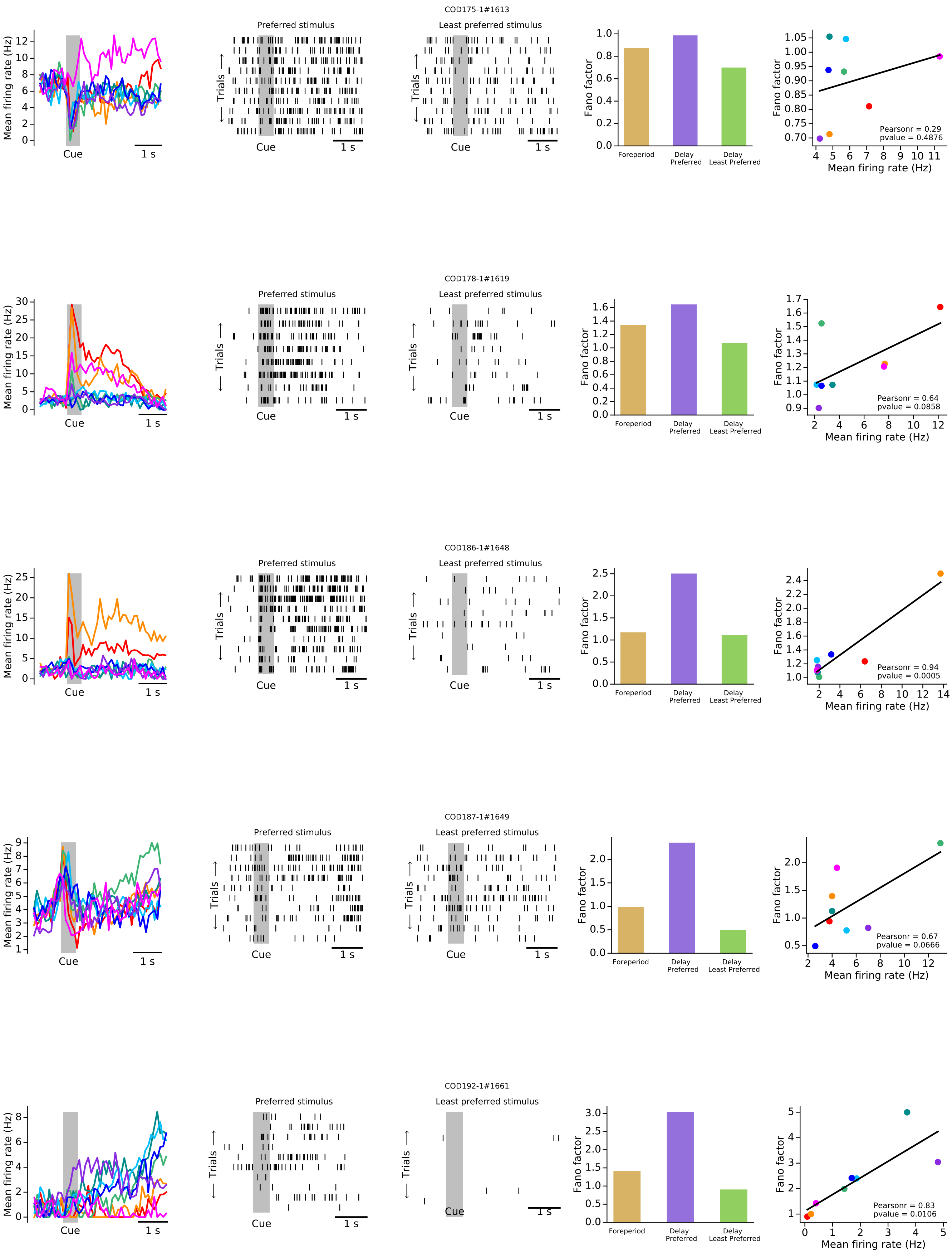
COD175-1#1613
Preferred stimulus
Least preferred stimulus
Mean firing rate (Hz)
Trials
Cue
1 s
Fano factor
Foreperiod
Delay Preferred
Delay Least Preferred
Pearsonr = 0.29
pvalue = 0.4876
Mean firing rate (Hz)
COD178-1#1619
Pearsonr = 0.64
pvalue = 0.0858
COD186-1#1648
Pearsonr = 0.94
pvalue = 0.0005
COD187-1#1649
Pearsonr = 0.67
pvalue = 0.0666
COD192-1#1661
Pearsonr = 0.83
pvalue = 0.0106

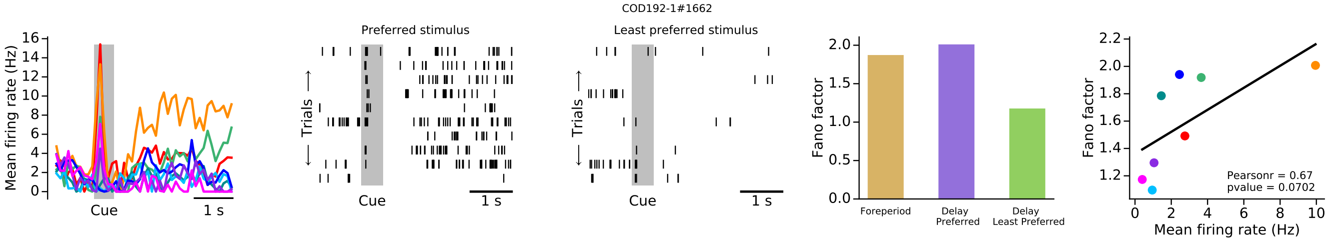
COD192-1#1662
Preferred stimulus
Least preferred stimulus
Mean firing rate (Hz)
Trials
Cue
1 s
Fano factor
Foreperiod
Delay Preferred
Delay Least Preferred
Mean firing rate (Hz)
Pearsonr = 0.67
pvalue = 0.0702

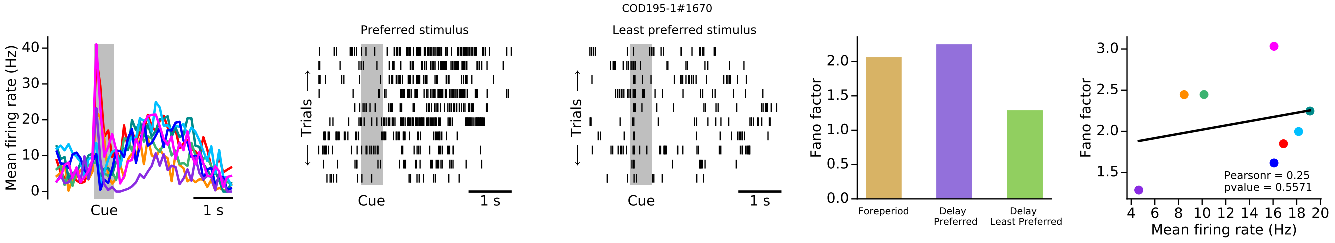
COD195-1#1670
Preferred stimulus
Least preferred stimulus
Mean firing rate (Hz)
Trials
Cue
1 s
Fano factor
Foreperiod
Delay Preferred
Delay Least Preferred
Mean firing rate (Hz)
Pearsonr = 0.25
pvalue = 0.5571

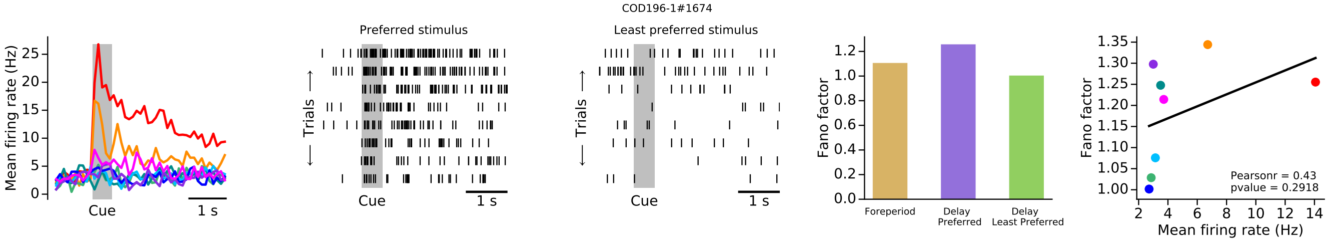
COD196-1#1674
Preferred stimulus
Least preferred stimulus
Mean firing rate (Hz)
Trials
Cue
1 s
Fano factor
Foreperiod
Delay Preferred
Delay Least Preferred
Mean firing rate (Hz)
Pearsonr = 0.43
pvalue = 0.2918

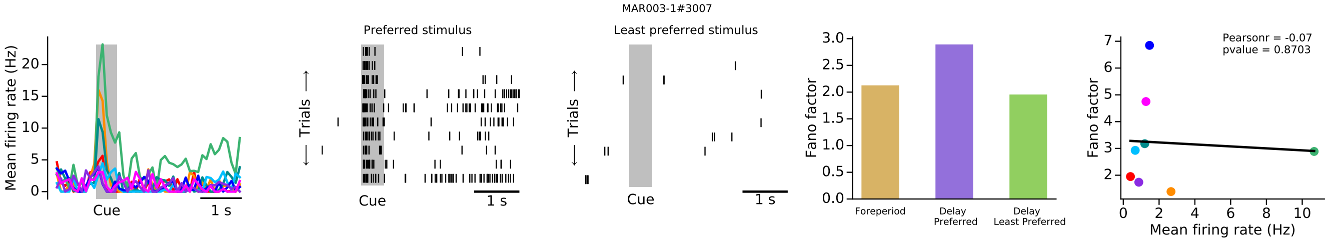
MAR003-1#3007
Preferred stimulus
Least preferred stimulus
Mean firing rate (Hz)
Trials
Cue
1 s
Fano factor
Foreperiod
Delay Preferred
Delay Least Preferred
Mean firing rate (Hz)
Pearsonr = -0.07
pvalue = 0.8703

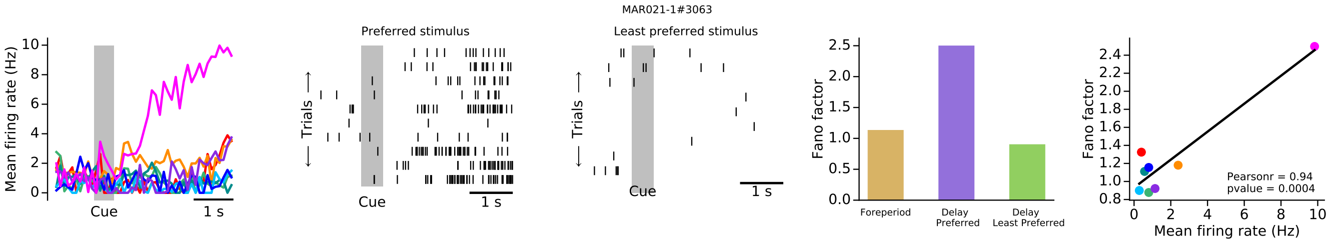
MAR021-1#3063
Preferred stimulus
Least preferred stimulus
Mean firing rate (Hz)
Trials
Cue
1 s
Fano factor
Foreperiod
Delay Preferred
Delay Least Preferred
Mean firing rate (Hz)
Pearsonr = 0.94
pvalue = 0.0004

MAR021-1#3065

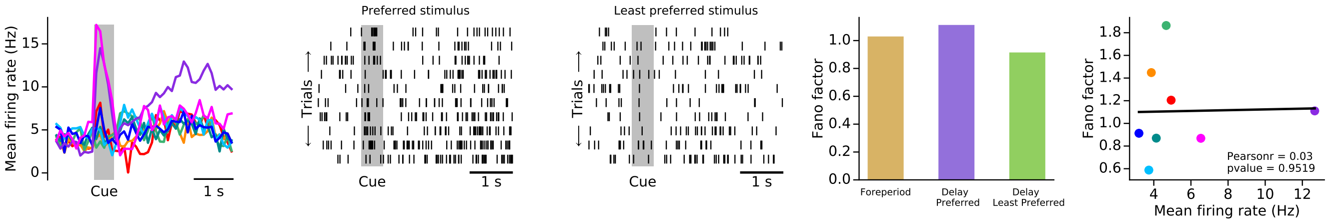


MAR022-1#3067

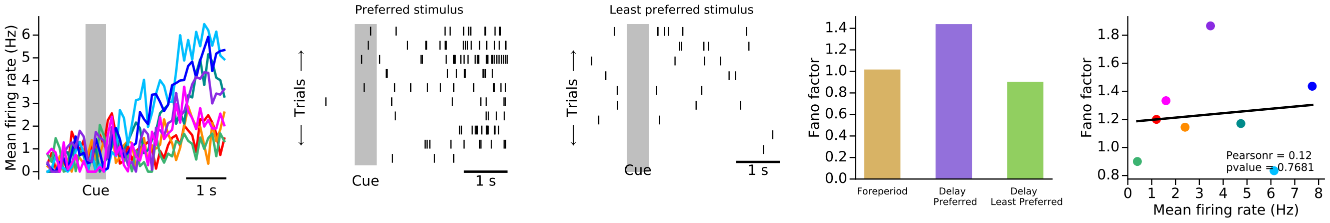


MAR023-1#3072

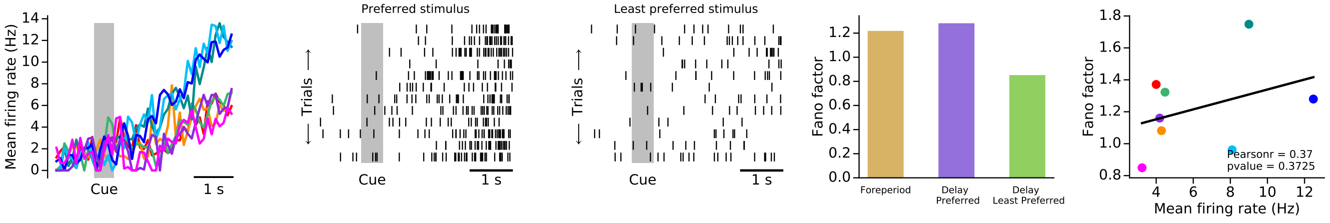


MAR024-1#3078

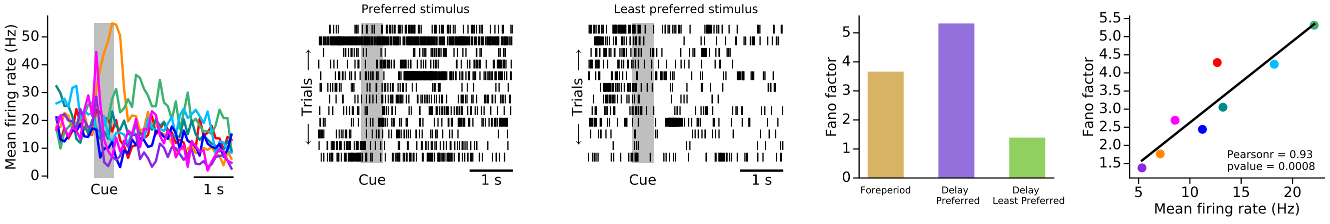


MAR025-1#3081

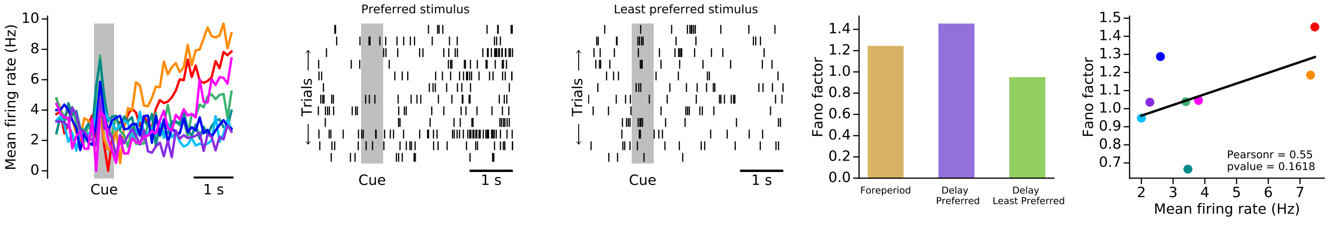

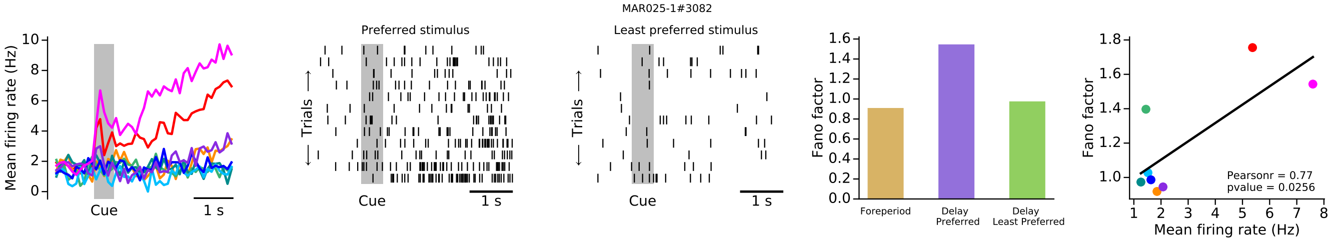
MAR025-1#3082
Mean firing rate (Hz)
Cue
1 s
Preferred stimulus
Trials
Least preferred stimulus
Fano factor
Foreperiod
Delay Preferred
Delay Least Preferred
Pearsonr = 0.77
pvalue = 0.0256
Mean firing rate (Hz)

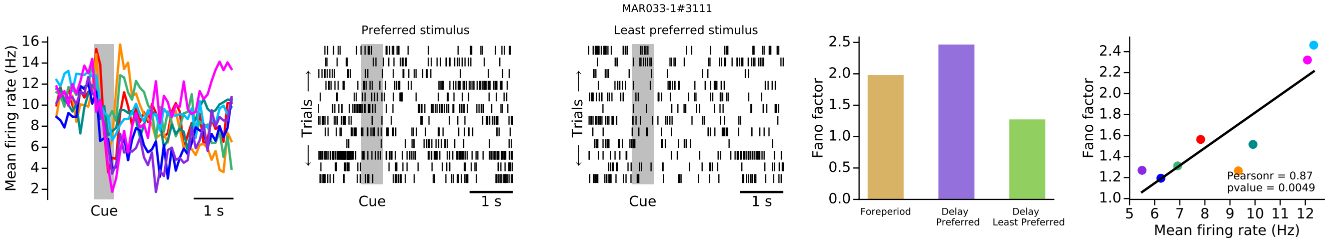
MAR033-1#3111
Mean firing rate (Hz)
Cue
1 s
Preferred stimulus
Trials
Least preferred stimulus
Fano factor
Foreperiod
Delay Preferred
Delay Least Preferred
Pearsonr = 0.87
pvalue = 0.0049
Mean firing rate (Hz)

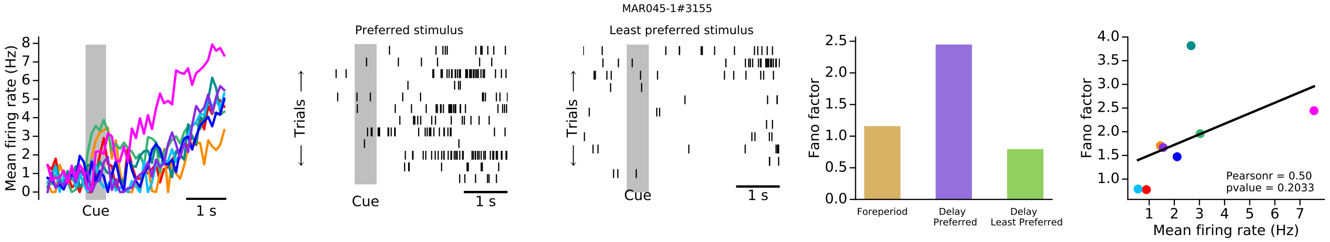
MAR045-1#3155
Mean firing rate (Hz)
Cue
1 s
Preferred stimulus
Trials
Least preferred stimulus
Fano factor
Foreperiod
Delay Preferred
Delay Least Preferred
Pearsonr = 0.50
pvalue = 0.2033
Mean firing rate (Hz)

### Group 2

COD008-1#1014

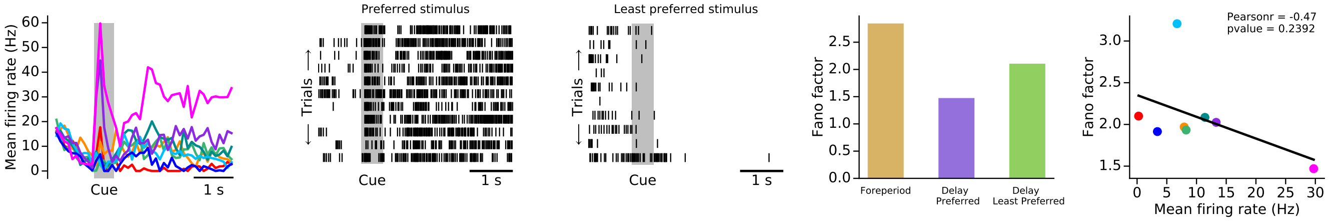


COD015-1#1024

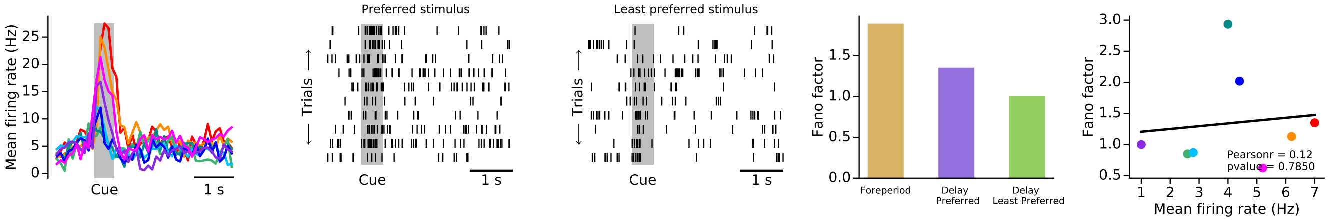


COD028-1#1054

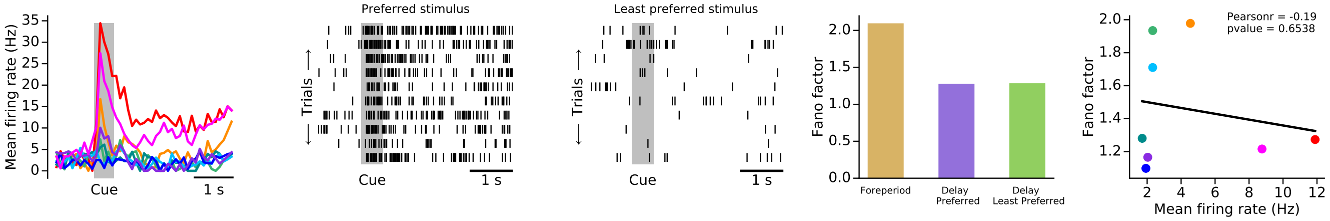


COD035-1#1065

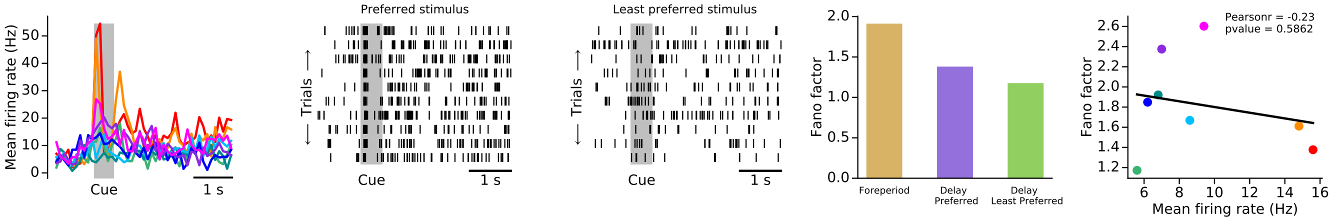


COD036-1#1068

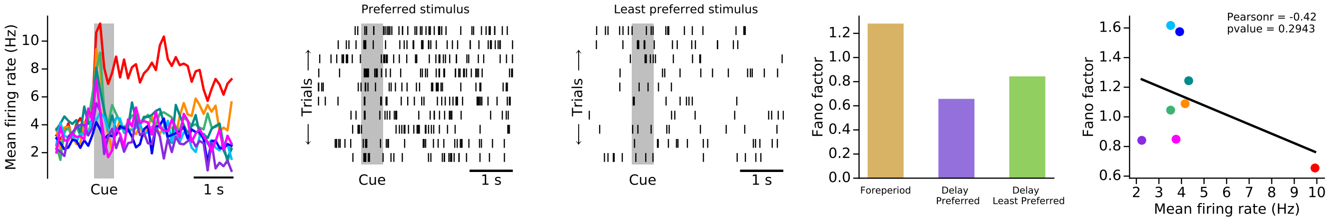

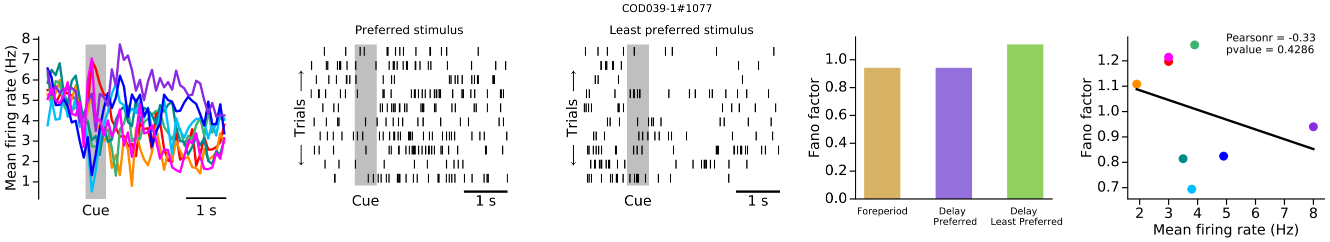
COD039-1#1077
Mean firing rate (Hz)
Cue
1 s
Preferred stimulus
Least preferred stimulus
Trials
Fano factor
Foreperiod
Delay Preferred
Delay Least Preferred
Pearsonr = -0.33
pvalue = 0.4286
Mean firing rate (Hz)

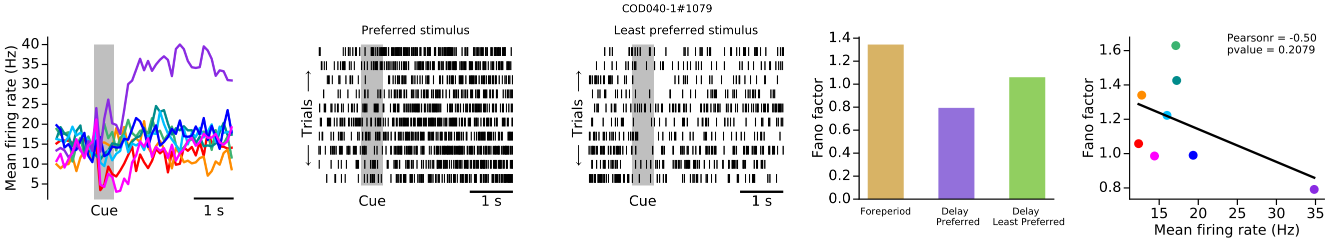
COD040-1#1079
Mean firing rate (Hz)
Cue
1 s
Preferred stimulus
Least preferred stimulus
Trials
Fano factor
Foreperiod
Delay Preferred
Delay Least Preferred
Pearsonr = -0.50
pvalue = 0.2079
Mean firing rate (Hz)

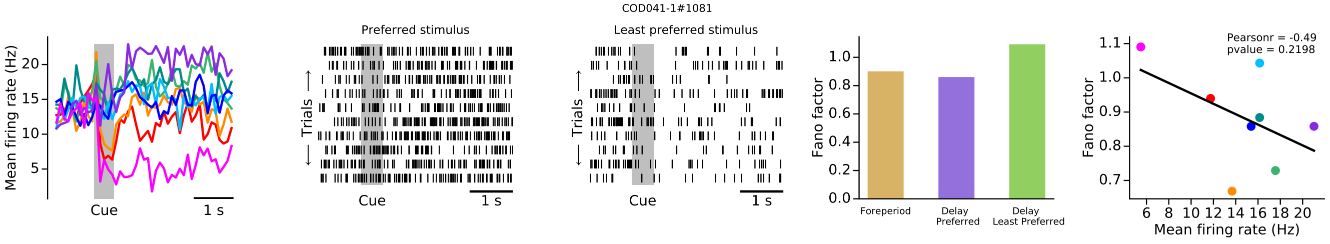
COD041-1#1081
Mean firing rate (Hz)
Cue
1 s
Preferred stimulus
Least preferred stimulus
Trials
Fano factor
Foreperiod
Delay Preferred
Delay Least Preferred
Pearsonr = -0.49
pvalue = 0.2198
Mean firing rate (Hz)

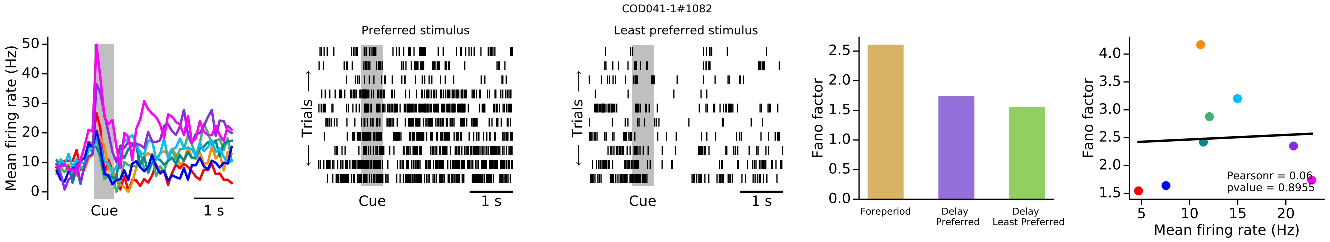
COD041-1#1082
Mean firing rate (Hz)
Cue
1 s
Preferred stimulus
Least preferred stimulus
Trials
Fano factor
Foreperiod
Delay Preferred
Delay Least Preferred
Pearsonr = 0.06
pvalue = 0.8955
Mean firing rate (Hz)

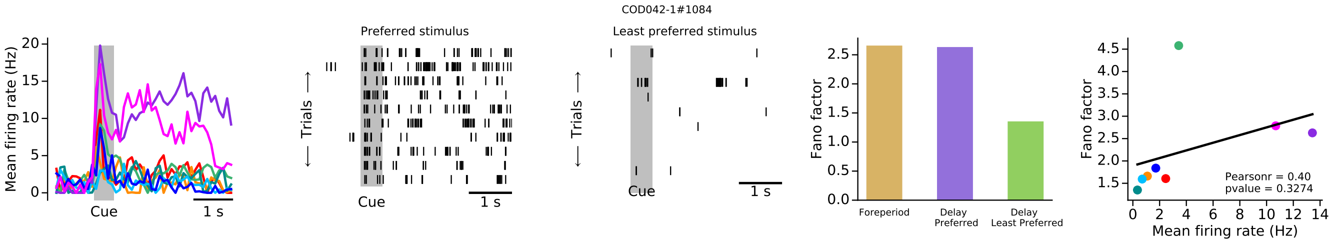
COD042-1#1084
Mean firing rate (Hz)
Cue
1 s
Preferred stimulus
Least preferred stimulus
Trials
Fano factor
Foreperiod
Delay Preferred
Delay Least Preferred
Pearsonr = 0.40
pvalue = 0.3274
Mean firing rate (Hz)

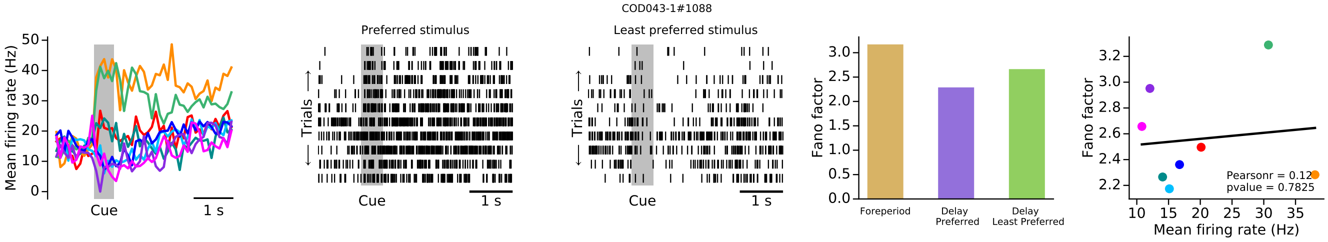
COD043-1#1088
Mean firing rate (Hz)
Cue
1 s
Preferred stimulus
Least preferred stimulus
Trials
Fano factor
Foreperiod
Delay Preferred
Delay Least Preferred
Pearsonr = 0.12
pvalue = 0.7825
Mean firing rate (Hz)

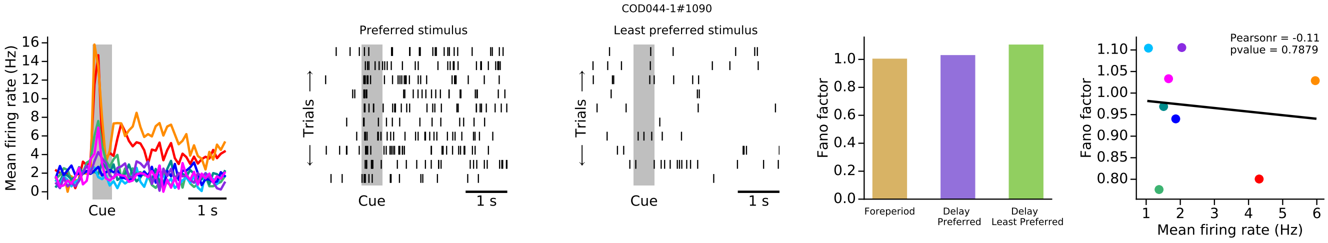
COD044-1#1090
Mean firing rate (Hz)
Cue
1 s
Preferred stimulus
Least preferred stimulus
Trials
Fano factor
Foreperiod
Delay Preferred
Delay Least Preferred
Pearsonr = -0.11
pvalue = 0.7879
Mean firing rate (Hz)

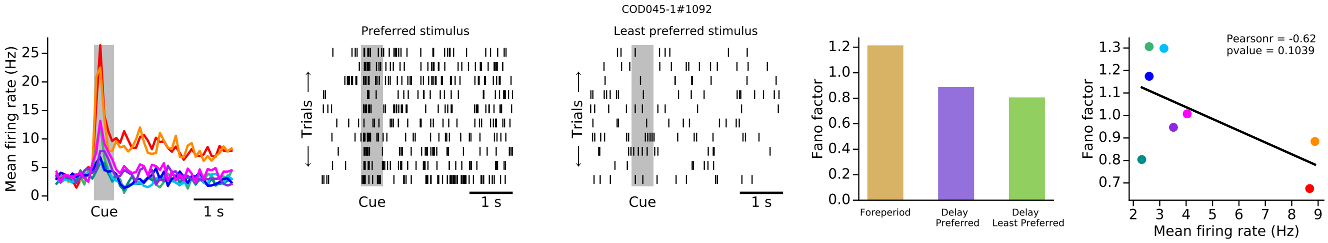
COD045-1#1092
Mean firing rate (Hz)
Cue
1 s
Preferred stimulus
Least preferred stimulus
Trials
Fano factor
Foreperiod
Delay Preferred
Delay Least Preferred
Pearsonr = -0.62
pvalue = 0.1039
Mean firing rate (Hz)

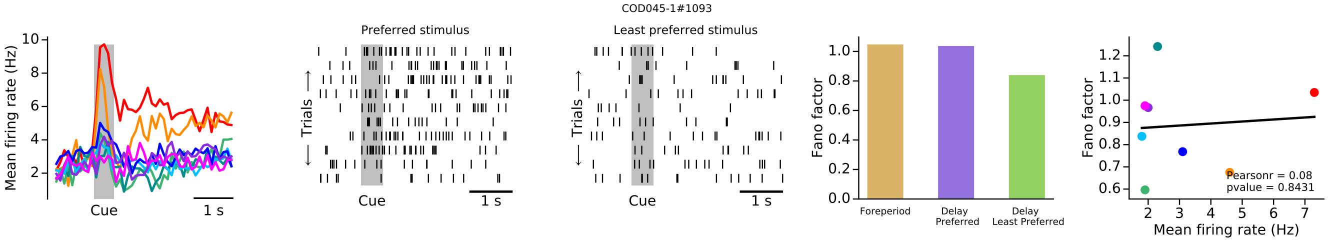
COD045-1#1093
Mean firing rate (Hz)
Cue
1 s
Preferred stimulus
Least preferred stimulus
Trials
Fano factor
Foreperiod
Delay Preferred
Delay Least Preferred
Pearsonr = 0.08
pvalue = 0.8431
Mean firing rate (Hz)

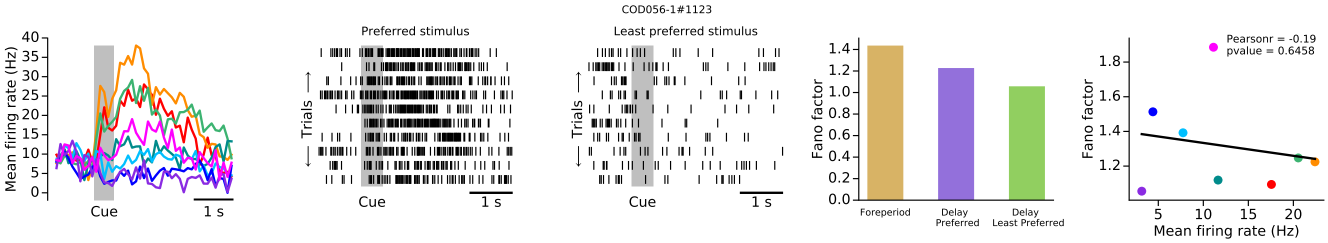
COD056-1#1123
Mean firing rate (Hz)
Cue
1 s
Preferred stimulus
Least preferred stimulus
Trials
Fano factor
Foreperiod
Delay Preferred
Delay Least Preferred
Pearsonr = -0.19
pvalue = 0.6458
Mean firing rate (Hz)

COD056-1#1124

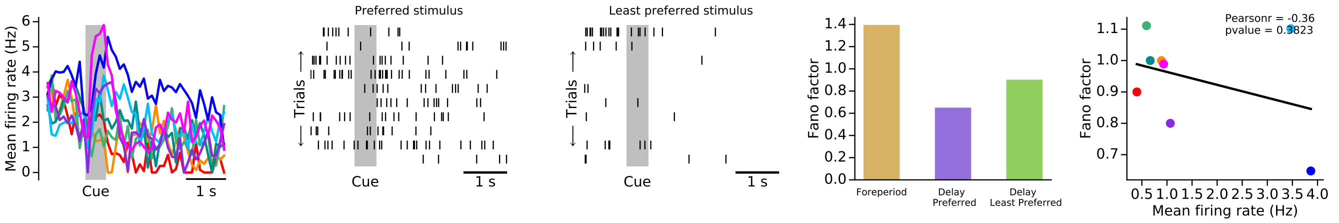


COD058-1#1133

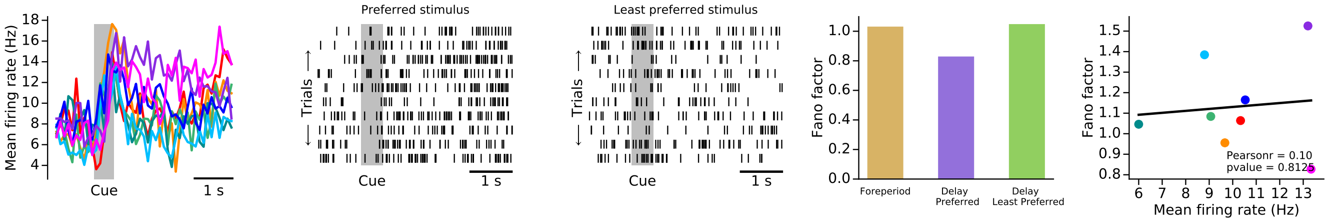


COD058-1#1134

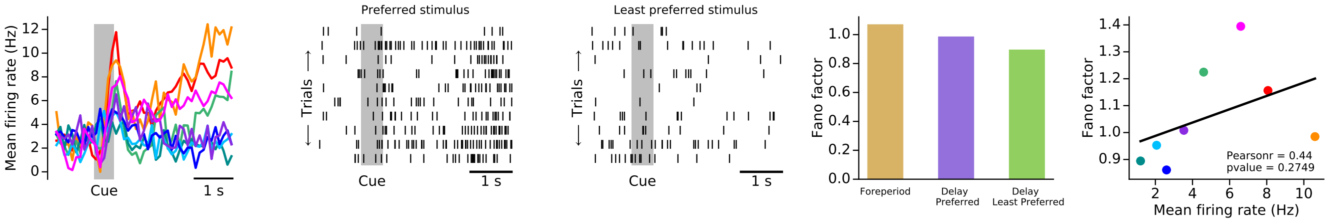


COD060-1#1138

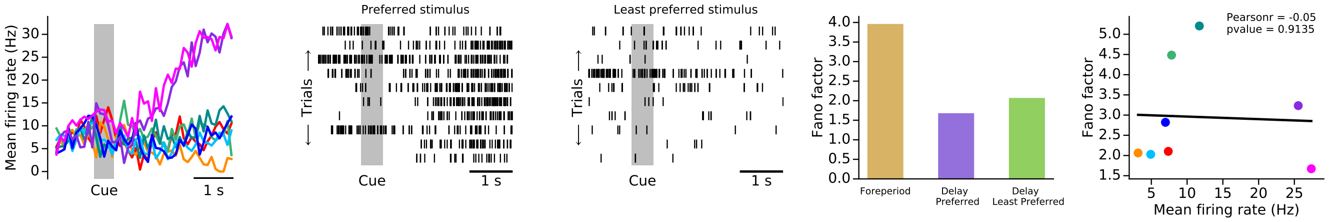


COD062-1#1147

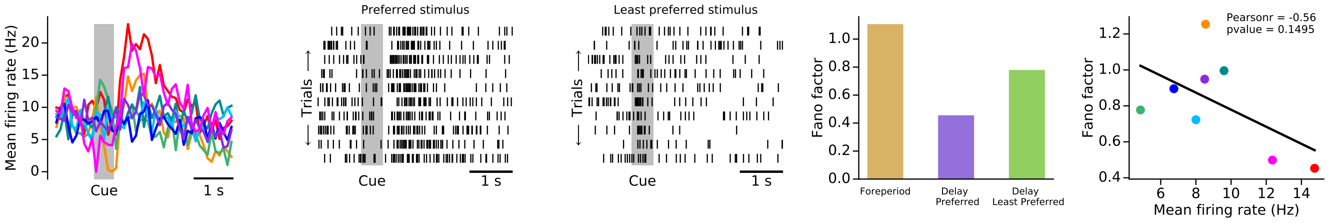

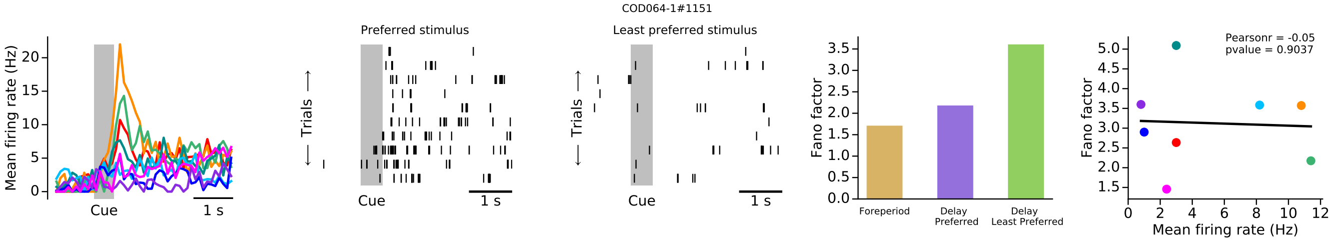
COD064-1#1151
Mean firing rate (Hz)
Cue
1 s
Preferred stimulus
Least preferred stimulus
Trials
Fano factor
Foreperiod
Delay Preferred
Delay Least Preferred
Pearsonr = -0.05
pvalue = 0.9037
Mean firing rate (Hz)

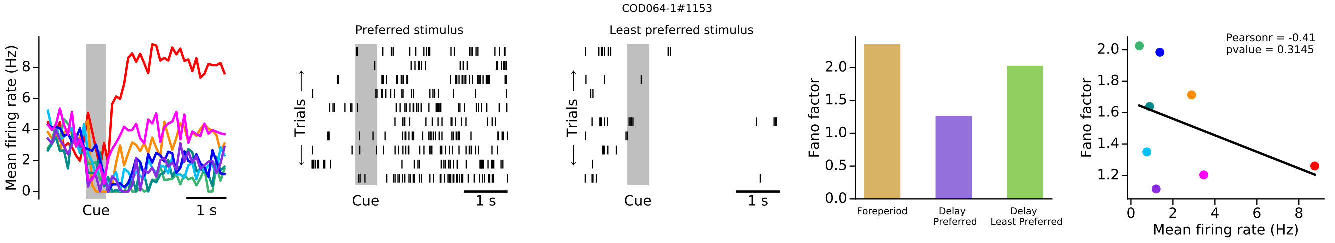
COD064-1#1153
Mean firing rate (Hz)
Cue
1 s
Preferred stimulus
Least preferred stimulus
Trials
Fano factor
Foreperiod
Delay Preferred
Delay Least Preferred
Pearsonr = -0.41
pvalue = 0.3145
Mean firing rate (Hz)

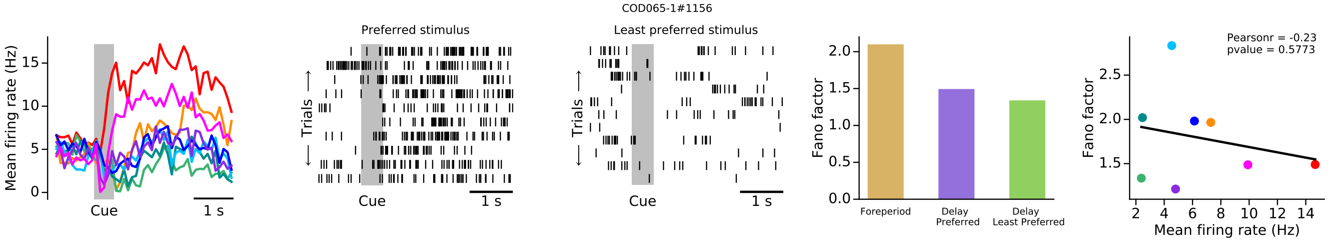
COD065-1#1156
Mean firing rate (Hz)
Cue
1 s
Preferred stimulus
Least preferred stimulus
Trials
Fano factor
Foreperiod
Delay Preferred
Delay Least Preferred
Pearsonr = -0.23
pvalue = 0.5773
Mean firing rate (Hz)

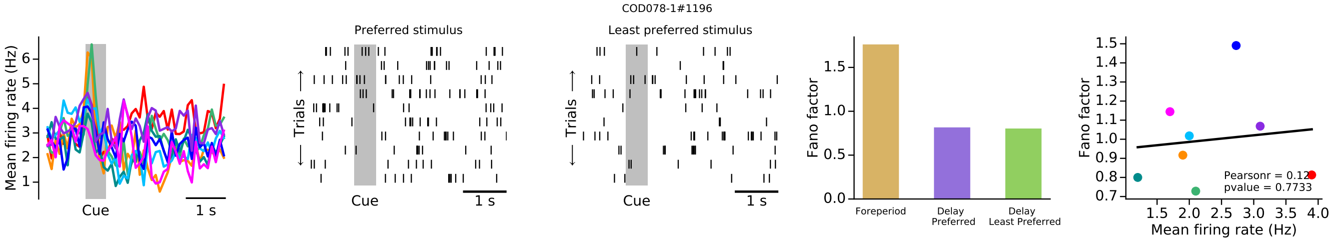
COD078-1#1196
Mean firing rate (Hz)
Cue
1 s
Preferred stimulus
Least preferred stimulus
Trials
Fano factor
Foreperiod
Delay Preferred
Delay Least Preferred
pvalue = 0.7733
Mean firing rate (Hz)

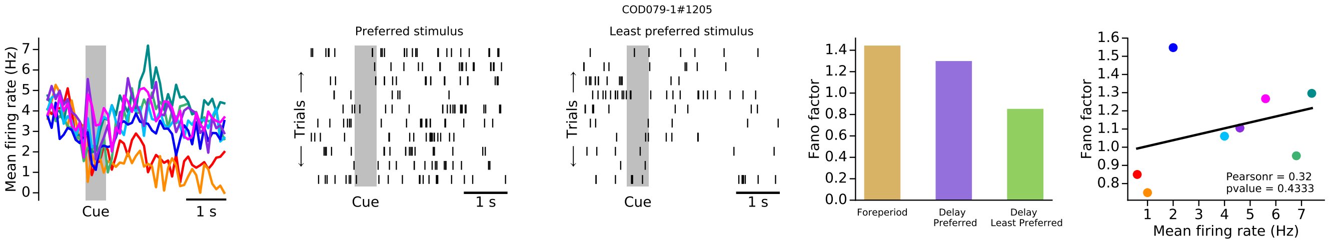
COD079-1#1205
Mean firing rate (Hz)
Cue
1 s
Preferred stimulus
Least preferred stimulus
Trials
Fano factor
Foreperiod
Delay Preferred
Delay Least Preferred
Pearsonr = 0.32
pvalue = 0.4333
Mean firing rate (Hz)

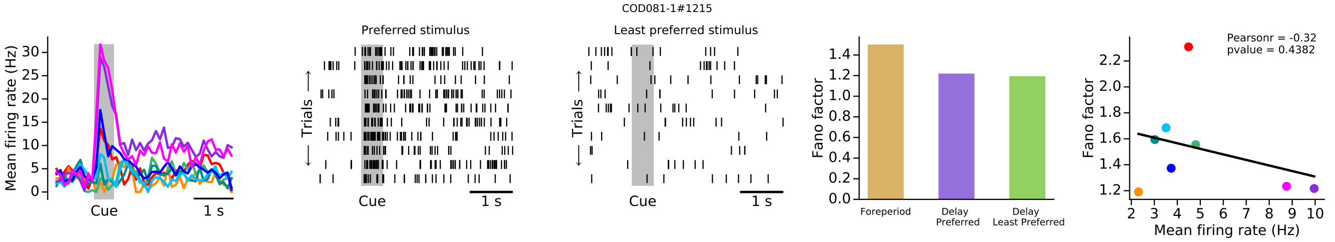
COD081-1#1215
Preferred stimulus
Least preferred stimulus
Mean firing rate (Hz)
Trials
Cue
1 s
Fano factor
Foreperiod
Delay Preferred
Delay Least Preferred
Pearsonr = -0.32
pvalue = 0.4382
Mean firing rate (Hz)

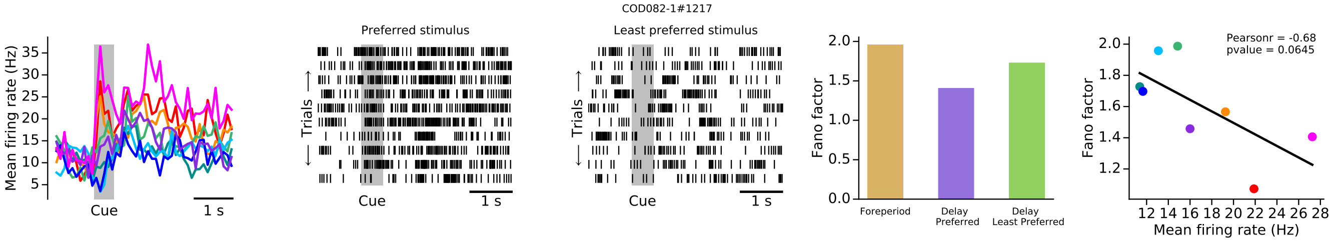
COD082-1#1217
Preferred stimulus
Least preferred stimulus
Mean firing rate (Hz)
Trials
Cue
1 s
Fano factor
Foreperiod
Delay Preferred
Delay Least Preferred
Pearsonr = -0.68
pvalue = 0.0645
Mean firing rate (Hz)

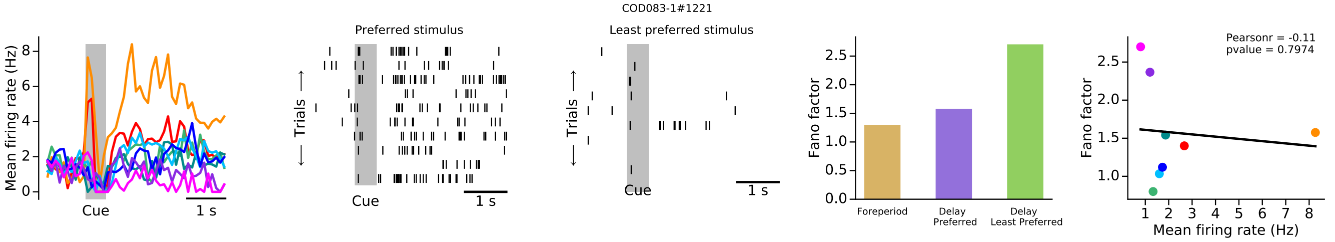
COD083-1#1221
Preferred stimulus
Least preferred stimulus
Mean firing rate (Hz)
Trials
Cue
1 s
Fano factor
Foreperiod
Delay Preferred
Delay Least Preferred
Pearsonr = -0.11
pvalue = 0.7974
Mean firing rate (Hz)

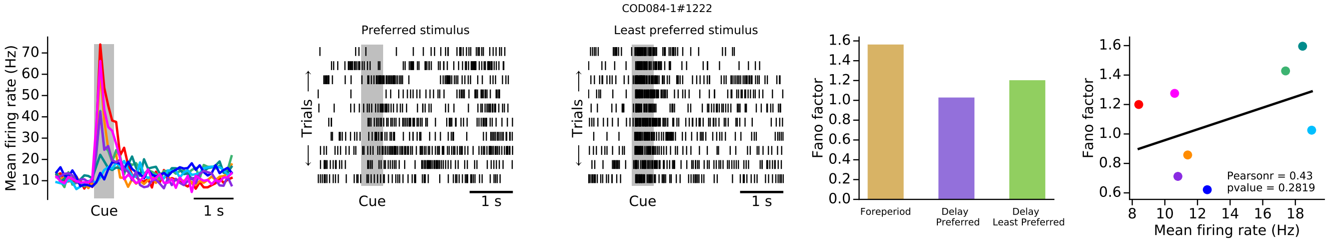
COD084-1#1222
Preferred stimulus
Least preferred stimulus
Mean firing rate (Hz)
Trials
Cue
1 s
Fano factor
Foreperiod
Delay Preferred
Delay Least Preferred
Pearsonr = 0.43
pvalue = 0.2819
Mean firing rate (Hz)

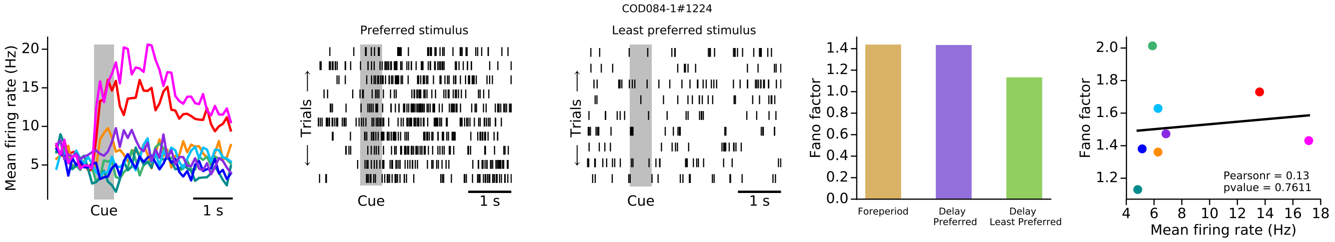
COD084-1#1224
Preferred stimulus
Least preferred stimulus
Mean firing rate (Hz)
Trials
Cue
1 s
Fano factor
Foreperiod
Delay Preferred
Delay Least Preferred
Pearsonr = 0.13
pvalue = 0.7611
Mean firing rate (Hz)

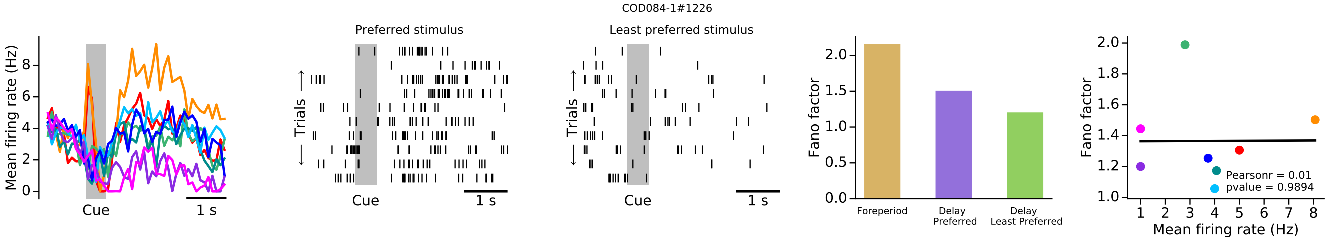
COD084-1#1226
Mean firing rate (Hz)
Cue
1 s
Preferred stimulus
Least preferred stimulus
Trials
Fano factor
Foreperiod
Delay Preferred
Delay Least Preferred
Pearsonr = 0.01
pvalue = 0.9894
Mean firing rate (Hz)

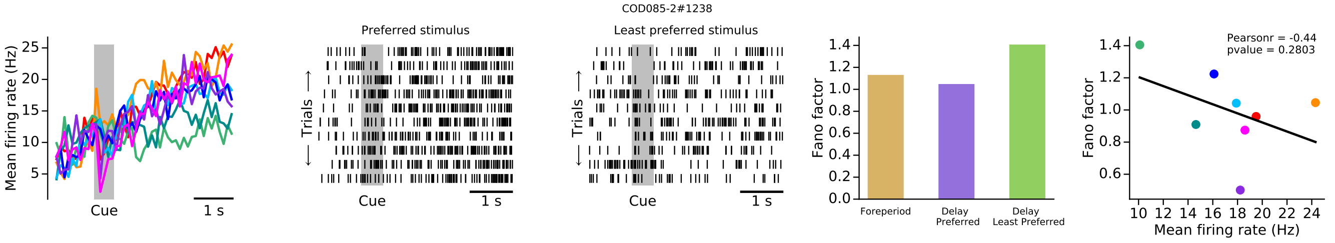
COD085-2#1238
Mean firing rate (Hz)
Cue
1 s
Preferred stimulus
Least preferred stimulus
Trials
Fano factor
Foreperiod
Delay Preferred
Delay Least Preferred
Pearsonr = -0.44
pvalue = 0.2803
Mean firing rate (Hz)

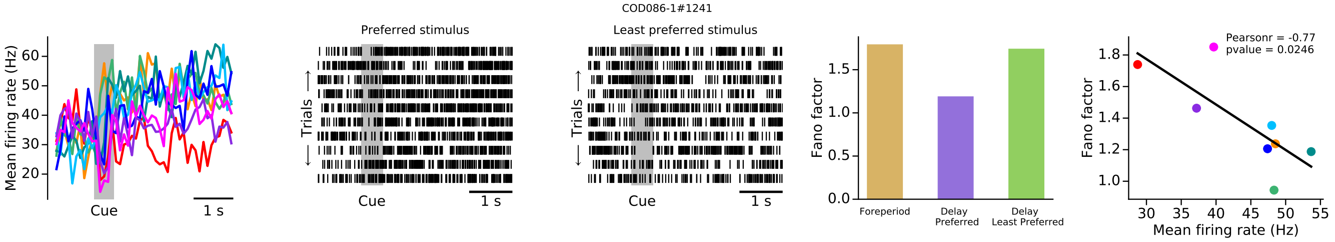
COD086-1#1241
Mean firing rate (Hz)
Cue
1 s
Preferred stimulus
Least preferred stimulus
Trials
Fano factor
Foreperiod
Delay Preferred
Delay Least Preferred
Pearsonr = -0.77
pvalue = 0.0246
Mean firing rate (Hz)

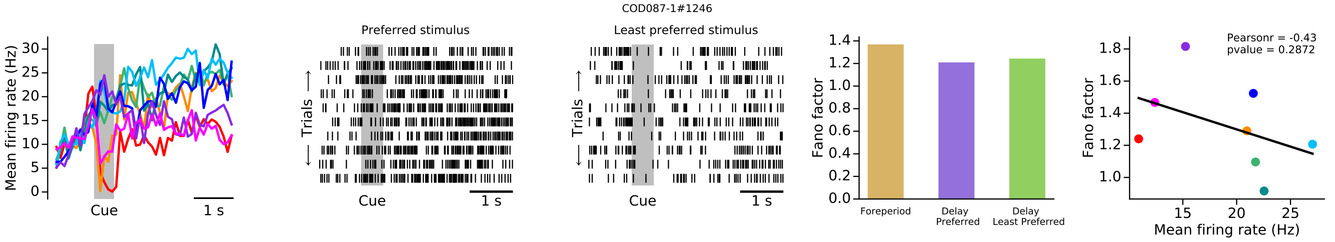
COD087-1#1246
Mean firing rate (Hz)
Cue
1 s
Preferred stimulus
Least preferred stimulus
Trials
Fano factor
Foreperiod
Delay Preferred
Delay Least Preferred
Pearsonr = -0.43
pvalue = 0.2872
Mean firing rate (Hz)

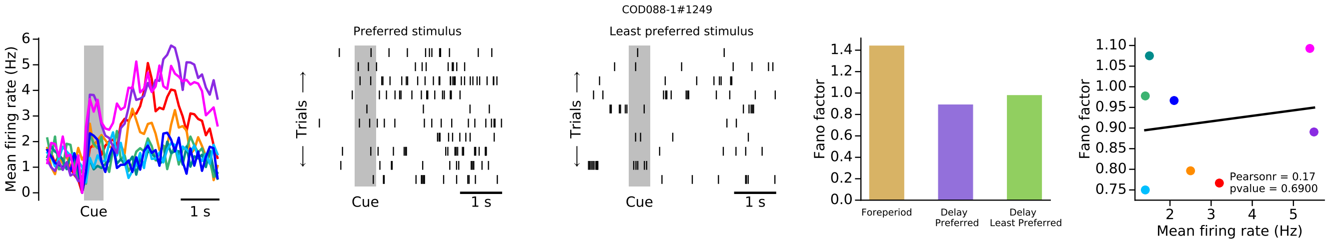
COD088-1#1249
Mean firing rate (Hz)
Cue
1 s
Preferred stimulus
Least preferred stimulus
Trials
Fano factor
Foreperiod
Delay Preferred
Delay Least Preferred
Pearsonr = 0.17
pvalue = 0.6900
Mean firing rate (Hz)

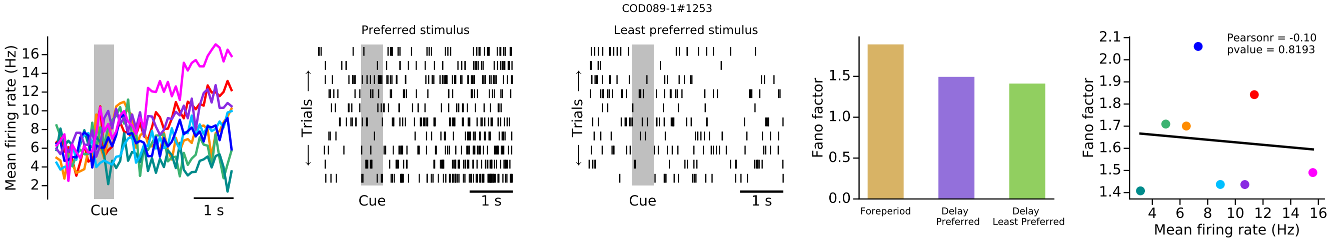
COD089-1#1253
Preferred stimulus
Least preferred stimulus
Mean firing rate (Hz)
Cue
1 s
Trials
Fano factor
Foreperiod
Delay Preferred
Delay Least Preferred
Pearsonr = -0.10
pvalue = 0.8193
Mean firing rate (Hz)

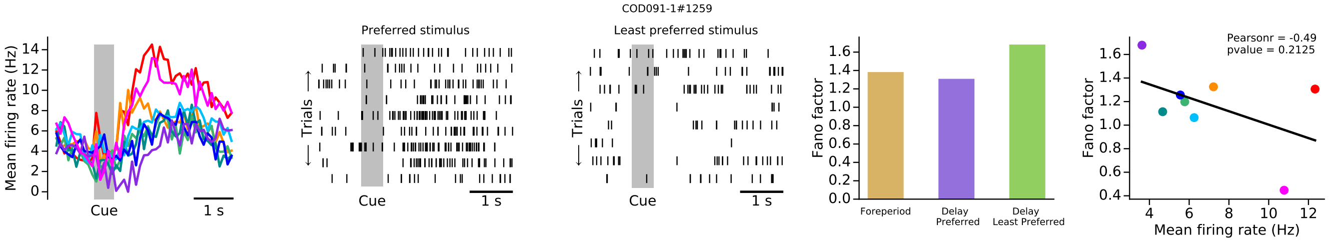
COD091-1#1259
Preferred stimulus
Least preferred stimulus
Mean firing rate (Hz)
Cue
1 s
Trials
Fano factor
Foreperiod
Delay Preferred
Delay Least Preferred
Pearsonr = -0.49
pvalue = 0.2125
Mean firing rate (Hz)

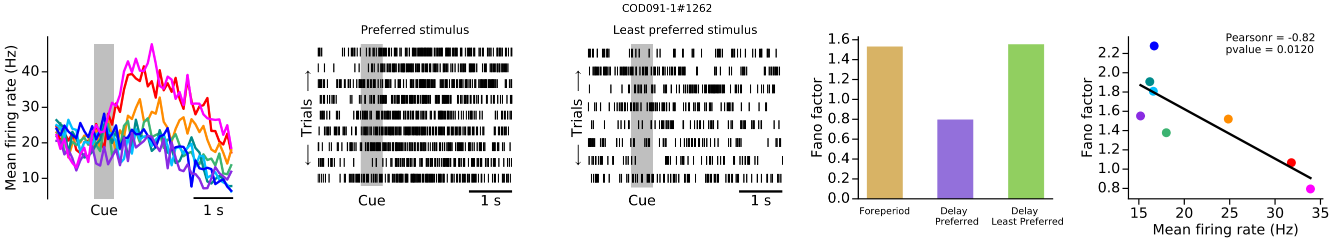
COD091-1#1262
Preferred stimulus
Least preferred stimulus
Mean firing rate (Hz)
Cue
1 s
Trials
Fano factor
Foreperiod
Delay Preferred
Delay Least Preferred
Pearsonr = -0.82
pvalue = 0.0120
Mean firing rate (Hz)

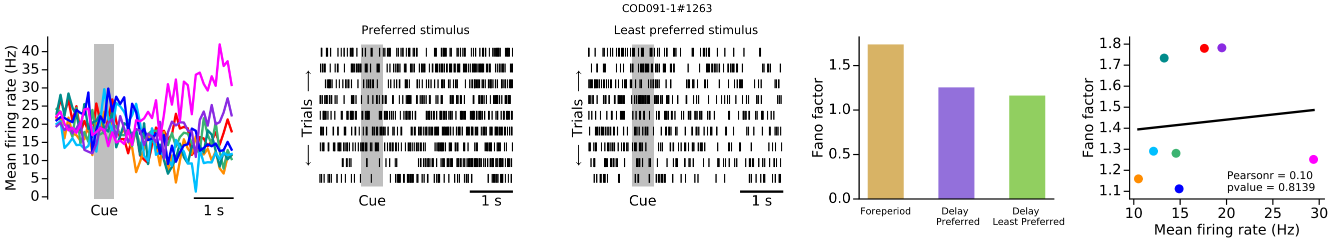
COD091-1#1263
Preferred stimulus
Least preferred stimulus
Mean firing rate (Hz)
Cue
1 s
Trials
Fano factor
Foreperiod
Delay Preferred
Delay Least Preferred
Pearsonr = 0.10
pvalue = 0.8139
Mean firing rate (Hz)

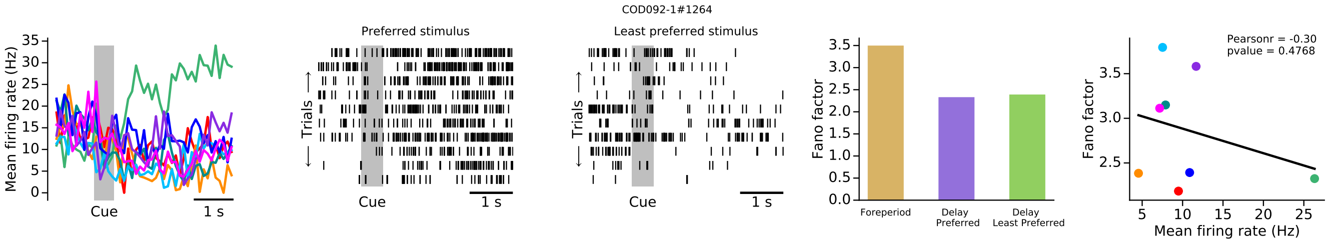
COD092-1#1264
Preferred stimulus
Least preferred stimulus
Mean firing rate (Hz)
Cue
1 s
Trials
Fano factor
Foreperiod
Delay Preferred
Delay Least Preferred
Pearsonr = -0.30
pvalue = 0.4768
Mean firing rate (Hz)

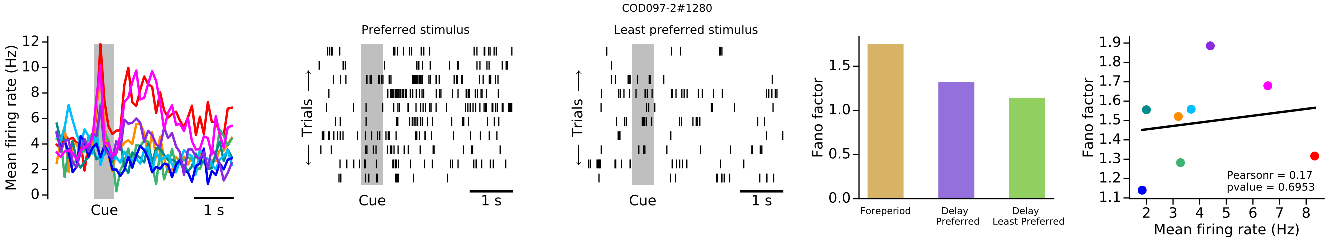
COD097-2#1280
Mean firing rate (Hz)
Cue
1 s
Preferred stimulus
Least preferred stimulus
Trials
Fano factor
Foreperiod
Delay Preferred
Delay Least Preferred
Pearsonr = 0.17
pvalue = 0.6953
Mean firing rate (Hz)

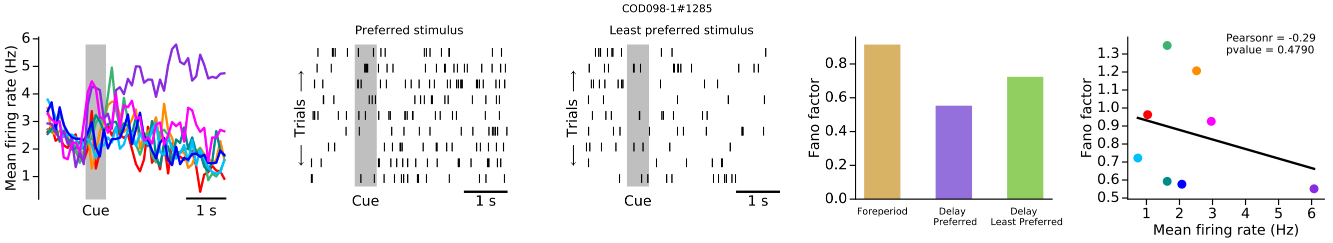
COD098-1#1285
Mean firing rate (Hz)
Cue
1 s
Preferred stimulus
Least preferred stimulus
Trials
Fano factor
Foreperiod
Delay Preferred
Delay Least Preferred
Pearsonr = -0.29
pvalue = 0.4790
Mean firing rate (Hz)

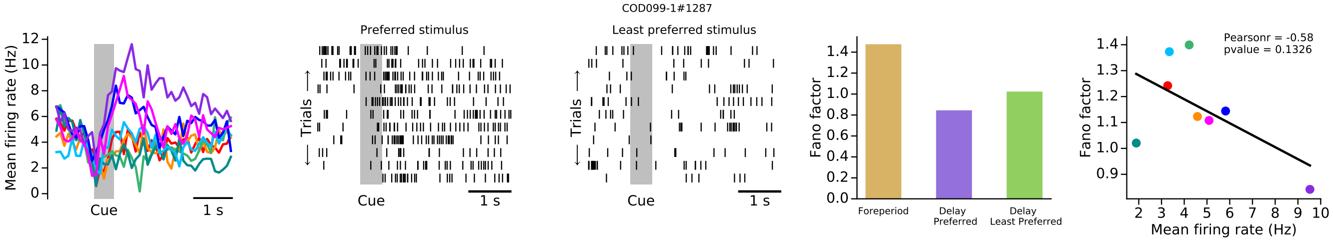
COD099-1#1287
Mean firing rate (Hz)
Cue
1 s
Preferred stimulus
Least preferred stimulus
Trials
Fano factor
Foreperiod
Delay Preferred
Delay Least Preferred
Pearsonr = -0.58
pvalue = 0.1326
Mean firing rate (Hz)

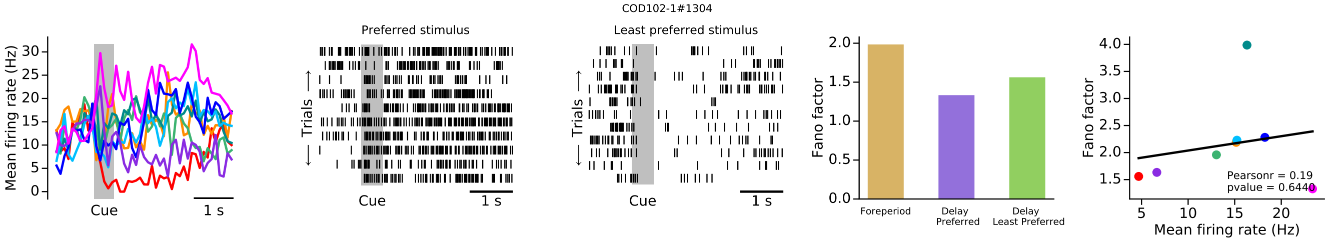
COD102-1#1304
Mean firing rate (Hz)
Cue
1 s
Preferred stimulus
Least preferred stimulus
Trials
Fano factor
Foreperiod
Delay Preferred
Delay Least Preferred
Pearsonr = 0.19
pvalue = 0.6440
Mean firing rate (Hz)

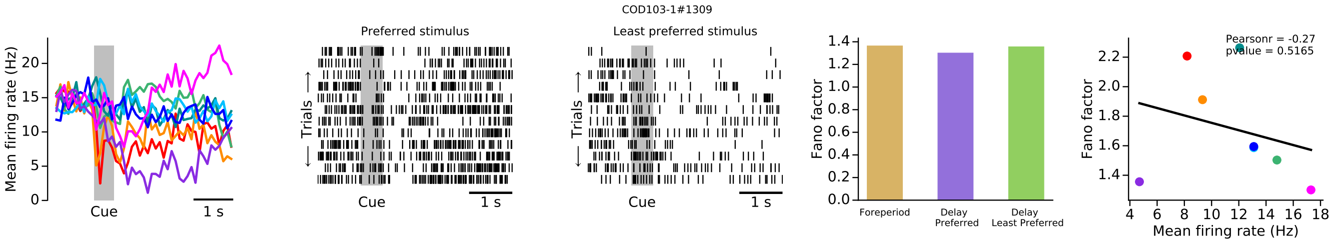
COD103-1#1309
Mean firing rate (Hz)
Cue
1 s
Preferred stimulus
Least preferred stimulus
Trials
Fano factor
Foreperiod
Delay Preferred
Delay Least Preferred
Pearsonr = -0.27
pvalue = 0.5165
Mean firing rate (Hz)

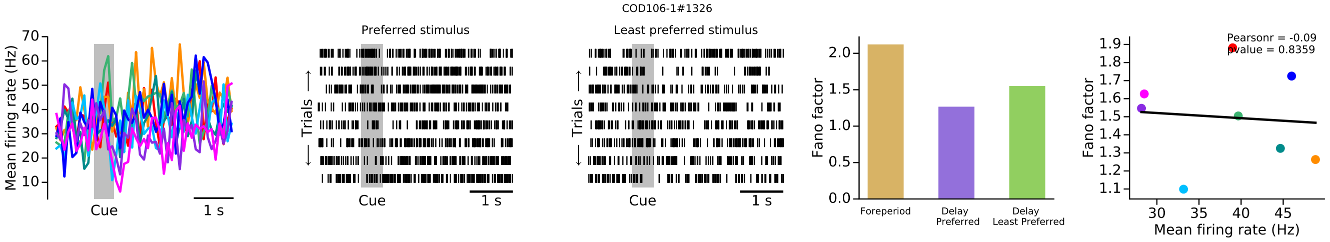
COD106-1#1326
Mean firing rate (Hz)
Cue
1 s
Preferred stimulus
Trials
Least preferred stimulus
Fano factor
Foreperiod
Delay Preferred
Delay Least Preferred
Pearsonr = -0.09
pvalue = 0.8359
Mean firing rate (Hz)

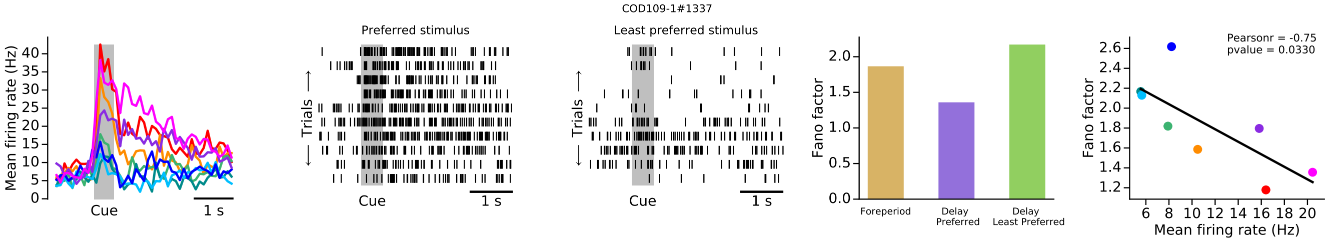
COD109-1#1337
Mean firing rate (Hz)
Cue
1 s
Preferred stimulus
Trials
Least preferred stimulus
Fano factor
Foreperiod
Delay Preferred
Delay Least Preferred
Pearsonr = -0.75
pvalue = 0.0330
Mean firing rate (Hz)

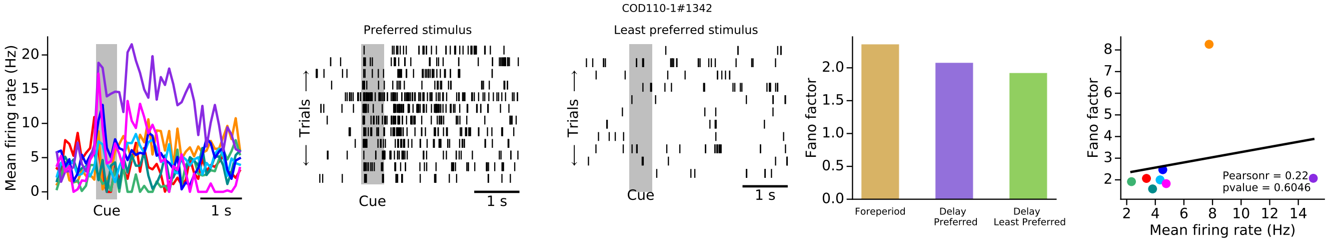
COD110-1#1342
Mean firing rate (Hz)
Cue
1 s
Preferred stimulus
Trials
Least preferred stimulus
Fano factor
Foreperiod
Delay Preferred
Delay Least Preferred
Pearsonr = 0.22
pvalue = 0.6046
Mean firing rate (Hz)

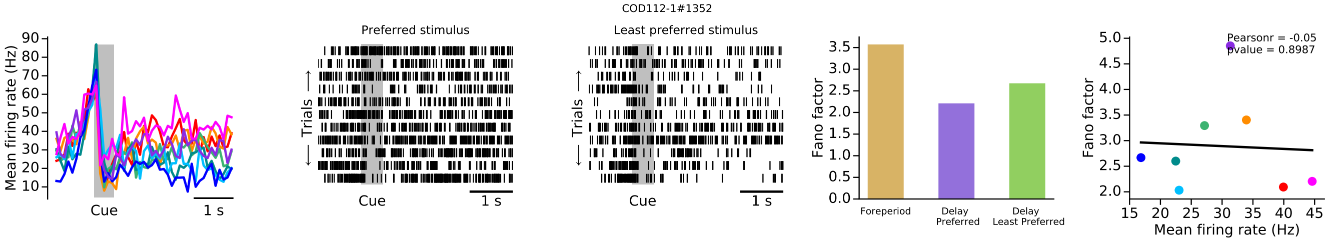
COD112-1#1352
Mean firing rate (Hz)
Cue
1 s
Preferred stimulus
Trials
Least preferred stimulus
Fano factor
Foreperiod
Delay Preferred
Delay Least Preferred
Pearsonr = -0.05
pvalue = 0.8987
Mean firing rate (Hz)

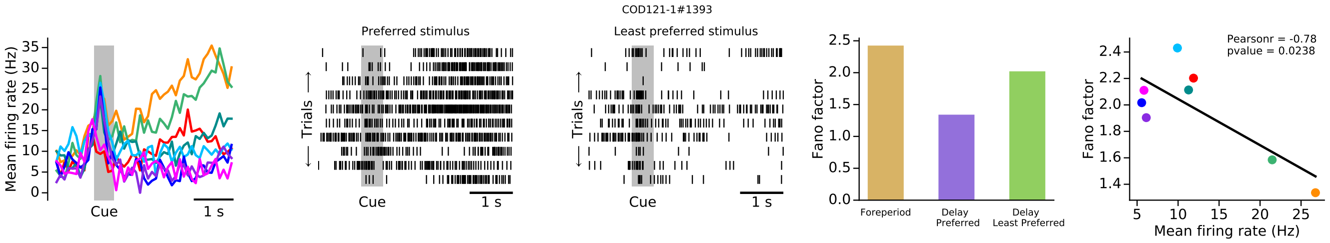
COD121-1#1393
Mean firing rate (Hz)
Cue
1 s
Preferred stimulus
Trials
Least preferred stimulus
Fano factor
Foreperiod
Delay Preferred
Delay Least Preferred
Pearsonr = -0.78
pvalue = 0.0238
Mean firing rate (Hz)

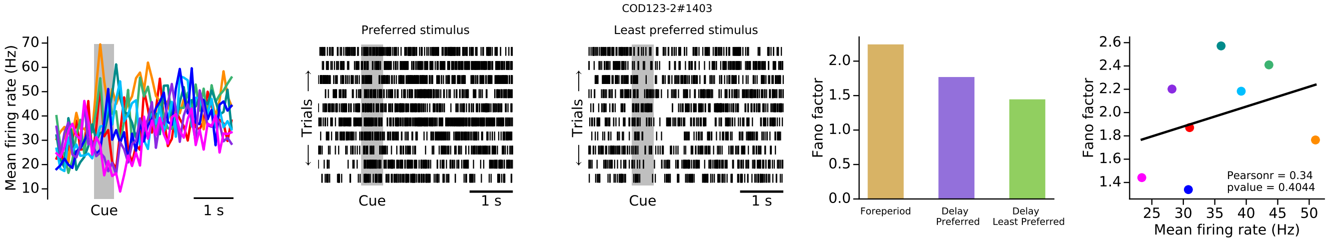
COD123-2#1403
Mean firing rate (Hz)
Cue
1 s
Preferred stimulus
Least preferred stimulus
Trials
Fano factor
Foreperiod
Delay Preferred
Delay Least Preferred
Pearsonr = 0.34
pvalue = 0.4044
Mean firing rate (Hz)

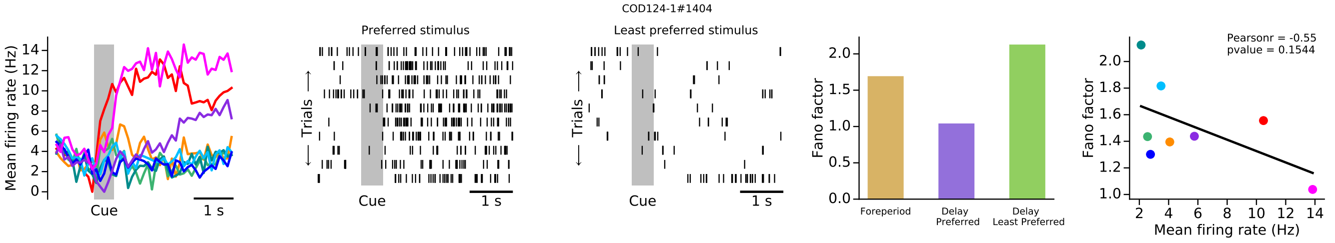
COD124-1#1404
Mean firing rate (Hz)
Cue
1 s
Preferred stimulus
Least preferred stimulus
Trials
Fano factor
Foreperiod
Delay Preferred
Delay Least Preferred
Pearsonr = -0.55
pvalue = 0.1544
Mean firing rate (Hz)

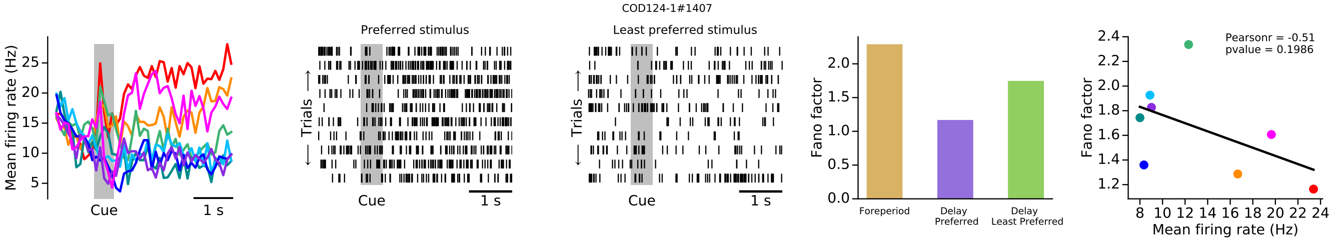
COD124-1#1407
Mean firing rate (Hz)
Cue
1 s
Preferred stimulus
Least preferred stimulus
Trials
Fano factor
Foreperiod
Delay Preferred
Delay Least Preferred
Pearsonr = -0.51
pvalue = 0.1986
Mean firing rate (Hz)

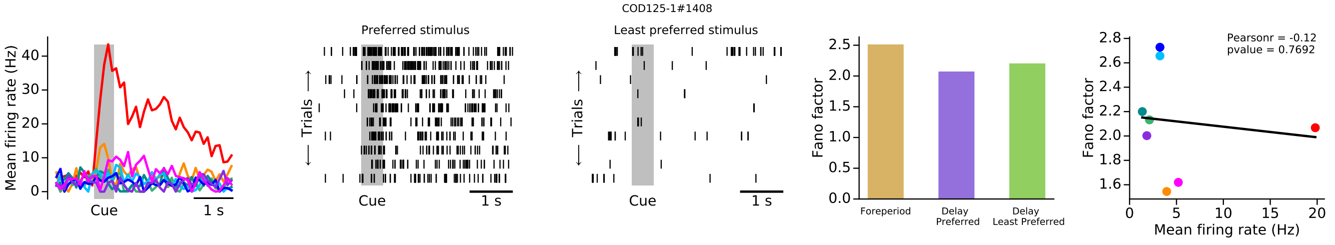
COD125-1#1408
Mean firing rate (Hz)
Cue
1 s
Preferred stimulus
Least preferred stimulus
Trials
Fano factor
Foreperiod
Delay Preferred
Delay Least Preferred
Pearsonr = -0.12
pvalue = 0.7692
Mean firing rate (Hz)

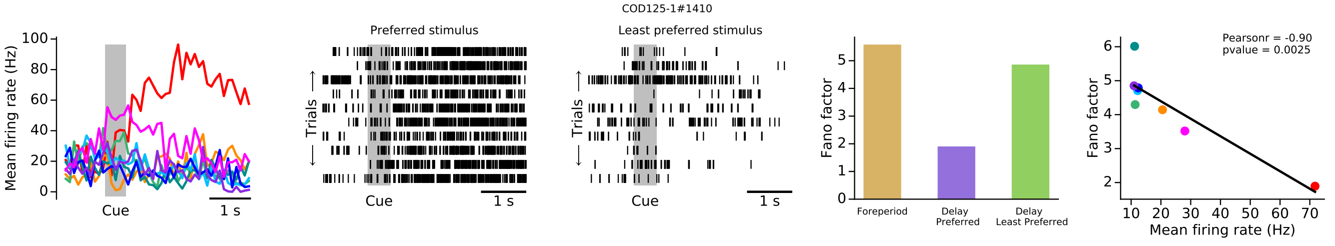
COD125-1#1410
Mean firing rate (Hz)
Cue
1 s
Preferred stimulus
Least preferred stimulus
Trials
Fano factor
Foreperiod
Delay Preferred
Delay Least Preferred
Pearsonr = -0.90
pvalue = 0.0025
Mean firing rate (Hz)

COD128-1#1422

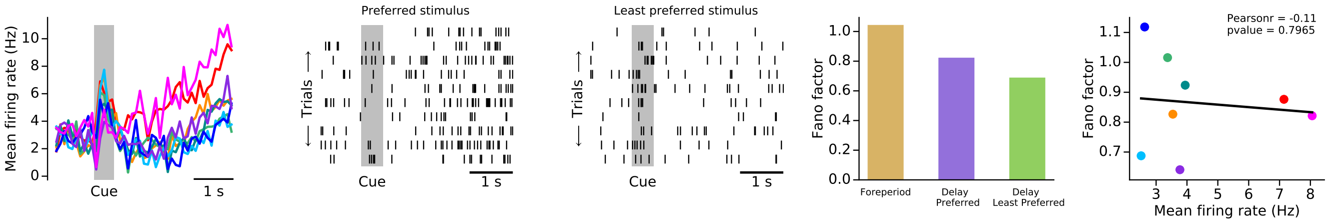


COD129-1#1426

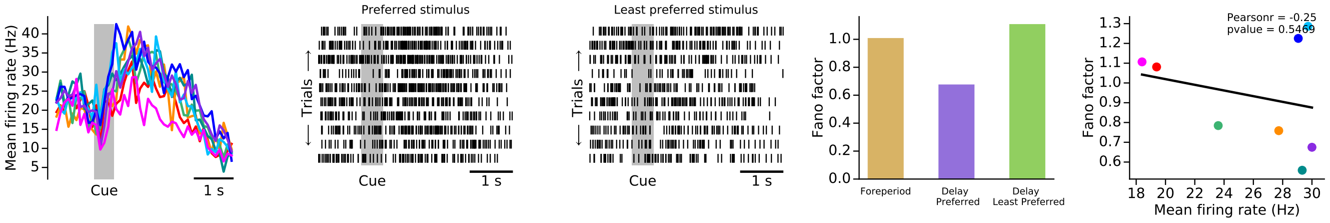


COD129-1#1427

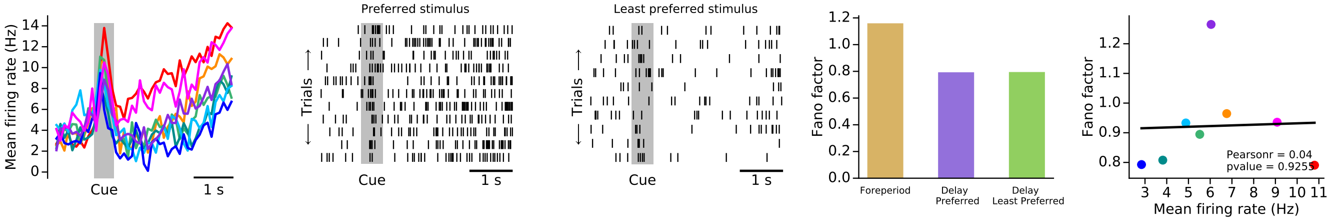


COD130-1#1432

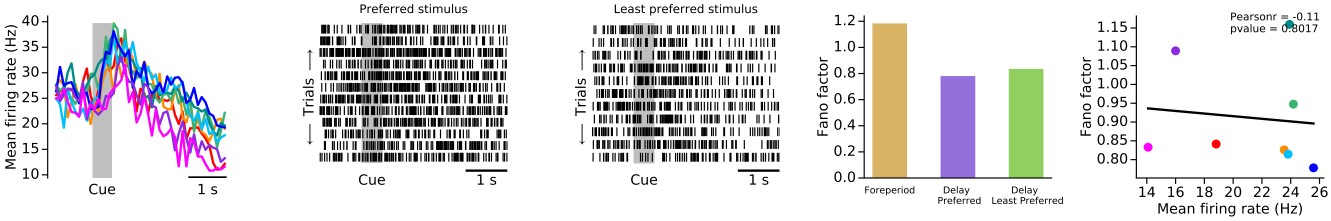


COD130-1#1434

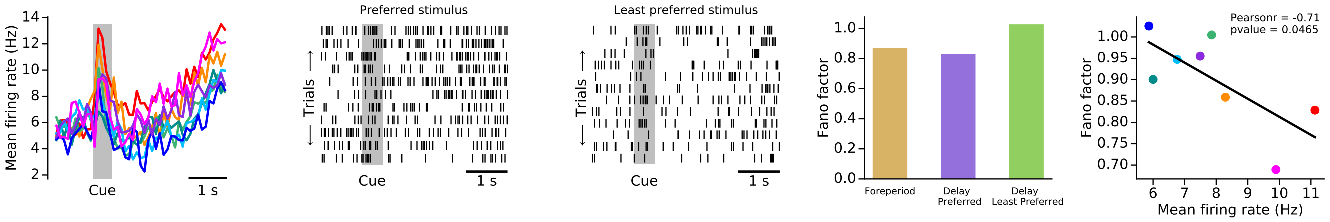

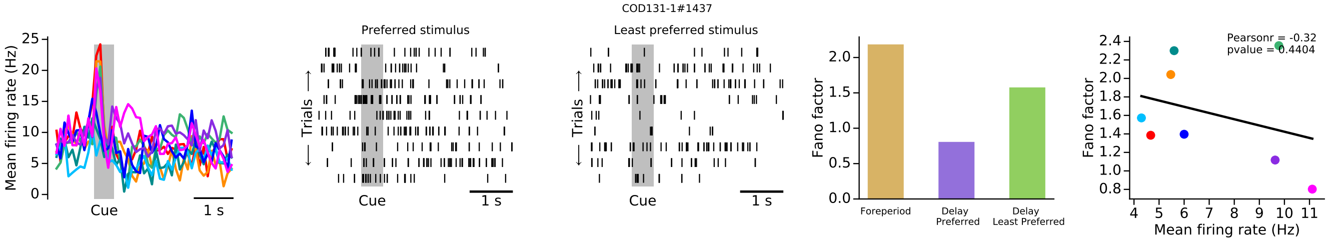
COD131-1#1437
Preferred stimulus
Least preferred stimulus
Mean firing rate (Hz)
Trials
Cue
1 s
Fano factor
Foreperiod
Delay Preferred
Delay Least Preferred
Pearsonr = -0.32
pvalue = 0.4404
Mean firing rate (Hz)

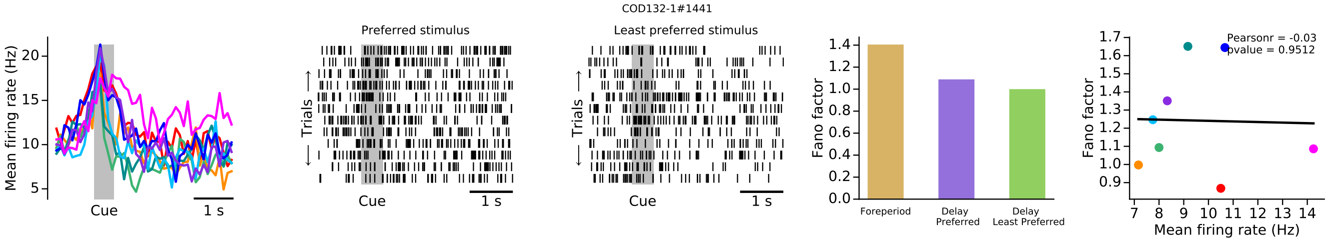
COD132-1#1441
Preferred stimulus
Least preferred stimulus
Mean firing rate (Hz)
Trials
Cue
1 s
Fano factor
Foreperiod
Delay Preferred
Delay Least Preferred
Pearsonr = -0.03
pvalue = 0.9512
Mean firing rate (Hz)

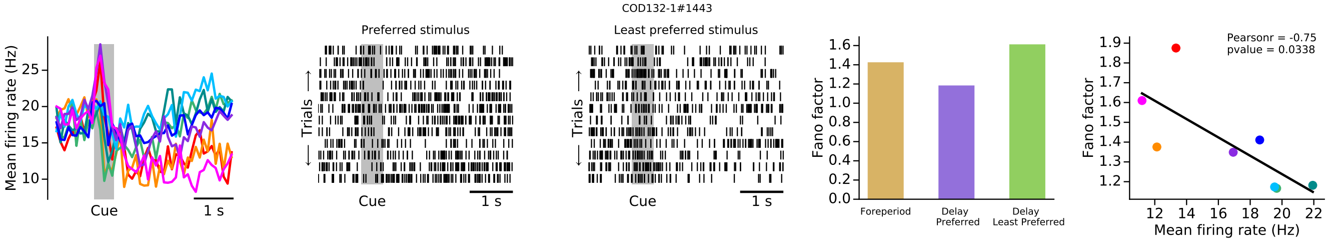
COD132-1#1443
Preferred stimulus
Least preferred stimulus
Mean firing rate (Hz)
Trials
Cue
1 s
Fano factor
Foreperiod
Delay Preferred
Delay Least Preferred
Pearsonr = -0.75
pvalue = 0.0338
Mean firing rate (Hz)

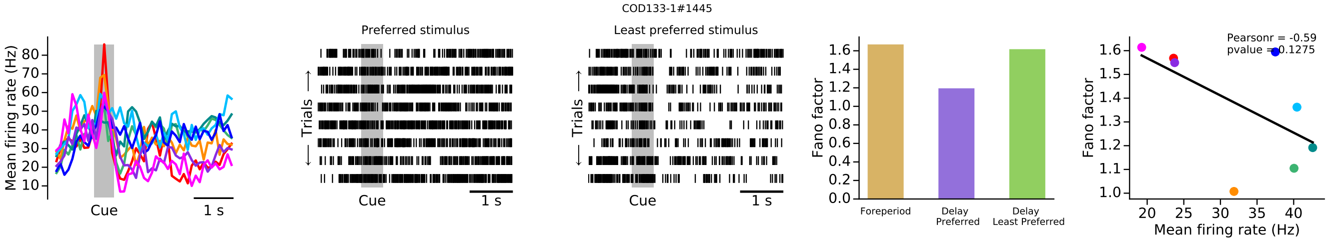
COD133-1#1445
Preferred stimulus
Least preferred stimulus
Mean firing rate (Hz)
Trials
Cue
1 s
Fano factor
Foreperiod
Delay Preferred
Delay Least Preferred
Pearsonr = -0.59
pvalue = 0.1275
Mean firing rate (Hz)

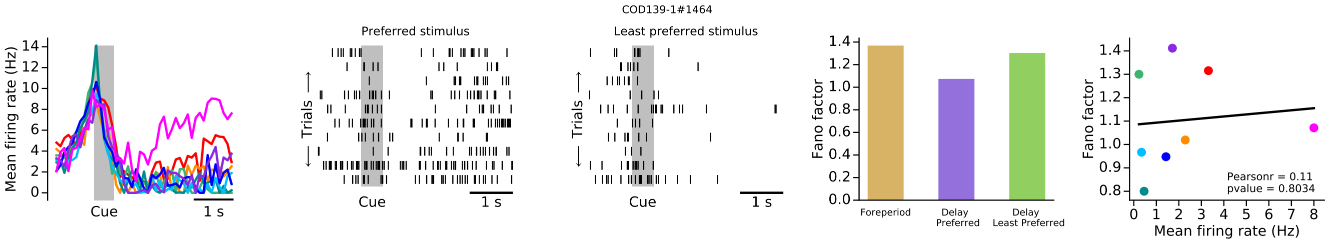
COD139-1#1464
Preferred stimulus
Least preferred stimulus
Mean firing rate (Hz)
Trials
Cue
1 s
Fano factor
Foreperiod
Delay Preferred
Delay Least Preferred
Pearsonr = 0.11
pvalue = 0.8034
Mean firing rate (Hz)

COD142-1#1477

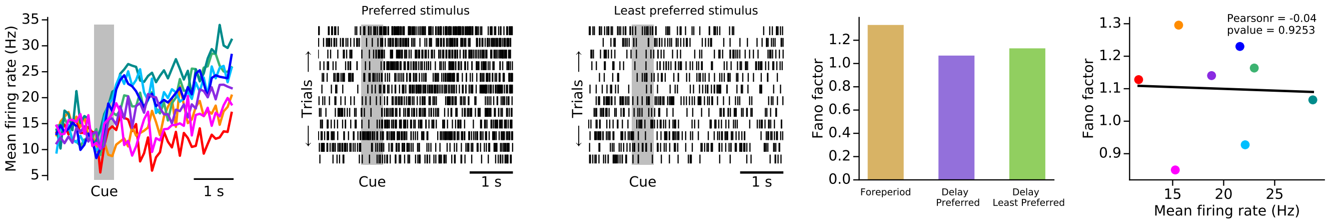


COD143-1#1481

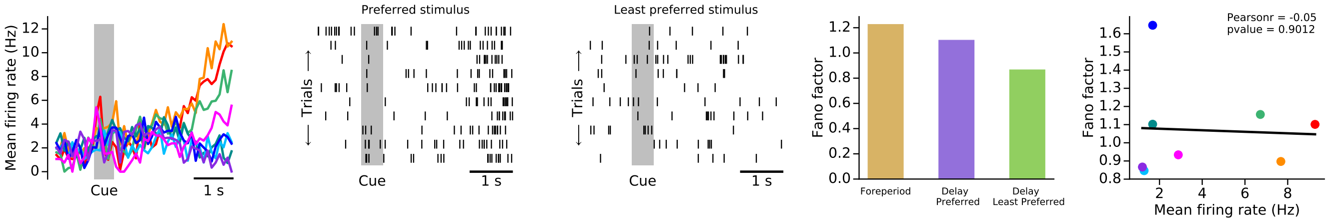


COD144-1#1486

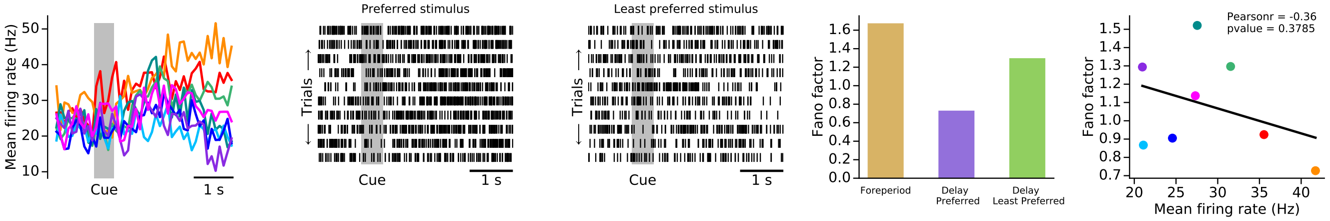


COD145-1#1493

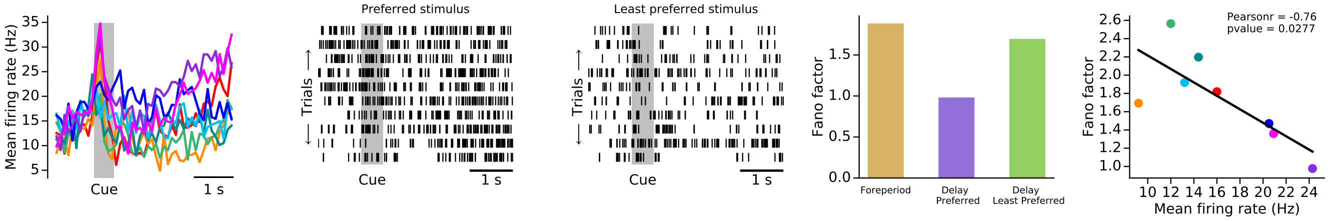


COD147-1#1505

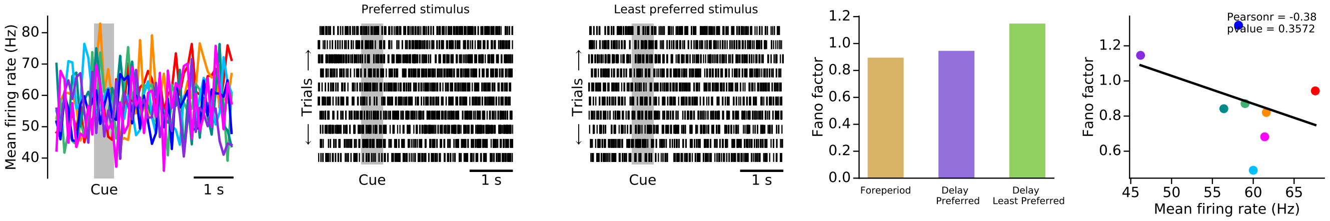

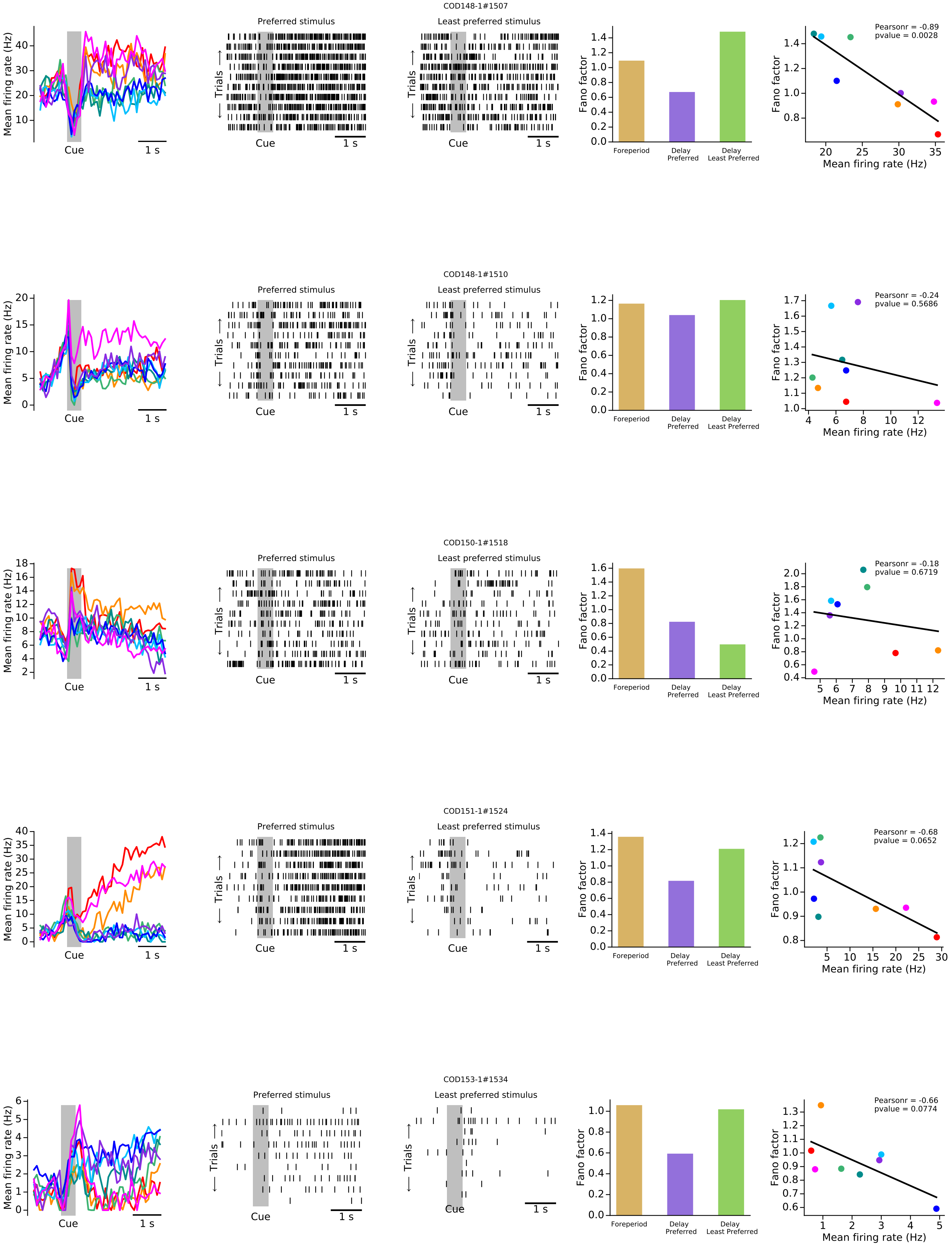
COD148-1#1507
Preferred stimulus
Least preferred stimulus
Mean firing rate (Hz)
Trials
Cue
1 s
Fano factor
Foreperiod
Delay Preferred
Delay Least Preferred
Pearsonr = -0.89
pvalue = 0.0028
Mean firing rate (Hz)
COD148-1#1510
Pearsonr = -0.24
pvalue = 0.5686
COD150-1#1518
Pearsonr = -0.18
pvalue = 0.6719
COD151-1#1524
Pearsonr = -0.68
pvalue = 0.0652
COD153-1#1534
Pearsonr = -0.66
pvalue = 0.0774

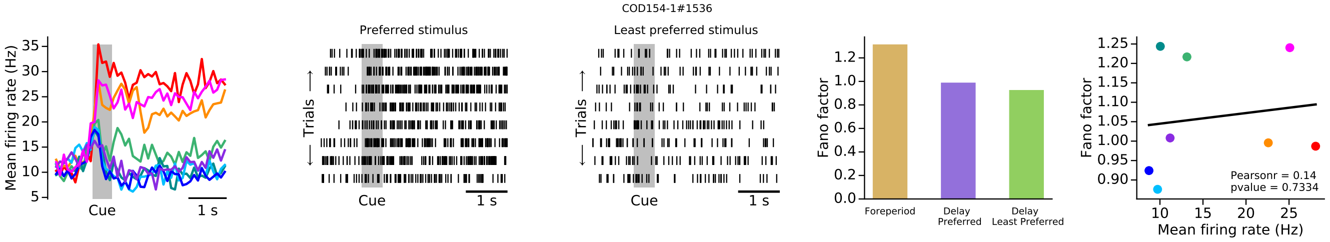
COD154-1#1536
Mean firing rate (Hz)
Cue
1 s
Preferred stimulus
Least preferred stimulus
Trials
Fano factor
Foreperiod
Delay Preferred
Delay Least Preferred
Pearsonr = 0.14
pvalue = 0.7334
Mean firing rate (Hz)

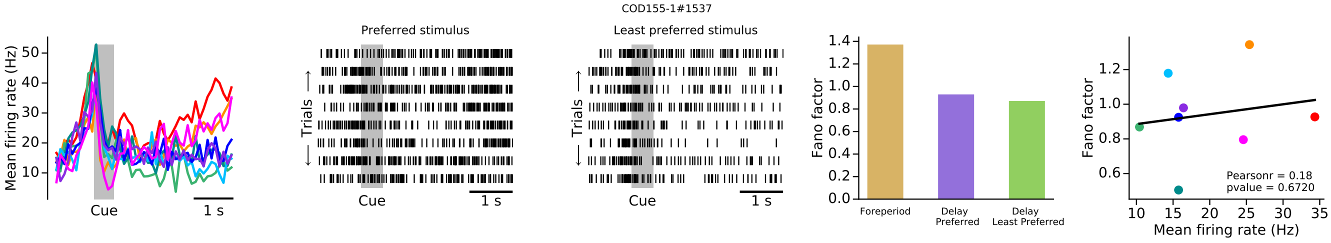
COD155-1#1537
Mean firing rate (Hz)
Cue
1 s
Preferred stimulus
Least preferred stimulus
Trials
Fano factor
Foreperiod
Delay Preferred
Delay Least Preferred
Pearsonr = 0.18
pvalue = 0.6720
Mean firing rate (Hz)

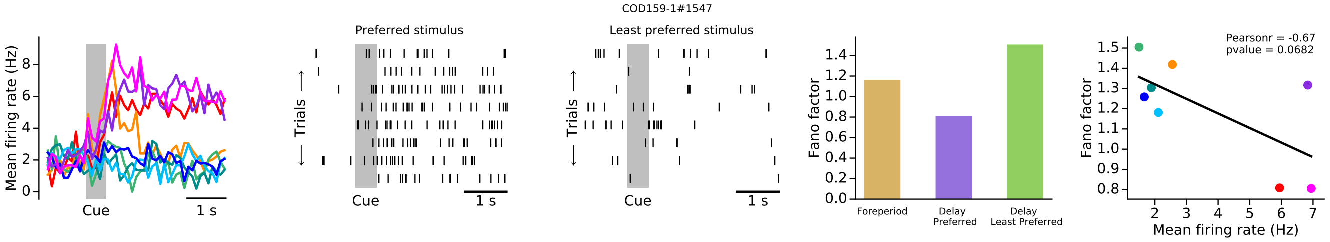
COD159-1#1547
Mean firing rate (Hz)
Cue
1 s
Preferred stimulus
Least preferred stimulus
Trials
Fano factor
Foreperiod
Delay Preferred
Delay Least Preferred
Pearsonr = -0.67
pvalue = 0.0682
Mean firing rate (Hz)

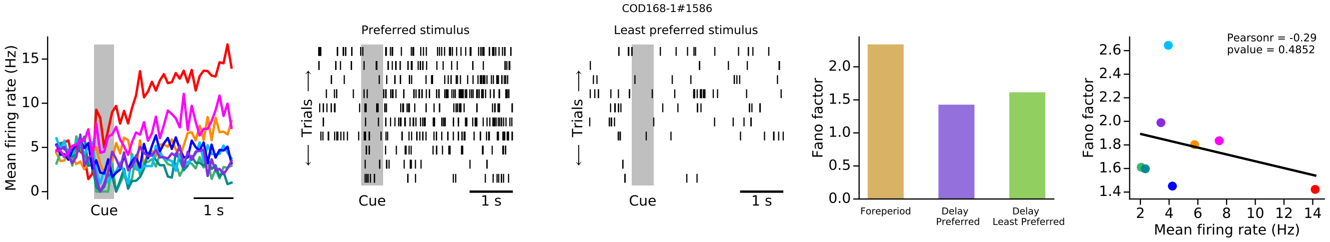
COD168-1#1586
Mean firing rate (Hz)
Cue
1 s
Preferred stimulus
Least preferred stimulus
Trials
Fano factor
Foreperiod
Delay Preferred
Delay Least Preferred
Pearsonr = -0.29
pvalue = 0.4852
Mean firing rate (Hz)

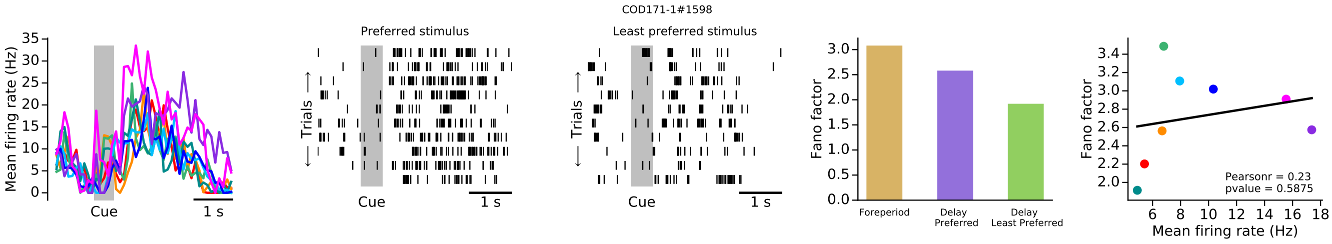
COD171-1#1598
Mean firing rate (Hz)
Cue
1 s
Preferred stimulus
Least preferred stimulus
Trials
Fano factor
Foreperiod
Delay Preferred
Delay Least Preferred
Pearsonr = 0.23
pvalue = 0.5875
Mean firing rate (Hz)

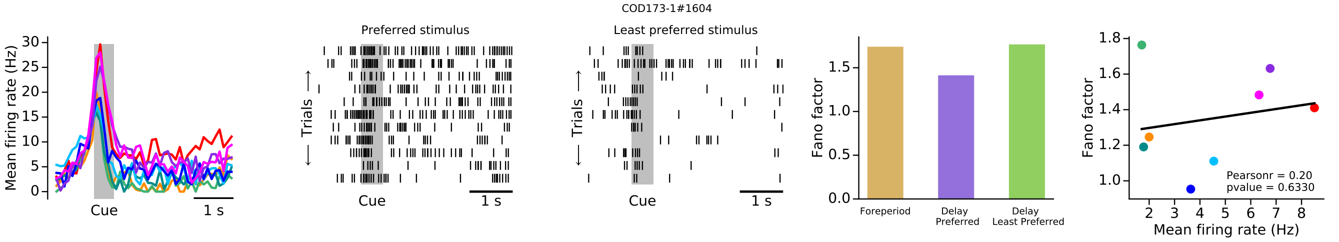
COD173-1#1604
Preferred stimulus
Least preferred stimulus
Mean firing rate (Hz)
Cue
1 s
Trials
Fano factor
Foreperiod
Delay Preferred
Delay Least Preferred
Pearsonr = 0.20
pvalue = 0.6330
Mean firing rate (Hz)

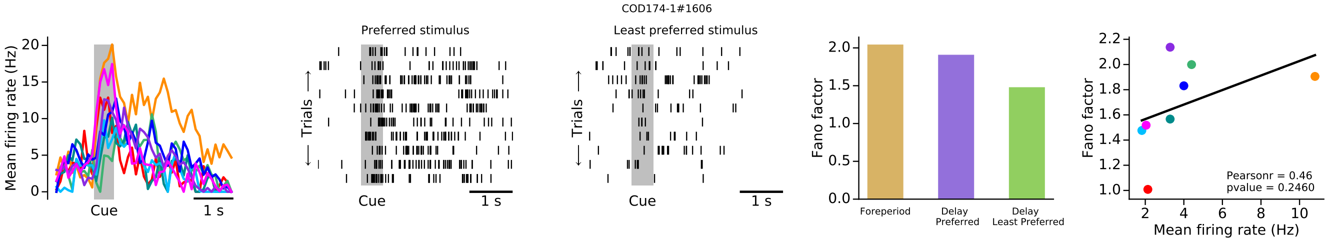
COD174-1#1606
Preferred stimulus
Least preferred stimulus
Mean firing rate (Hz)
Cue
1 s
Trials
Fano factor
Foreperiod
Delay Preferred
Delay Least Preferred
Pearsonr = 0.46
pvalue = 0.2460
Mean firing rate (Hz)

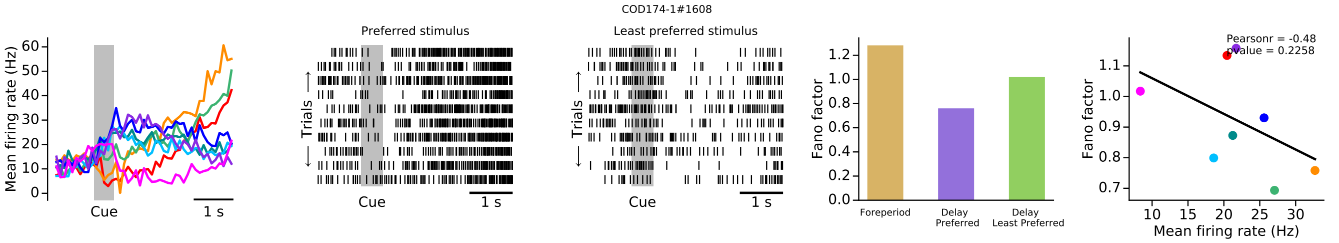
COD174-1#1608
Preferred stimulus
Least preferred stimulus
Mean firing rate (Hz)
Cue
1 s
Trials
Fano factor
Foreperiod
Delay Preferred
Delay Least Preferred
Pearsonr = -0.48
pvalue = 0.2258
Mean firing rate (Hz)

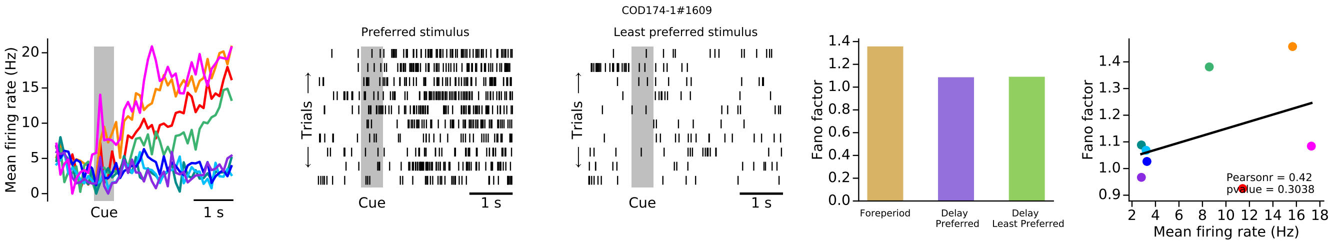
COD174-1#1609
Preferred stimulus
Least preferred stimulus
Mean firing rate (Hz)
Cue
1 s
Trials
Fano factor
Foreperiod
Delay Preferred
Delay Least Preferred
Pearsonr = 0.42
pvalue = 0.3038
Mean firing rate (Hz)

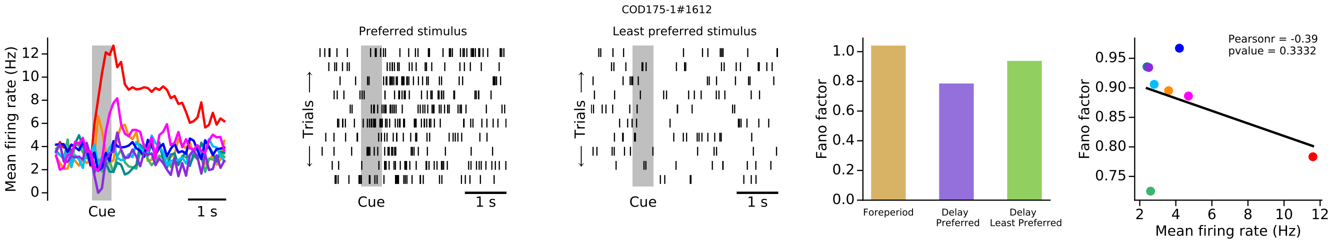
COD175-1#1612
Preferred stimulus
Least preferred stimulus
Mean firing rate (Hz)
Cue
1 s
Trials
Fano factor
Foreperiod
Delay Preferred
Delay Least Preferred
Pearsonr = -0.39
pvalue = 0.3332
Mean firing rate (Hz)

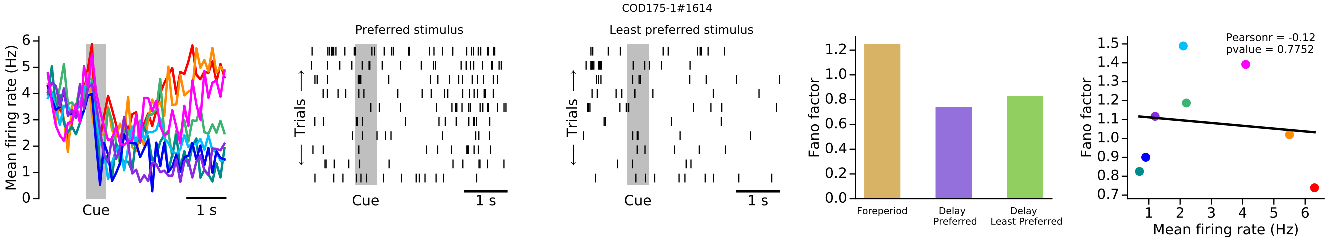
COD175-1#1614
Mean firing rate (Hz)
Cue
1 s
Preferred stimulus
Least preferred stimulus
Trials
Fano factor
Foreperiod
Delay Preferred
Delay Least Preferred
Pearsonr = -0.12
pvalue = 0.7752
Mean firing rate (Hz)

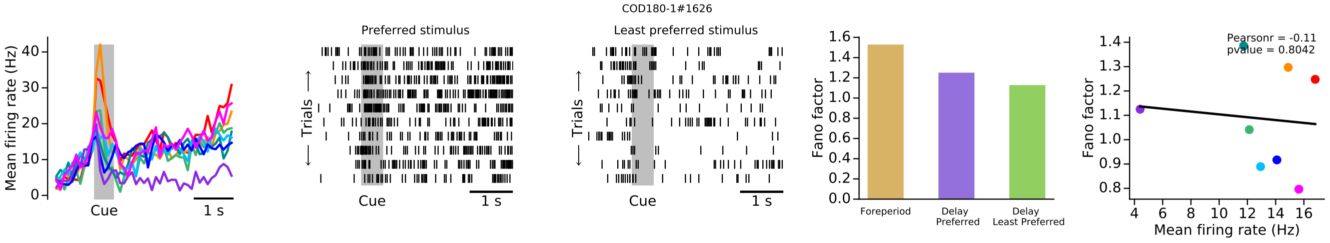
COD180-1#1626
Mean firing rate (Hz)
Cue
1 s
Preferred stimulus
Least preferred stimulus
Trials
Fano factor
Foreperiod
Delay Preferred
Delay Least Preferred
Pearsonr = -0.11
pvalue = 0.8042
Mean firing rate (Hz)

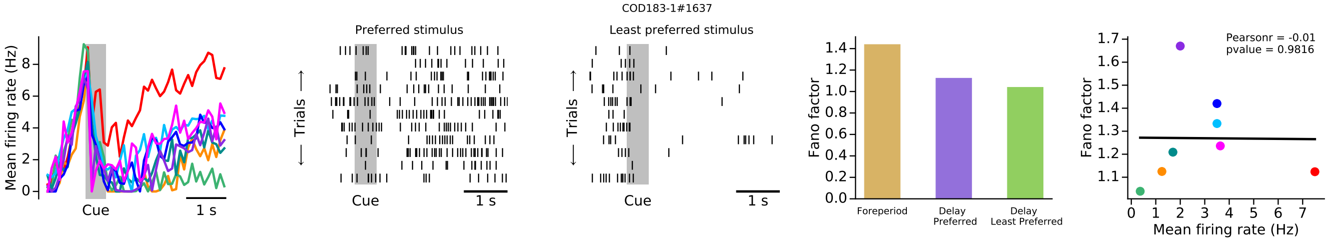
COD183-1#1637
Mean firing rate (Hz)
Cue
1 s
Preferred stimulus
Least preferred stimulus
Trials
Fano factor
Foreperiod
Delay Preferred
Delay Least Preferred
Pearsonr = -0.01
pvalue = 0.9816
Mean firing rate (Hz)

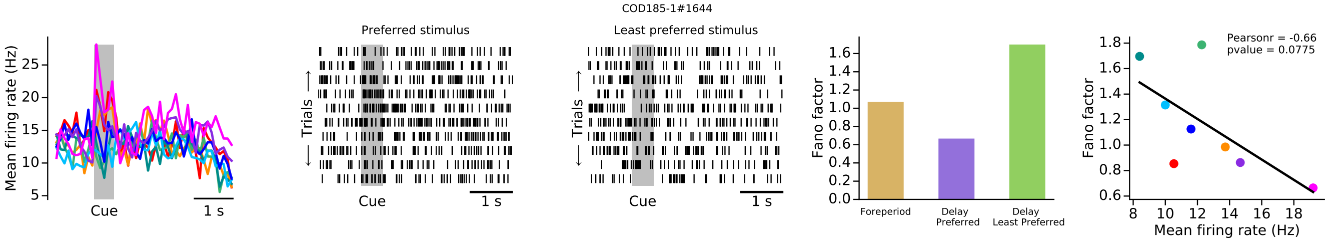
COD185-1#1644
Mean firing rate (Hz)
Cue
1 s
Preferred stimulus
Least preferred stimulus
Trials
Fano factor
Foreperiod
Delay Preferred
Delay Least Preferred
Pearsonr = -0.66
pvalue = 0.0775
Mean firing rate (Hz)

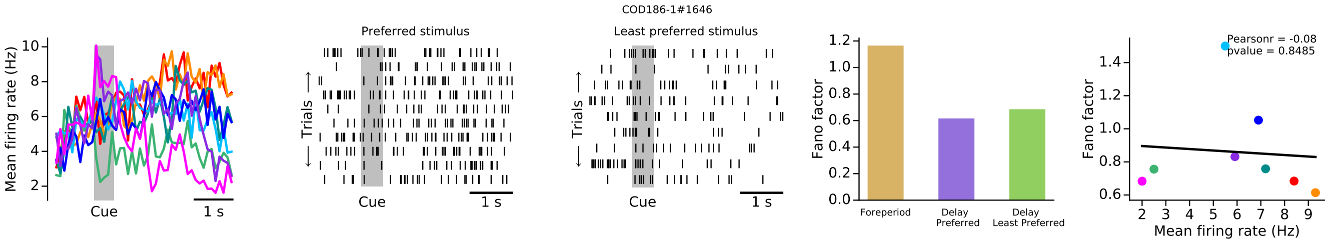
COD186-1#1646
Mean firing rate (Hz)
Cue
1 s
Preferred stimulus
Least preferred stimulus
Trials
Fano factor
Foreperiod
Delay Preferred
Delay Least Preferred
Pearsonr = -0.08
pvalue = 0.8485
Mean firing rate (Hz)

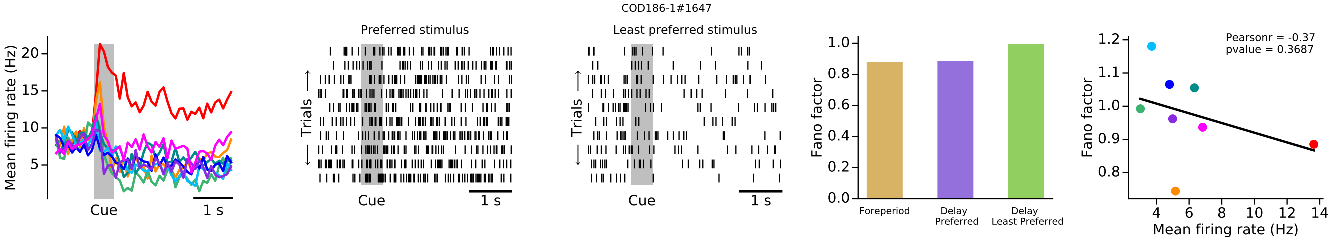
COD186-1#1647
Preferred stimulus
Least preferred stimulus
Mean firing rate (Hz)
Trials
Cue
1 s
Fano factor
Foreperiod
Delay Preferred
Delay Least Preferred
Pearsonr = -0.37
pvalue = 0.3687
Mean firing rate (Hz)

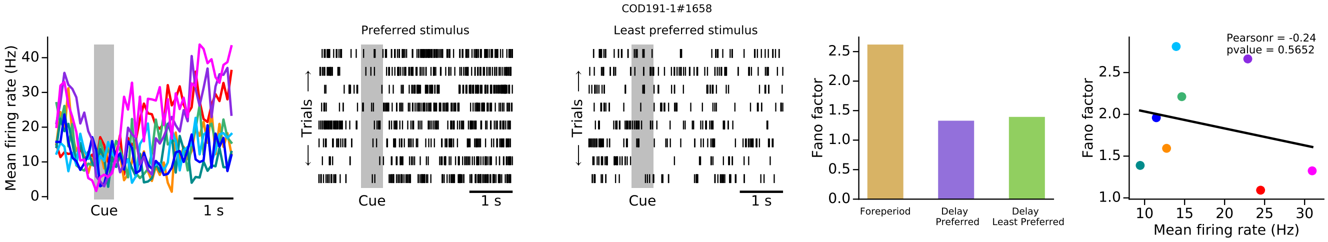
COD191-1#1658
Preferred stimulus
Least preferred stimulus
Mean firing rate (Hz)
Trials
Cue
1 s
Fano factor
Foreperiod
Delay Preferred
Delay Least Preferred
Pearsonr = -0.24
pvalue = 0.5652
Mean firing rate (Hz)

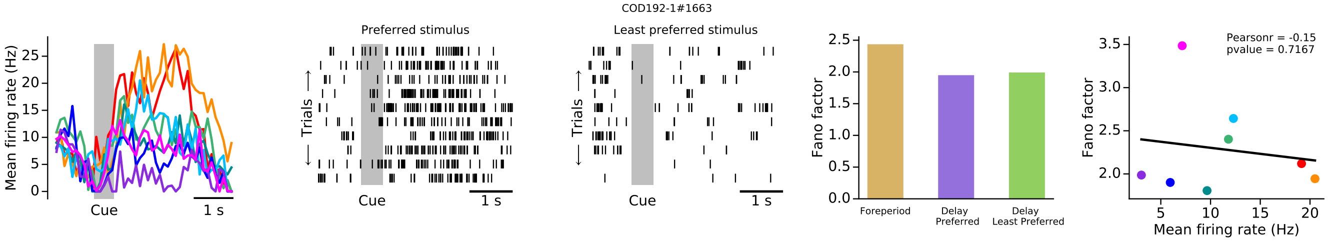
COD192-1#1663
Preferred stimulus
Least preferred stimulus
Mean firing rate (Hz)
Trials
Cue
1 s
Fano factor
Foreperiod
Delay Preferred
Delay Least Preferred
Pearsonr = -0.15
pvalue = 0.7167
Mean firing rate (Hz)

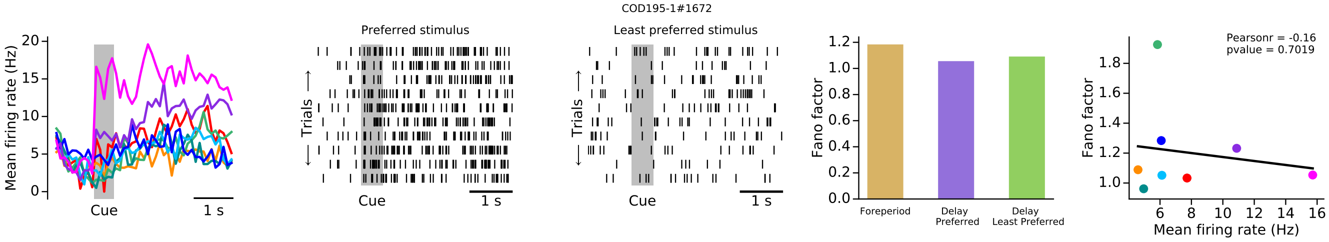
COD195-1#1672
Preferred stimulus
Least preferred stimulus
Mean firing rate (Hz)
Trials
Cue
1 s
Fano factor
Foreperiod
Delay Preferred
Delay Least Preferred
Pearsonr = -0.16
pvalue = 0.7019
Mean firing rate (Hz)

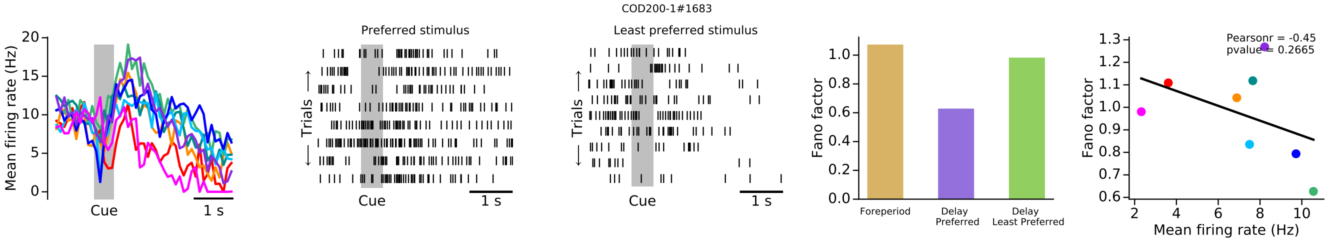
COD200-1#1683
Preferred stimulus
Least preferred stimulus
Mean firing rate (Hz)
Trials
Cue
1 s
Fano factor
Foreperiod
Delay Preferred
Delay Least Preferred
Pearsonr = -0.45
pvalue = 0.2665
Mean firing rate (Hz)

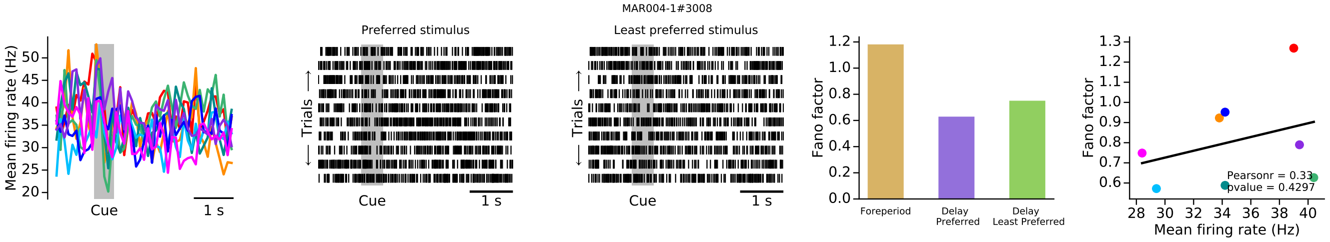
MAR004-1#3008
Preferred stimulus
Least preferred stimulus
Mean firing rate (Hz)
Cue
1 s
Trials
Fano factor
Foreperiod
Delay Preferred
Delay Least Preferred
Pearsonr = 0.33
pvalue = 0.4297
Mean firing rate (Hz)

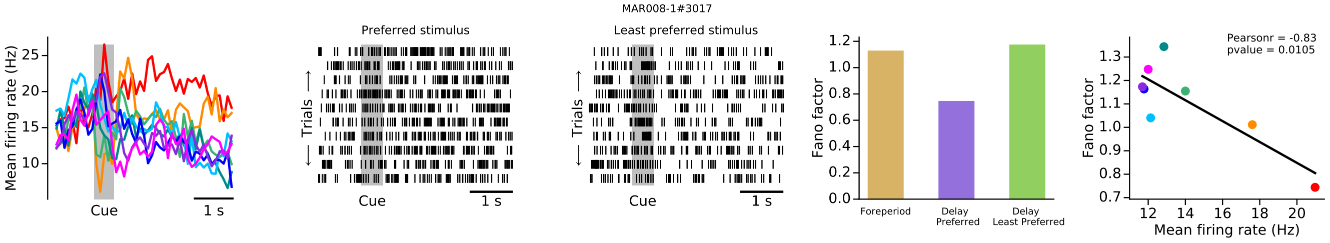
MAR008-1#3017
Preferred stimulus
Least preferred stimulus
Mean firing rate (Hz)
Cue
1 s
Trials
Fano factor
Foreperiod
Delay Preferred
Delay Least Preferred
Pearsonr = -0.83
pvalue = 0.0105
Mean firing rate (Hz)

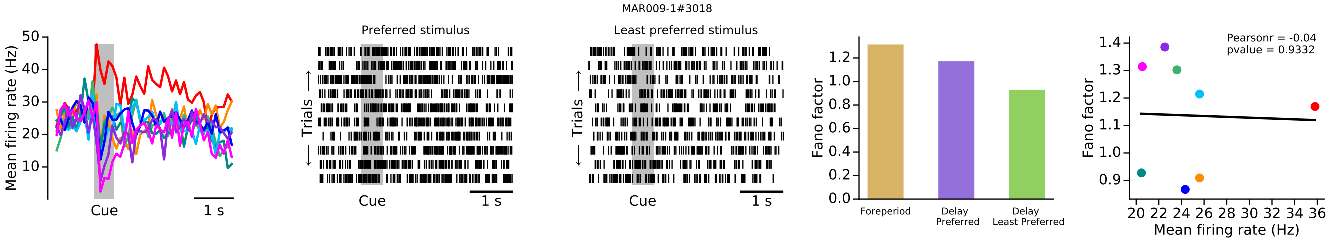
MAR009-1#3018
Preferred stimulus
Least preferred stimulus
Mean firing rate (Hz)
Cue
1 s
Trials
Fano factor
Foreperiod
Delay Preferred
Delay Least Preferred
Pearsonr = -0.04
pvalue = 0.9332
Mean firing rate (Hz)

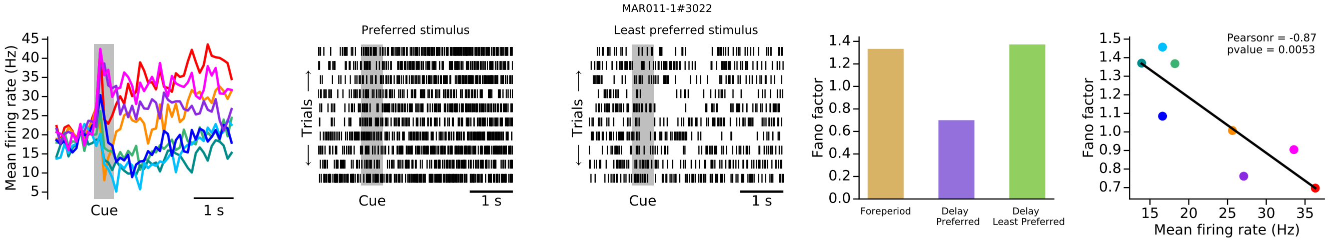
MAR011-1#3022
Preferred stimulus
Least preferred stimulus
Mean firing rate (Hz)
Cue
1 s
Trials
Fano factor
Foreperiod
Delay Preferred
Delay Least Preferred
Pearsonr = -0.87
pvalue = 0.0053
Mean firing rate (Hz)

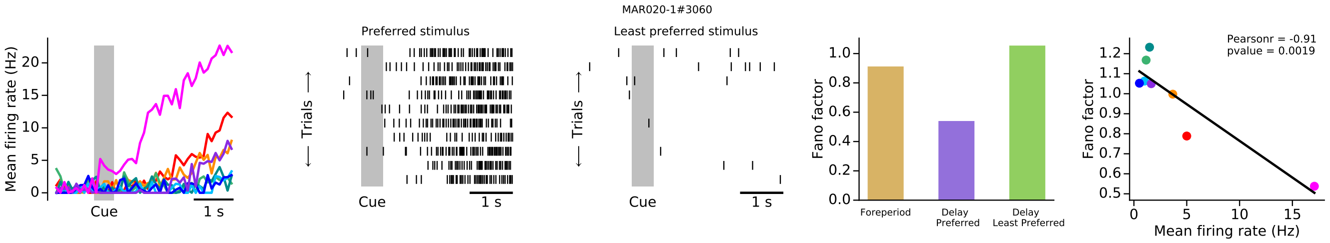
MAR020-1#3060
Preferred stimulus
Least preferred stimulus
Mean firing rate (Hz)
Cue
1 s
Trials
Fano factor
Foreperiod
Delay Preferred
Delay Least Preferred
Pearsonr = -0.91
pvalue = 0.0019
Mean firing rate (Hz)

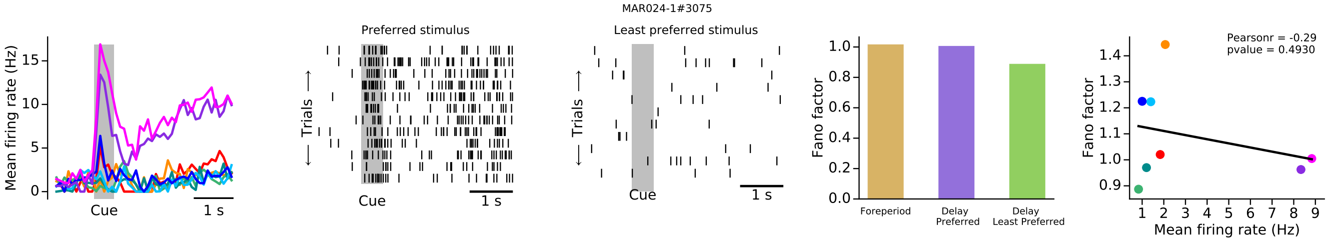
MAR024-1#3075
Mean firing rate (Hz)
Cue
1 s
Preferred stimulus
Least preferred stimulus
Trials
Fano factor
Foreperiod
Delay Preferred
Delay Least Preferred
Pearsonr = -0.29
pvalue = 0.4930
Mean firing rate (Hz)

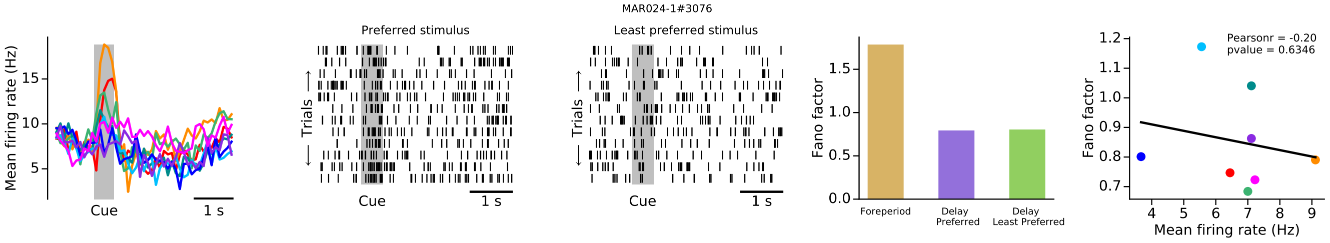
MAR024-1#3076
Mean firing rate (Hz)
Cue
1 s
Preferred stimulus
Least preferred stimulus
Trials
Fano factor
Foreperiod
Delay Preferred
Delay Least Preferred
Pearsonr = -0.20
pvalue = 0.6346
Mean firing rate (Hz)

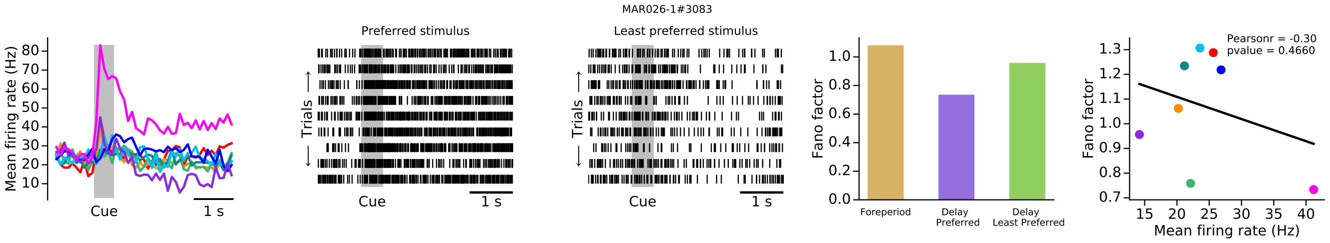
MAR026-1#3083
Mean firing rate (Hz)
Cue
1 s
Preferred stimulus
Least preferred stimulus
Trials
Fano factor
Foreperiod
Delay Preferred
Delay Least Preferred
Pearsonr = -0.30
pvalue = 0.4660
Mean firing rate (Hz)

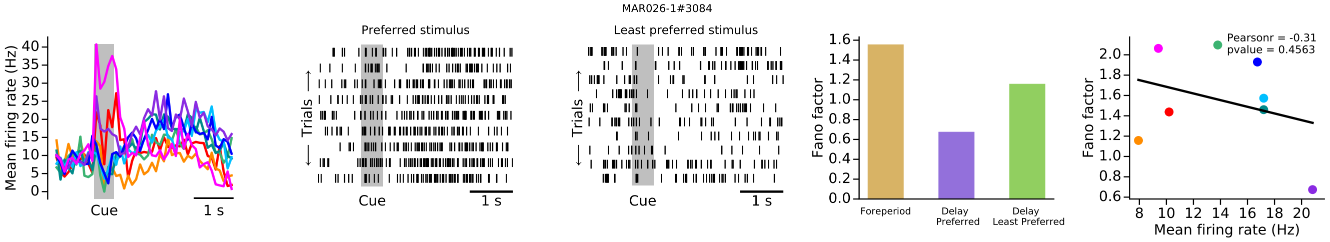
MAR026-1#3084
Mean firing rate (Hz)
Cue
1 s
Preferred stimulus
Least preferred stimulus
Trials
Fano factor
Foreperiod
Delay Preferred
Delay Least Preferred
Pearsonr = -0.31
pvalue = 0.4563
Mean firing rate (Hz)

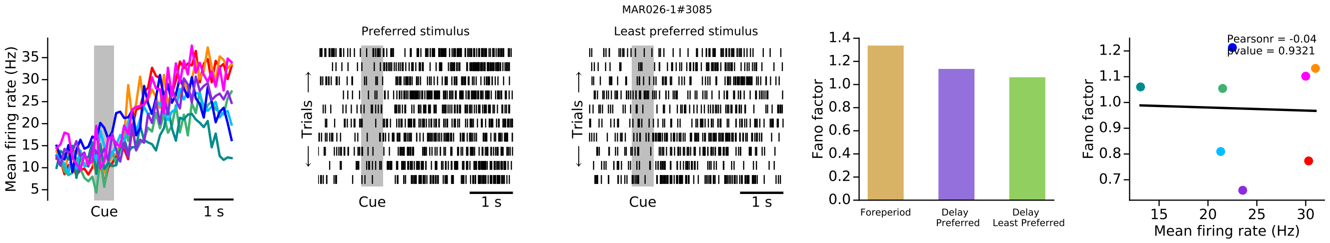
MAR026-1#3085
Mean firing rate (Hz)
Cue
1 s
Preferred stimulus
Least preferred stimulus
Trials
Fano factor
Foreperiod
Delay Preferred
Delay Least Preferred
Pearsonr = -0.04
pvalue = 0.9321
Mean firing rate (Hz)

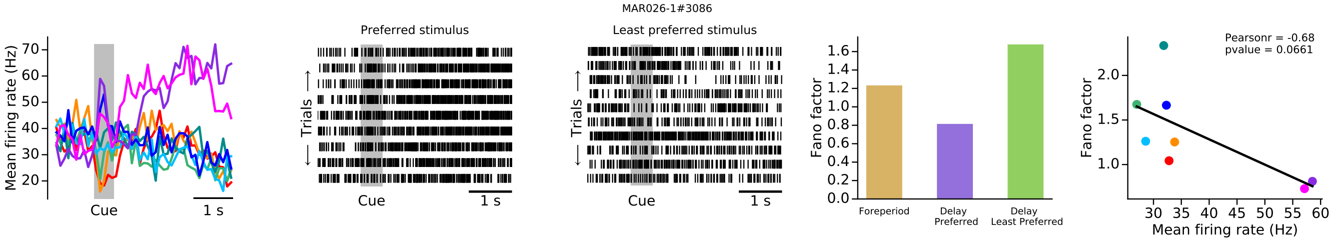
MAR026-1#3086
Preferred stimulus
Least preferred stimulus
Mean firing rate (Hz)
Trials
Cue
1 s
Fano factor
Foreperiod
Delay Preferred
Delay Least Preferred
Pearsonr = -0.68
pvalue = 0.0661
Mean firing rate (Hz)

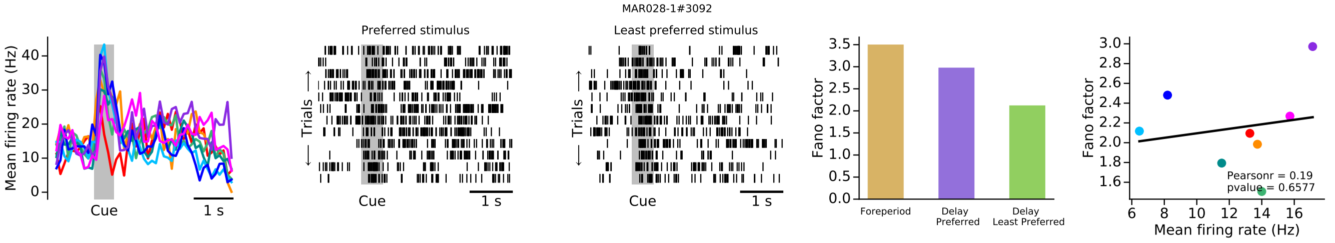
MAR028-1#3092
Preferred stimulus
Least preferred stimulus
Mean firing rate (Hz)
Trials
Cue
1 s
Fano factor
Foreperiod
Delay Preferred
Delay Least Preferred
Pearsonr = 0.19
pvalue = 0.6577
Mean firing rate (Hz)

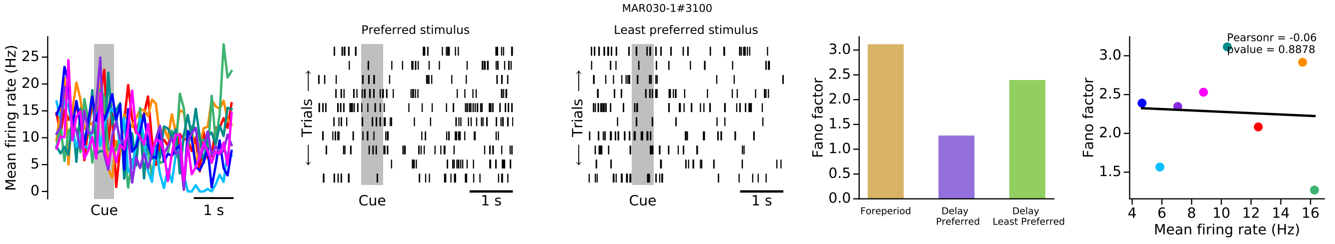
MAR030-1#3100
Preferred stimulus
Least preferred stimulus
Mean firing rate (Hz)
Trials
Cue
1 s
Fano factor
Foreperiod
Delay Preferred
Delay Least Preferred
Pearsonr = -0.06
pvalue = 0.8878
Mean firing rate (Hz)

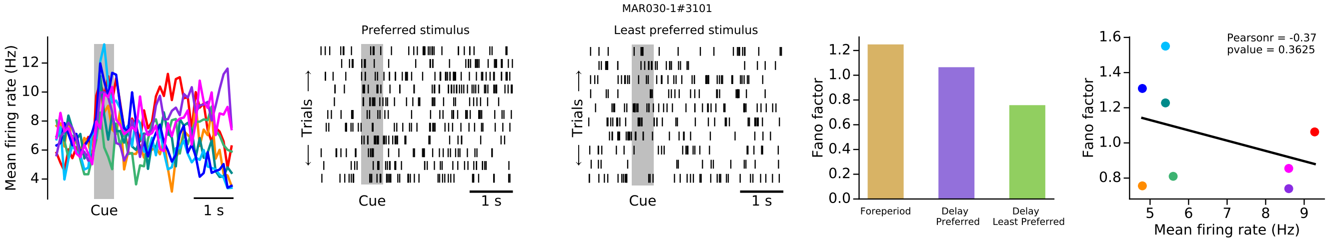
MAR030-1#3101
Preferred stimulus
Least preferred stimulus
Mean firing rate (Hz)
Trials
Cue
1 s
Fano factor
Foreperiod
Delay Preferred
Delay Least Preferred
Pearsonr = -0.37
pvalue = 0.3625
Mean firing rate (Hz)

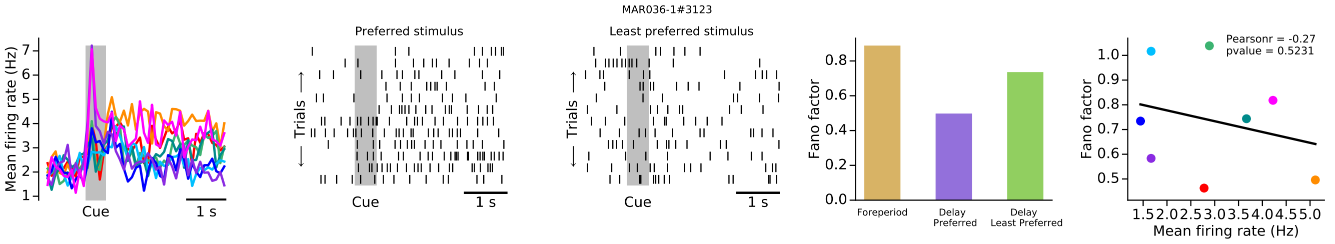
MAR036-1#3123
Preferred stimulus
Least preferred stimulus
Mean firing rate (Hz)
Trials
Cue
1 s
Fano factor
Foreperiod
Delay Preferred
Delay Least Preferred
Pearsonr = -0.27
pvalue = 0.5231
Mean firing rate (Hz)

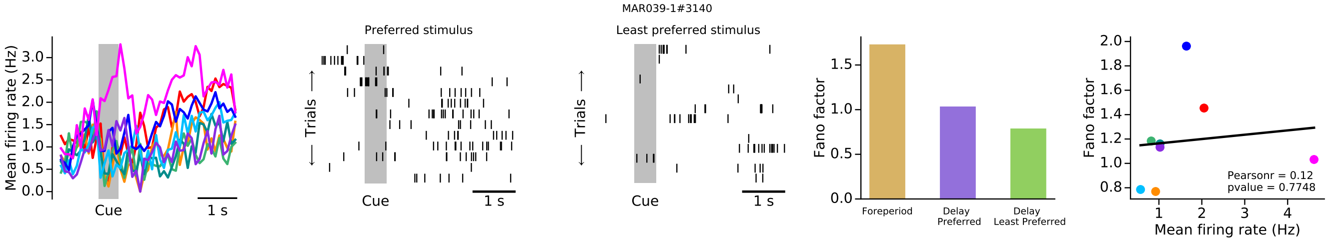
MAR039-1#3140
Mean firing rate (Hz)
Cue
1 s
Preferred stimulus
Trials
Least preferred stimulus
Fano factor
Foreperiod
Delay Preferred
Delay Least Preferred
Pearsonr = 0.12
pvalue = 0.7748
Mean firing rate (Hz)

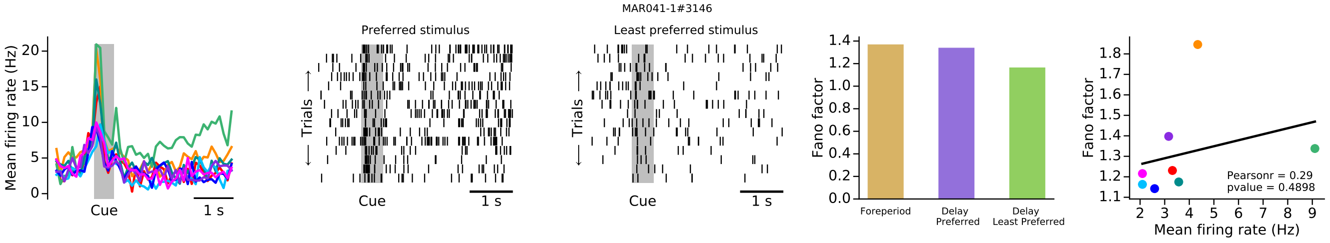
MAR041-1#3146
Mean firing rate (Hz)
Cue
1 s
Preferred stimulus
Trials
Least preferred stimulus
Fano factor
Foreperiod
Delay Preferred
Delay Least Preferred
Pearsonr = 0.29
pvalue = 0.4898
Mean firing rate (Hz)

### VDD NEURONS

#### Group 1

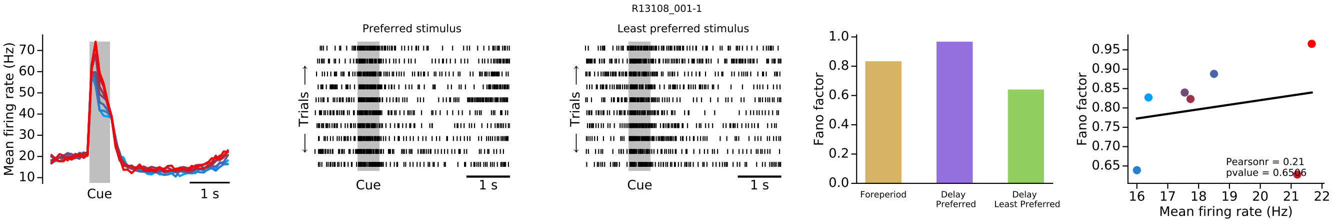


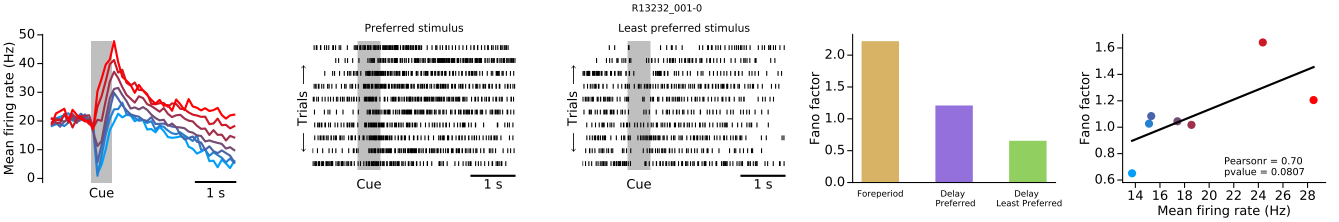


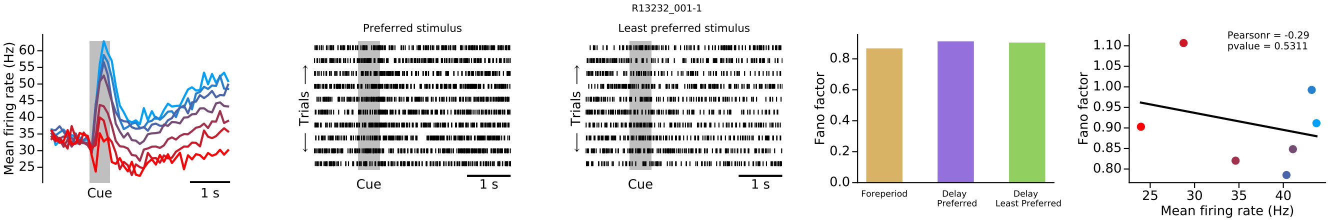


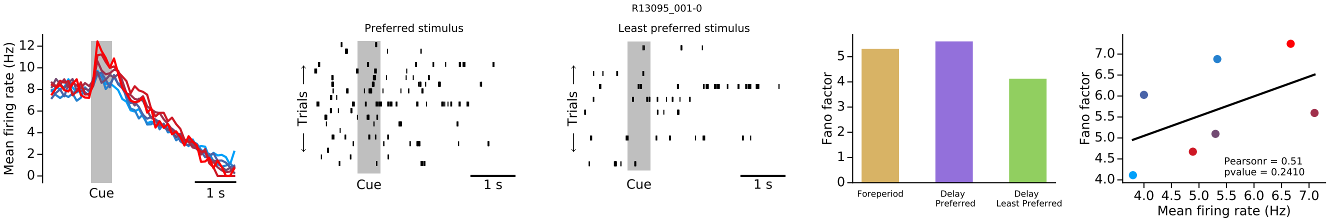


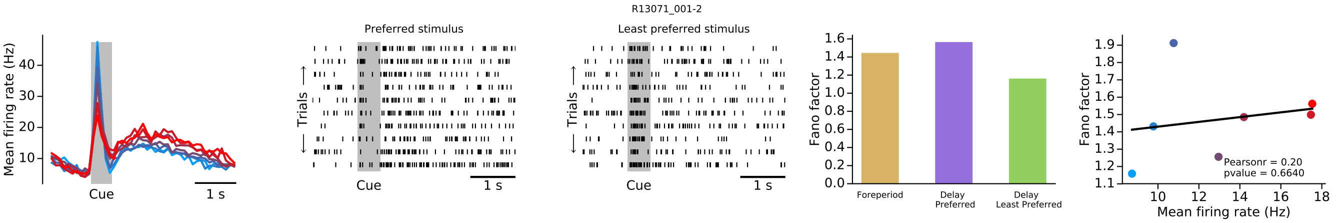

R13214_001-2

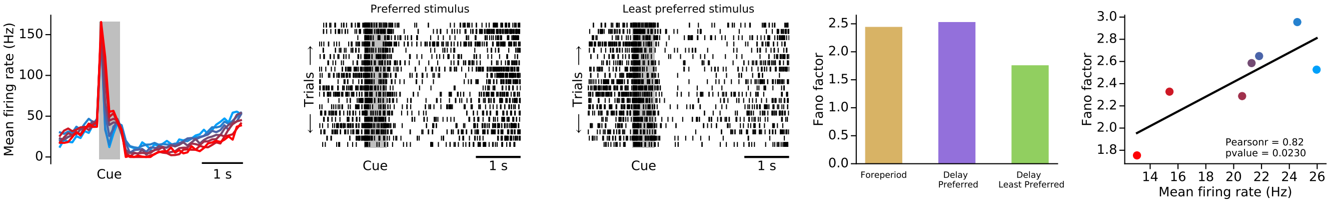


R13037_001-0

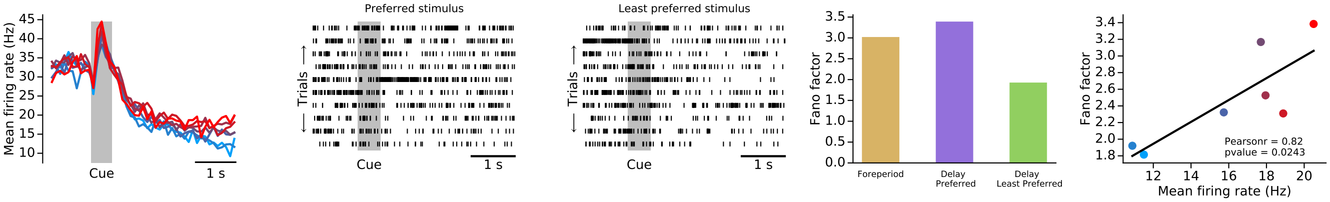


R13102_001-6

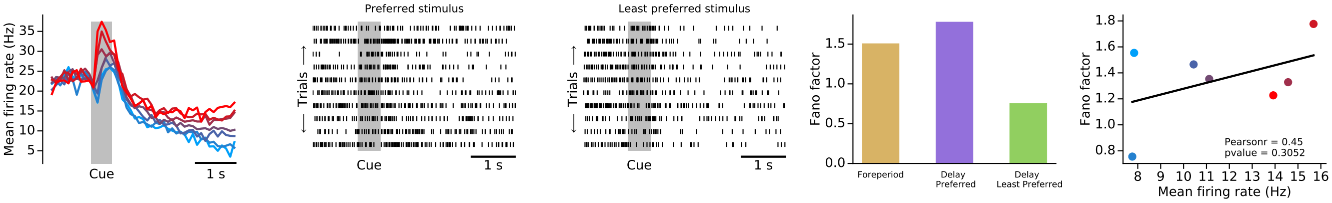


R13018_001-2

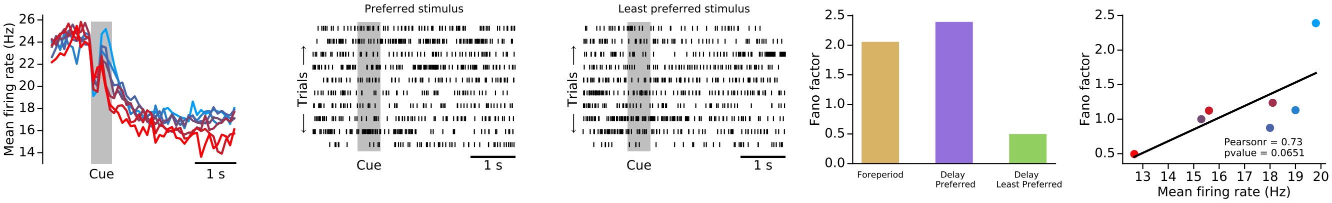


R13217_001-5

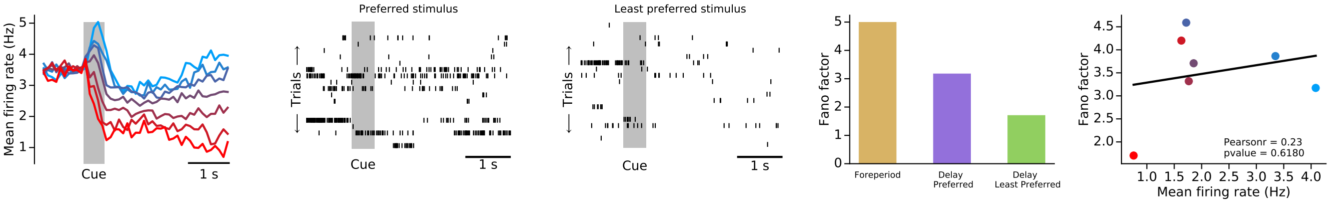

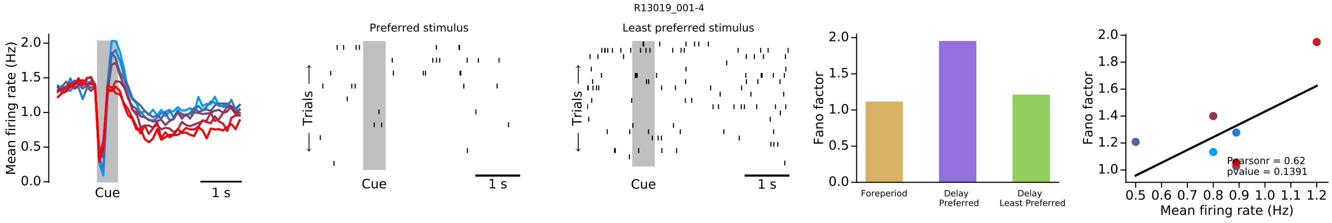
R13019_001-4
Preferred stimulus
Least preferred stimulus
Mean firing rate (Hz)
Trials
Cue
1 s
Fano factor
Foreperiod
Delay Preferred
Delay Least Preferred
Mean firing rate (Hz)
Pearsonr = 0.62
pvalue = 0.1391

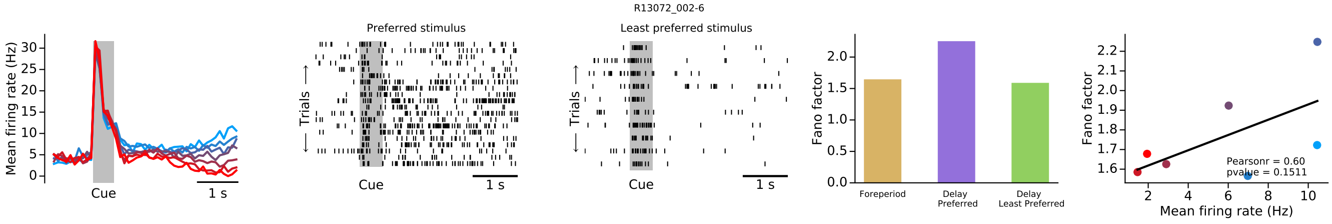
R13072_002-6
Preferred stimulus
Least preferred stimulus
Mean firing rate (Hz)
Trials
Cue
1 s
Fano factor
Foreperiod
Delay Preferred
Delay Least Preferred
Mean firing rate (Hz)
Pearsonr = 0.60
pvalue = 0.1511

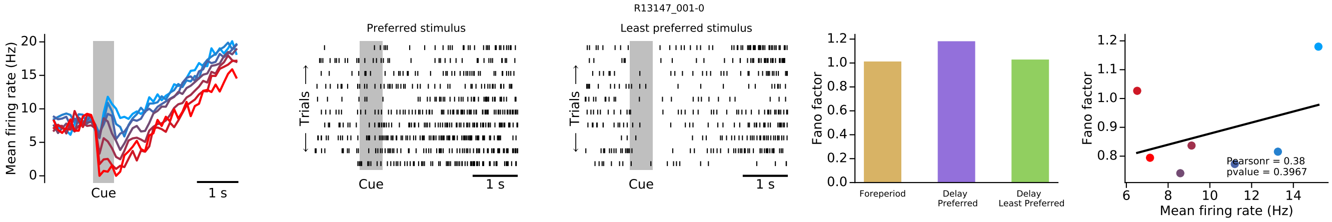
R13147_001-0
Preferred stimulus
Least preferred stimulus
Mean firing rate (Hz)
Trials
Cue
1 s
Fano factor
Foreperiod
Delay Preferred
Delay Least Preferred
Mean firing rate (Hz)
Pearsonr = 0.38
pvalue = 0.3967

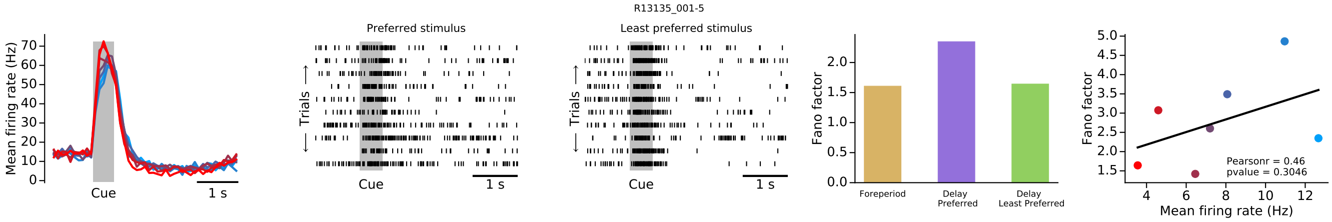
R13135_001-5
Preferred stimulus
Least preferred stimulus
Mean firing rate (Hz)
Trials
Cue
1 s
Fano factor
Foreperiod
Delay Preferred
Delay Least Preferred
Mean firing rate (Hz)
Pearsonr = 0.46
pvalue = 0.3046

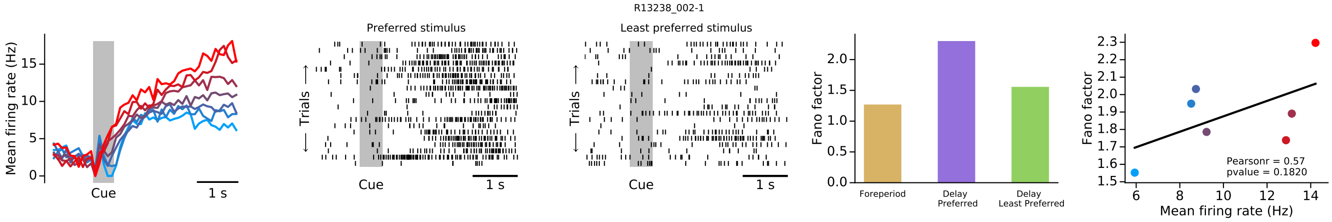
R13238_002-1
Preferred stimulus
Least preferred stimulus
Mean firing rate (Hz)
Trials
Cue
1 s
Fano factor
Foreperiod
Delay Preferred
Delay Least Preferred
Mean firing rate (Hz)
Pearsonr = 0.57
pvalue = 0.1820

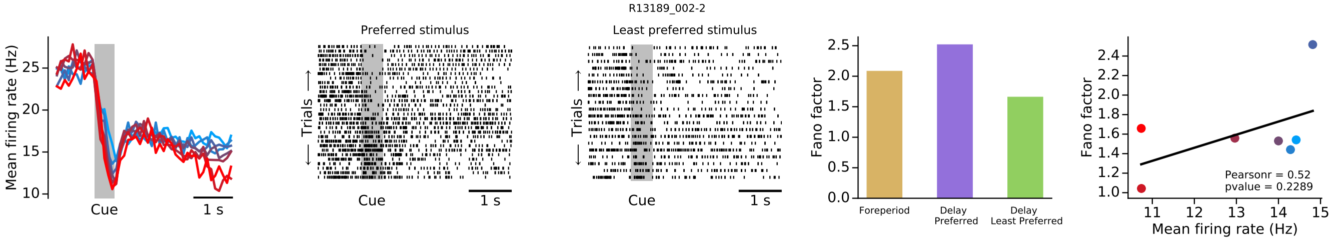
R13189_002-2
Mean firing rate (Hz)
Cue
1 s
Preferred stimulus
Least preferred stimulus
Trials
Fano factor
Foreperiod
Delay Preferred
Delay Least Preferred
Pearsonr = 0.52
pvalue = 0.2289
Mean firing rate (Hz)

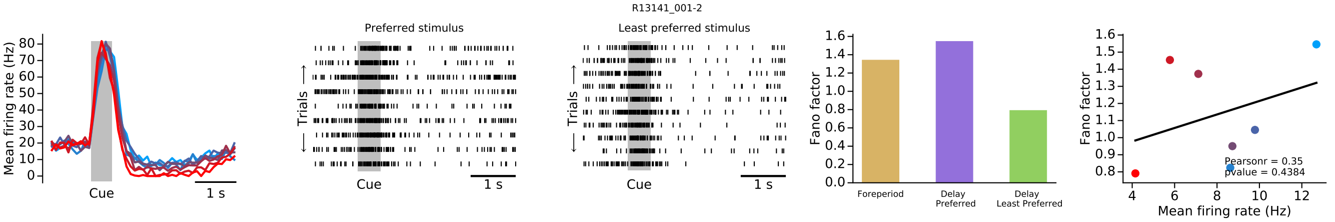
R13141_001-2
Mean firing rate (Hz)
Cue
1 s
Preferred stimulus
Least preferred stimulus
Trials
Fano factor
Foreperiod
Delay Preferred
Delay Least Preferred
Pearsonr = 0.35
pvalue = 0.4384
Mean firing rate (Hz)

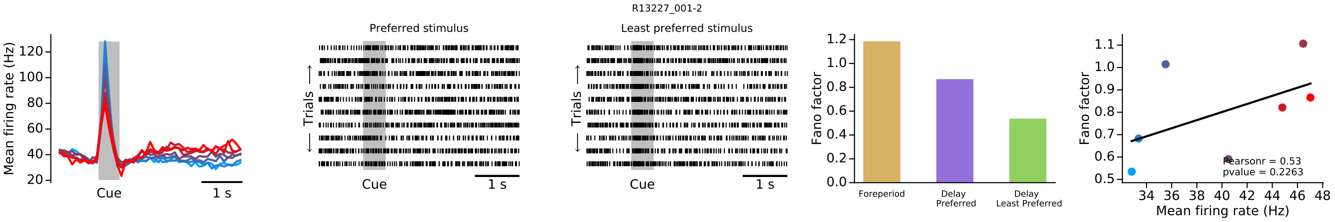
R13227_001-2
Mean firing rate (Hz)
Cue
1 s
Preferred stimulus
Least preferred stimulus
Trials
Fano factor
Foreperiod
Delay Preferred
Delay Least Preferred
Pearsonr = 0.53
pvalue = 0.2263
Mean firing rate (Hz)

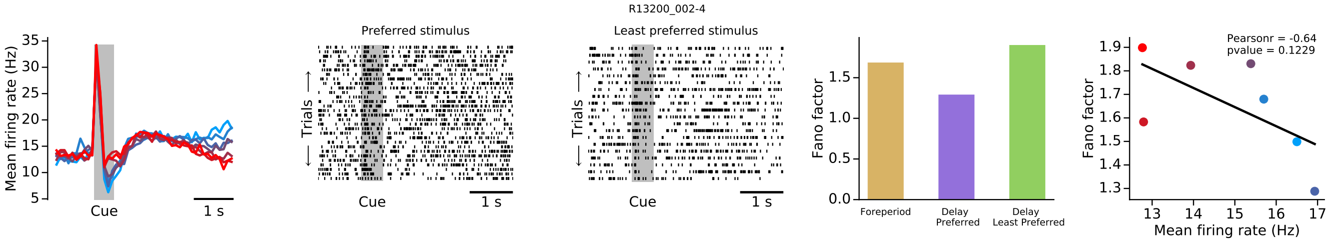
R13200_002-4
Mean firing rate (Hz)
Cue
1 s
Preferred stimulus
Least preferred stimulus
Trials
Fano factor
Foreperiod
Delay Preferred
Delay Least Preferred
Pearsonr = -0.64
pvalue = 0.1229
Mean firing rate (Hz)

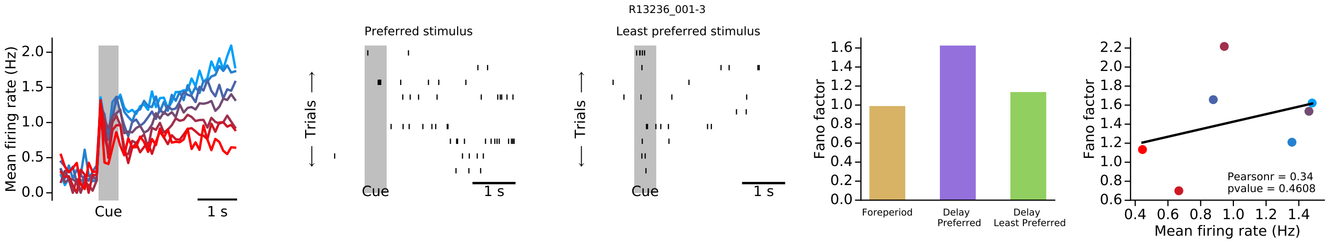
R13236_001-3
Mean firing rate (Hz)
Cue
1 s
Preferred stimulus
Least preferred stimulus
Trials
Fano factor
Foreperiod
Delay Preferred
Delay Least Preferred
Pearsonr = 0.34
pvalue = 0.4608
Mean firing rate (Hz)

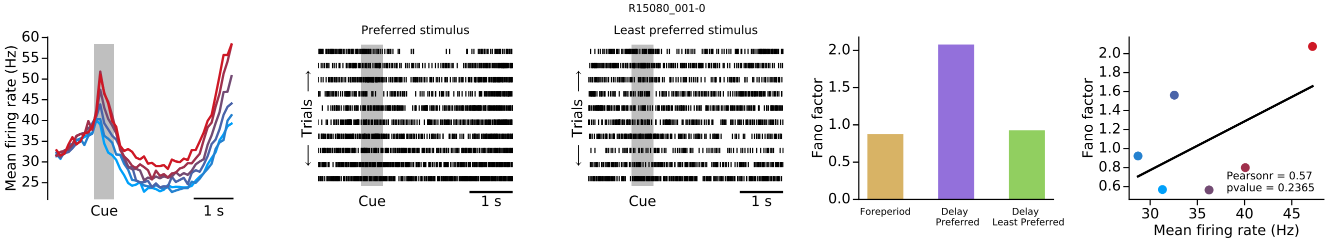
R15080_001-0
Mean firing rate (Hz)
Cue
1 s
Preferred stimulus
Least preferred stimulus
Trials
Fano factor
Foreperiod
Delay Preferred
Delay Least Preferred
Pearsonr = 0.57
pvalue = 0.2365
Mean firing rate (Hz)

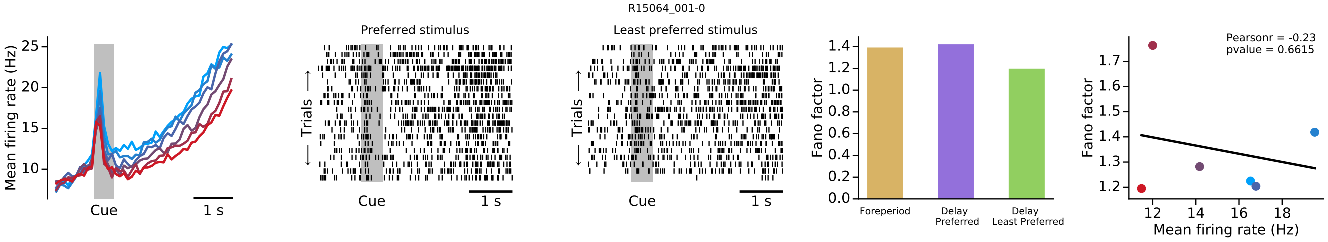
R15064_001-0
Mean firing rate (Hz)
Cue
1 s
Preferred stimulus
Least preferred stimulus
Trials
Fano factor
Foreperiod
Delay Preferred
Delay Least Preferred
Pearsonr = -0.23
pvalue = 0.6615
Mean firing rate (Hz)

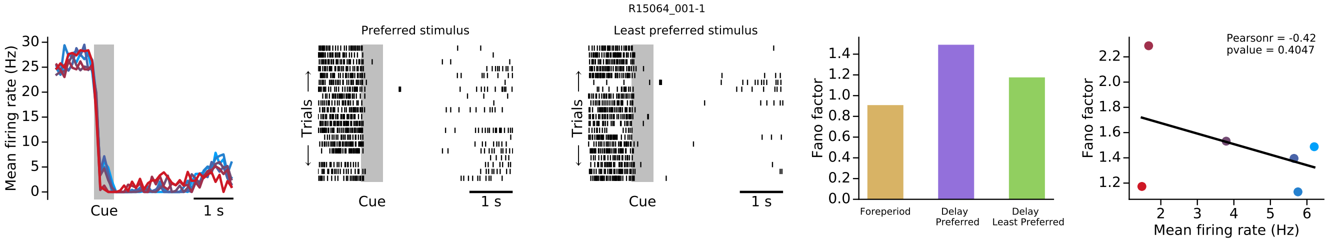
R15064_001-1
Mean firing rate (Hz)
Cue
1 s
Preferred stimulus
Least preferred stimulus
Trials
Fano factor
Foreperiod
Delay Preferred
Delay Least Preferred
Pearsonr = -0.42
pvalue = 0.4047
Mean firing rate (Hz)

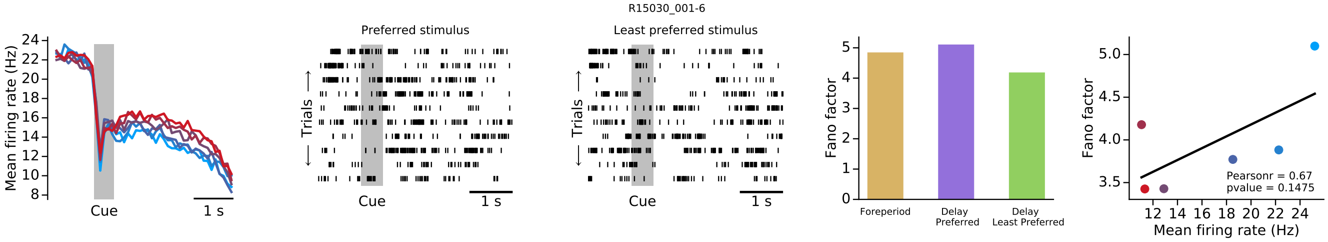
R15030_001-6
Mean firing rate (Hz)
Cue
1 s
Preferred stimulus
Least preferred stimulus
Trials
Fano factor
Foreperiod
Delay Preferred
Delay Least Preferred
Pearsonr = 0.67
pvalue = 0.1475
Mean firing rate (Hz)

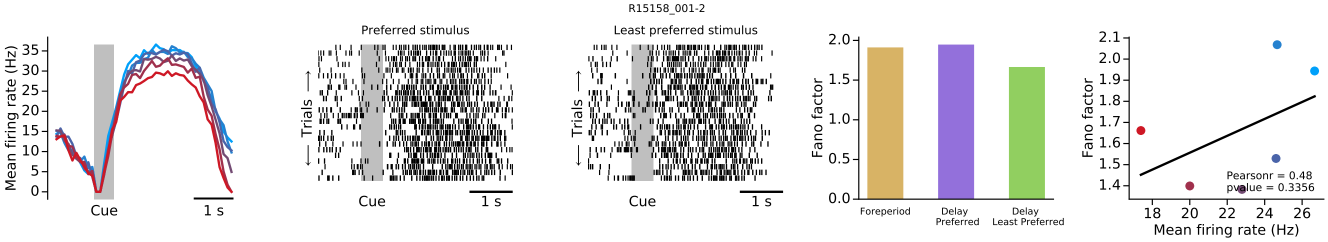
R15158_001-2
Mean firing rate (Hz)
Cue
1 s
Preferred stimulus
Least preferred stimulus
Trials
Fano factor
Foreperiod
Delay Preferred
Delay Least Preferred
Pearsonr = 0.48
pvalue = 0.3356
Mean firing rate (Hz)

R15042_001-0

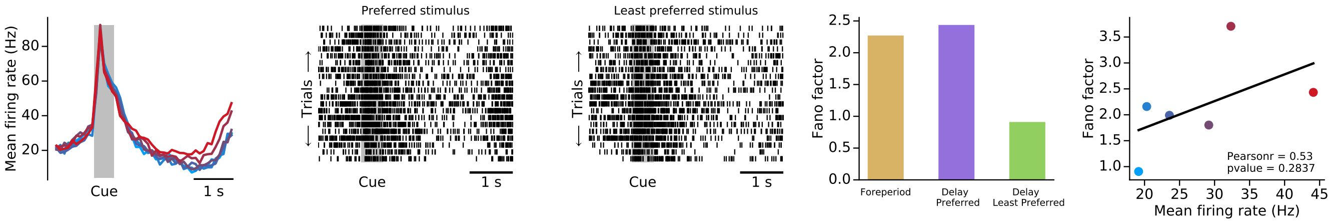


R15132_001-2

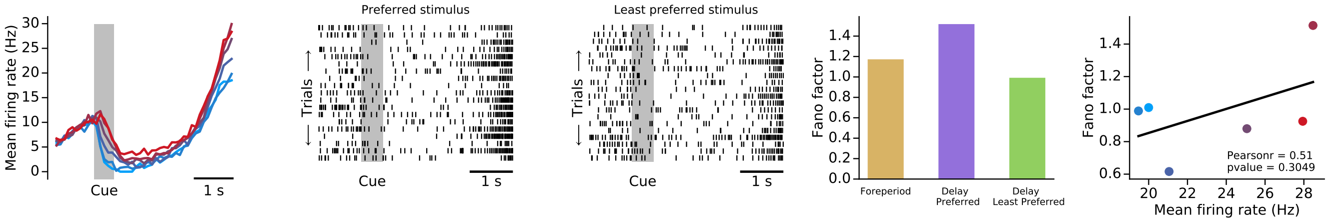


R15007_001-4

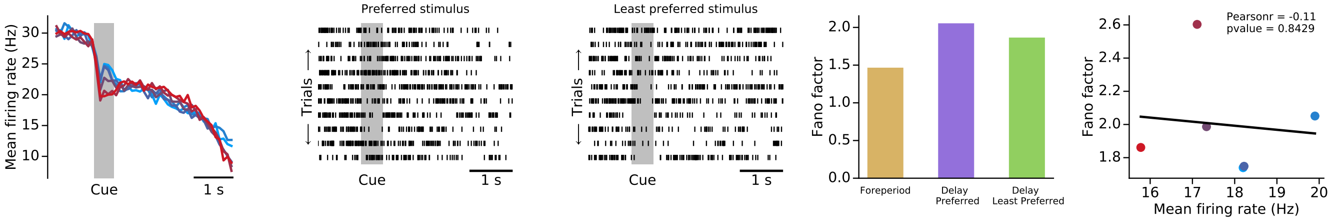


R15033_001-0

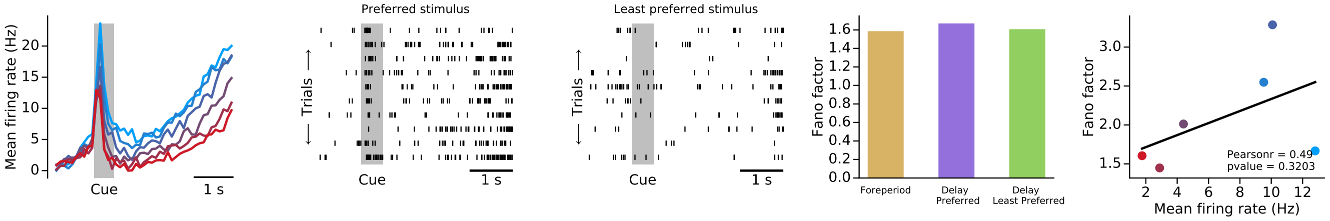


R15058_001-1

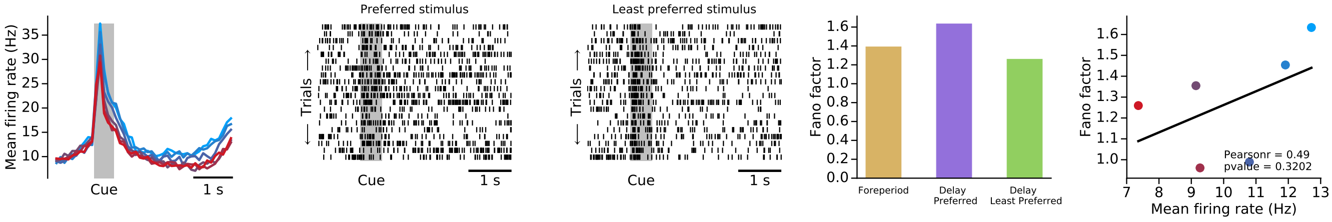

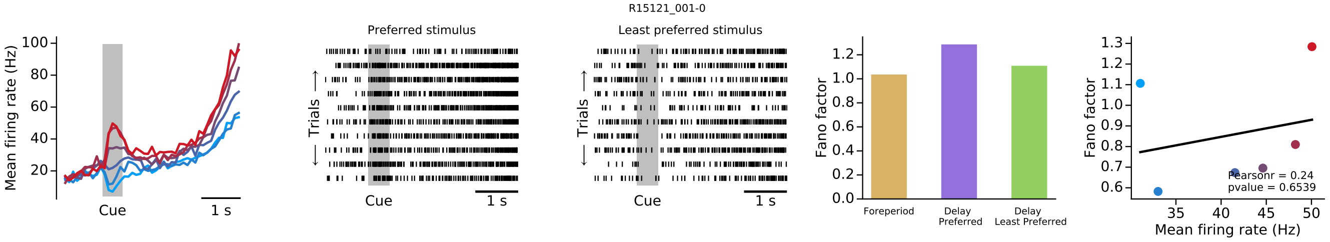
R15121_001-0
Mean firing rate (Hz)
Cue
1 s
Preferred stimulus
Least preferred stimulus
Trials
Fano factor
Foreperiod
Delay Preferred
Delay Least Preferred
Pearsonr = 0.24
pvalue = 0.6539
Mean firing rate (Hz)

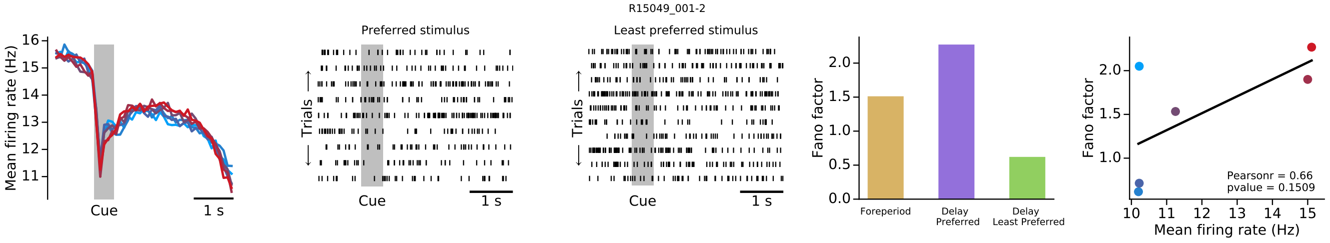
R15049_001-2
Mean firing rate (Hz)
Cue
1 s
Preferred stimulus
Least preferred stimulus
Trials
Fano factor
Foreperiod
Delay Preferred
Delay Least Preferred
Pearsonr = 0.66
pvalue = 0.1509
Mean firing rate (Hz)

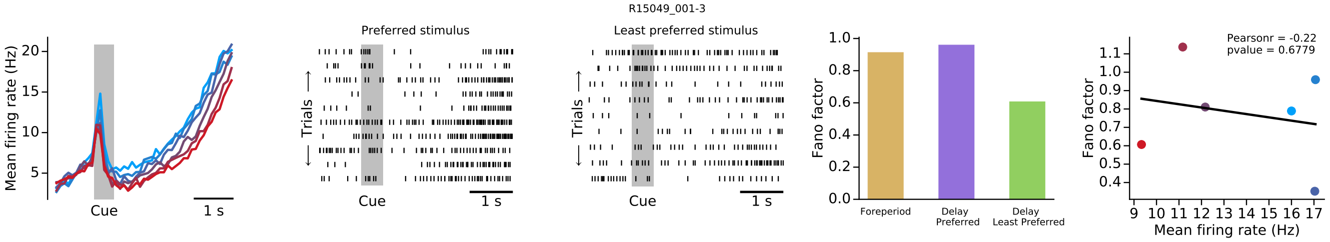
R15049_001-3
Mean firing rate (Hz)
Cue
1 s
Preferred stimulus
Least preferred stimulus
Trials
Fano factor
Foreperiod
Delay Preferred
Delay Least Preferred
Pearsonr = -0.22
pvalue = 0.6779
Mean firing rate (Hz)

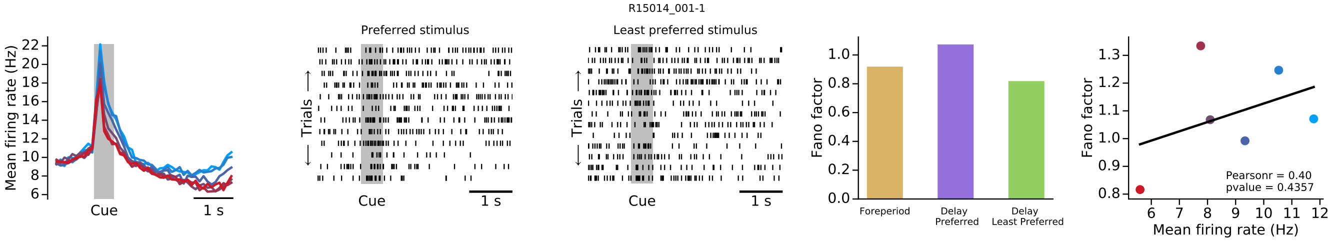
R15014_001-1
Mean firing rate (Hz)
Cue
1 s
Preferred stimulus
Least preferred stimulus
Trials
Fano factor
Foreperiod
Delay Preferred
Delay Least Preferred
Pearsonr = 0.40
pvalue = 0.4357
Mean firing rate (Hz)

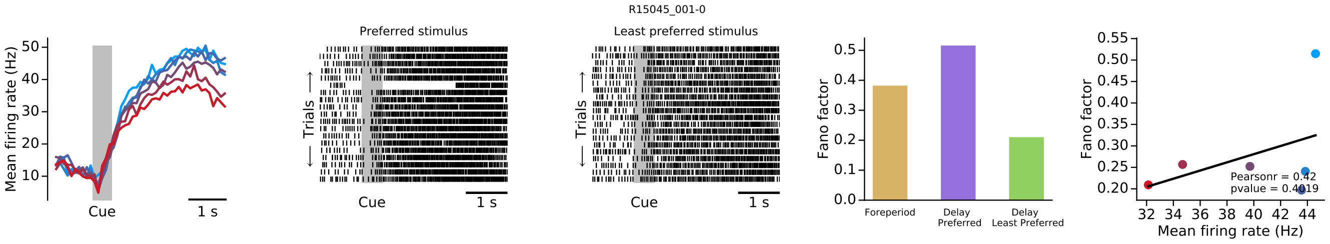
R15045_001-0
Mean firing rate (Hz)
Cue
1 s
Preferred stimulus
Least preferred stimulus
Trials
Fano factor
Foreperiod
Delay Preferred
Delay Least Preferred
Pearsonr = 0.42
Mean firing rate (Hz)

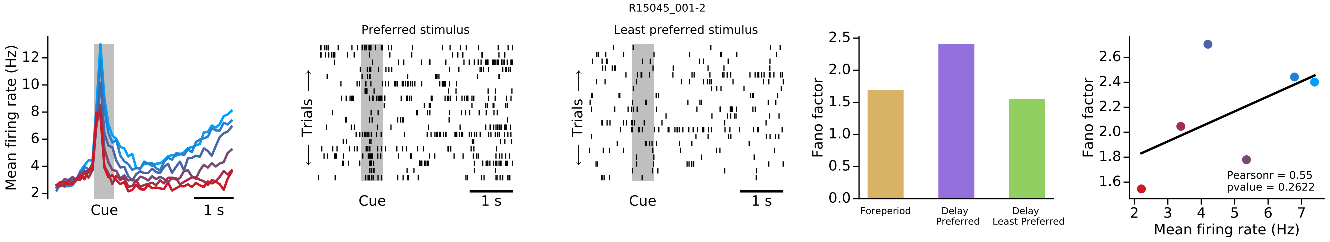
R15045_001-2
Mean firing rate (Hz)
Cue
1 s
Preferred stimulus
Least preferred stimulus
Trials
Fano factor
Foreperiod
Delay Preferred
Delay Least Preferred
Pearsonr = 0.55
pvalue = 0.2622
Mean firing rate (Hz)

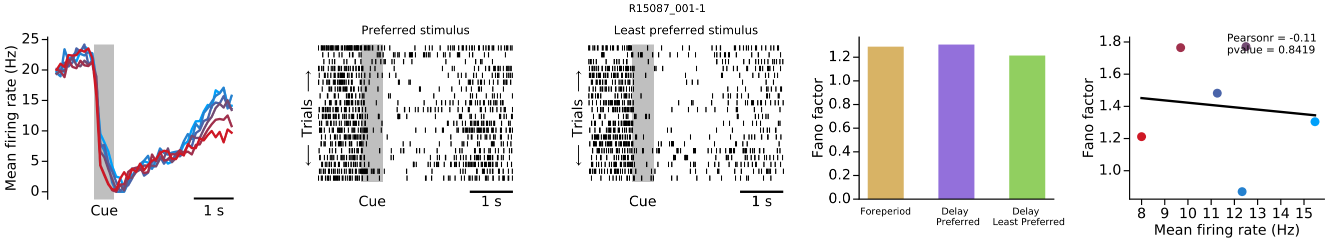
R15087_001-1
Mean firing rate (Hz)
Cue
1 s
Preferred stimulus
Least preferred stimulus
Trials
Fano factor
Foreperiod
Delay Preferred
Delay Least Preferred
Pearsonr = -0.11
pvalue = 0.8419
Mean firing rate (Hz)

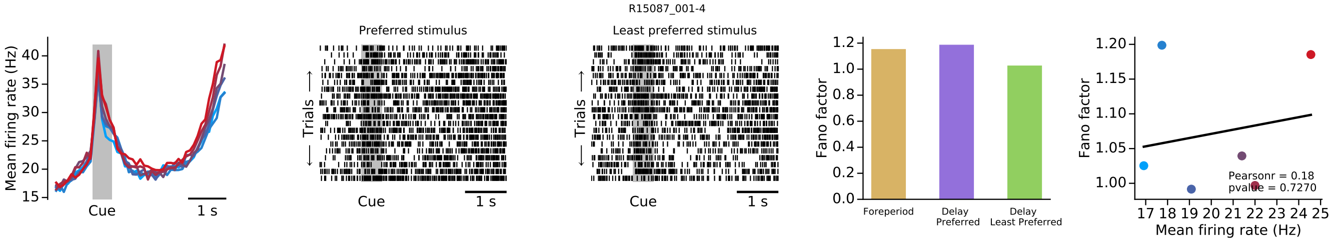
R15087_001-4
Mean firing rate (Hz)
Cue
1 s
Preferred stimulus
Least preferred stimulus
Trials
Fano factor
Foreperiod
Delay Preferred
Delay Least Preferred
Pearsonr = 0.18
pvalue = 0.7270
Mean firing rate (Hz)

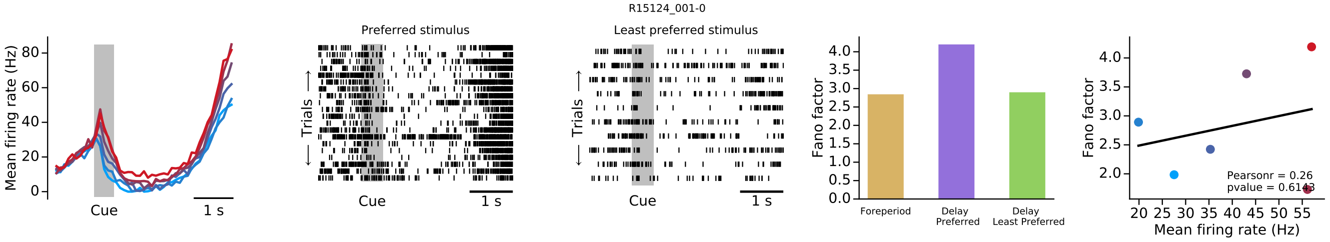
R15124_001-0
Mean firing rate (Hz)
Cue
1 s
Preferred stimulus
Least preferred stimulus
Trials
Fano factor
Foreperiod
Delay Preferred
Delay Least Preferred
Pearsonr = 0.26
pvalue = 0.6143
Mean firing rate (Hz)

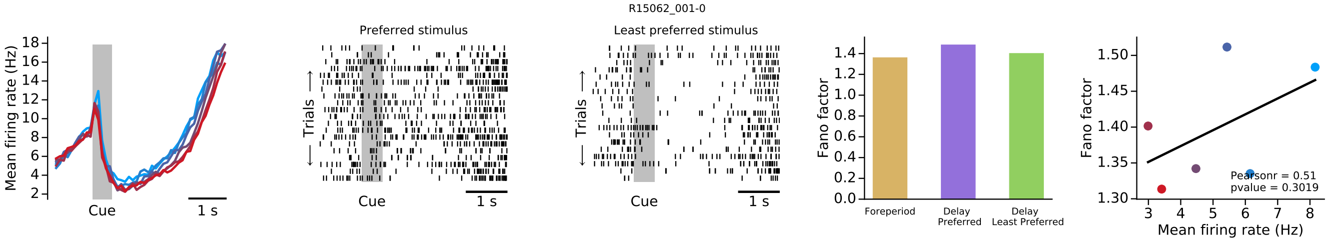
R15062_001-0
Mean firing rate (Hz)
Cue
1 s
Preferred stimulus
Least preferred stimulus
Trials
Fano factor
Foreperiod
Delay Preferred
Delay Least Preferred
Pearsonr = 0.51
pvalue = 0.3019
Mean firing rate (Hz)

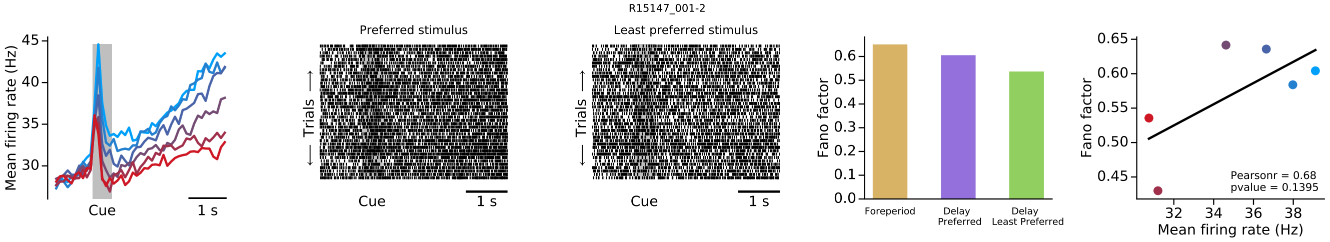
R15147_001-2
Mean firing rate (Hz)
Cue
1 s
Preferred stimulus
Least preferred stimulus
Trials
Fano factor
Foreperiod
Delay Preferred
Delay Least Preferred
Pearsonr = 0.68
pvalue = 0.1395
Mean firing rate (Hz)

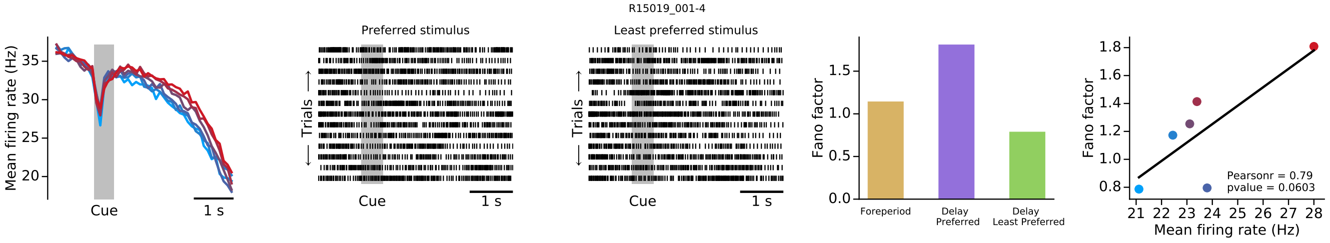
R15019_001-4
Mean firing rate (Hz)
Cue
1 s
Preferred stimulus
Least preferred stimulus
Trials
Fano factor
Foreperiod
Delay Preferred
Delay Least Preferred
Pearsonr = 0.79
pvalue = 0.0603
Mean firing rate (Hz)

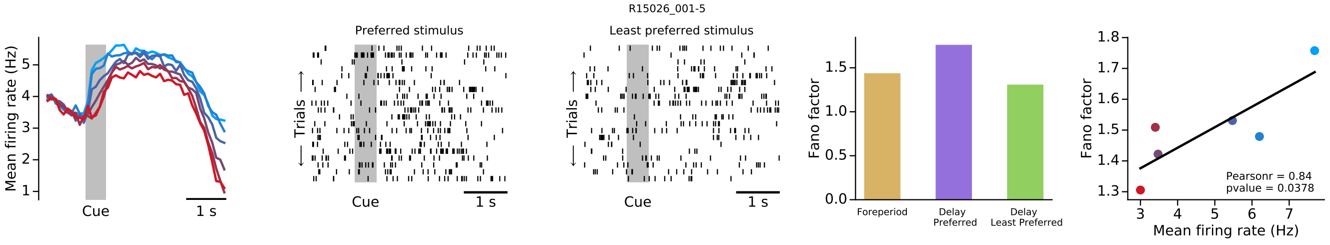
R15026_001-5
Mean firing rate (Hz)
Cue
1 s
Preferred stimulus
Least preferred stimulus
Trials
Fano factor
Foreperiod
Delay Preferred
Delay Least Preferred
Pearsonr = 0.84
pvalue = 0.0378
Mean firing rate (Hz)

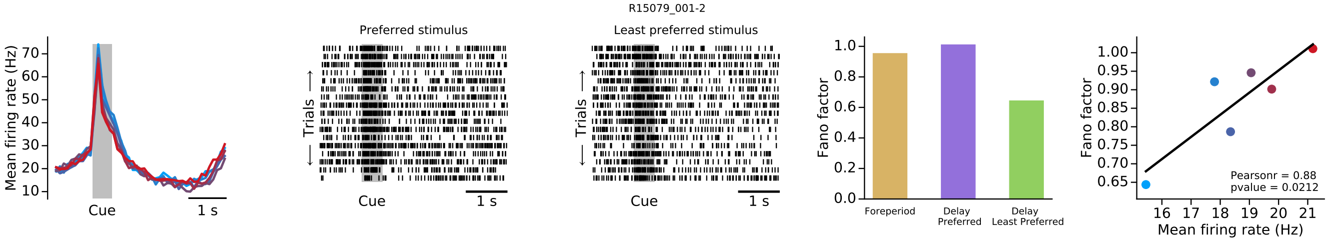
R15079_001-2
Mean firing rate (Hz)
Cue
1 s
Preferred stimulus
Least preferred stimulus
Trials
Fano factor
Foreperiod
Delay Preferred
Delay Least Preferred
Pearsonr = 0.88
pvalue = 0.0212
Mean firing rate (Hz)

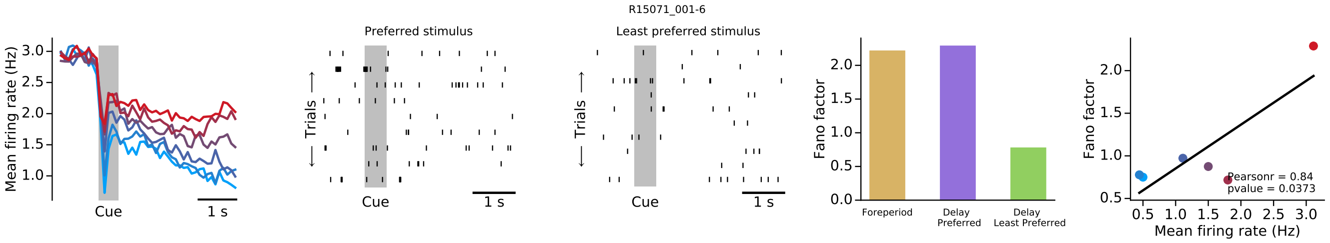
R15071_001-6
Mean firing rate (Hz)
Cue
1 s
Preferred stimulus
Least preferred stimulus
Trials
Fano factor
Foreperiod
Delay Preferred
Delay Least Preferred
Pearsonr = 0.84
pvalue = 0.0373
Mean firing rate (Hz)

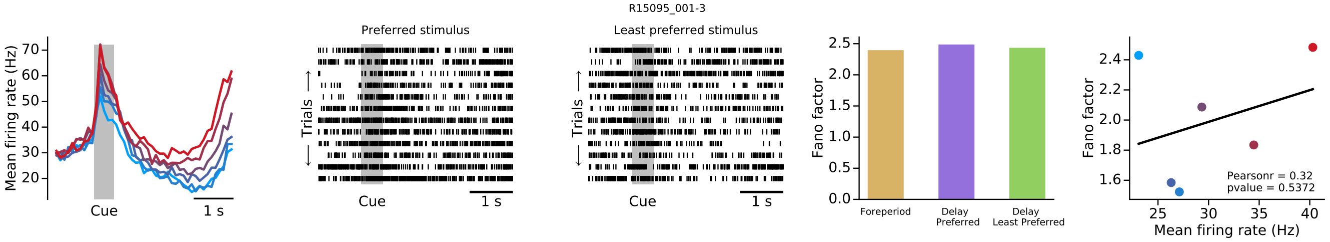
R15095_001-3
Mean firing rate (Hz)
70
60
50
40
30
20
Cue
1 s
Preferred stimulus
Trials
Cue
1 s
Least preferred stimulus
Trials
Cue
1 s
Fano factor
2.5
2.0
1.5
1.0
0.5
0.0
Foreperiod
Delay Preferred
Delay Least Preferred
Fano factor
2.4
2.2
2.0
1.8
1.6
25
30
35
40
Mean firing rate (Hz)
Pearsonr = 0.32
pvalue = 0.5372

### Group 2

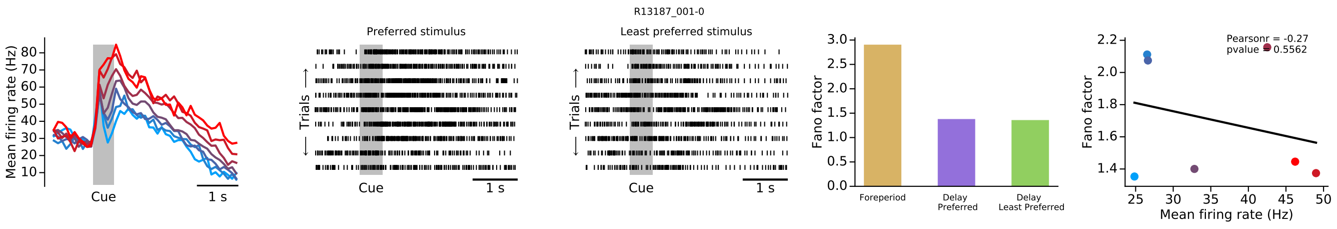
R13187_001-0
Preferred stimulus
Least preferred stimulus
Mean firing rate (Hz)
Trials
Cue
1 s
Fano factor
Foreperiod
Delay Preferred
Delay Least Preferred
Pearsonr = -0.27
pvalue = 0.5562
Mean firing rate (Hz)

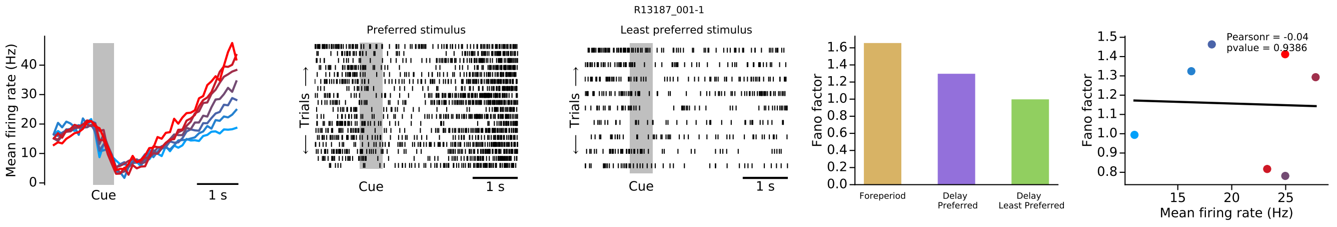
R13187_001-1
Preferred stimulus
Least preferred stimulus
Mean firing rate (Hz)
Trials
Cue
1 s
Fano factor
Foreperiod
Delay Preferred
Delay Least Preferred
Pearsonr = -0.04
pvalue = 0.9386
Mean firing rate (Hz)

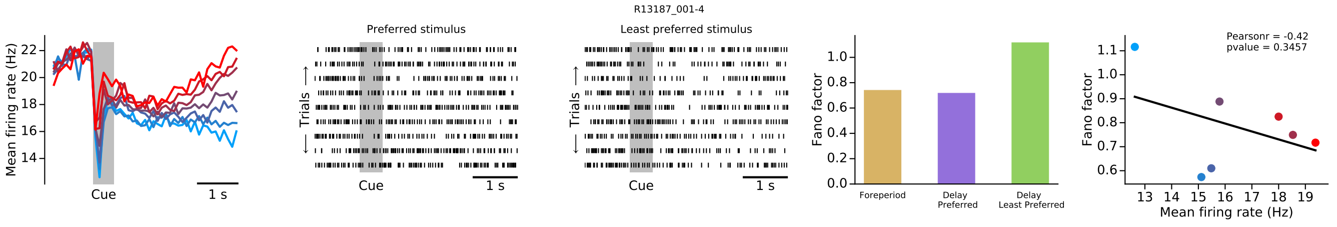
R13187_001-4
Preferred stimulus
Least preferred stimulus
Mean firing rate (Hz)
Trials
Cue
1 s
Fano factor
Foreperiod
Delay Preferred
Delay Least Preferred
Pearsonr = -0.42
pvalue = 0.3457
Mean firing rate (Hz)

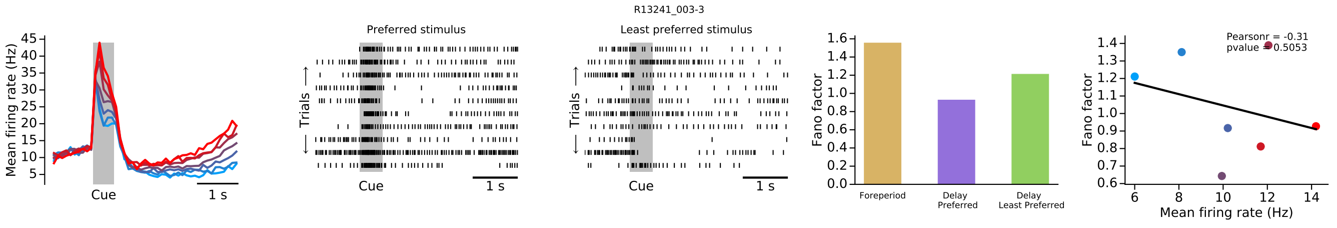
R13241_003-3
Preferred stimulus
Least preferred stimulus
Mean firing rate (Hz)
Trials
Cue
1 s
Fano factor
Foreperiod
Delay Preferred
Delay Least Preferred
Pearsonr = -0.31
pvalue = 0.5053
Mean firing rate (Hz)

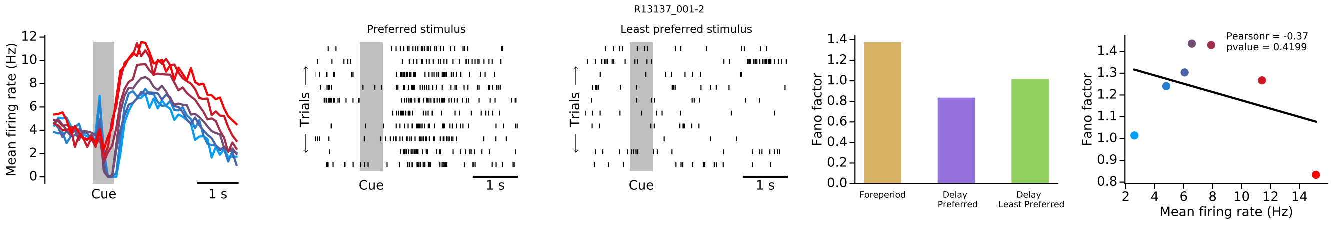
R13137_001-2
Preferred stimulus
Least preferred stimulus
Mean firing rate (Hz)
Trials
Cue
1 s
Fano factor
Foreperiod
Delay Preferred
Delay Least Preferred
Pearsonr = -0.37
pvalue = 0.4199
Mean firing rate (Hz)

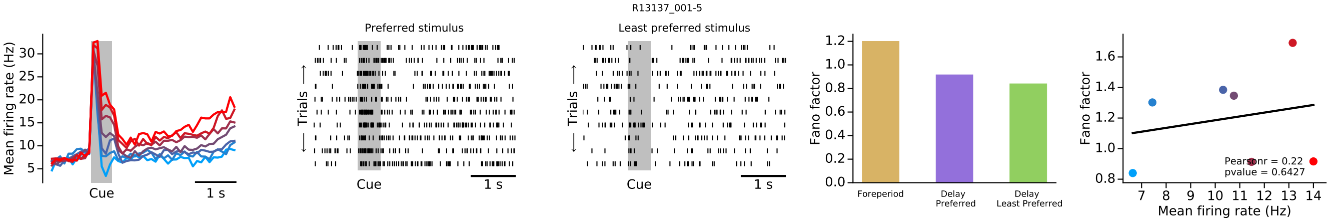
R13137_001-5
Mean firing rate (Hz)
Cue
1 s
Preferred stimulus
Least preferred stimulus
Trials
Fano factor
Foreperiod
Delay Preferred
Delay Least Preferred
Pearsonr = 0.22
pvalue = 0.6427
Mean firing rate (Hz)

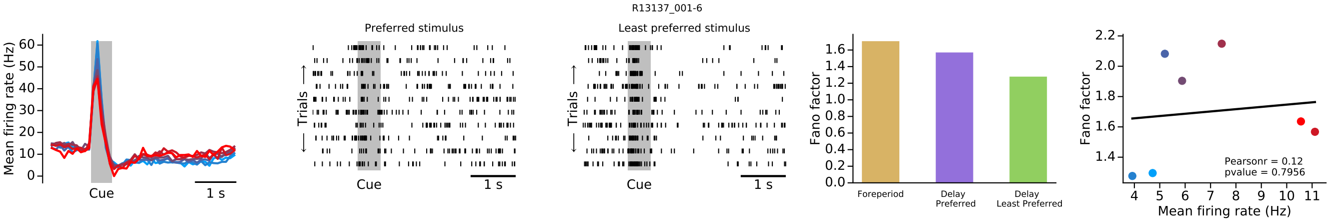
R13137_001-6
Mean firing rate (Hz)
Cue
1 s
Preferred stimulus
Least preferred stimulus
Trials
Fano factor
Foreperiod
Delay Preferred
Delay Least Preferred
Pearsonr = 0.12
pvalue = 0.7956
Mean firing rate (Hz)

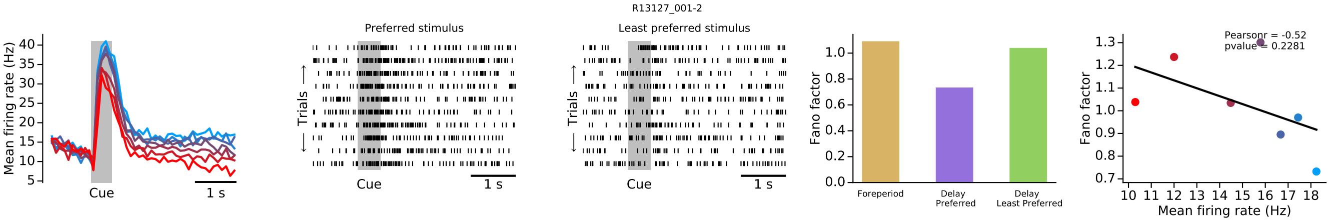
R13127_001-2
Mean firing rate (Hz)
Cue
1 s
Preferred stimulus
Least preferred stimulus
Trials
Fano factor
Foreperiod
Delay Preferred
Delay Least Preferred
Pearsonr = -0.52
pvalue = 0.2281
Mean firing rate (Hz)

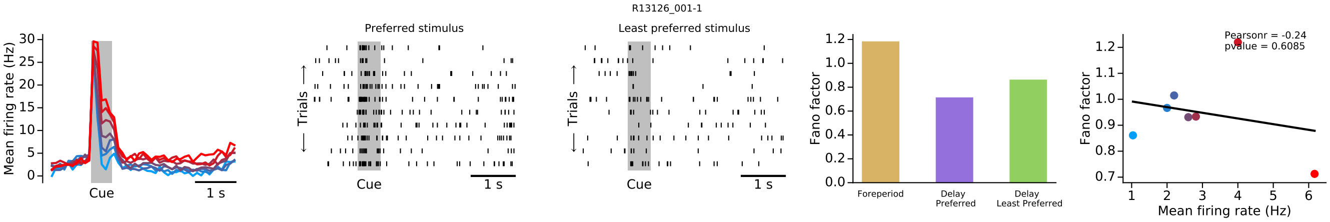
R13126_001-1
Mean firing rate (Hz)
Cue
1 s
Preferred stimulus
Least preferred stimulus
Trials
Fano factor
Foreperiod
Delay Preferred
Delay Least Preferred
Pearsonr = -0.24
pvalue = 0.6085
Mean firing rate (Hz)

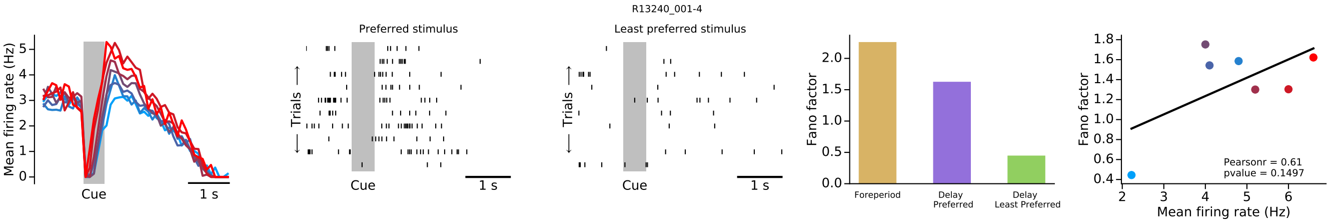
R13240_001-4
Mean firing rate (Hz)
Cue
1 s
Preferred stimulus
Least preferred stimulus
Trials
Fano factor
Foreperiod
Delay Preferred
Delay Least Preferred
Pearsonr = 0.61
pvalue = 0.1497
Mean firing rate (Hz)

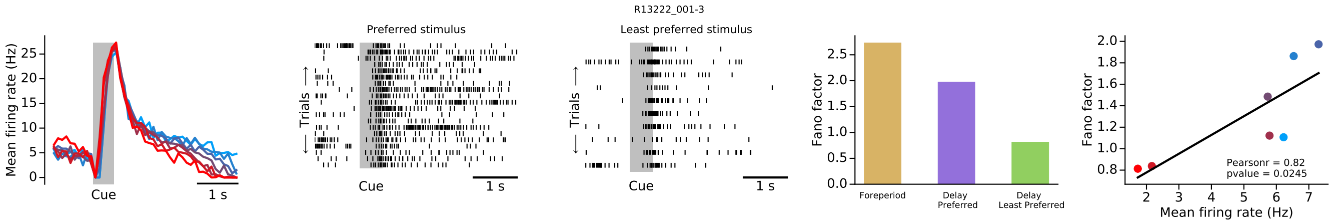
R13222_001-3
Preferred stimulus
Least preferred stimulus
Mean firing rate (Hz)
Cue
1 s
Trials
Fano factor
Foreperiod
Delay Preferred
Delay Least Preferred
Pearsonr = 0.82
pvalue = 0.0245
Mean firing rate (Hz)

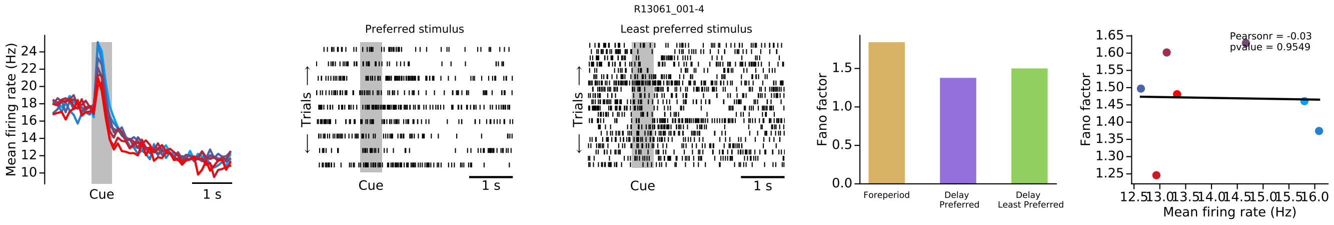
R13061_001-4
Preferred stimulus
Least preferred stimulus
Mean firing rate (Hz)
Cue
1 s
Trials
Fano factor
Foreperiod
Delay Preferred
Delay Least Preferred
Pearsonr = -0.03
pvalue = 0.9549
Mean firing rate (Hz)

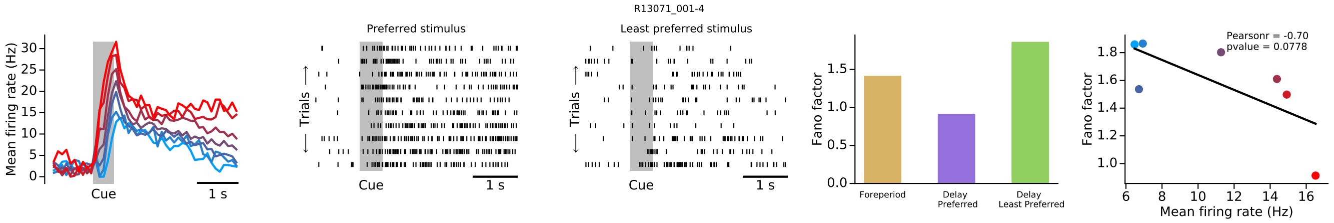
R13071_001-4
Preferred stimulus
Least preferred stimulus
Mean firing rate (Hz)
Cue
1 s
Trials
Fano factor
Foreperiod
Delay Preferred
Delay Least Preferred
Pearsonr = -0.70
pvalue = 0.0778
Mean firing rate (Hz)

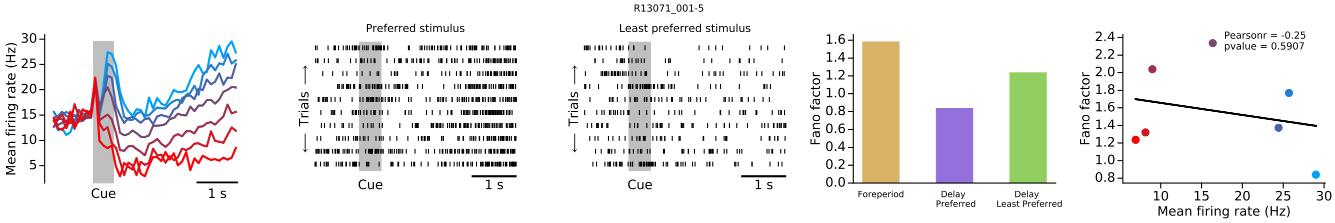
R13071_001-5
Preferred stimulus
Least preferred stimulus
Mean firing rate (Hz)
Cue
1 s
Trials
Fano factor
Foreperiod
Delay Preferred
Delay Least Preferred
Pearsonr = -0.25
pvalue = 0.5907
Mean firing rate (Hz)

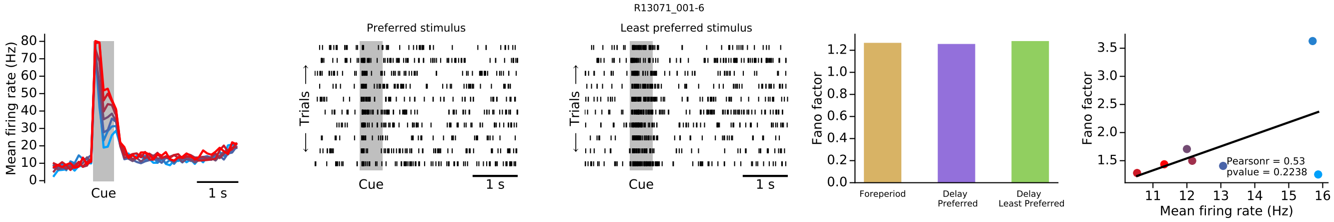
R13071_001-6
Preferred stimulus
Least preferred stimulus
Mean firing rate (Hz)
Cue
1 s
Trials
Fano factor
Foreperiod
Delay Preferred
Delay Least Preferred
Pearsonr = 0.53
pvalue = 0.2238
Mean firing rate (Hz)

R13186_001-4

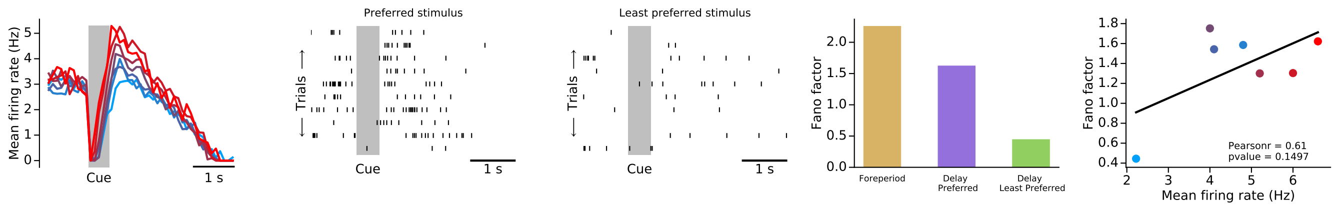


R13087_001-6

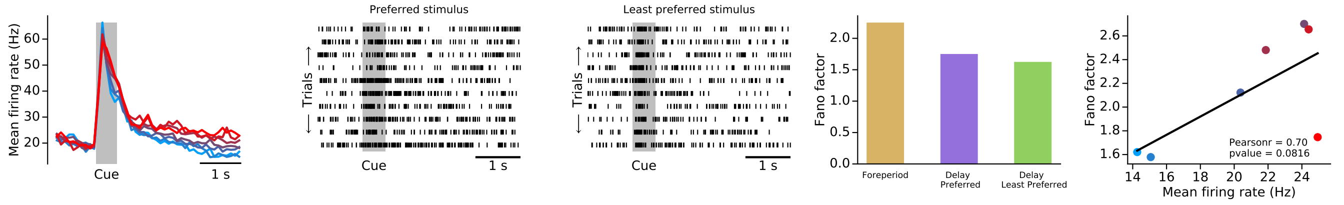


R13179_001-4

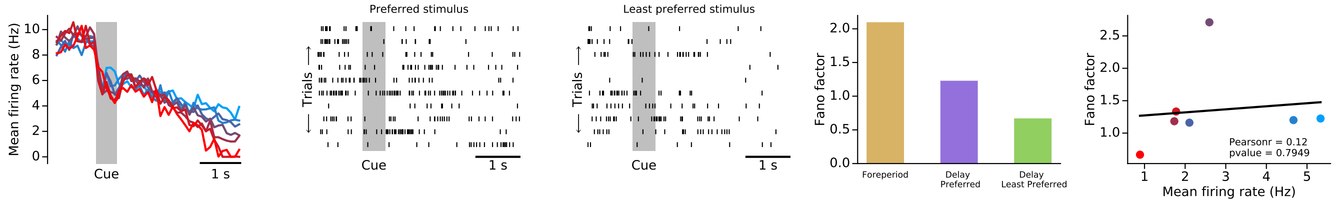


R13194_001-3

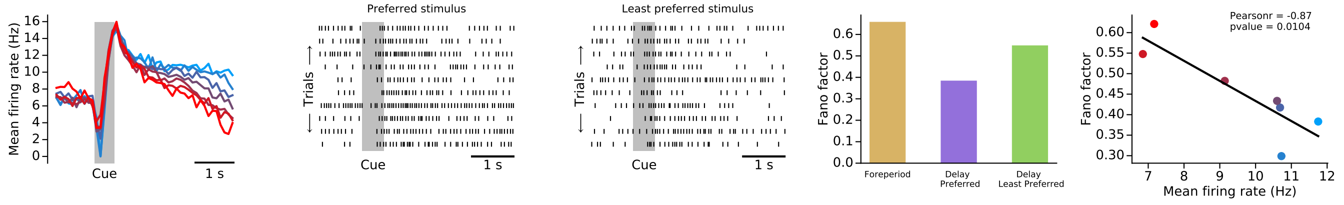


R13194_001-4

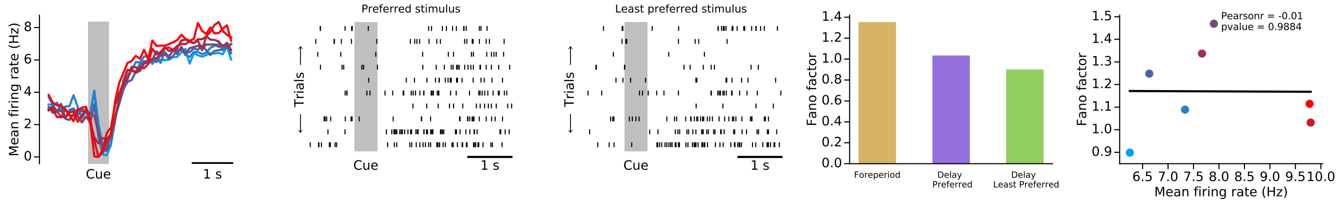

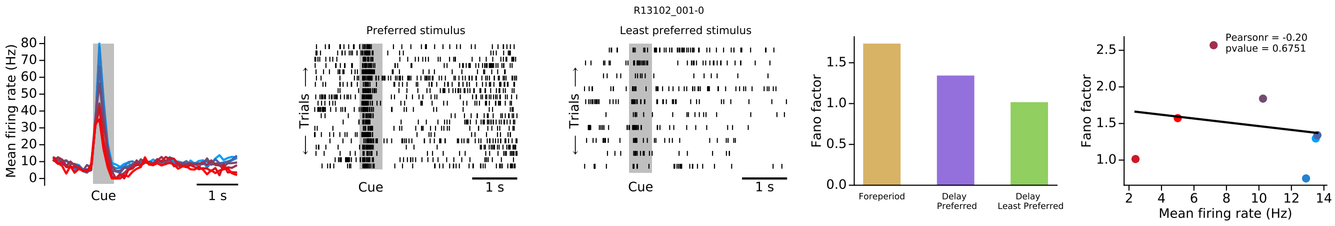
R13102_001-0
Preferred stimulus
Least preferred stimulus
Mean firing rate (Hz)
Cue
1 s
Trials
Fano factor
Foreperiod
Delay Preferred
Delay Least Preferred
Pearsonr = -0.20
pvalue = 0.6751
Mean firing rate (Hz)

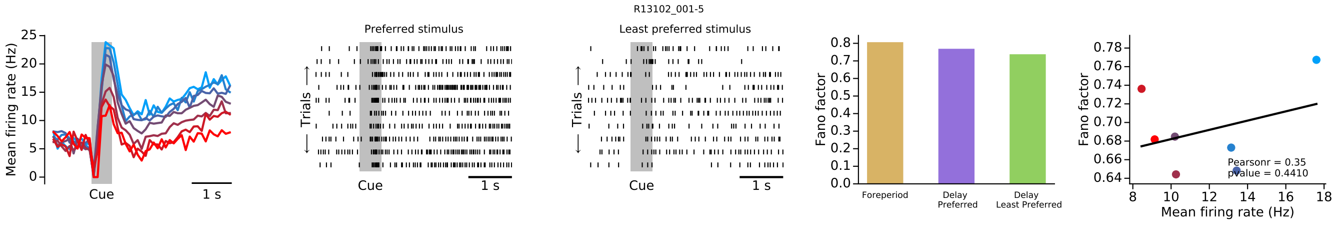
R13102_001-5
Preferred stimulus
Least preferred stimulus
Mean firing rate (Hz)
Cue
1 s
Trials
Fano factor
Foreperiod
Delay Preferred
Delay Least Preferred
Pearsonr = 0.35
pvalue = 0.4410
Mean firing rate (Hz)

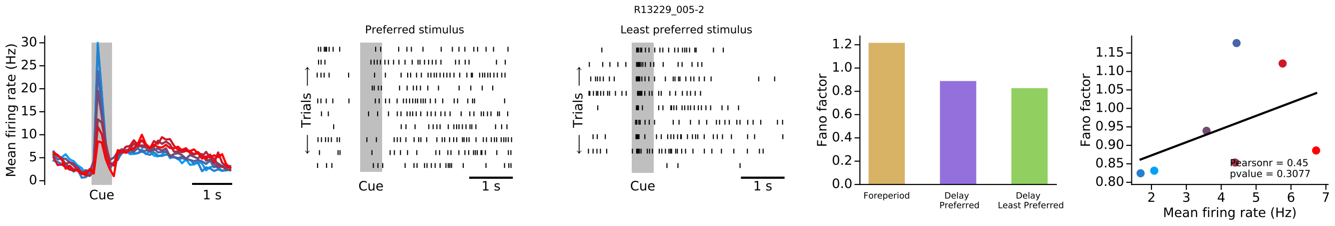
R13229_005-2
Preferred stimulus
Least preferred stimulus
Mean firing rate (Hz)
Cue
1 s
Trials
Fano factor
Foreperiod
Delay Preferred
Delay Least Preferred
Pearsonr = 0.45
pvalue = 0.3077
Mean firing rate (Hz)

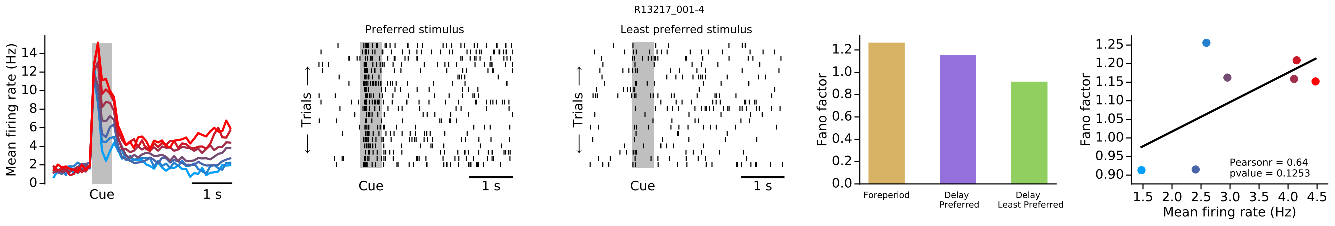
R13217_001-4
Preferred stimulus
Least preferred stimulus
Mean firing rate (Hz)
Cue
1 s
Trials
Fano factor
Foreperiod
Delay Preferred
Delay Least Preferred
Pearsonr = 0.64
pvalue = 0.1253
Mean firing rate (Hz)

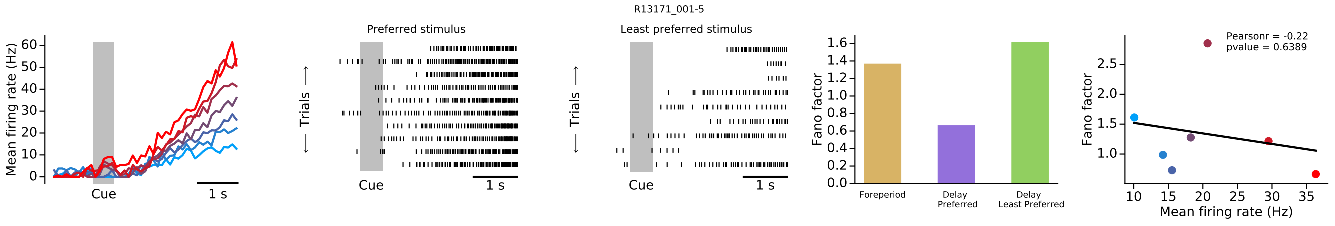
R13171_001-5
Preferred stimulus
Least preferred stimulus
Mean firing rate (Hz)
Cue
1 s
Trials
Fano factor
Foreperiod
Delay Preferred
Delay Least Preferred
Pearsonr = -0.22
pvalue = 0.6389
Mean firing rate (Hz)

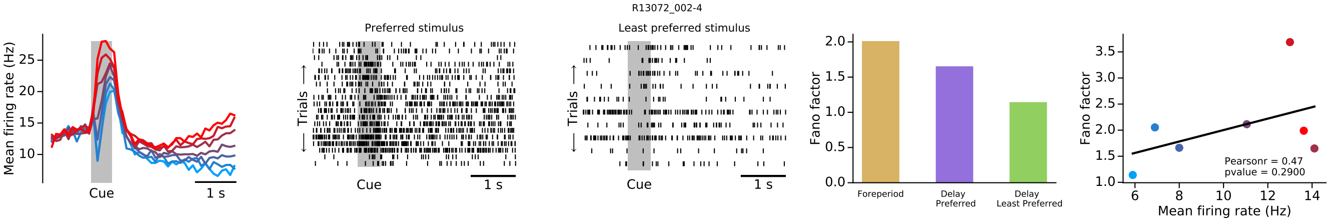
R13072_002-4
Mean firing rate (Hz)
Cue
1 s
Preferred stimulus
Least preferred stimulus
Trials
Fano factor
Foreperiod
Delay Preferred
Delay Least Preferred
Pearsonr = 0.47
pvalue = 0.2900
Mean firing rate (Hz)

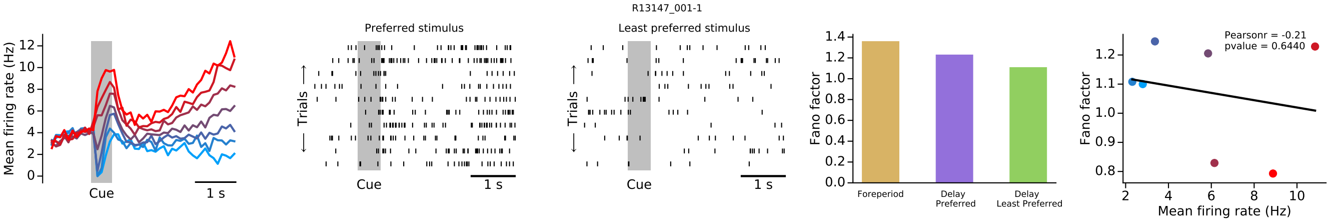
R13147_001-1
Mean firing rate (Hz)
Cue
1 s
Preferred stimulus
Least preferred stimulus
Trials
Fano factor
Foreperiod
Delay Preferred
Delay Least Preferred
Pearsonr = -0.21
pvalue = 0.6440
Mean firing rate (Hz)

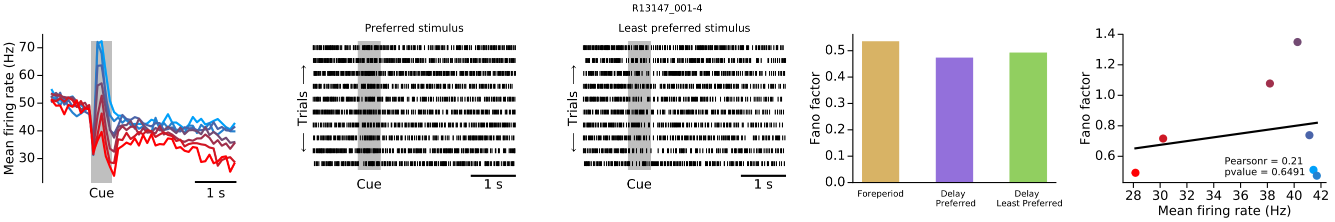
R13147_001-4
Mean firing rate (Hz)
Cue
1 s
Preferred stimulus
Least preferred stimulus
Trials
Fano factor
Foreperiod
Delay Preferred
Delay Least Preferred
Pearsonr = 0.21
pvalue = 0.6491
Mean firing rate (Hz)

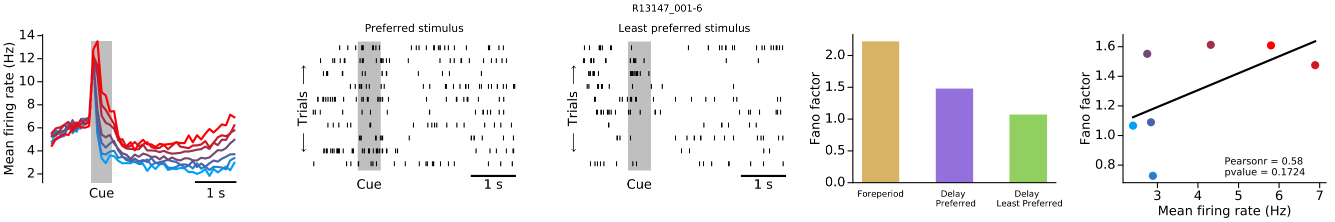
R13147_001-6
Mean firing rate (Hz)
Cue
1 s
Preferred stimulus
Least preferred stimulus
Trials
Fano factor
Foreperiod
Delay Preferred
Delay Least Preferred
Pearsonr = 0.58
pvalue = 0.1724
Mean firing rate (Hz)

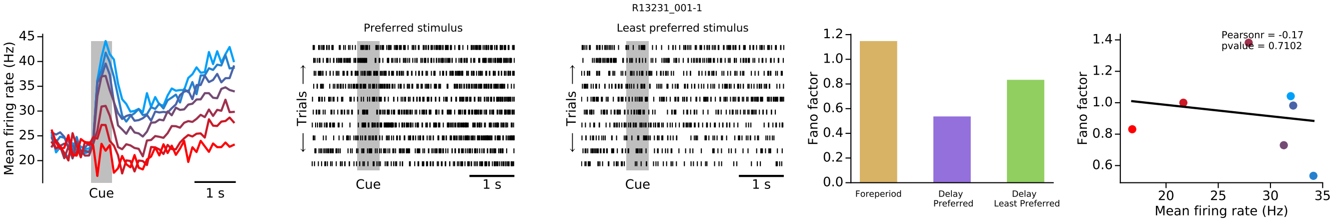
R13231_001-1
Mean firing rate (Hz)
Cue
1 s
Preferred stimulus
Least preferred stimulus
Trials
Fano factor
Foreperiod
Delay Preferred
Delay Least Preferred
Pearsonr = -0.17
pvalue = 0.7102
Mean firing rate (Hz)

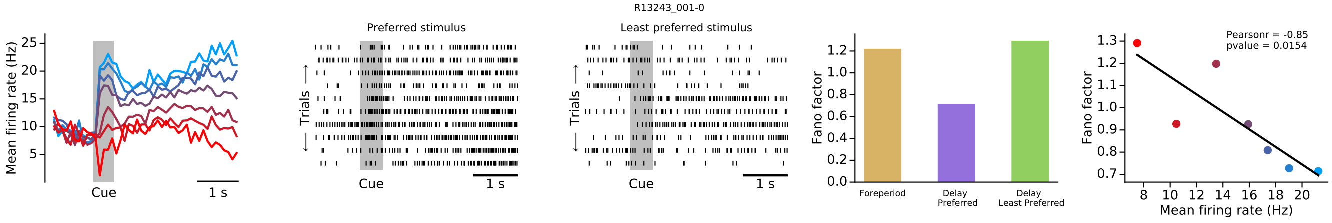
R13243_001-0
Mean firing rate (Hz)
Cue
1 s
Preferred stimulus
Least preferred stimulus
Trials
Fano factor
Foreperiod
Delay Preferred
Delay Least Preferred
Pearsonr = -0.85
pvalue = 0.0154
Mean firing rate (Hz)

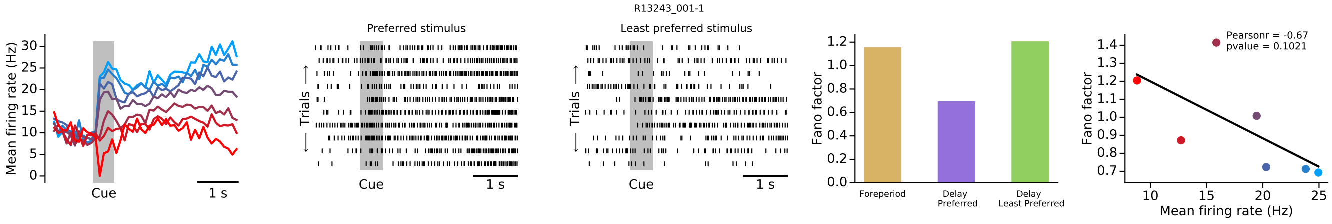
R13243_001-1
Mean firing rate (Hz)
Cue
1 s
Preferred stimulus
Least preferred stimulus
Trials
Fano factor
Foreperiod
Delay Preferred
Delay Least Preferred
Pearsonr = -0.67
pvalue = 0.1021
Mean firing rate (Hz)

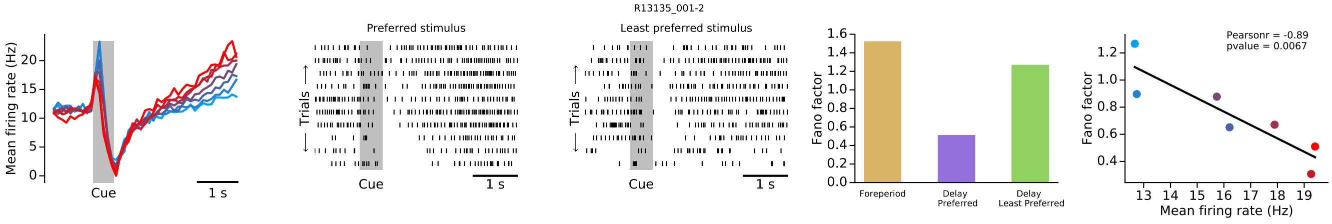
R13135_001-2
Mean firing rate (Hz)
Cue
1 s
Preferred stimulus
Least preferred stimulus
Trials
Fano factor
Foreperiod
Delay Preferred
Delay Least Preferred
Pearsonr = -0.89
pvalue = 0.0067
Mean firing rate (Hz)

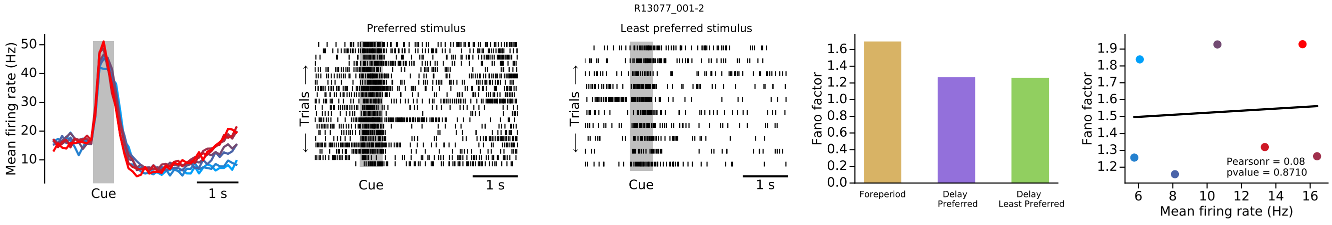
R13077_001-2
Mean firing rate (Hz)
Cue
1 s
Preferred stimulus
Least preferred stimulus
Trials
Fano factor
Foreperiod
Delay Preferred
Delay Least Preferred
Pearsonr = 0.08
pvalue = 0.8710
Mean firing rate (Hz)

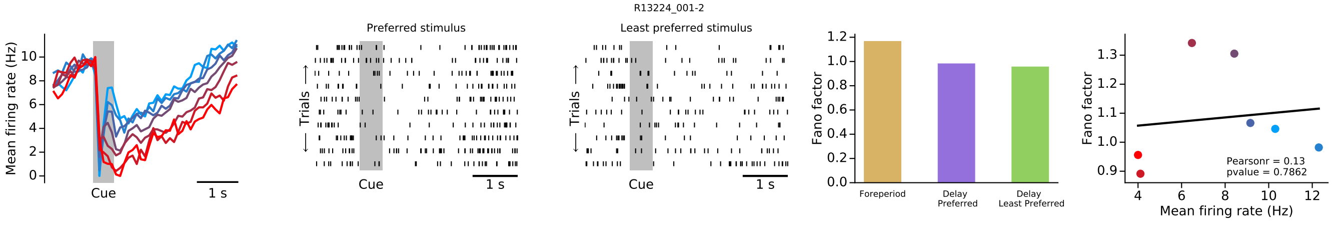
R13224_001-2
Mean firing rate (Hz)
Cue
1 s
Preferred stimulus
Least preferred stimulus
Trials
Fano factor
Foreperiod
Delay Preferred
Delay Least Preferred
Pearsonr = 0.13
pvalue = 0.7862
Mean firing rate (Hz)

R13117_001-2

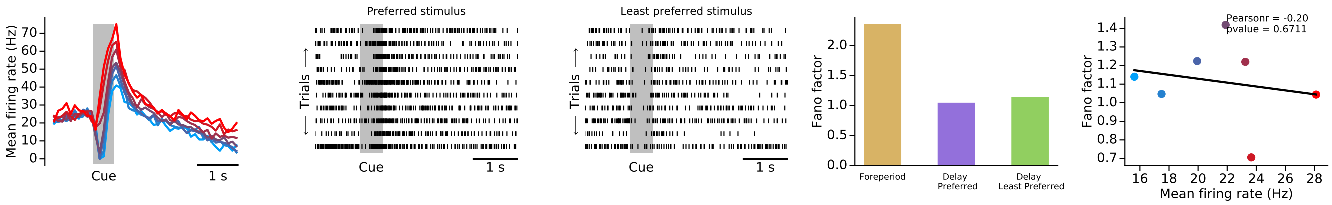


R13138_001-1

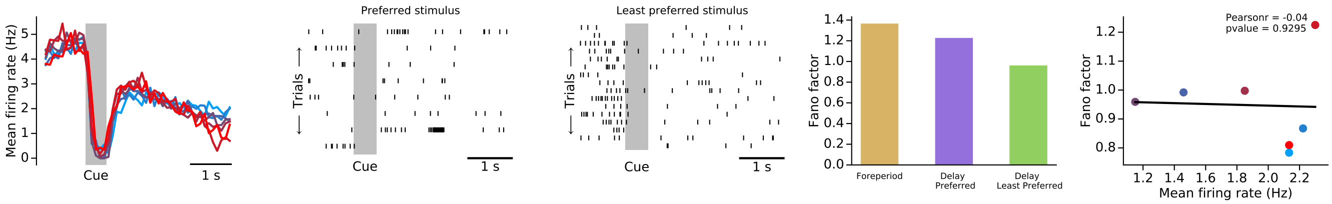


R13138_001-4

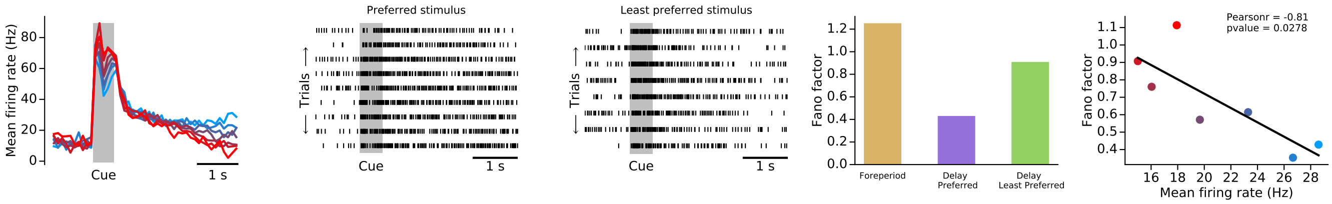


R13224_005-3

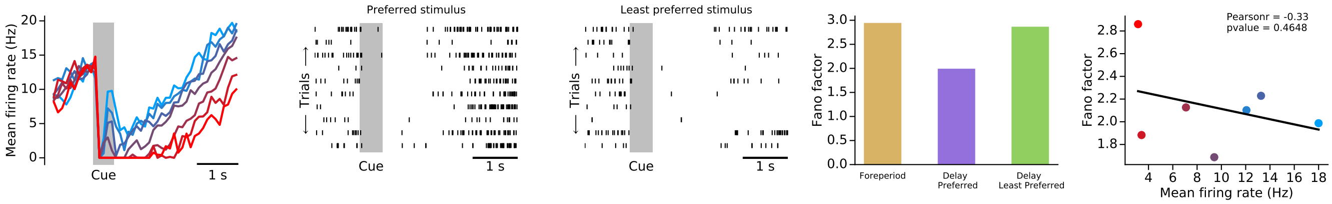


R13143_001-2

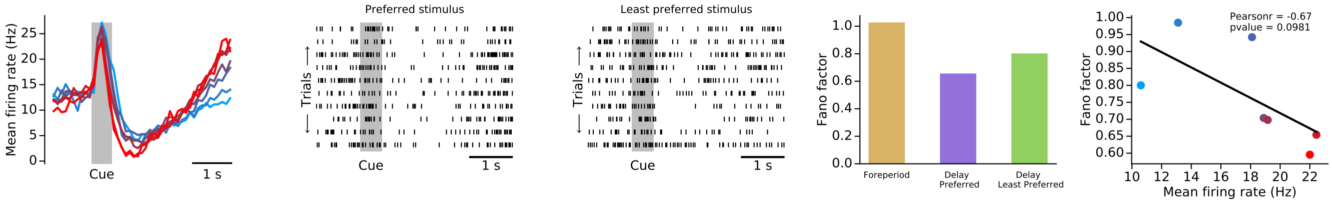

R13143_001-5

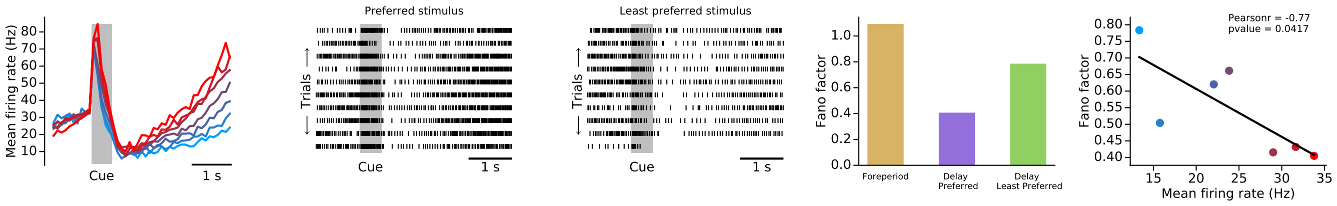


R13158_001-5

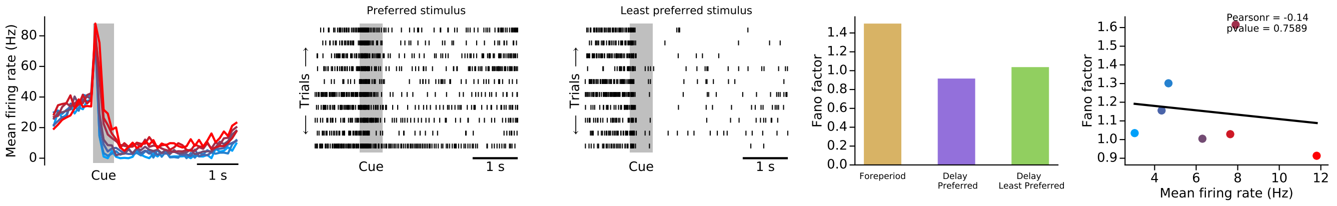


R13089_001-5

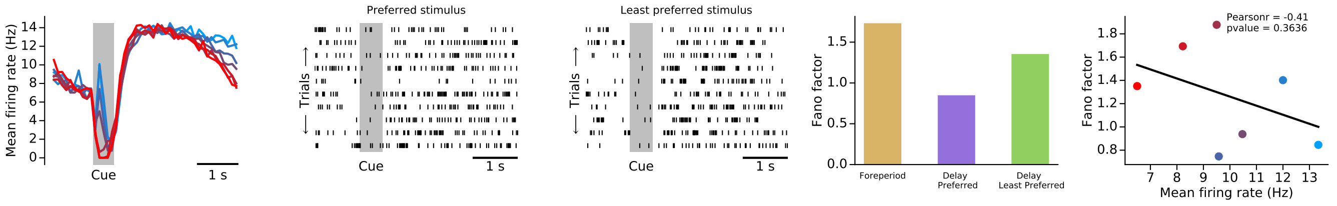


R13208_001-2

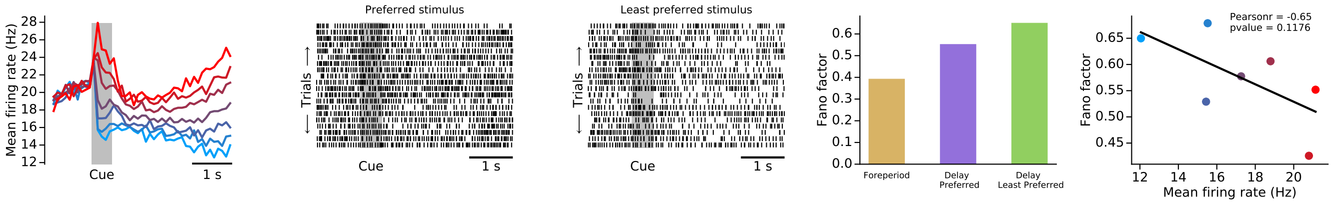


R13158_003-5

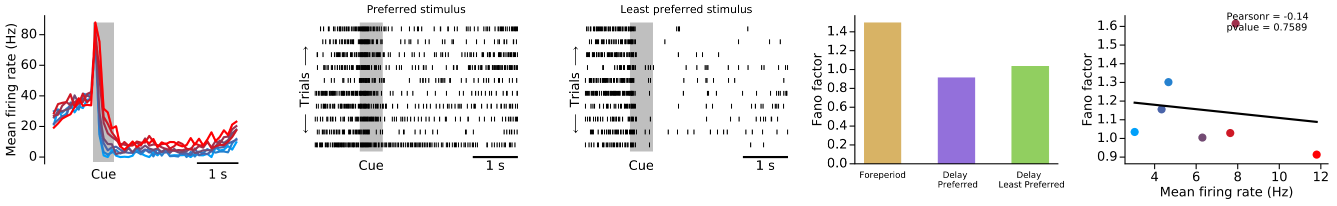

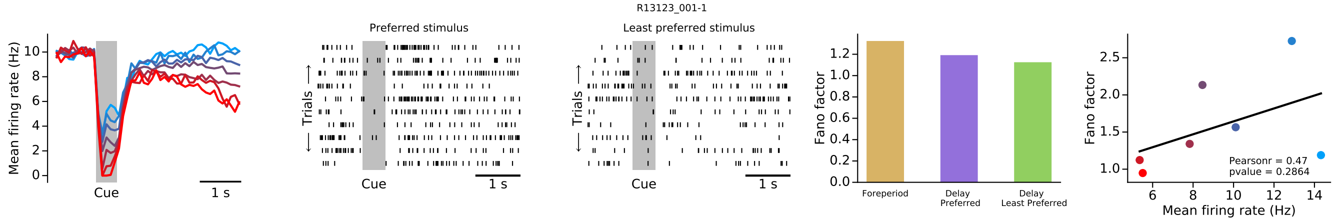
R13123_001-1
Mean firing rate (Hz)
Cue
1 s
Preferred stimulus
Least preferred stimulus
Trials
Fano factor
Foreperiod
Delay Preferred
Delay Least Preferred
Pearsonr = 0.47
pvalue = 0.2864
Mean firing rate (Hz)

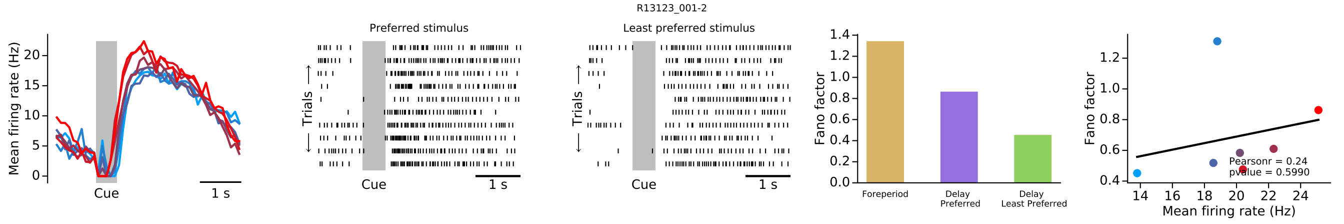
R13123_001-2
Mean firing rate (Hz)
Cue
1 s
Preferred stimulus
Least preferred stimulus
Trials
Fano factor
Foreperiod
Delay Preferred
Delay Least Preferred
Pearsonr = 0.24
pvalue = 0.5990
Mean firing rate (Hz)

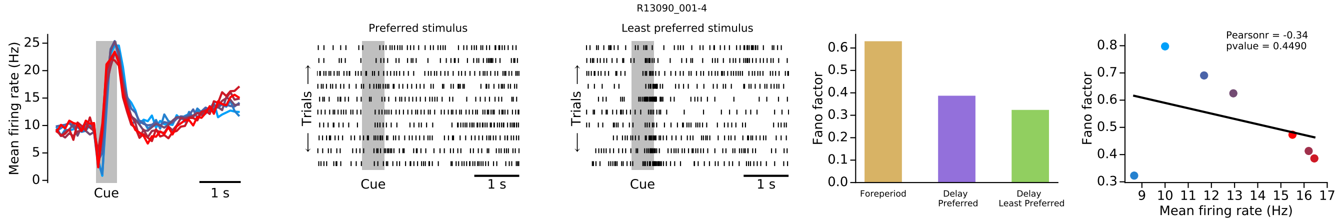
R13090_001-4
Mean firing rate (Hz)
Cue
1 s
Preferred stimulus
Least preferred stimulus
Trials
Fano factor
Foreperiod
Delay Preferred
Delay Least Preferred
Pearsonr = -0.34
pvalue = 0.4490
Mean firing rate (Hz)

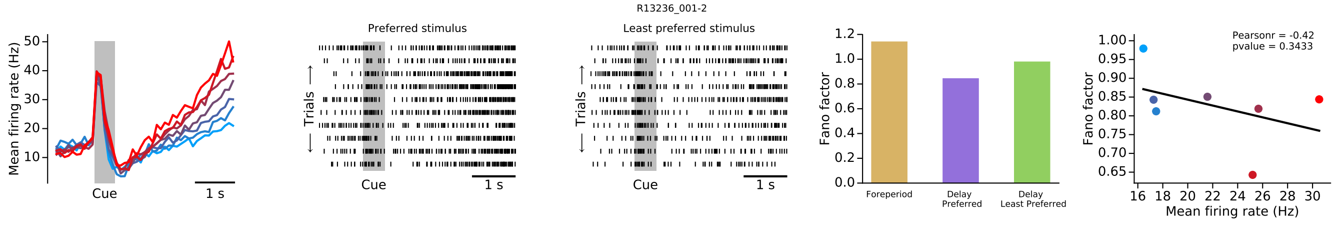
R13236_001-2
Mean firing rate (Hz)
Cue
1 s
Preferred stimulus
Least preferred stimulus
Trials
Fano factor
Foreperiod
Delay Preferred
Delay Least Preferred
Pearsonr = -0.42
pvalue = 0.3433
Mean firing rate (Hz)

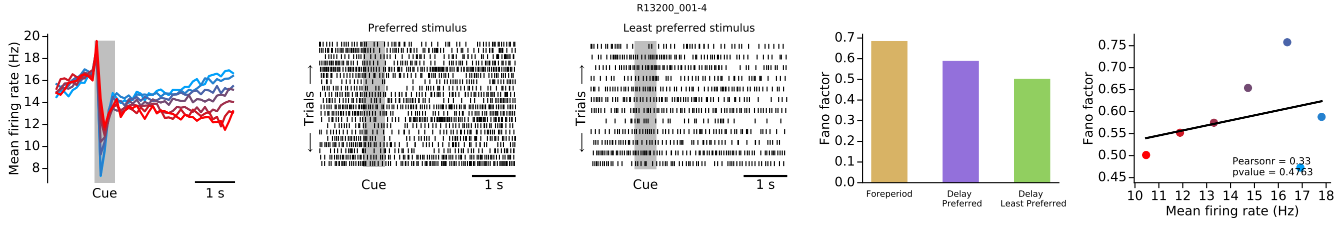
R13200_001-4
Mean firing rate (Hz)
Cue
1 s
Preferred stimulus
Least preferred stimulus
Trials
Fano factor
Foreperiod
Delay Preferred
Delay Least Preferred
Pearsonr = 0.33
Mean firing rate (Hz)

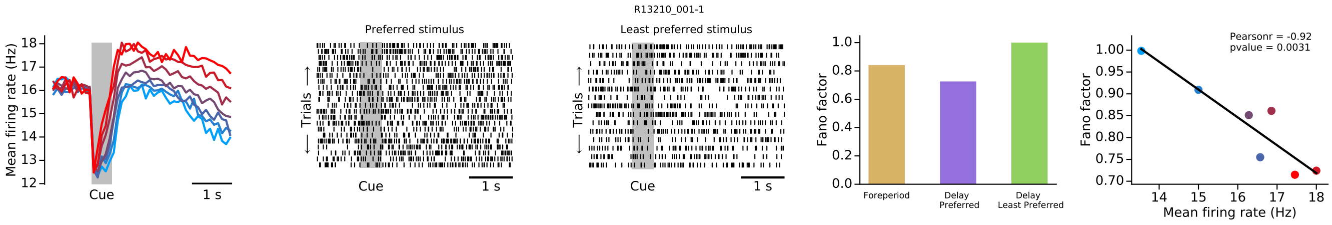
R13210_001-1
Mean firing rate (Hz)
Cue
1 s
Preferred stimulus
Least preferred stimulus
Trials
Fano factor
Foreperiod
Delay Preferred
Delay Least Preferred
Pearsonr = -0.92
pvalue = 0.0031
Mean firing rate (Hz)

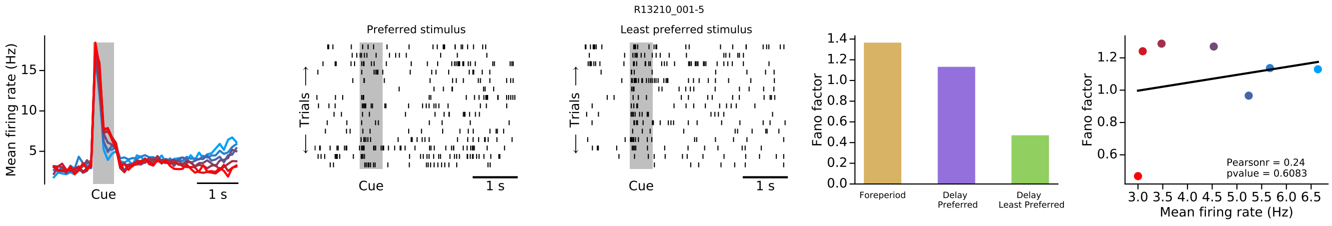
R13210_001-5
Mean firing rate (Hz)
Cue
1 s
Preferred stimulus
Least preferred stimulus
Trials
Fano factor
Foreperiod
Delay Preferred
Delay Least Preferred
Pearsonr = 0.24
pvalue = 0.6083
Mean firing rate (Hz)

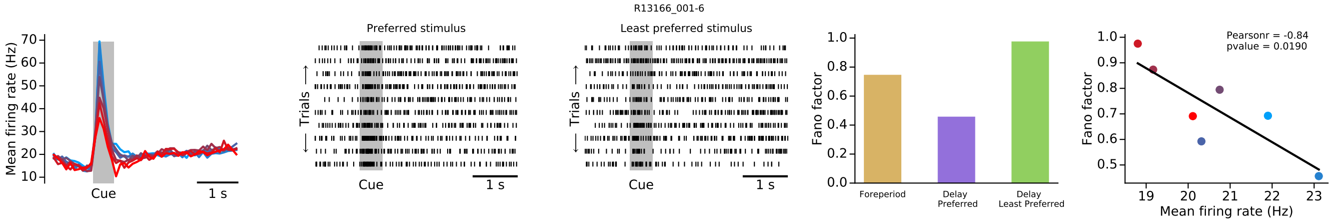
R13166_001-6
Mean firing rate (Hz)
Cue
1 s
Preferred stimulus
Least preferred stimulus
Trials
Fano factor
Foreperiod
Delay Preferred
Delay Least Preferred
Pearsonr = -0.84
pvalue = 0.0190
Mean firing rate (Hz)

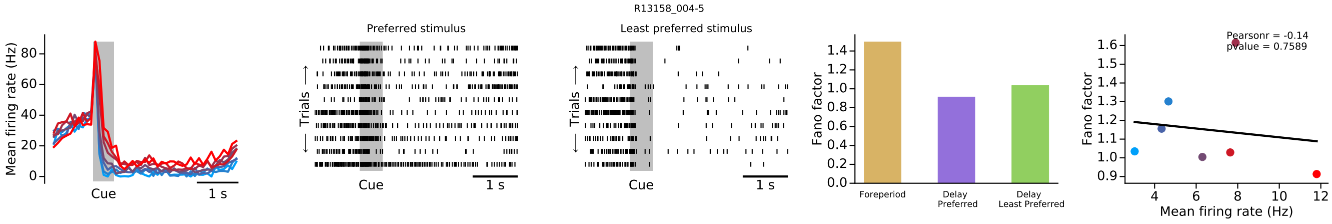
R13158_004-5
Mean firing rate (Hz)
Cue
1 s
Preferred stimulus
Least preferred stimulus
Trials
Fano factor
Foreperiod
Delay Preferred
Delay Least Preferred
Pearsonr = -0.14
pvalue = 0.7589
Mean firing rate (Hz)

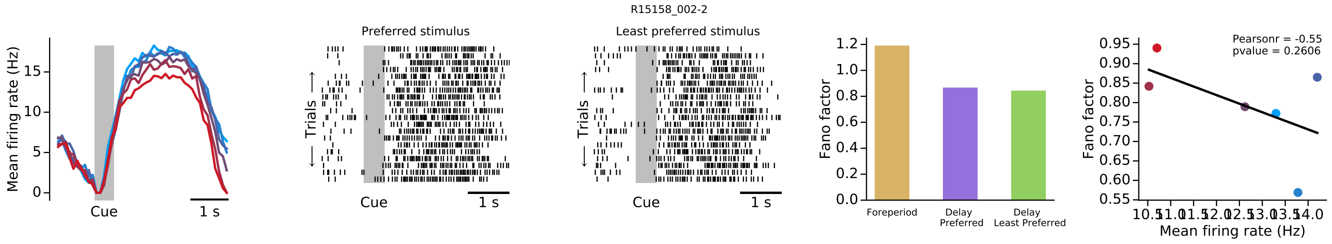
R15158_002-2
Mean firing rate (Hz)
Cue
1 s
Preferred stimulus
Least preferred stimulus
Trials
Fano factor
Foreperiod
Delay Preferred
Delay Least Preferred
Pearsonr = -0.55
pvalue = 0.2606
Mean firing rate (Hz)

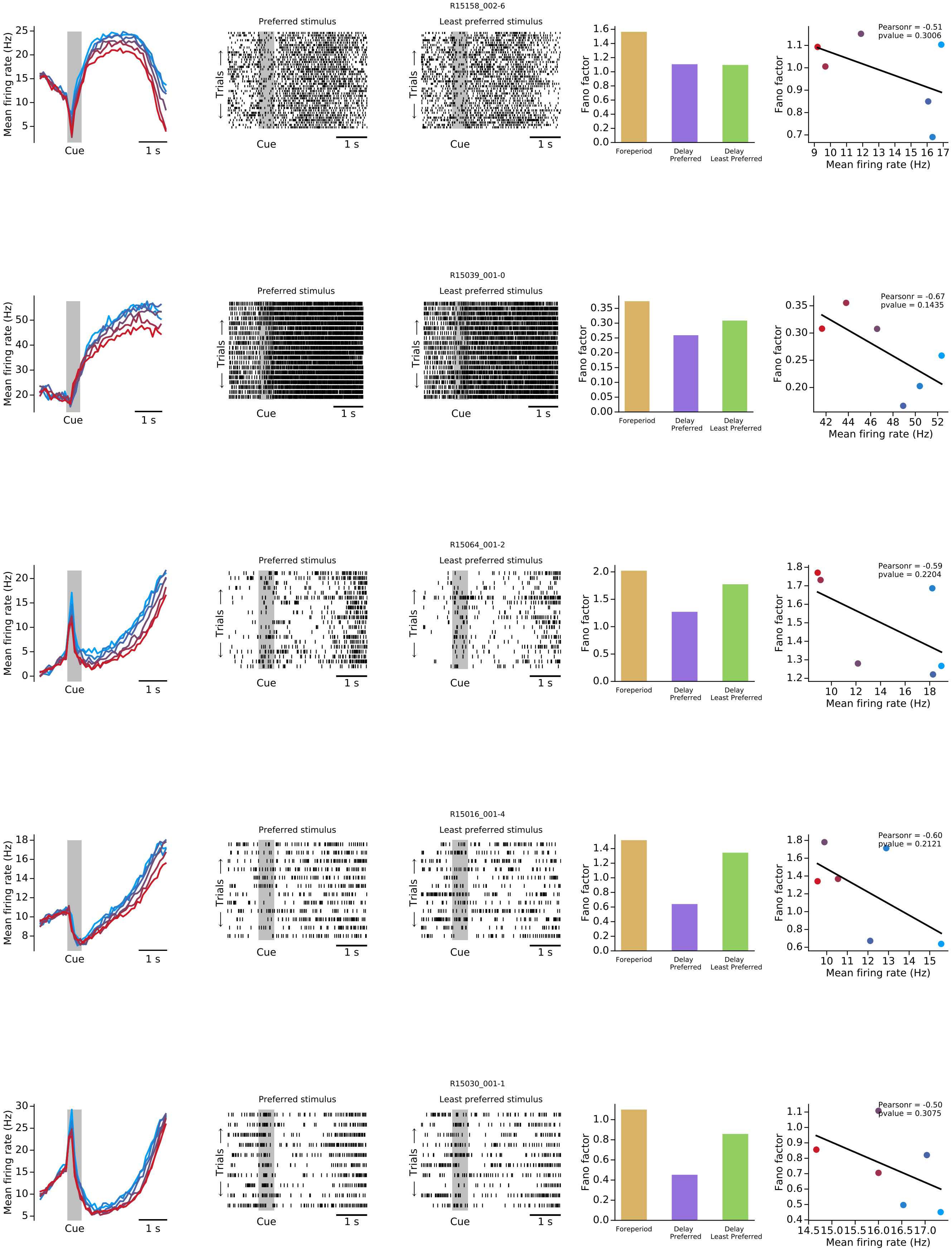
R15158_002-6
Preferred stimulus
Least preferred stimulus
Mean firing rate (Hz)
Cue
1 s
Trials
Fano factor
Foreperiod
Delay Preferred
Delay Least Preferred
Pearsonr = -0.51
pvalue = 0.3006
Mean firing rate (Hz)
R15039_001-0
Preferred stimulus
Least preferred stimulus
Pearsonr = -0.67
pvalue = 0.1435
R15064_001-2
Preferred stimulus
Least preferred stimulus
Pearsonr = -0.59
pvalue = 0.2204
R15016_001-4
Preferred stimulus
Least preferred stimulus
Pearsonr = -0.60
pvalue = 0.2121
R15030_001-1
Preferred stimulus
Least preferred stimulus
Pearsonr = -0.50
pvalue = 0.3075

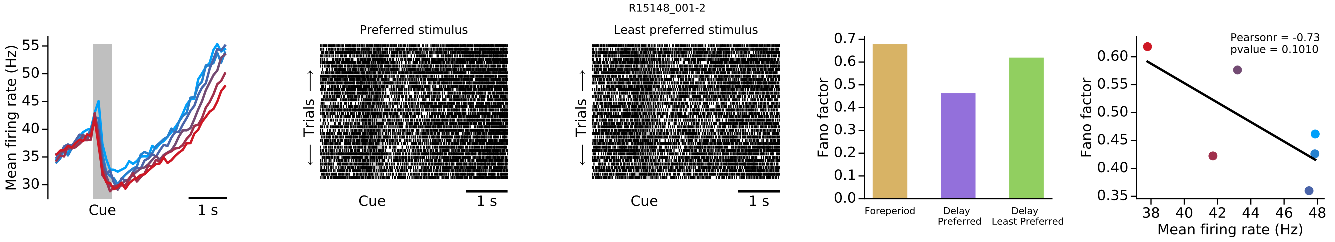
R15148_001-2
Mean firing rate (Hz)
Cue
1 s
Preferred stimulus
Least preferred stimulus
Trials
Fano factor
Foreperiod
Delay Preferred
Delay Least Preferred
Pearsonr = -0.73
pvalue = 0.1010
Mean firing rate (Hz)

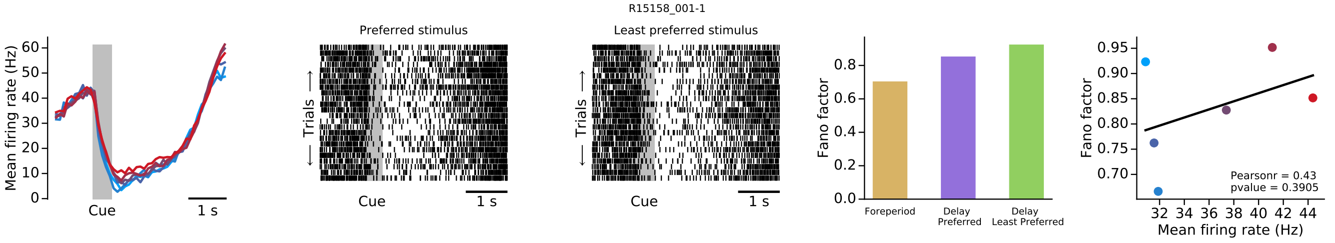
R15158_001-1
Mean firing rate (Hz)
Cue
1 s
Preferred stimulus
Least preferred stimulus
Trials
Fano factor
Foreperiod
Delay Preferred
Delay Least Preferred
Pearsonr = 0.43
pvalue = 0.3905
Mean firing rate (Hz)

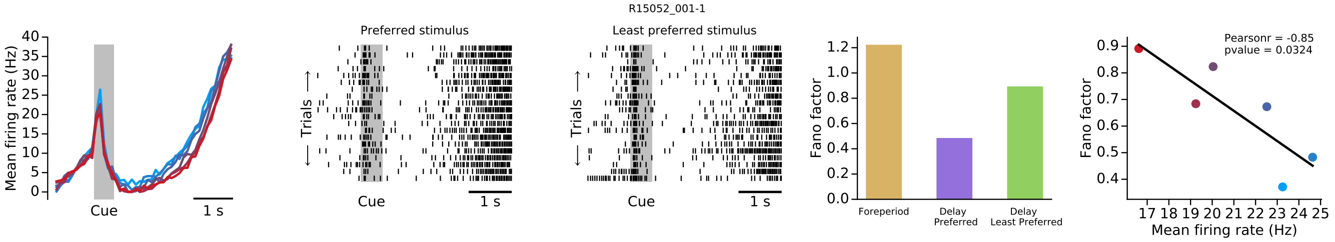
R15052_001-1
Mean firing rate (Hz)
Cue
1 s
Preferred stimulus
Least preferred stimulus
Trials
Fano factor
Foreperiod
Delay Preferred
Delay Least Preferred
Pearsonr = -0.85
pvalue = 0.0324
Mean firing rate (Hz)

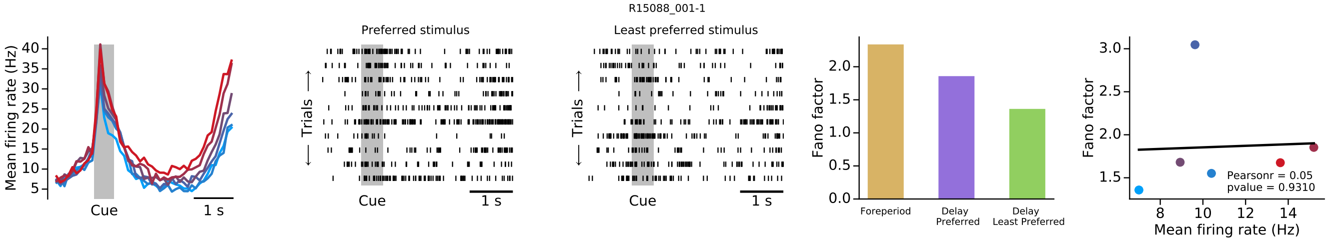
R15088_001-1
Mean firing rate (Hz)
Cue
1 s
Preferred stimulus
Least preferred stimulus
Trials
Fano factor
Foreperiod
Delay Preferred
Delay Least Preferred
Pearsonr = 0.05
pvalue = 0.9310
Mean firing rate (Hz)

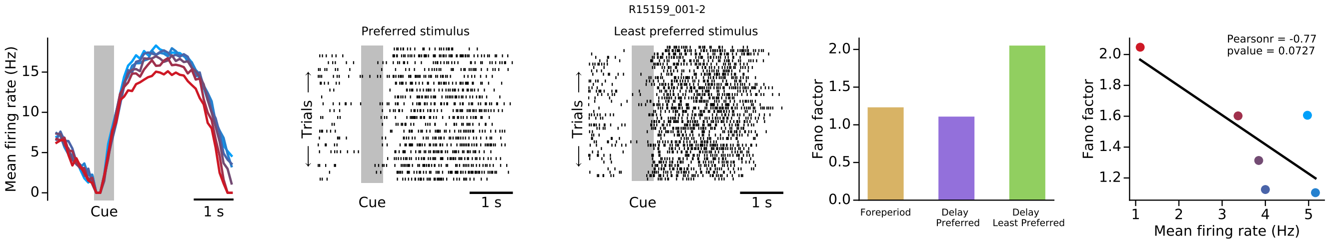
R15159_001-2
Mean firing rate (Hz)
Cue
1 s
Preferred stimulus
Least preferred stimulus
Trials
Fano factor
Foreperiod
Delay Preferred
Delay Least Preferred
Pearsonr = -0.77
pvalue = 0.0727
Mean firing rate (Hz)

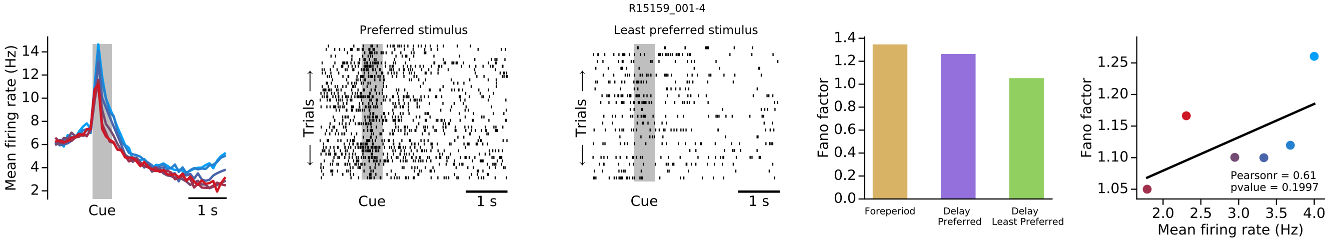
R15159_001-4
Mean firing rate (Hz)
Cue
1 s
Preferred stimulus
Least preferred stimulus
Trials
Fano factor
Foreperiod
Delay Preferred
Delay Least Preferred
Pearsonr = 0.61
pvalue = 0.1997
Mean firing rate (Hz)

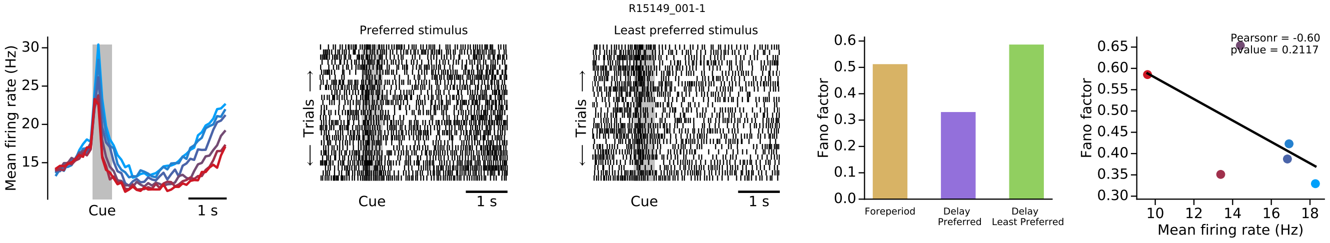
R15149_001-1
Mean firing rate (Hz)
Cue
1 s
Preferred stimulus
Least preferred stimulus
Trials
Fano factor
Foreperiod
Delay Preferred
Delay Least Preferred
Pearsonr = -0.60
pvalue = 0.2117
Mean firing rate (Hz)

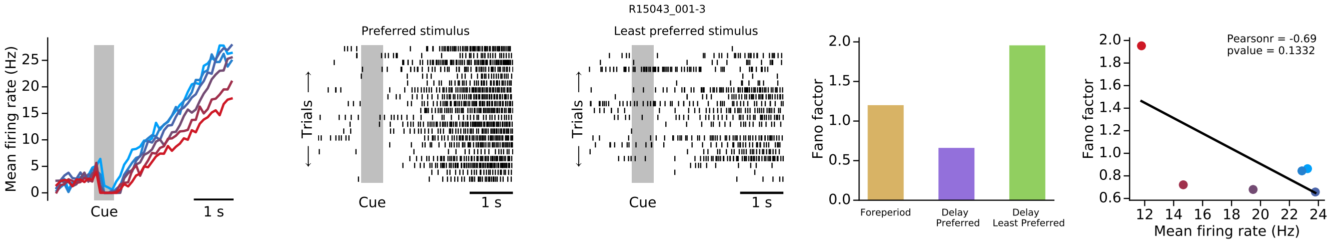
R15043_001-3
Mean firing rate (Hz)
Cue
1 s
Preferred stimulus
Least preferred stimulus
Trials
Fano factor
Foreperiod
Delay Preferred
Delay Least Preferred
Pearsonr = -0.69
pvalue = 0.1332
Mean firing rate (Hz)

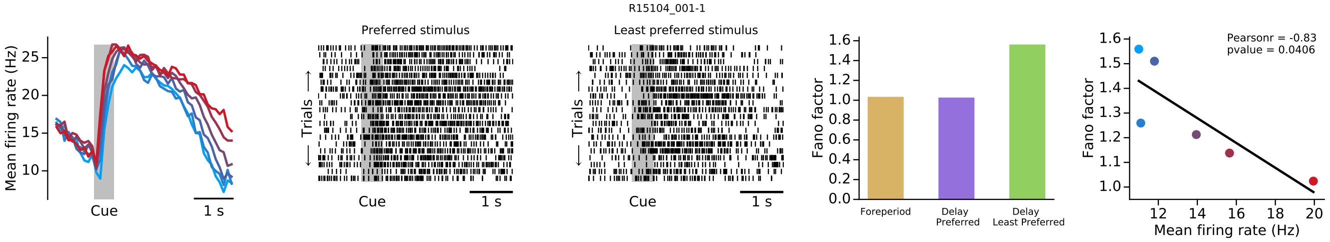
R15104_001-1
Mean firing rate (Hz)
Cue
1 s
Preferred stimulus
Least preferred stimulus
Trials
Fano factor
Foreperiod
Delay Preferred
Delay Least Preferred
Pearsonr = -0.83
pvalue = 0.0406
Mean firing rate (Hz)

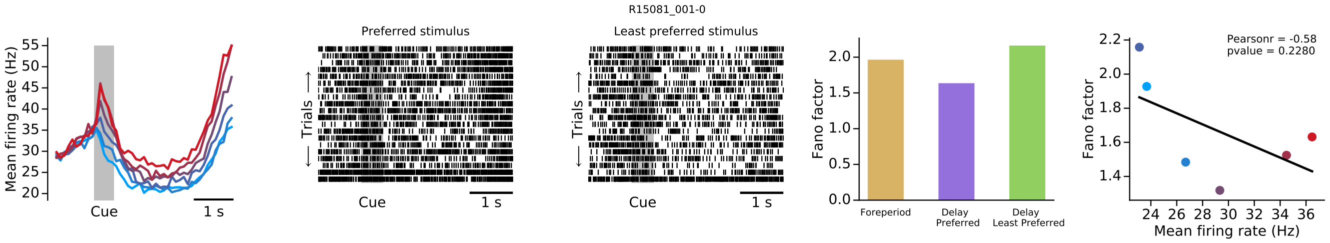
R15081_001-0
Mean firing rate (Hz)
Cue
1 s
Preferred stimulus
Least preferred stimulus
Trials
Fano factor
Foreperiod
Delay Preferred
Delay Least Preferred
Pearsonr = -0.58
pvalue = 0.2280
Mean firing rate (Hz)

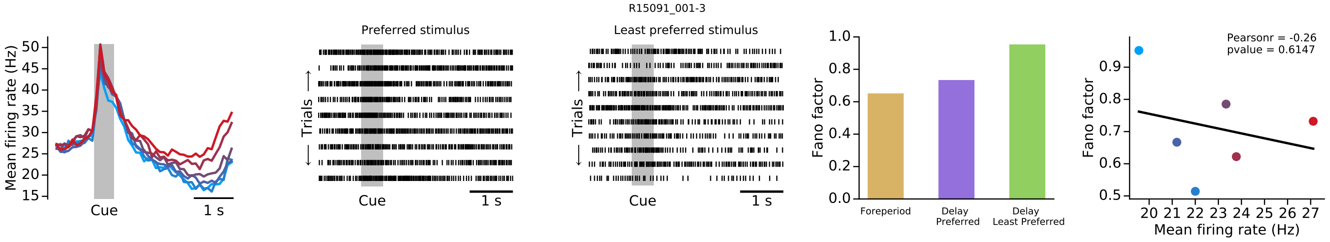
R15091_001-3
Mean firing rate (Hz)
Cue
1 s
Preferred stimulus
Least preferred stimulus
Trials
Fano factor
Foreperiod
Delay Preferred
Delay Least Preferred
Pearsonr = -0.26
pvalue = 0.6147
Mean firing rate (Hz)

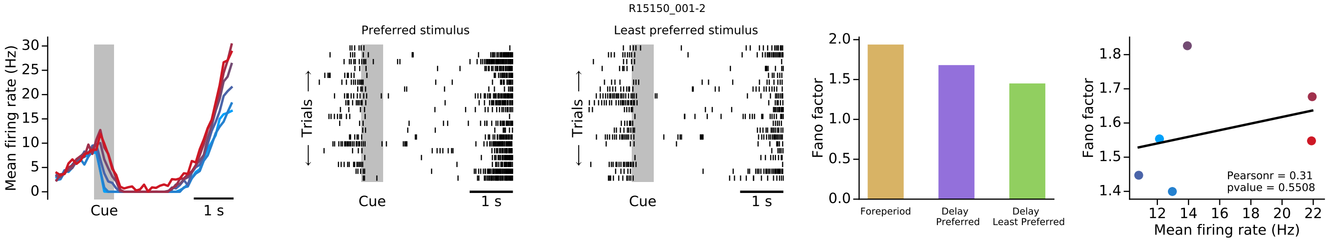
R15150_001-2
Mean firing rate (Hz)
Cue
1 s
Preferred stimulus
Least preferred stimulus
Trials
Fano factor
Foreperiod
Delay Preferred
Delay Least Preferred
Pearsonr = 0.31
pvalue = 0.5508
Mean firing rate (Hz)

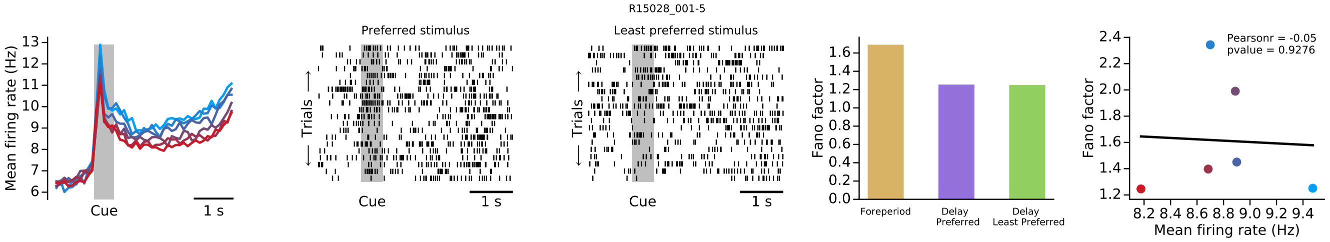
R15028_001-5
Mean firing rate (Hz)
Cue
1 s
Preferred stimulus
Least preferred stimulus
Trials
Fano factor
Foreperiod
Delay Preferred
Delay Least Preferred
Pearsonr = -0.05
pvalue = 0.9276
Mean firing rate (Hz)

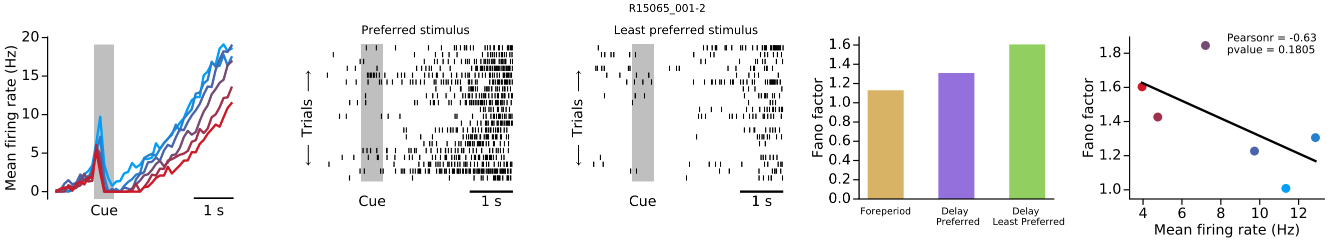
R15065_001-2
Mean firing rate (Hz)
Cue
1 s
Preferred stimulus
Least preferred stimulus
Trials
Fano factor
Foreperiod
Delay Preferred
Delay Least Preferred
Pearsonr = -0.63
pvalue = 0.1805
Mean firing rate (Hz)

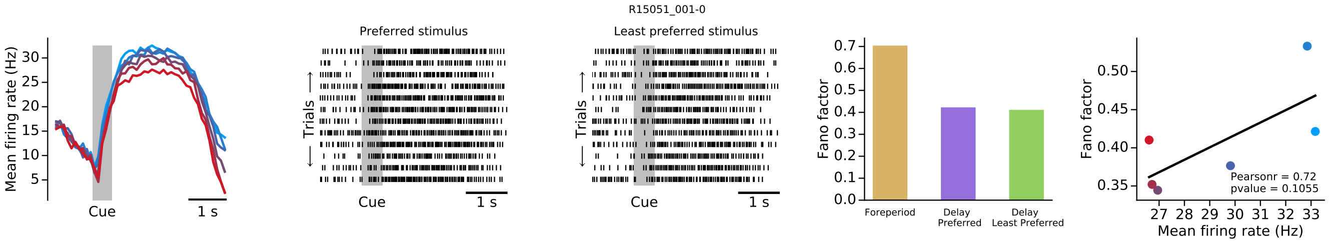
R15051_001-0
Mean firing rate (Hz)
Cue
1 s
Preferred stimulus
Least preferred stimulus
Trials
Fano factor
Foreperiod
Delay Preferred
Delay Least Preferred
Pearsonr = 0.72
pvalue = 0.1055
Mean firing rate (Hz)

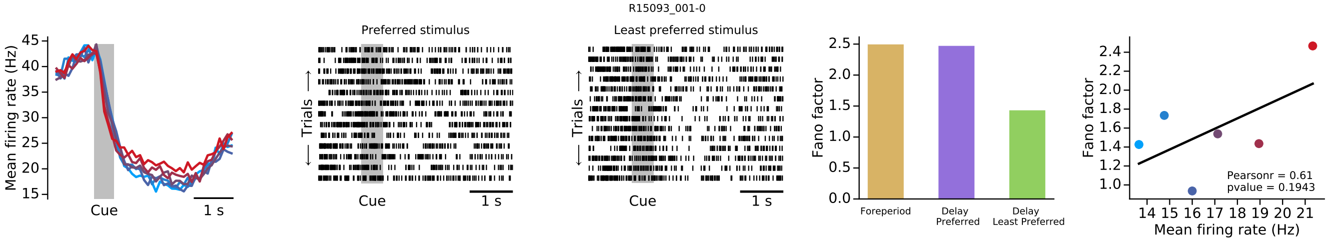
R15093_001-0
Preferred stimulus
Least preferred stimulus
Mean firing rate (Hz)
Trials
Cue
1 s
Fano factor
Foreperiod
Delay Preferred
Delay Least Preferred
Pearsonr = 0.61
pvalue = 0.1943
Mean firing rate (Hz)

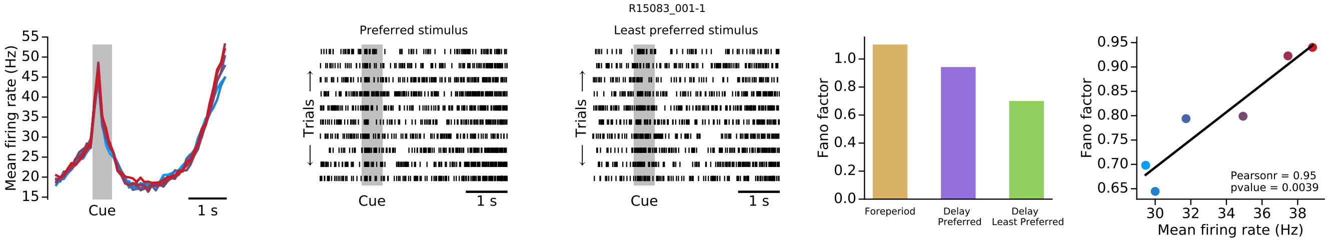
R15083_001-1
Preferred stimulus
Least preferred stimulus
Mean firing rate (Hz)
Trials
Cue
1 s
Fano factor
Foreperiod
Delay Preferred
Delay Least Preferred
Pearsonr = 0.95
pvalue = 0.0039
Mean firing rate (Hz)

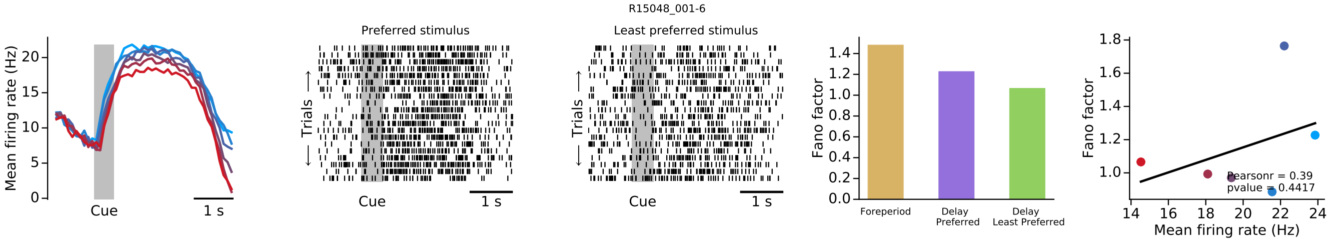
R15048_001-6
Preferred stimulus
Least preferred stimulus
Mean firing rate (Hz)
Trials
Cue
1 s
Fano factor
Foreperiod
Delay Preferred
Delay Least Preferred
Pearsonr = 0.39
pvalue = 0.4417
Mean firing rate (Hz)

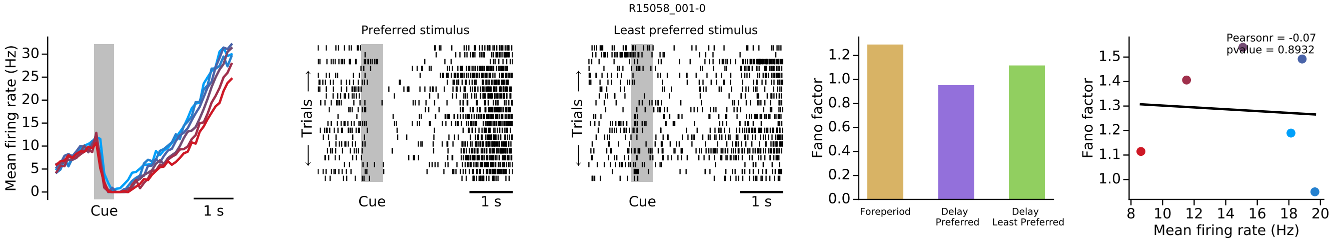
R15058_001-0
Preferred stimulus
Least preferred stimulus
Mean firing rate (Hz)
Trials
Cue
1 s
Fano factor
Foreperiod
Delay Preferred
Delay Least Preferred
Pearsonr = -0.07
pvalue = 0.8932
Mean firing rate (Hz)

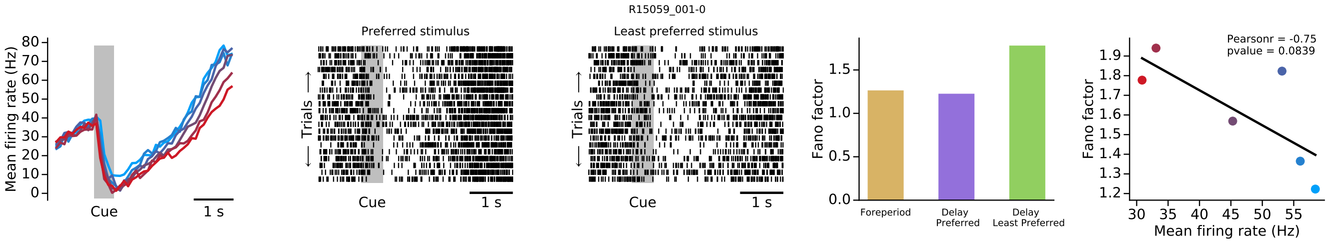
R15059_001-0
Preferred stimulus
Least preferred stimulus
Mean firing rate (Hz)
Trials
Cue
1 s
Fano factor
Foreperiod
Delay Preferred
Delay Least Preferred
Pearsonr = -0.75
pvalue = 0.0839
Mean firing rate (Hz)

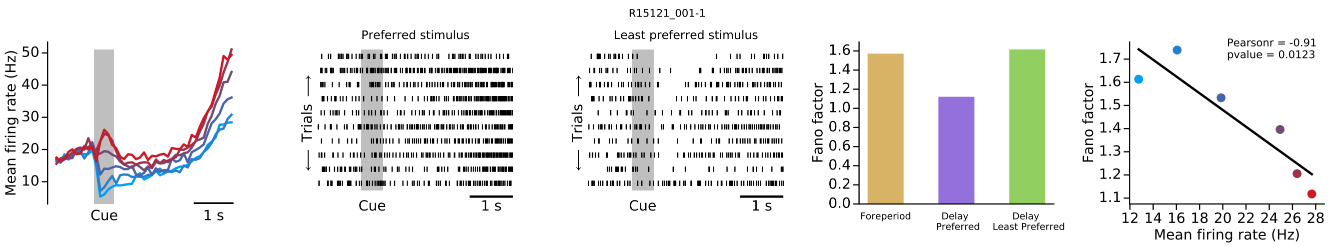
R15121_001-1
Preferred stimulus
Least preferred stimulus
Mean firing rate (Hz)
Cue
1 s
Trials
Fano factor
Foreperiod
Delay Preferred
Delay Least Preferred
Pearsonr = -0.91
pvalue = 0.0123
Mean firing rate (Hz)

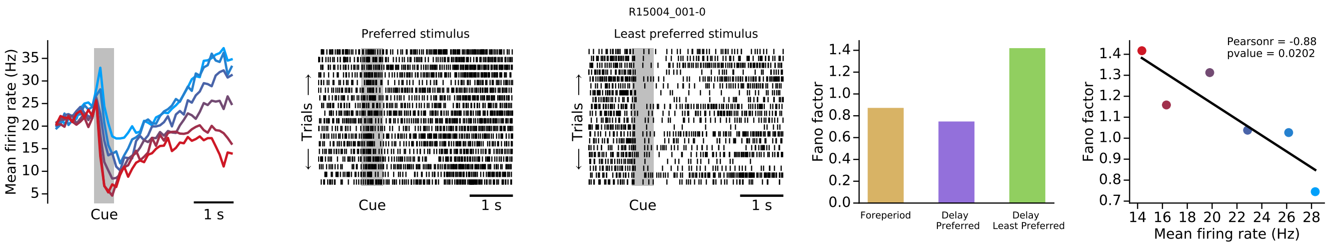
R15004_001-0
Preferred stimulus
Least preferred stimulus
Mean firing rate (Hz)
Cue
1 s
Trials
Fano factor
Foreperiod
Delay Preferred
Delay Least Preferred
Pearsonr = -0.88
pvalue = 0.0202
Mean firing rate (Hz)

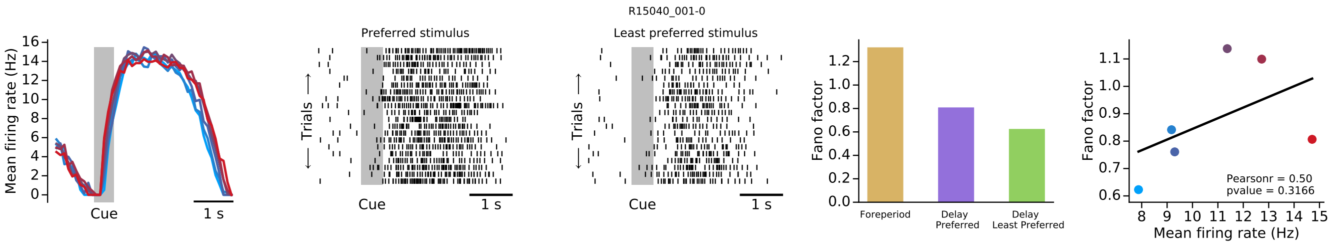
R15040_001-0
Preferred stimulus
Least preferred stimulus
Mean firing rate (Hz)
Cue
1 s
Trials
Fano factor
Foreperiod
Delay Preferred
Delay Least Preferred
Pearsonr = 0.50
pvalue = 0.3166
Mean firing rate (Hz)

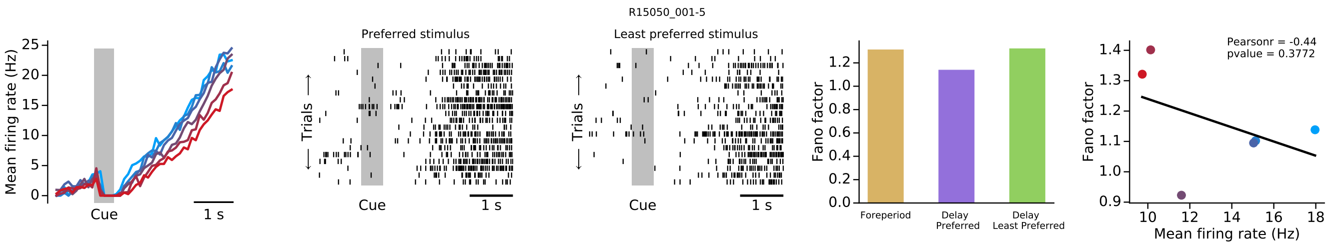
R15050_001-5
Preferred stimulus
Least preferred stimulus
Mean firing rate (Hz)
Cue
1 s
Trials
Fano factor
Foreperiod
Delay Preferred
Delay Least Preferred
Pearsonr = -0.44
pvalue = 0.3772
Mean firing rate (Hz)

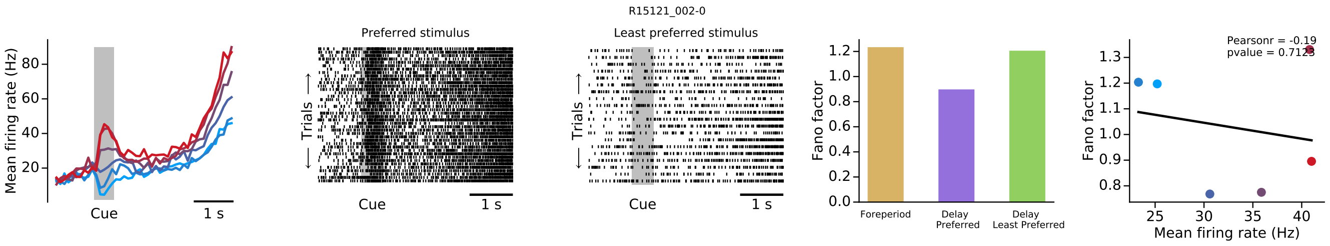
R15121_002-0
Preferred stimulus
Least preferred stimulus
Mean firing rate (Hz)
Cue
1 s
Trials
Fano factor
Foreperiod
Delay Preferred
Delay Least Preferred
Pearsonr = -0.19
pvalue = 0.7123
Mean firing rate (Hz)

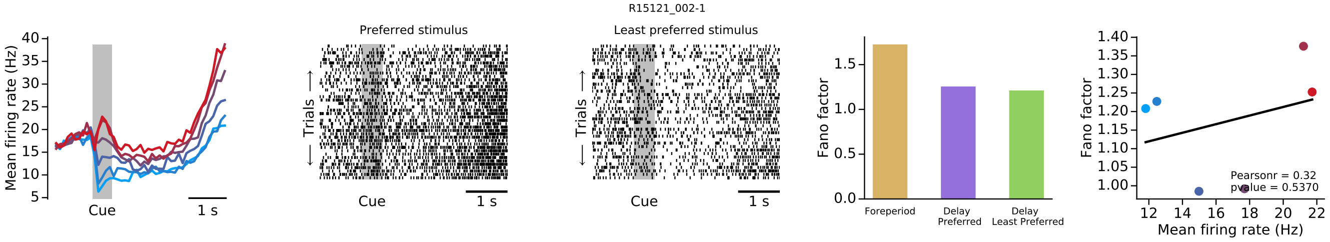
R15121_002-1
Mean firing rate (Hz)
Cue
1 s
Preferred stimulus
Least preferred stimulus
Trials
Fano factor
Foreperiod
Delay Preferred
Delay Least Preferred
Pearsonr = 0.32
pvalue = 0.5370
Mean firing rate (Hz)

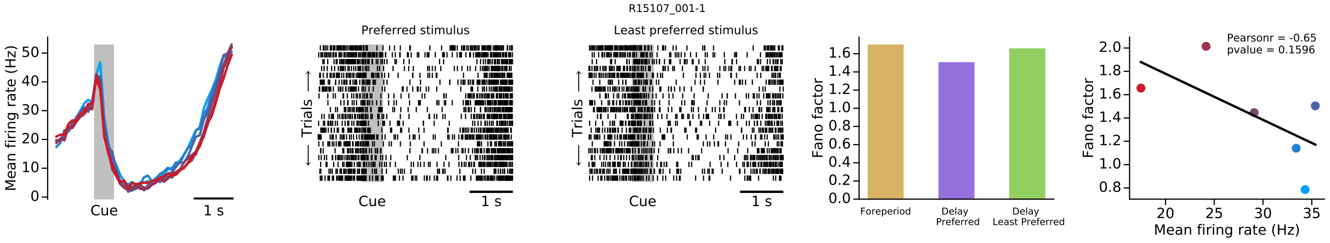
R15107_001-1
Mean firing rate (Hz)
Cue
1 s
Preferred stimulus
Least preferred stimulus
Trials
Fano factor
Foreperiod
Delay Preferred
Delay Least Preferred
Pearsonr = -0.65
pvalue = 0.1596
Mean firing rate (Hz)

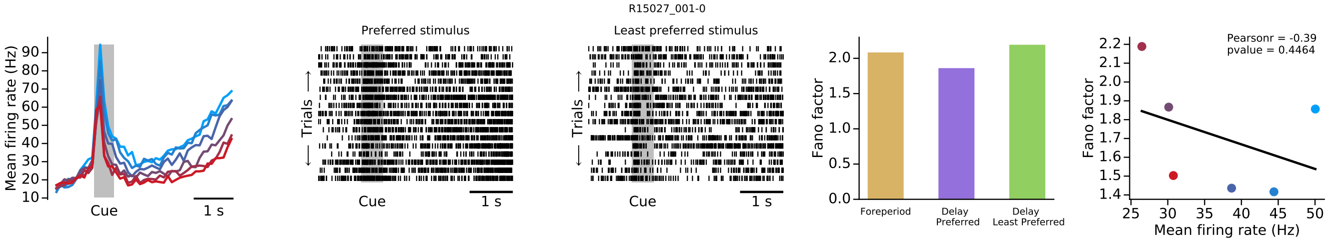
R15027_001-0
Mean firing rate (Hz)
Cue
1 s
Preferred stimulus
Least preferred stimulus
Trials
Fano factor
Foreperiod
Delay Preferred
Delay Least Preferred
Pearsonr = -0.39
pvalue = 0.4464
Mean firing rate (Hz)

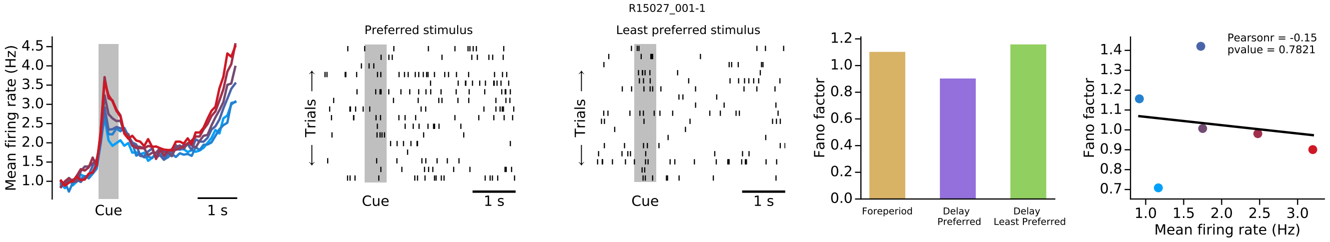
R15027_001-1
Mean firing rate (Hz)
Cue
1 s
Preferred stimulus
Least preferred stimulus
Trials
Fano factor
Foreperiod
Delay Preferred
Delay Least Preferred
Pearsonr = -0.15
pvalue = 0.7821
Mean firing rate (Hz)

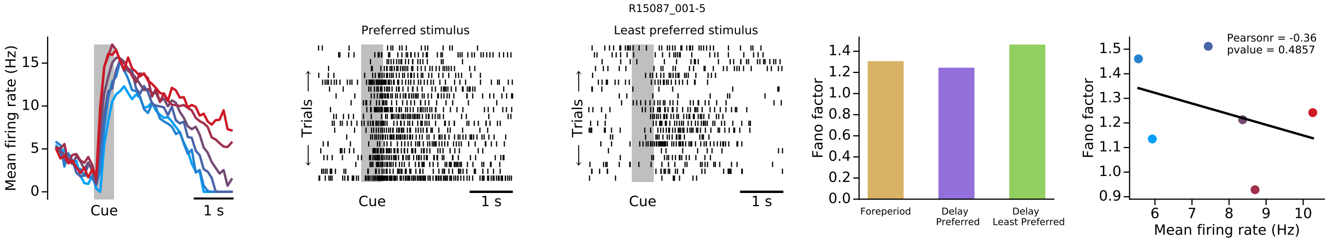
R15087_001-5
Mean firing rate (Hz)
Cue
1 s
Preferred stimulus
Least preferred stimulus
Trials
Fano factor
Foreperiod
Delay Preferred
Delay Least Preferred
Pearsonr = -0.36
pvalue = 0.4857
Mean firing rate (Hz)

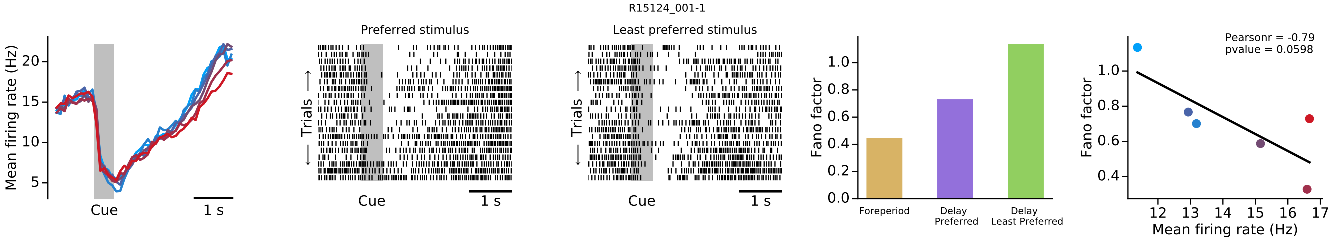
R15124_001-1
Mean firing rate (Hz)
Cue
1 s
Preferred stimulus
Least preferred stimulus
Trials
Fano factor
Foreperiod
Delay Preferred
Delay Least Preferred
Pearsonr = -0.79
pvalue = 0.0598
Mean firing rate (Hz)

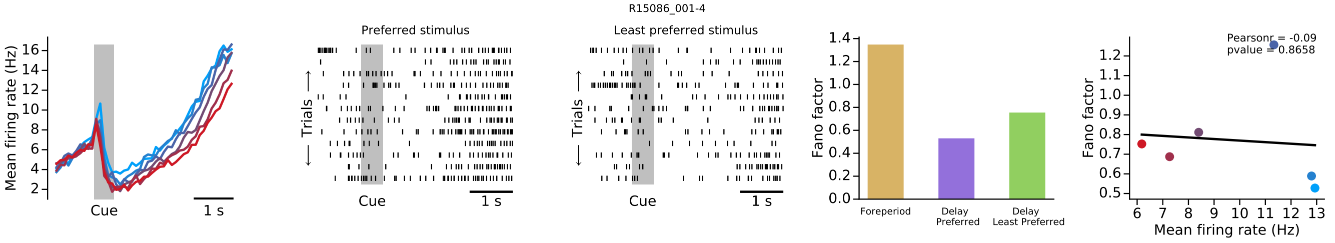
R15086_001-4
Mean firing rate (Hz)
Cue
1 s
Preferred stimulus
Least preferred stimulus
Trials
Fano factor
Foreperiod
Delay Preferred
Delay Least Preferred
Pearsonr = -0.09
pvalue = 0.8658
Mean firing rate (Hz)

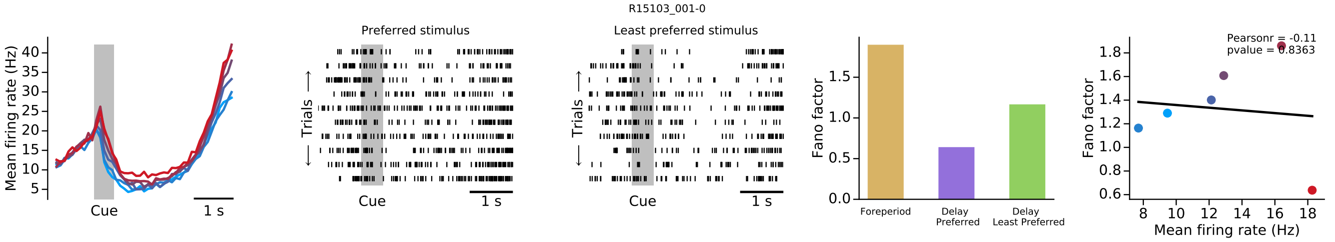
R15103_001-0
Mean firing rate (Hz)
Cue
1 s
Preferred stimulus
Least preferred stimulus
Trials
Fano factor
Foreperiod
Delay Preferred
Delay Least Preferred
Pearsonr = -0.11
pvalue = 0.8363
Mean firing rate (Hz)

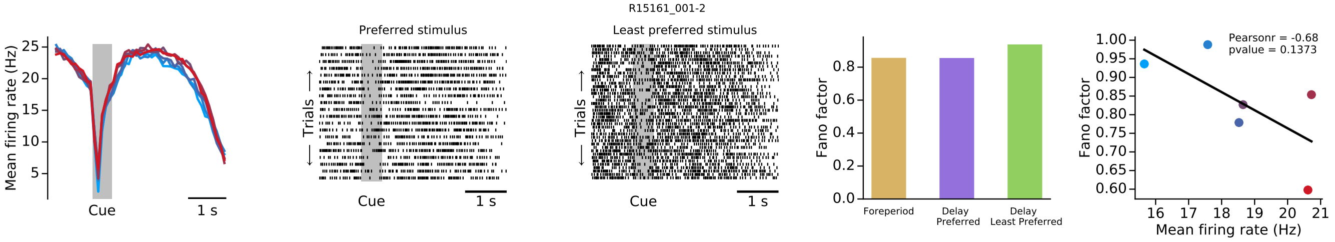
R15161_001-2
Mean firing rate (Hz)
Cue
1 s
Preferred stimulus
Least preferred stimulus
Trials
Fano factor
Foreperiod
Delay Preferred
Delay Least Preferred
Pearsonr = -0.68
pvalue = 0.1373
Mean firing rate (Hz)

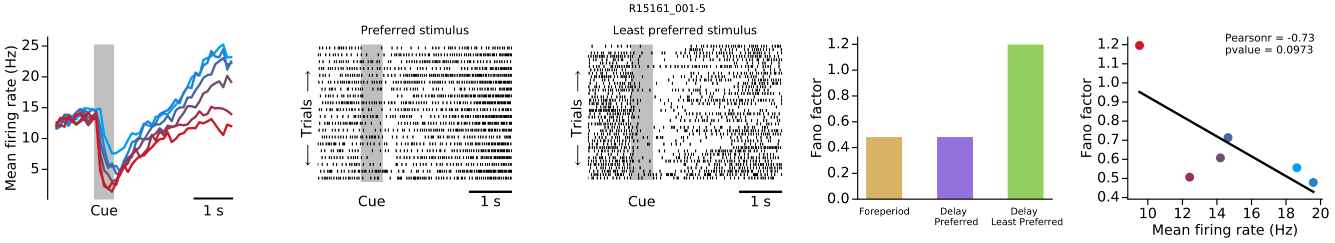
R15161_001-5
Mean firing rate (Hz)
Cue
1 s
Preferred stimulus
Least preferred stimulus
Trials
Fano factor
Foreperiod
Delay Preferred
Delay Least Preferred
Pearsonr = -0.73
pvalue = 0.0973
Mean firing rate (Hz)

R15054_001-1

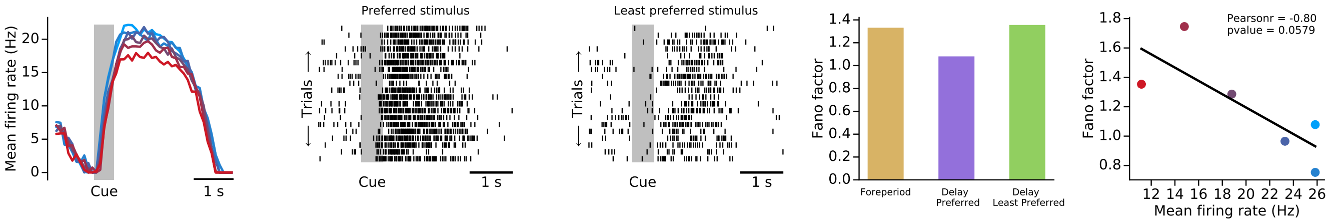


R15026_001-0

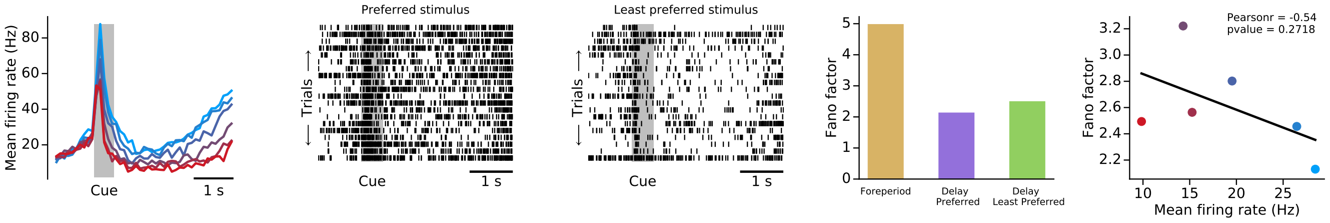


R15108_001-1

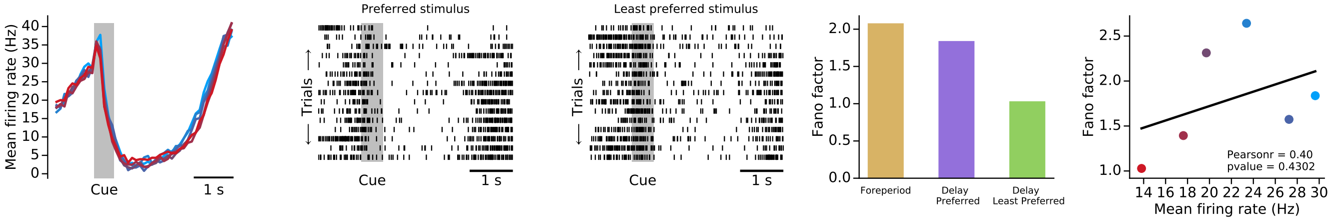


R15070_001-1

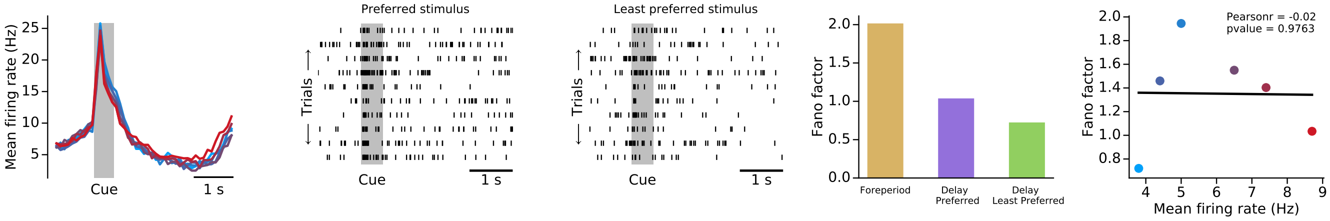


R15060_001-1

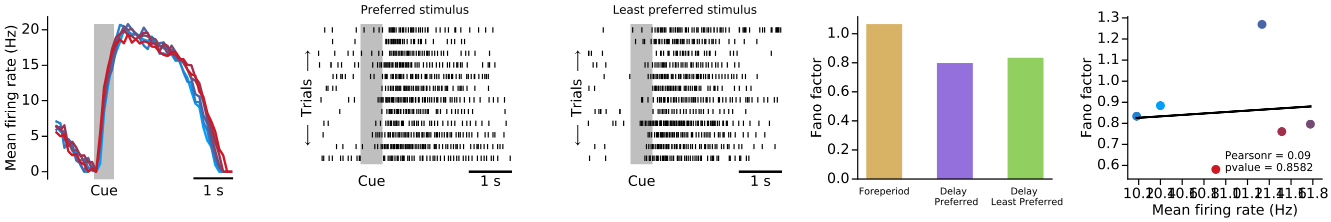

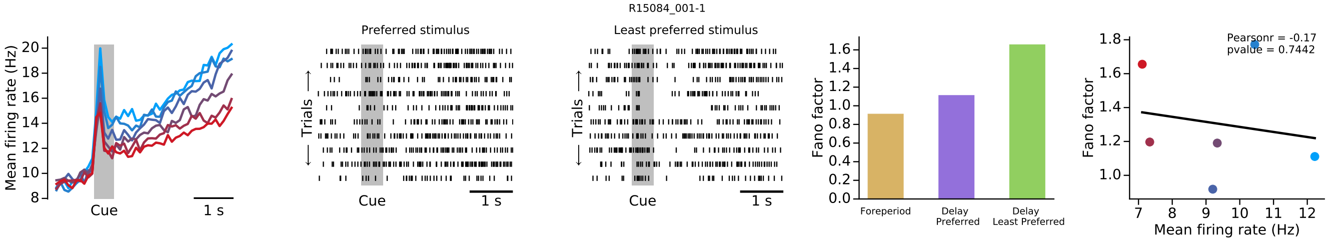
R15084_001-1
Mean firing rate (Hz)
Cue
1 s
Preferred stimulus
Least preferred stimulus
Trials
Fano factor
Foreperiod
Delay Preferred
Delay Least Preferred
Pearsonr = -0.17
pvalue = 0.7442
Mean firing rate (Hz)

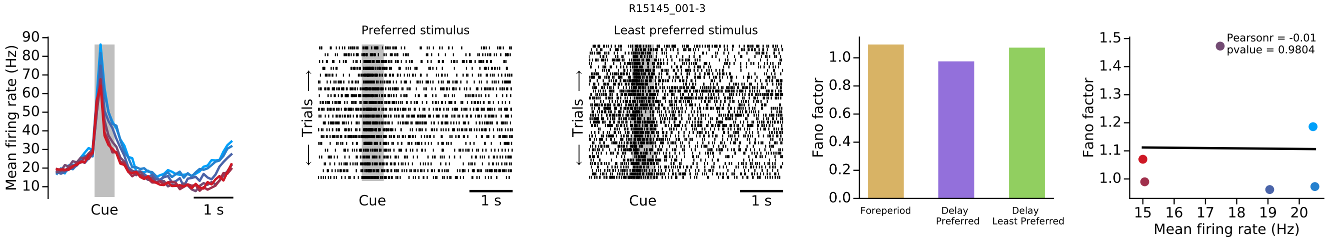
R15145_001-3
Mean firing rate (Hz)
Cue
1 s
Preferred stimulus
Least preferred stimulus
Trials
Fano factor
Foreperiod
Delay Preferred
Delay Least Preferred
Pearsonr = -0.01
pvalue = 0.9804
Mean firing rate (Hz)

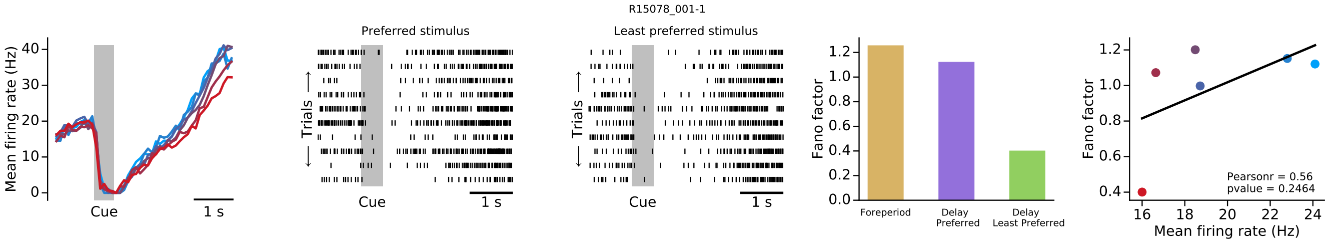
R15078_001-1
Mean firing rate (Hz)
Cue
1 s
Preferred stimulus
Least preferred stimulus
Trials
Fano factor
Foreperiod
Delay Preferred
Delay Least Preferred
Pearsonr = 0.56
pvalue = 0.2464
Mean firing rate (Hz)

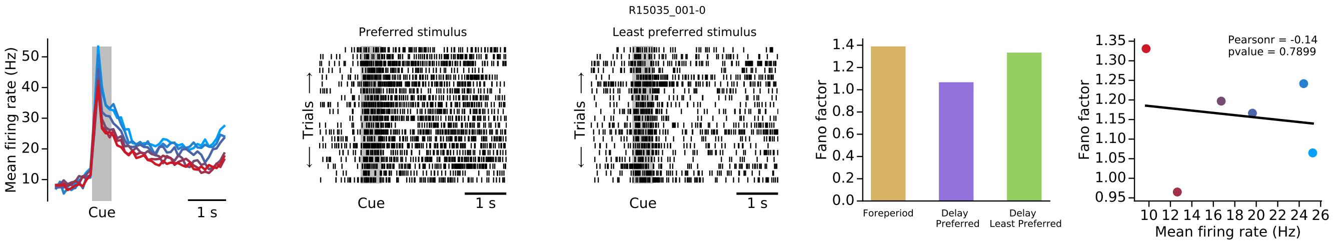
R15035_001-0
Mean firing rate (Hz)
Cue
1 s
Preferred stimulus
Least preferred stimulus
Trials
Fano factor
Foreperiod
Delay Preferred
Delay Least Preferred
Pearsonr = -0.14
pvalue = 0.7899
Mean firing rate (Hz)

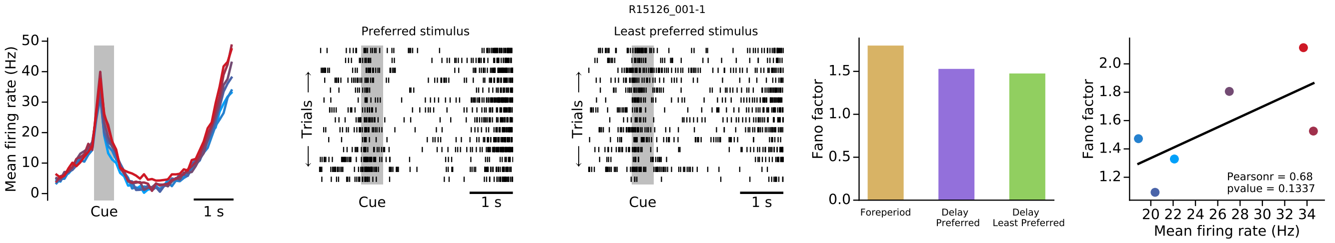
R15126_001-1
Mean firing rate (Hz)
Cue
1 s
Preferred stimulus
Least preferred stimulus
Trials
Fano factor
Foreperiod
Delay Preferred
Delay Least Preferred
Pearsonr = 0.68
pvalue = 0.1337
Mean firing rate (Hz)

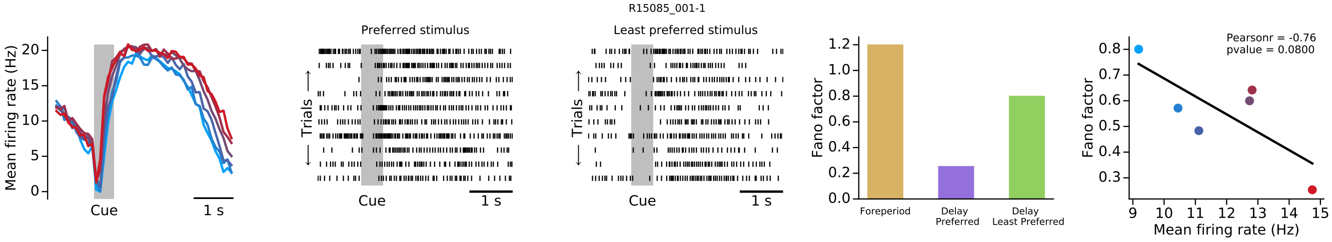
R15085_001-1
Mean firing rate (Hz)
Cue
1 s
Preferred stimulus
Trials
Least preferred stimulus
Fano factor
Foreperiod
Delay Preferred
Delay Least Preferred
Pearsonr = -0.76
pvalue = 0.0800
Mean firing rate (Hz)

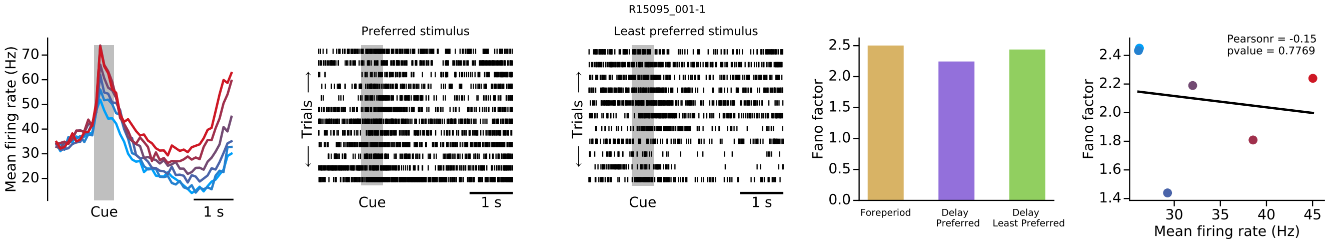
R15095_001-1
Mean firing rate (Hz)
Cue
1 s
Preferred stimulus
Trials
Least preferred stimulus
Fano factor
Foreperiod
Delay Preferred
Delay Least Preferred
Pearsonr = -0.15
pvalue = 0.7769
Mean firing rate (Hz)

### MNM NEURONS

#### Group 1

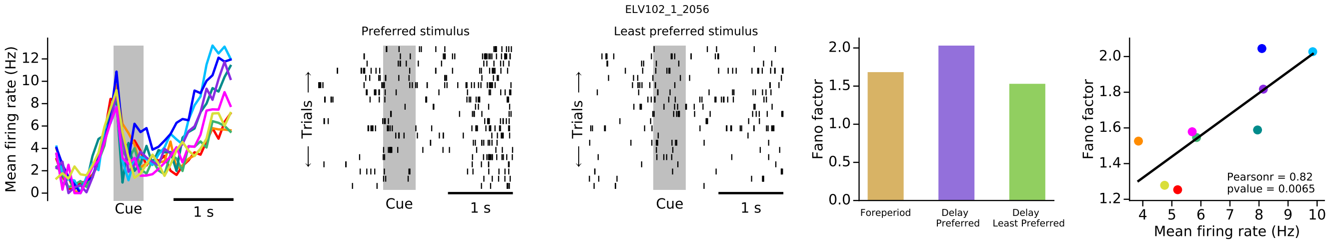


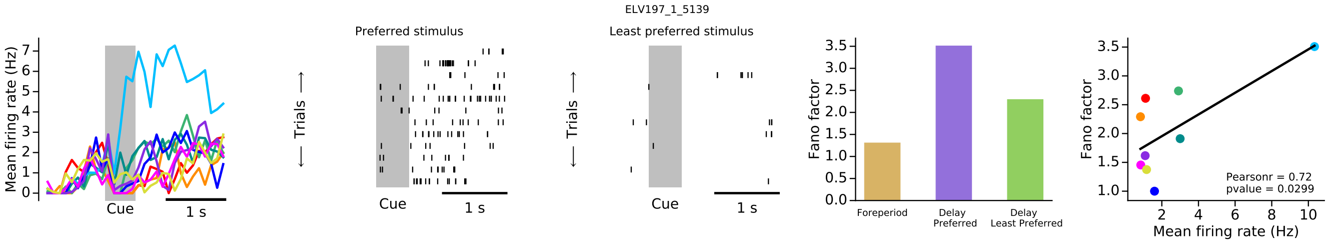


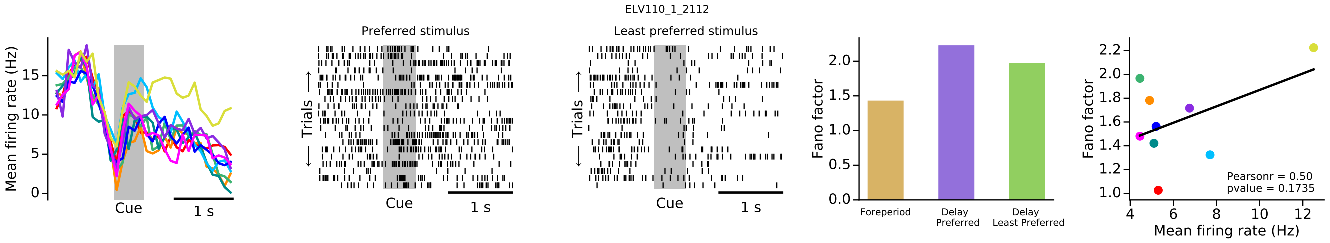


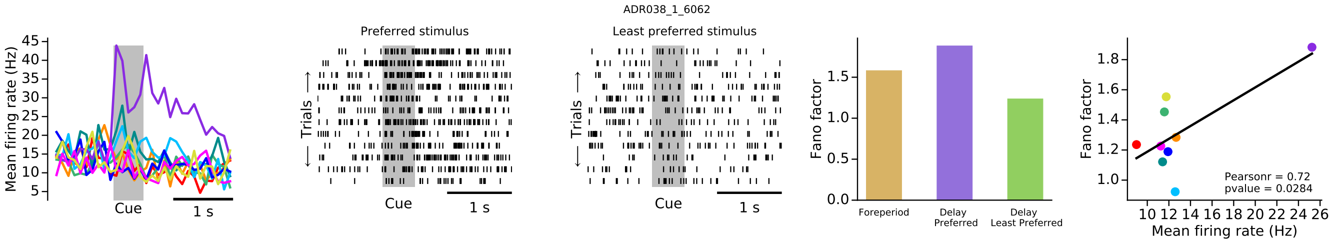

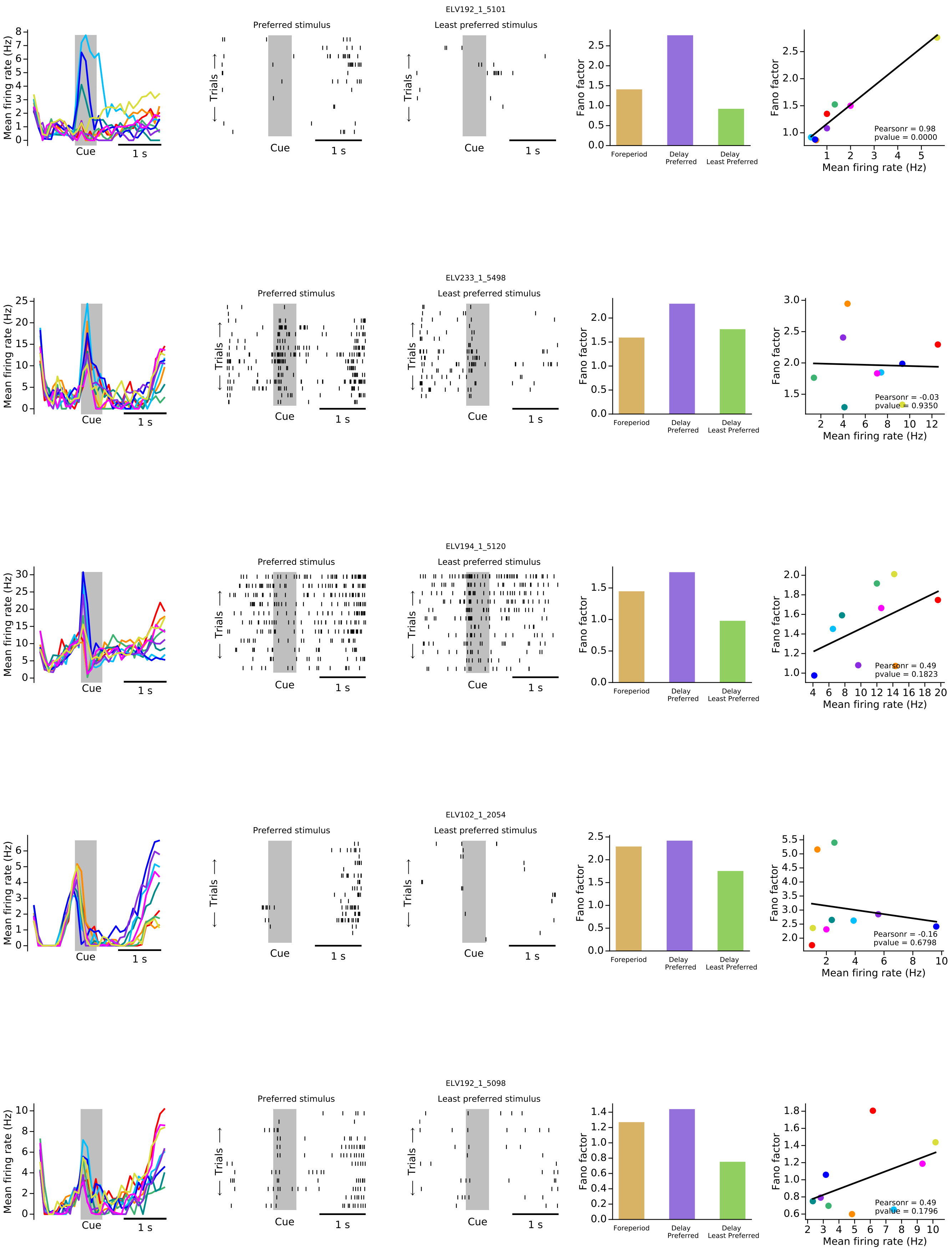
ELV192_1_5101
Preferred stimulus
Least preferred stimulus
Mean firing rate (Hz)
Trials
Cue
1 s
Fano factor
Foreperiod
Delay Preferred
Delay Least Preferred
Pearsonr = 0.98
pvalue = 0.0000
Mean firing rate (Hz)
ELV233_1_5498
Preferred stimulus
Least preferred stimulus
Pearsonr = -0.03
pvalue = 0.9350
ELV194_1_5120
Preferred stimulus
Least preferred stimulus
Pearsonr = 0.49
pvalue = 0.1823
ELV102_1_2054
Preferred stimulus
Least preferred stimulus
Pearsonr = -0.16
pvalue = 0.6798
ELV192_1_5098
Preferred stimulus
Least preferred stimulus
Pearsonr = 0.49
pvalue = 0.1796

ELV194_1_5118

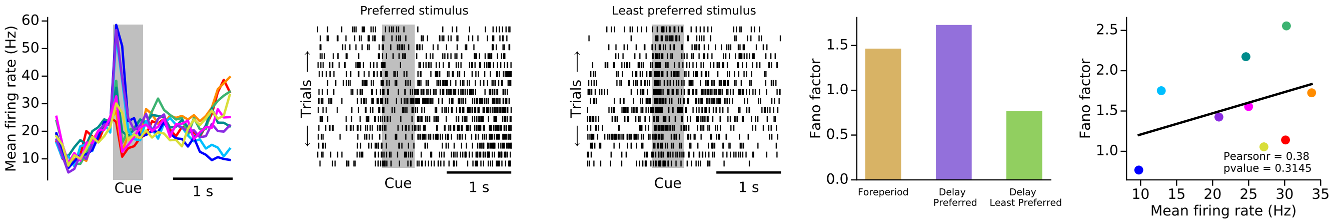


ELV198_1_5149

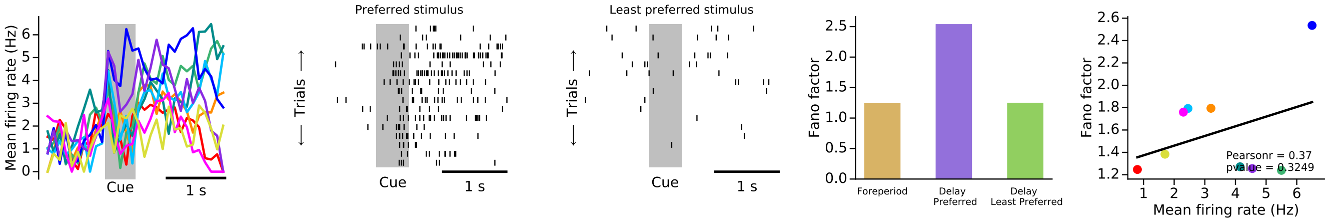


ELV123_1_2181

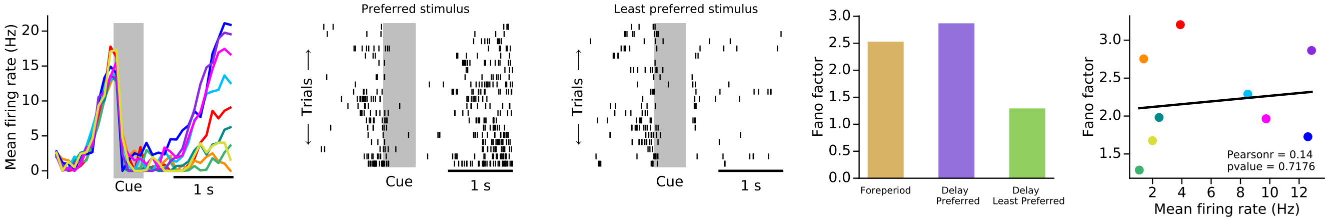


ELV192_1_5105

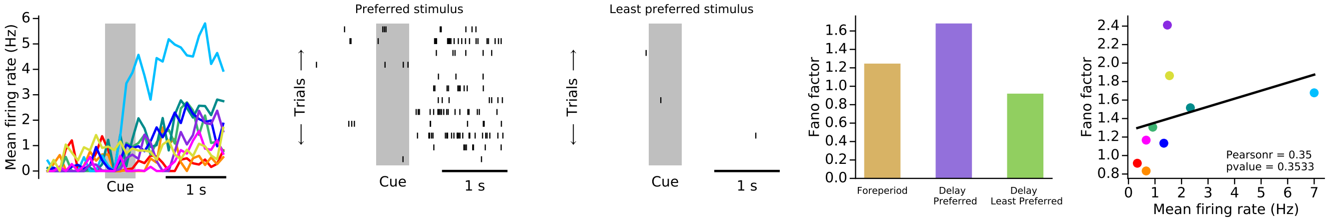


ELV130_1_2240

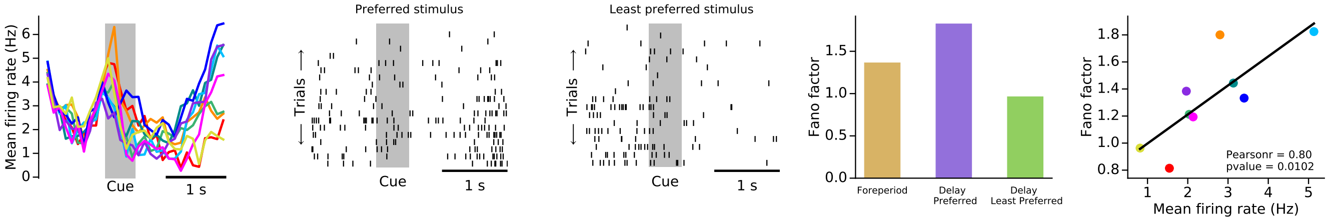

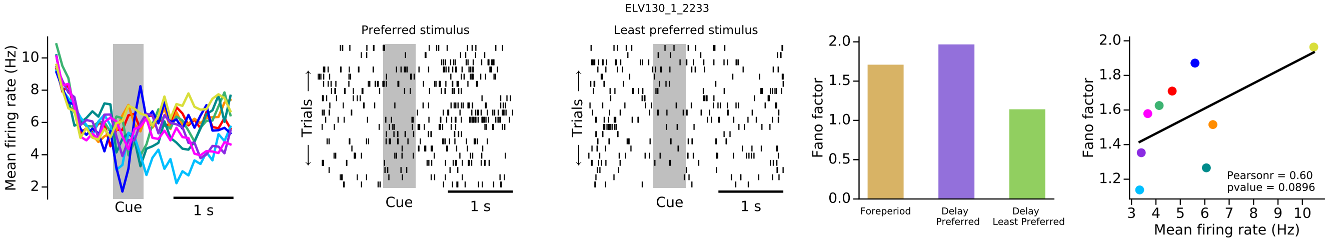
ELV130_1_2233
Preferred stimulus
Least preferred stimulus
Mean firing rate (Hz)
Trials
Cue
1 s
Fano factor
Foreperiod
Delay Preferred
Delay Least Preferred
Pearsonr = 0.60
pvalue = 0.0896
Mean firing rate (Hz)

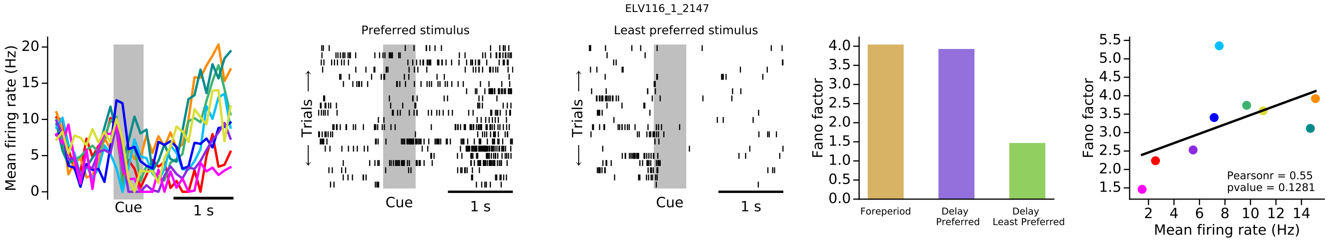
ELV116_1_2147
Preferred stimulus
Least preferred stimulus
Mean firing rate (Hz)
Trials
Cue
1 s
Fano factor
Foreperiod
Delay Preferred
Delay Least Preferred
Pearsonr = 0.55
pvalue = 0.1281
Mean firing rate (Hz)

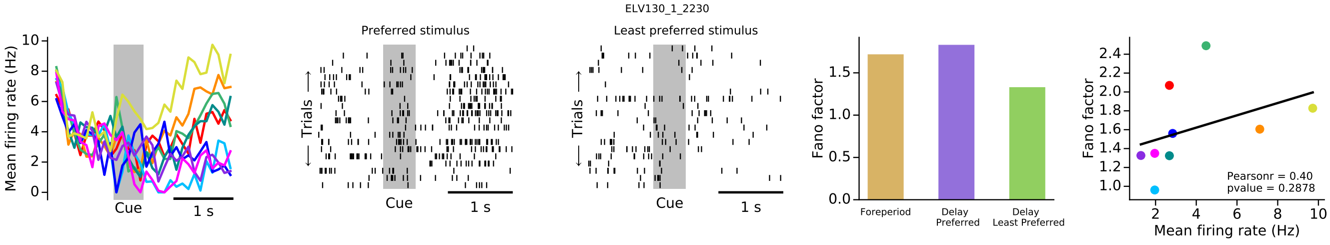
ELV130_1_2230
Preferred stimulus
Least preferred stimulus
Mean firing rate (Hz)
Trials
Cue
1 s
Fano factor
Foreperiod
Delay Preferred
Delay Least Preferred
Pearsonr = 0.40
pvalue = 0.2878
Mean firing rate (Hz)

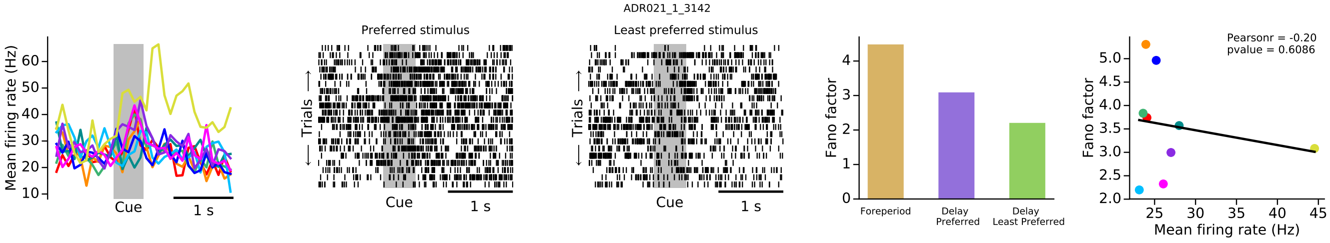
ADR021_1_3142
Preferred stimulus
Least preferred stimulus
Mean firing rate (Hz)
Trials
Cue
1 s
Fano factor
Foreperiod
Delay Preferred
Delay Least Preferred
Pearsonr = -0.20
pvalue = 0.6086
Mean firing rate (Hz)

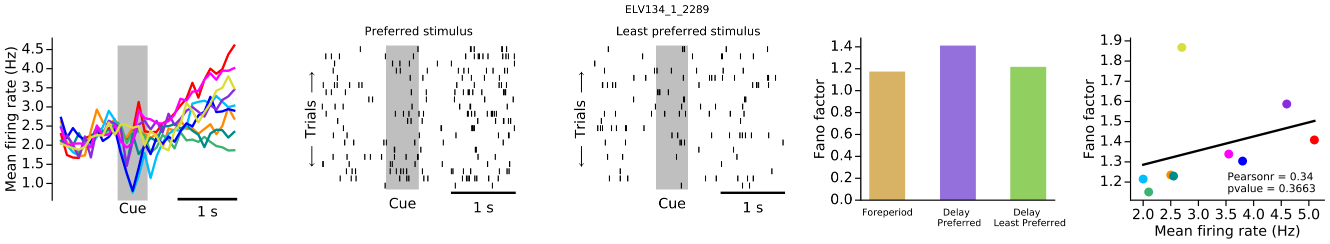
ELV134_1_2289
Preferred stimulus
Least preferred stimulus
Mean firing rate (Hz)
Trials
Cue
1 s
Fano factor
Foreperiod
Delay Preferred
Delay Least Preferred
Pearsonr = 0.34
pvalue = 0.3663
Mean firing rate (Hz)

ELV197_1_5144

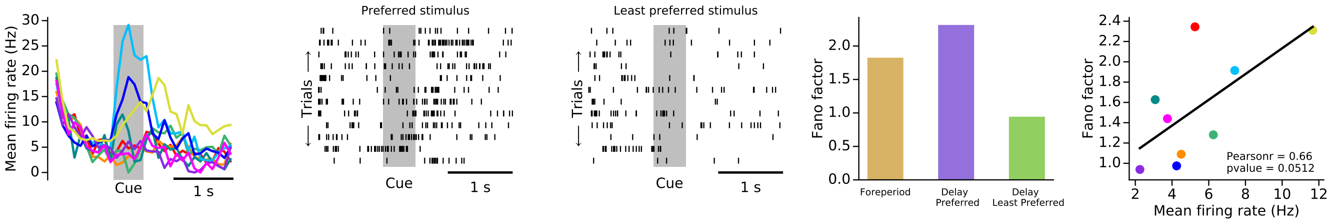


ELV196_1_5129

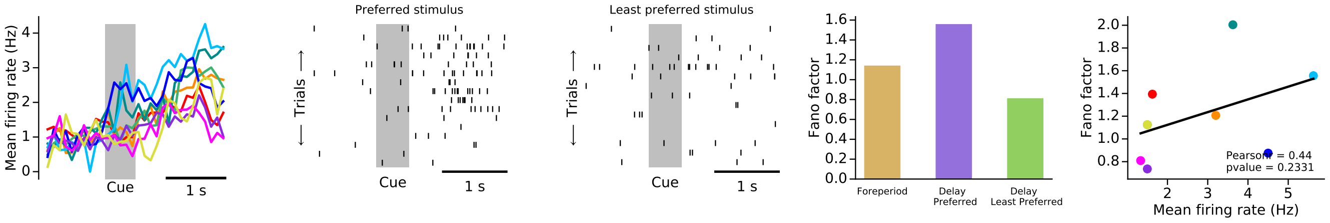


ELV224_1_5428

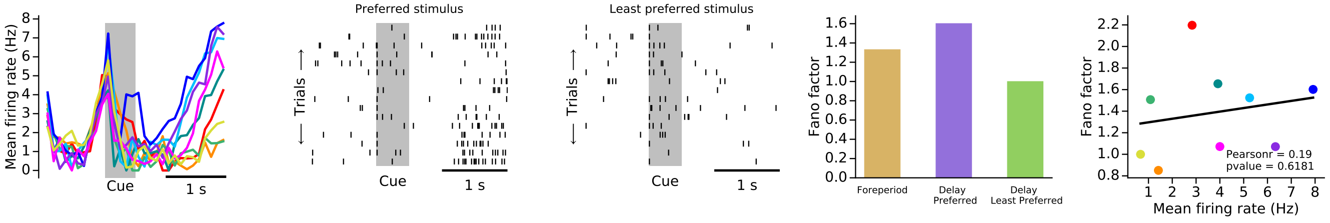


ADR042_1_6099

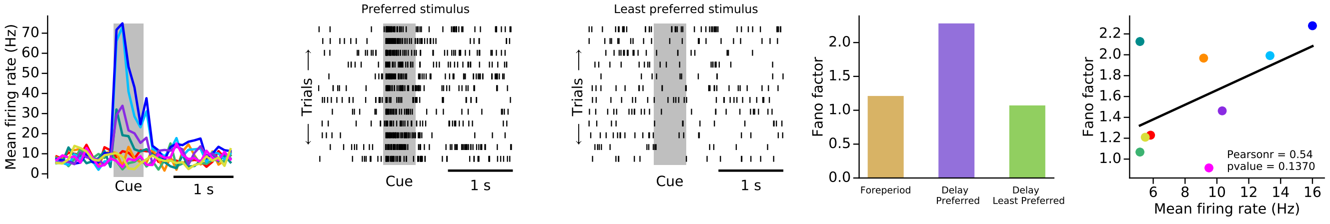


ELV222_1_5392

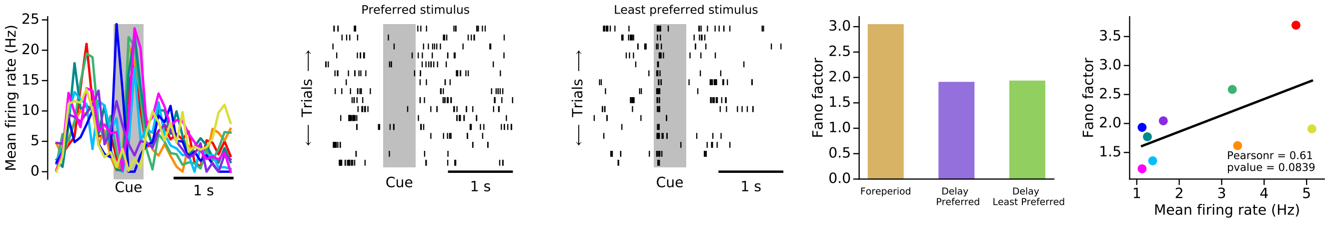

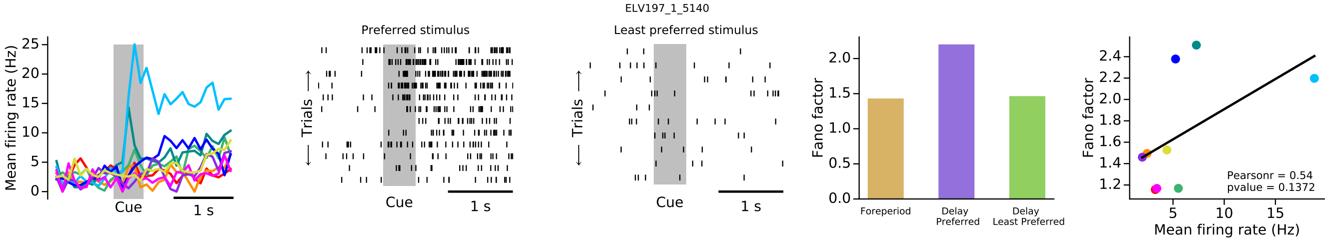
ELV197_1_5140
Preferred stimulus
Least preferred stimulus
Mean firing rate (Hz)
Trials
Cue
1 s
Fano factor
Foreperiod
Delay Preferred
Delay Least Preferred
Pearsonr = 0.54
pvalue = 0.1372
Mean firing rate (Hz)

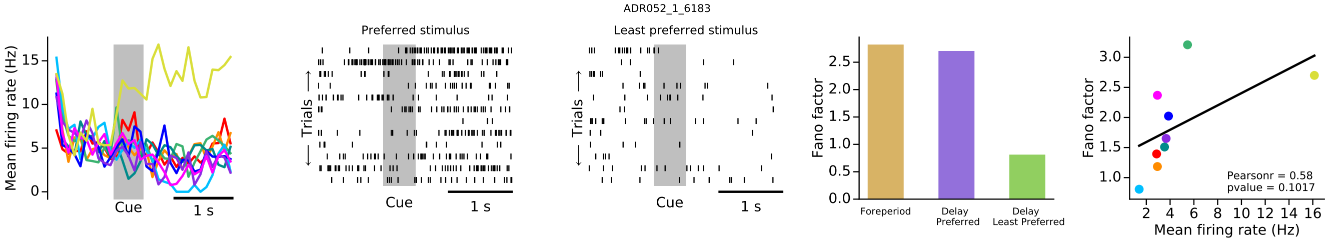
ADR052_1_6183
Preferred stimulus
Least preferred stimulus
Mean firing rate (Hz)
Trials
Cue
1 s
Fano factor
Foreperiod
Delay Preferred
Delay Least Preferred
Pearsonr = 0.58
pvalue = 0.1017
Mean firing rate (Hz)

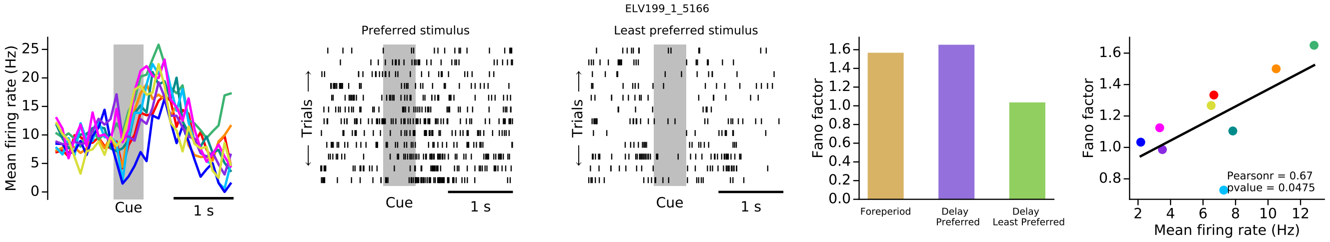
ELV199_1_5166
Preferred stimulus
Least preferred stimulus
Mean firing rate (Hz)
Trials
Cue
1 s
Fano factor
Foreperiod
Delay Preferred
Delay Least Preferred
Pearsonr = 0.67
pvalue = 0.0475
Mean firing rate (Hz)

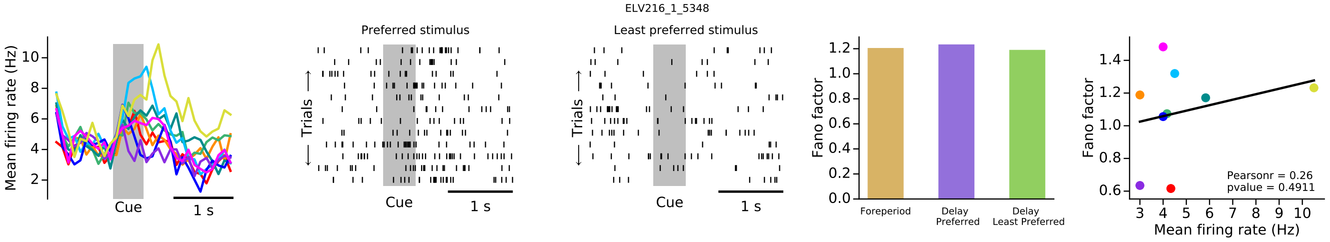
ELV216_1_5348
Preferred stimulus
Least preferred stimulus
Mean firing rate (Hz)
Trials
Cue
1 s
Fano factor
Foreperiod
Delay Preferred
Delay Least Preferred
Pearsonr = 0.26
pvalue = 0.4911
Mean firing rate (Hz)

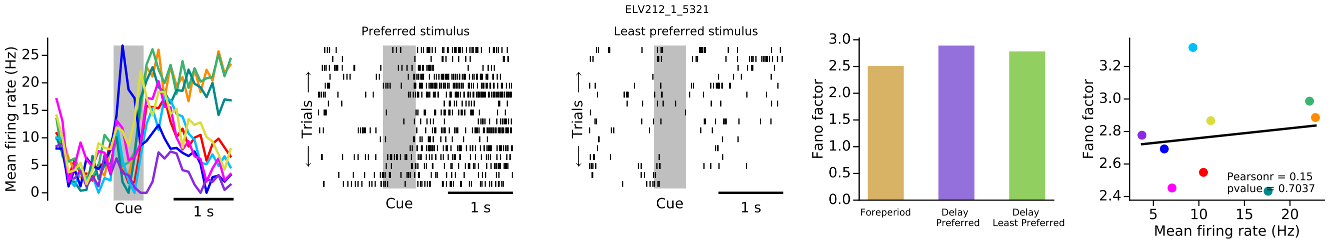
ELV212_1_5321
Preferred stimulus
Least preferred stimulus
Mean firing rate (Hz)
Trials
Cue
1 s
Fano factor
Foreperiod
Delay Preferred
Delay Least Preferred
Pearsonr = 0.15
pvalue = 0.7037
Mean firing rate (Hz)

ELV215_1_5420

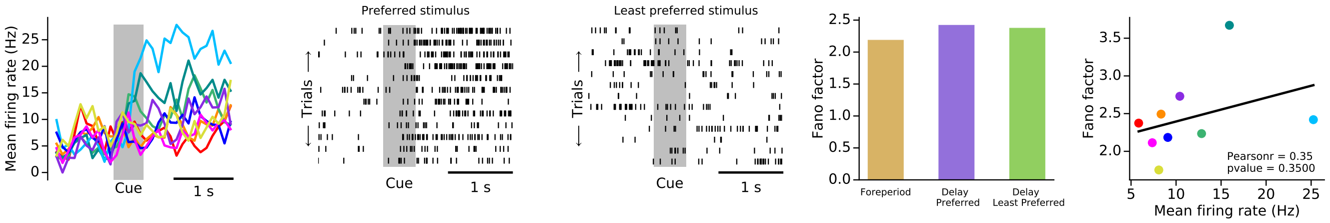


ADR032_1_6024

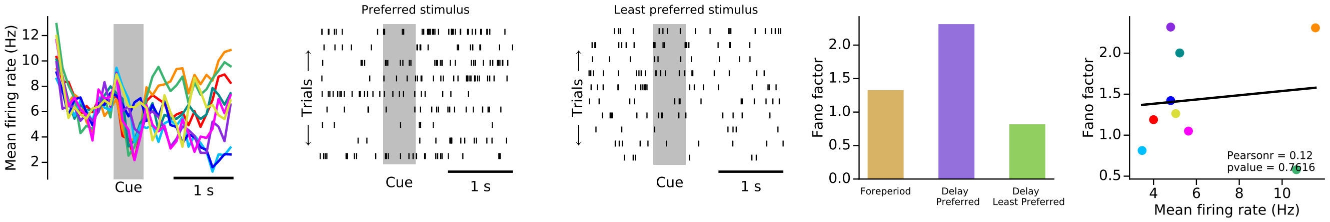


ELV197_1_5136

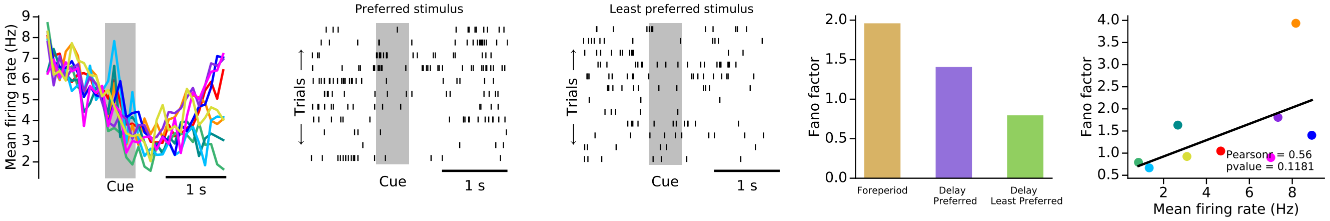


ELV194_1_5112

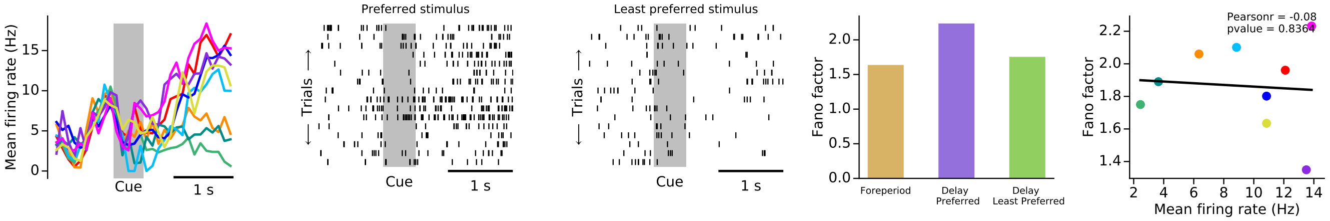


ELV198_1_5155

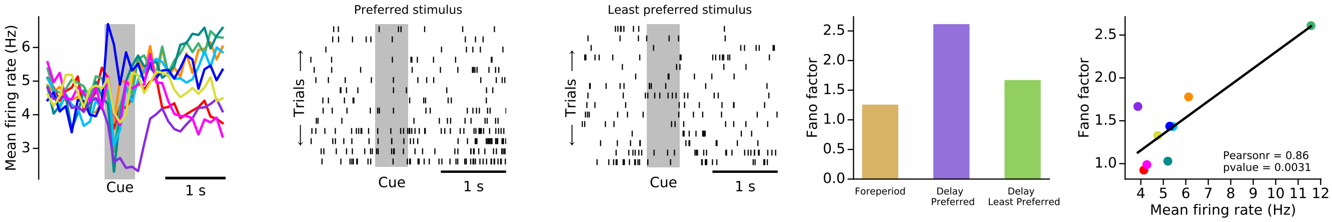

#### Group 2

ADR045_1_6130

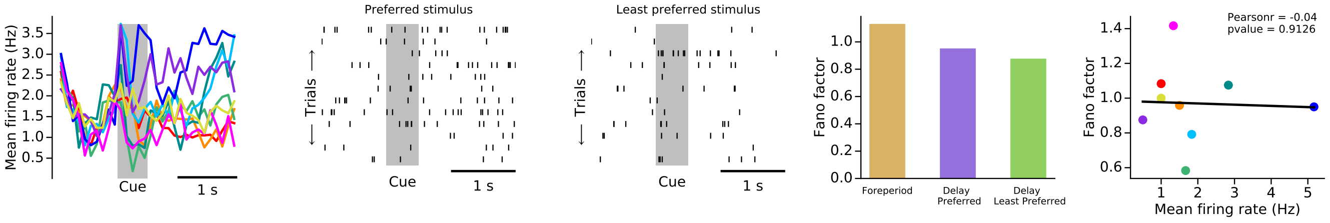


ELV191_1_5084

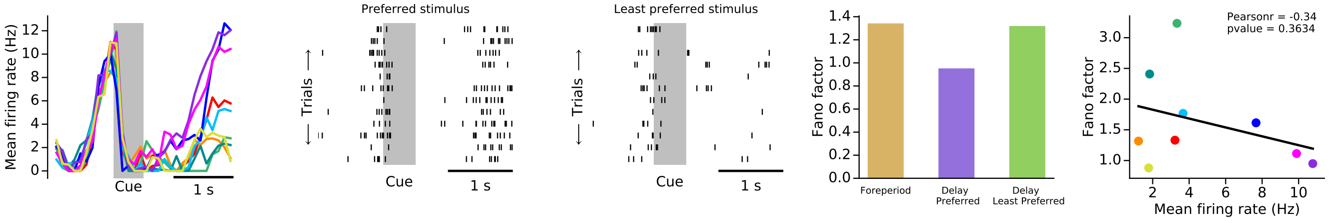


ELV192_1_5088

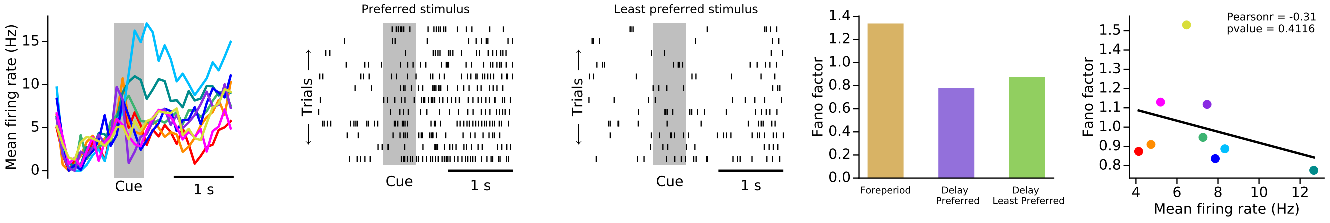


ELV194_1_5121

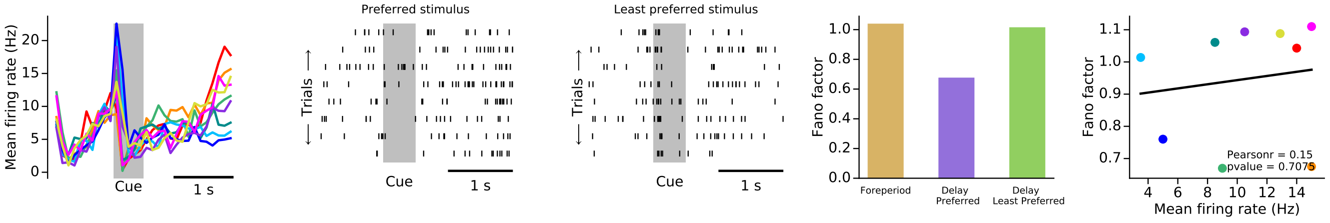


ELV265_1_5518

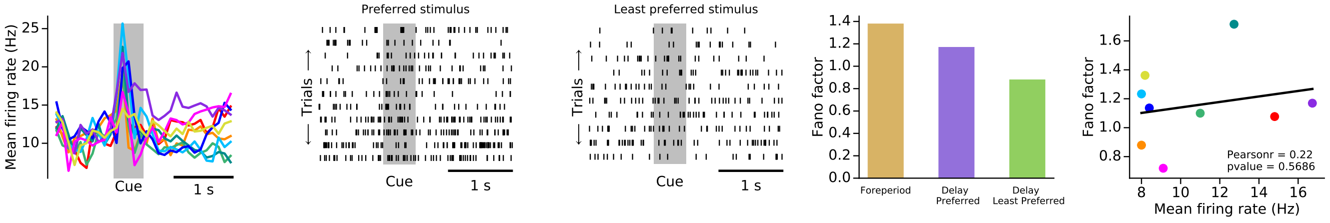

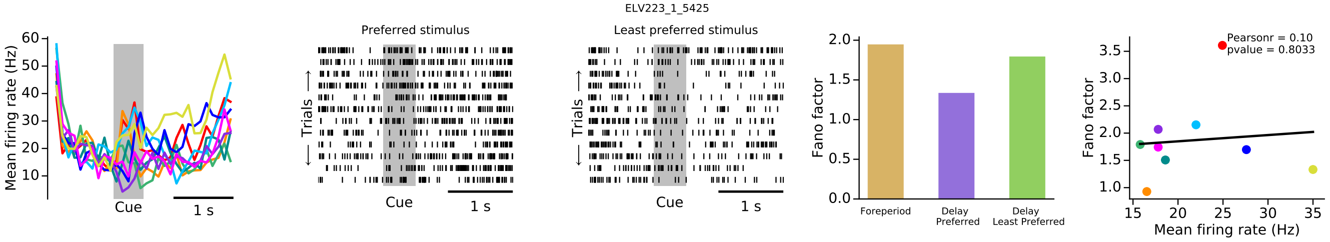
ELV223_1_5425
Preferred stimulus
Least preferred stimulus
Mean firing rate (Hz)
Trials
Cue
1 s
Fano factor
Foreperiod
Delay Preferred
Delay Least Preferred
Pearsonr = 0.10
pvalue = 0.8033
Mean firing rate (Hz)

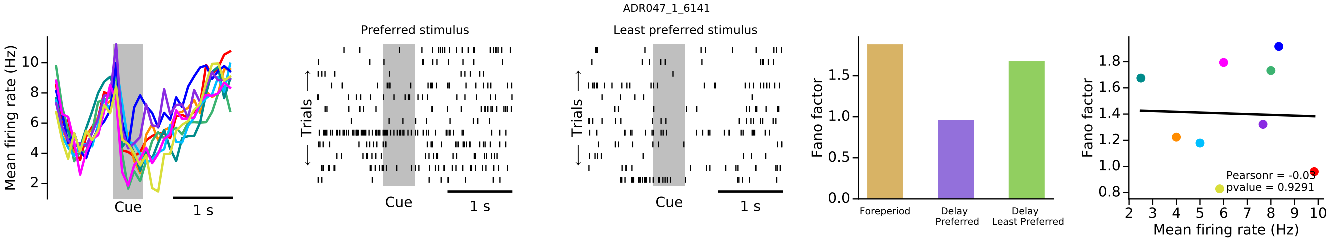
ADR047_1_6141
Preferred stimulus
Least preferred stimulus
Mean firing rate (Hz)
Trials
Cue
1 s
Fano factor
Foreperiod
Delay Preferred
Delay Least Preferred
Pearsonr = -0.03
pvalue = 0.9291
Mean firing rate (Hz)

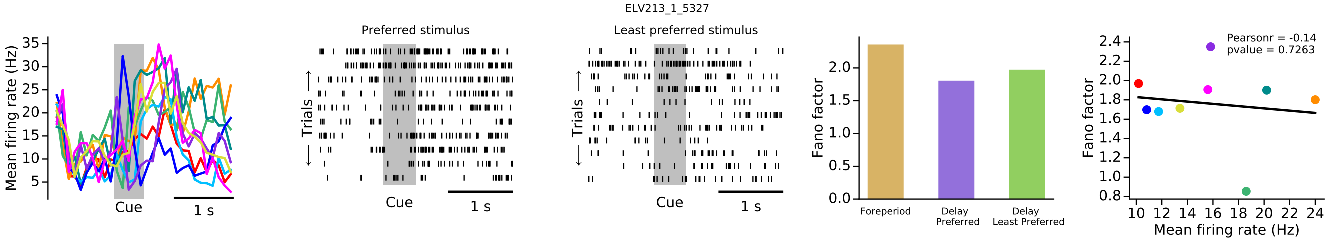
ELV213_1_5327
Preferred stimulus
Least preferred stimulus
Mean firing rate (Hz)
Trials
Cue
1 s
Fano factor
Foreperiod
Delay Preferred
Delay Least Preferred
Pearsonr = -0.14
pvalue = 0.7263
Mean firing rate (Hz)

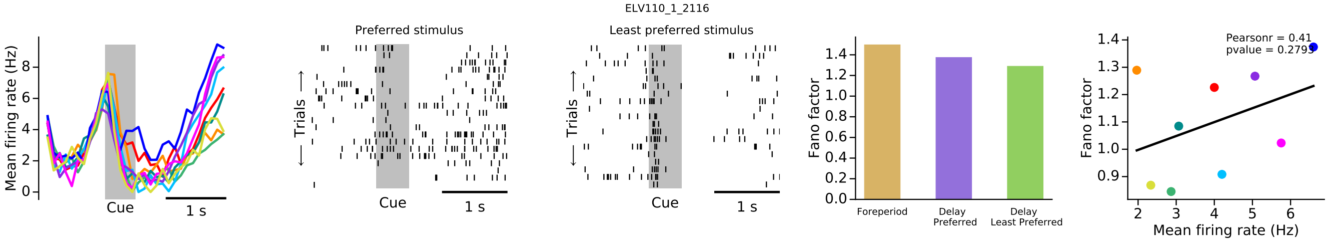
ELV110_1_2116
Preferred stimulus
Least preferred stimulus
Mean firing rate (Hz)
Trials
Cue
1 s
Fano factor
Foreperiod
Delay Preferred
Delay Least Preferred
Pearsonr = 0.41
pvalue = 0.2795
Mean firing rate (Hz)

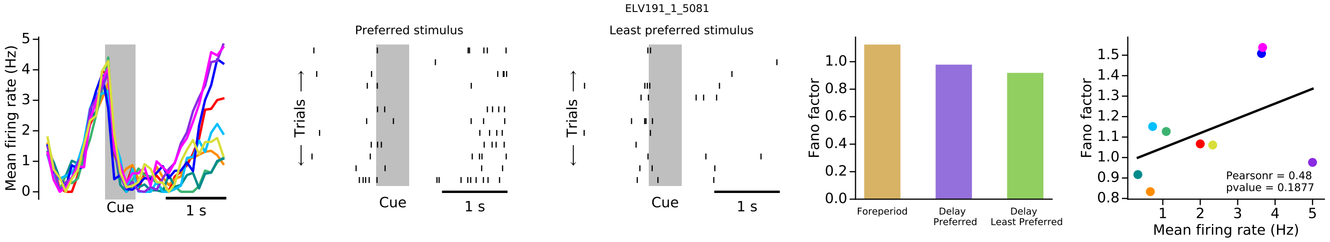
ELV191_1_5081
Preferred stimulus
Least preferred stimulus
Mean firing rate (Hz)
Trials
Cue
1 s
Fano factor
Foreperiod
Delay Preferred
Delay Least Preferred
Pearsonr = 0.48
pvalue = 0.1877
Mean firing rate (Hz)

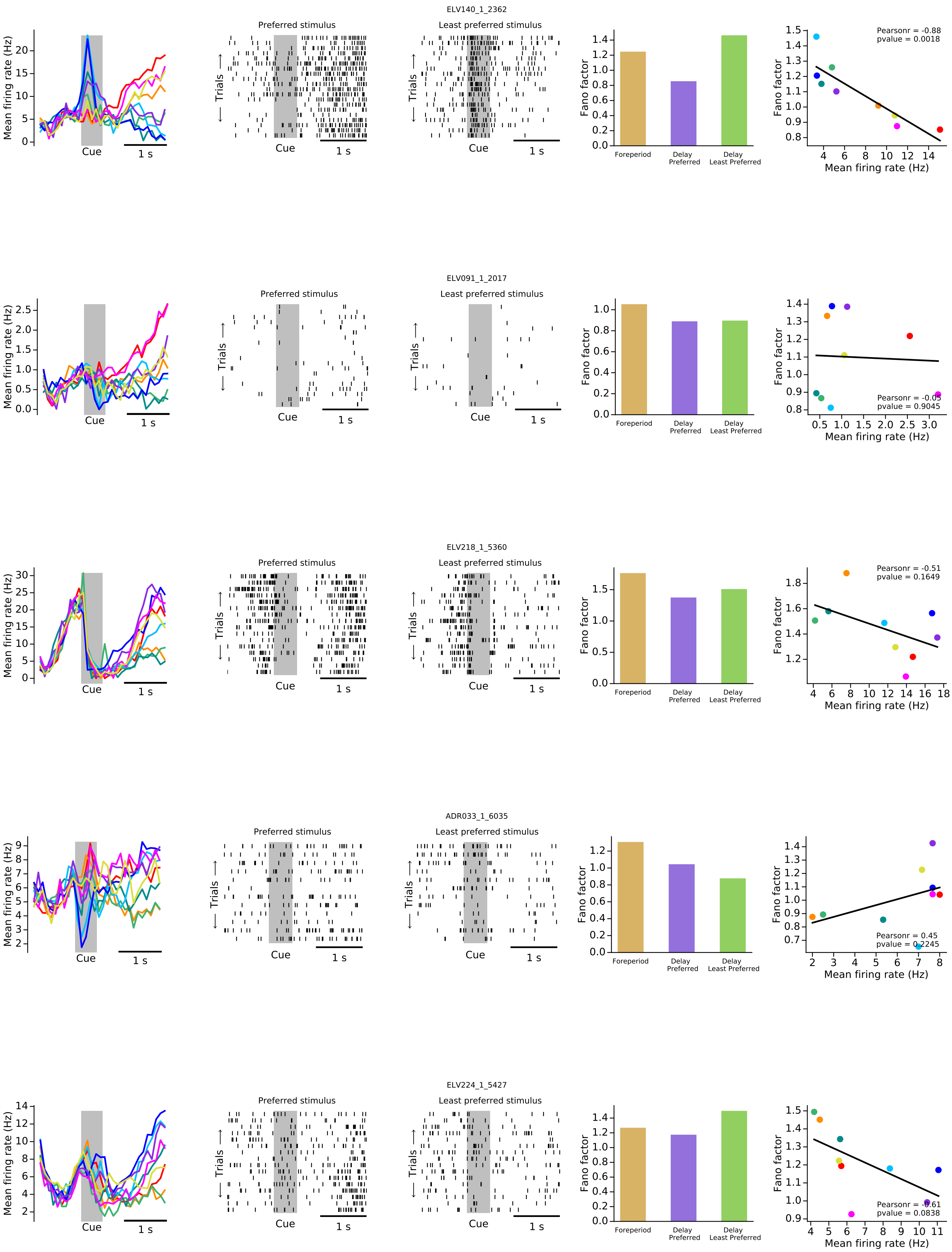
ELV140_1_2362
Preferred stimulus
Least preferred stimulus
Mean firing rate (Hz)
Trials
Cue
1 s
Fano factor
Foreperiod
Delay Preferred
Delay Least Preferred
Pearsonr = -0.88
pvalue = 0.0018
Mean firing rate (Hz)
ELV091_1_2017
Preferred stimulus
Least preferred stimulus
Pearsonr = -0.05
pvalue = 0.9045
ELV218_1_5360
Preferred stimulus
Least preferred stimulus
Pearsonr = -0.51
pvalue = 0.1649
ADR033_1_6035
Preferred stimulus
Least preferred stimulus
Pearsonr = 0.45
pvalue = 0.2245
ELV224_1_5427
Preferred stimulus
Least preferred stimulus
Pearsonr = -0.61
pvalue = 0.0838

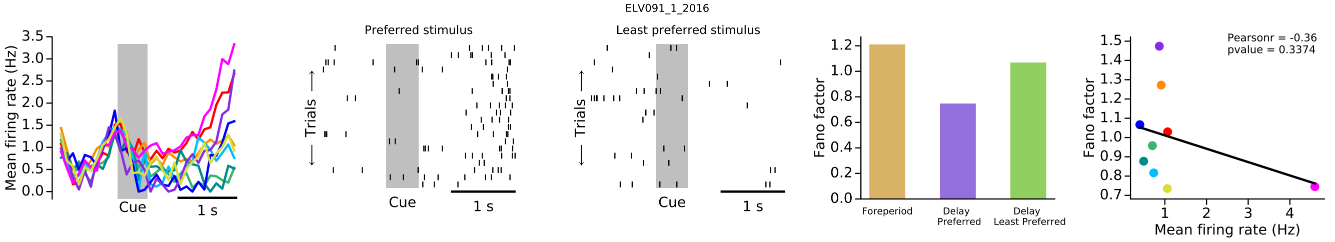
ELV091_1_2016
Preferred stimulus
Least preferred stimulus
Mean firing rate (Hz)
Trials
Cue
1 s
Fano factor
Foreperiod
Delay Preferred
Delay Least Preferred
Pearsonr = -0.36
pvalue = 0.3374
Mean firing rate (Hz)

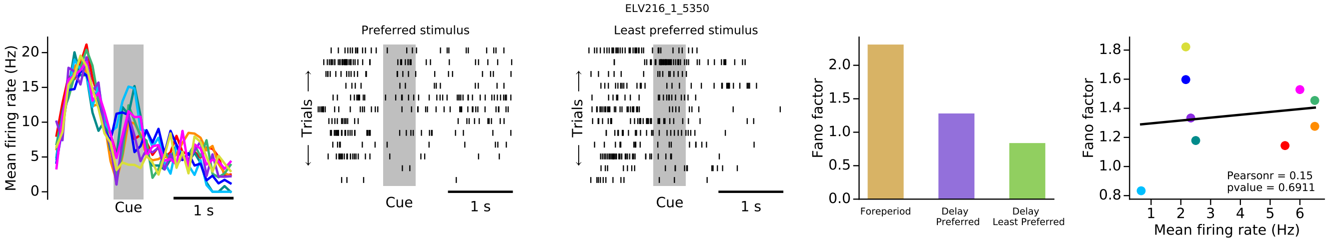
ELV216_1_5350
Preferred stimulus
Least preferred stimulus
Mean firing rate (Hz)
Trials
Cue
1 s
Fano factor
Foreperiod
Delay Preferred
Delay Least Preferred
Pearsonr = 0.15
pvalue = 0.6911
Mean firing rate (Hz)

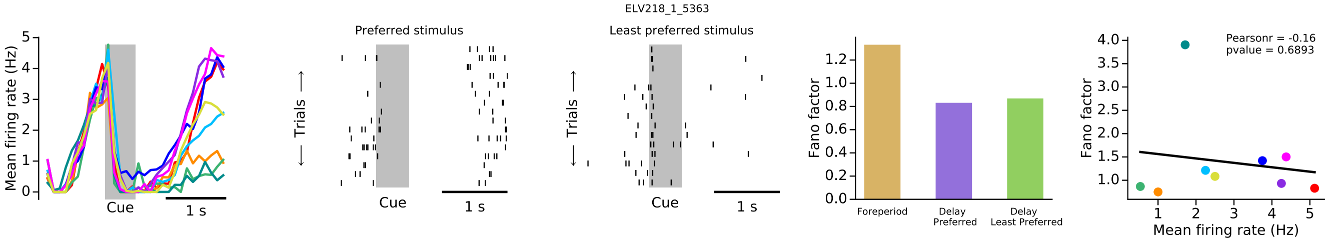
ELV218_1_5363
Preferred stimulus
Least preferred stimulus
Mean firing rate (Hz)
Trials
Cue
1 s
Fano factor
Foreperiod
Delay Preferred
Delay Least Preferred
Pearsonr = -0.16
pvalue = 0.6893
Mean firing rate (Hz)

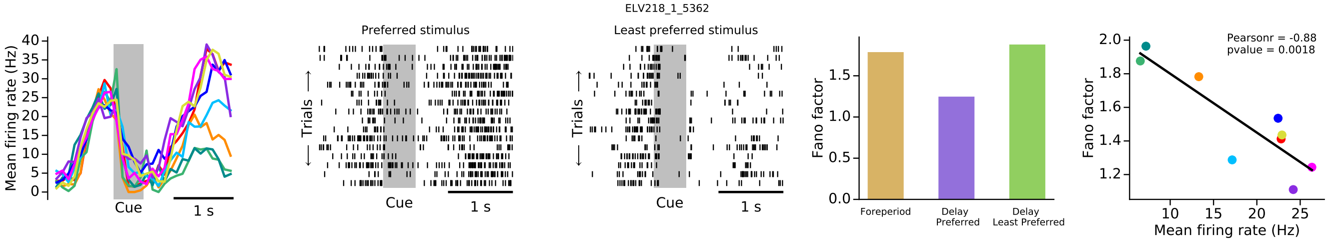
ELV218_1_5362
Preferred stimulus
Least preferred stimulus
Mean firing rate (Hz)
Trials
Cue
1 s
Fano factor
Foreperiod
Delay Preferred
Delay Least Preferred
Pearsonr = -0.88
pvalue = 0.0018
Mean firing rate (Hz)

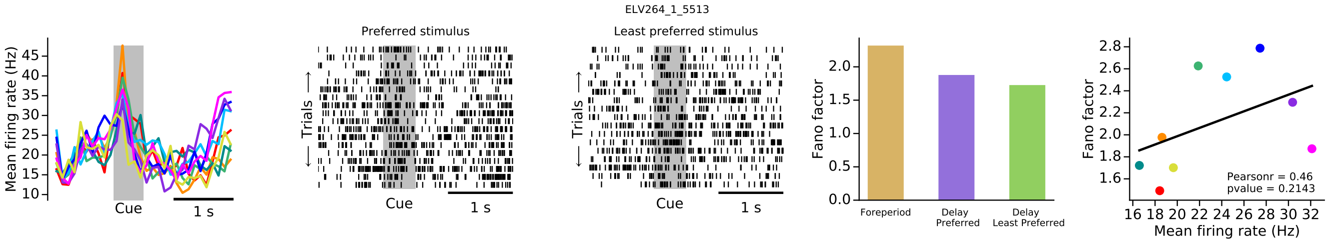
ELV264_1_5513
Preferred stimulus
Least preferred stimulus
Mean firing rate (Hz)
Trials
Cue
1 s
Fano factor
Foreperiod
Delay Preferred
Delay Least Preferred
Pearsonr = 0.46
pvalue = 0.2143
Mean firing rate (Hz)

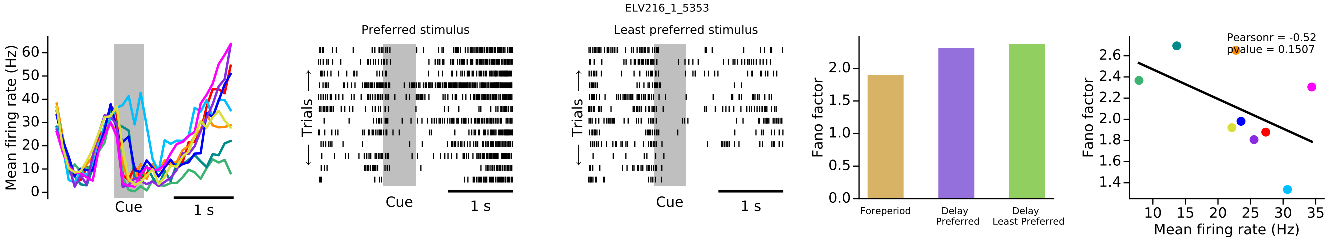
ELV216_1_5353
Mean firing rate (Hz)
Cue
1 s
Preferred stimulus
Trials
Least preferred stimulus
Fano factor
Foreperiod
Delay Preferred
Delay Least Preferred
Pearsonr = -0.52
pvalue = 0.1507
Mean firing rate (Hz)

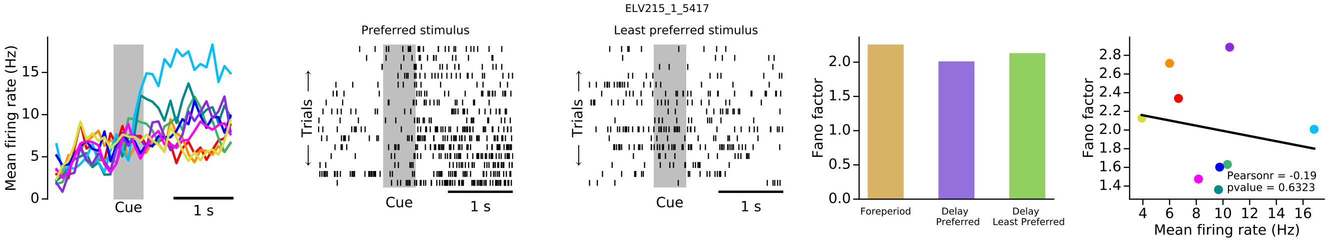
ELV215_1_5417
Mean firing rate (Hz)
Cue
1 s
Preferred stimulus
Trials
Least preferred stimulus
Fano factor
Foreperiod
Delay Preferred
Delay Least Preferred
Pearsonr = -0.19
pvalue = 0.6323
Mean firing rate (Hz)

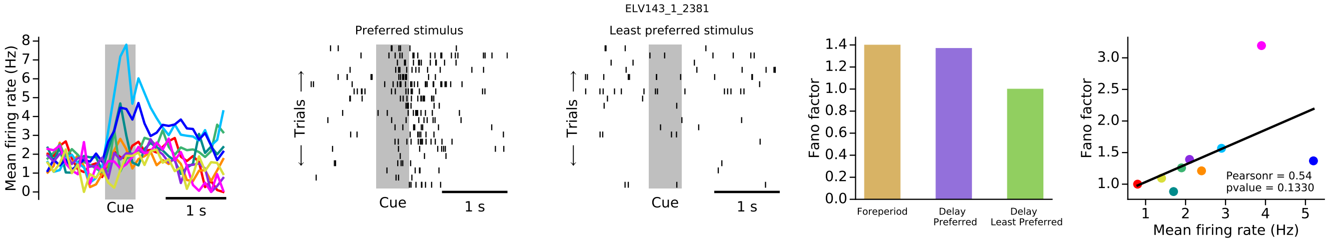
ELV143_1_2381
Mean firing rate (Hz)
Cue
1 s
Preferred stimulus
Trials
Least preferred stimulus
Fano factor
Foreperiod
Delay Preferred
Delay Least Preferred
Pearsonr = 0.54
pvalue = 0.1330
Mean firing rate (Hz)

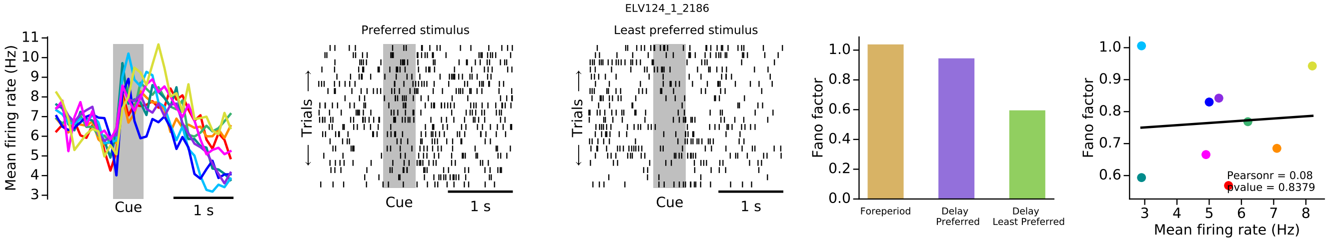
ELV124_1_2186
Mean firing rate (Hz)
Cue
1 s
Preferred stimulus
Trials
Least preferred stimulus
Fano factor
Foreperiod
Delay Preferred
Delay Least Preferred
Pearsonr = 0.08
pvalue = 0.8379
Mean firing rate (Hz)

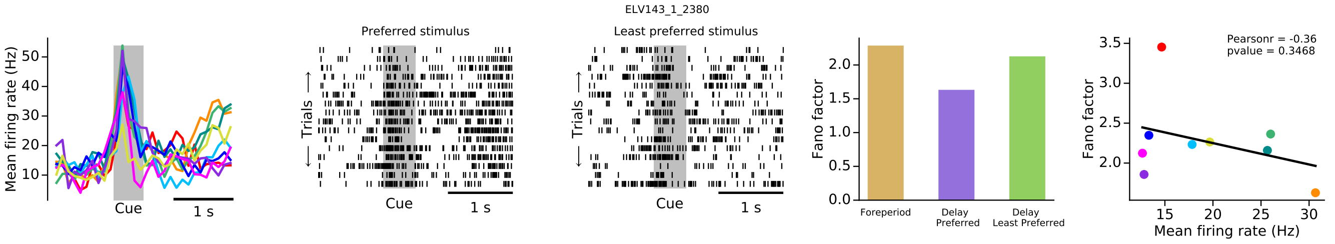
ELV143_1_2380
Mean firing rate (Hz)
Cue
1 s
Preferred stimulus
Trials
Least preferred stimulus
Fano factor
Foreperiod
Delay Preferred
Delay Least Preferred
Pearsonr = -0.36
pvalue = 0.3468
Mean firing rate (Hz)

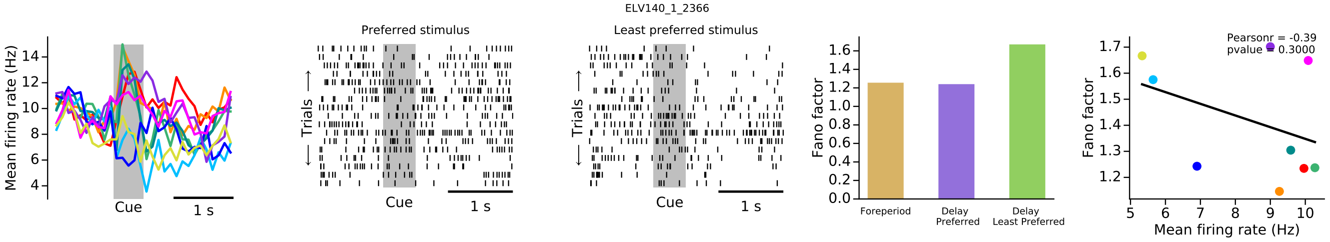
ELV140_1_2366
Preferred stimulus
Least preferred stimulus
Mean firing rate (Hz)
Cue
1 s
Trials
Fano factor
Foreperiod
Delay Preferred
Delay Least Preferred
Pearsonr = -0.39
pvalue = 0.3000
Mean firing rate (Hz)

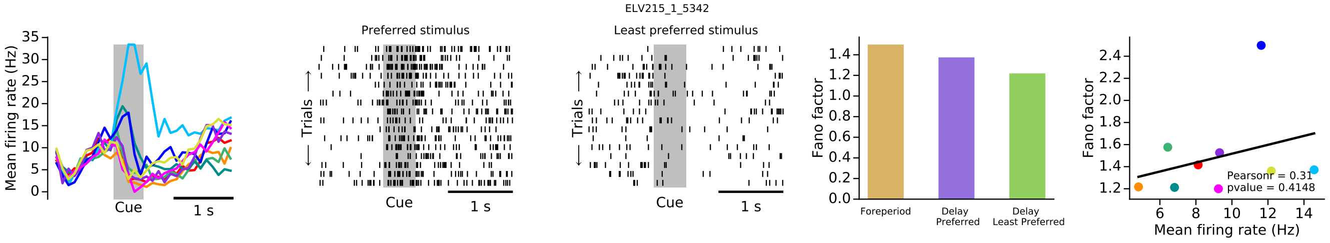
ELV215_1_5342
Preferred stimulus
Least preferred stimulus
Mean firing rate (Hz)
Cue
1 s
Trials
Fano factor
Foreperiod
Delay Preferred
Delay Least Preferred
Pearsonr = 0.31
pvalue = 0.4148
Mean firing rate (Hz)

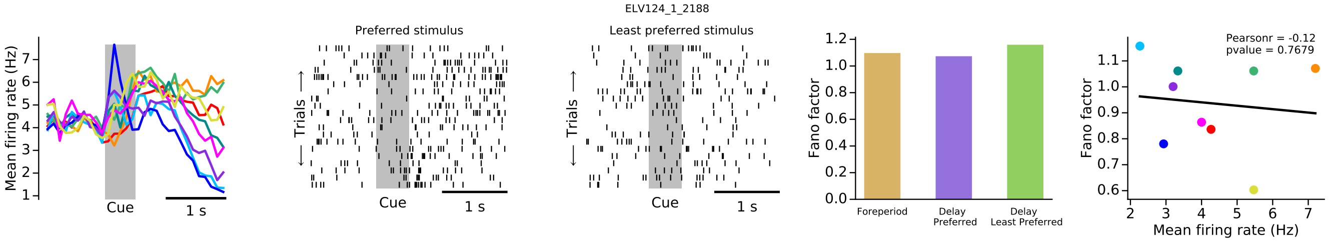
ELV124_1_2188
Preferred stimulus
Least preferred stimulus
Mean firing rate (Hz)
Cue
1 s
Trials
Fano factor
Foreperiod
Delay Preferred
Delay Least Preferred
Pearsonr = -0.12
pvalue = 0.7679
Mean firing rate (Hz)

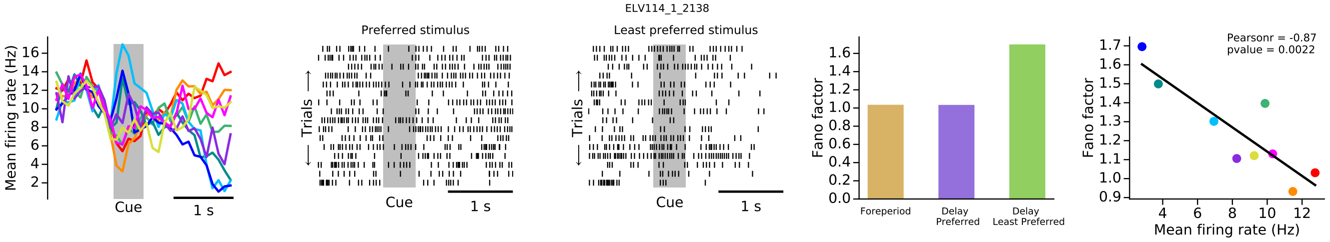
ELV114_1_2138
Preferred stimulus
Least preferred stimulus
Mean firing rate (Hz)
Cue
1 s
Trials
Fano factor
Foreperiod
Delay Preferred
Delay Least Preferred
Pearsonr = -0.87
pvalue = 0.0022
Mean firing rate (Hz)

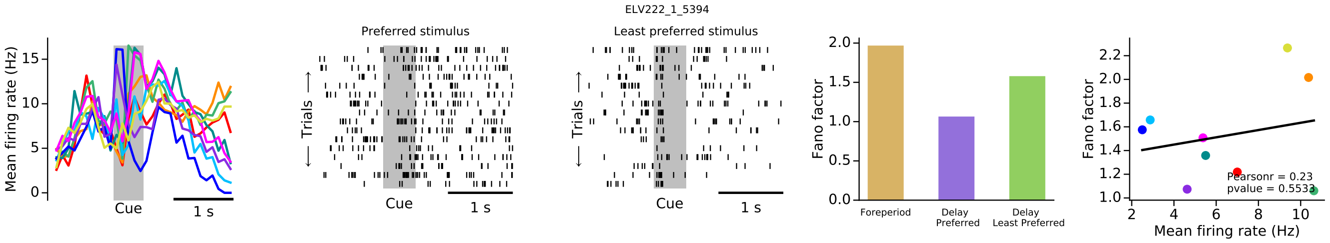
ELV222_1_5394
Preferred stimulus
Least preferred stimulus
Mean firing rate (Hz)
Cue
1 s
Trials
Fano factor
Foreperiod
Delay Preferred
Delay Least Preferred
Pearsonr = 0.23
pvalue = 0.5533
Mean firing rate (Hz)

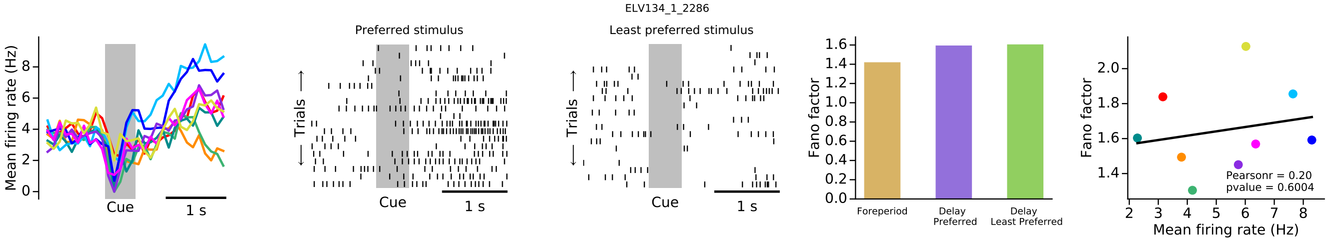
ELV134_1_2286
Preferred stimulus
Least preferred stimulus
Mean firing rate (Hz)
Trials
Cue
1 s
Fano factor
Foreperiod
Delay Preferred
Delay Least Preferred
Pearsonr = 0.20
pvalue = 0.6004
Mean firing rate (Hz)

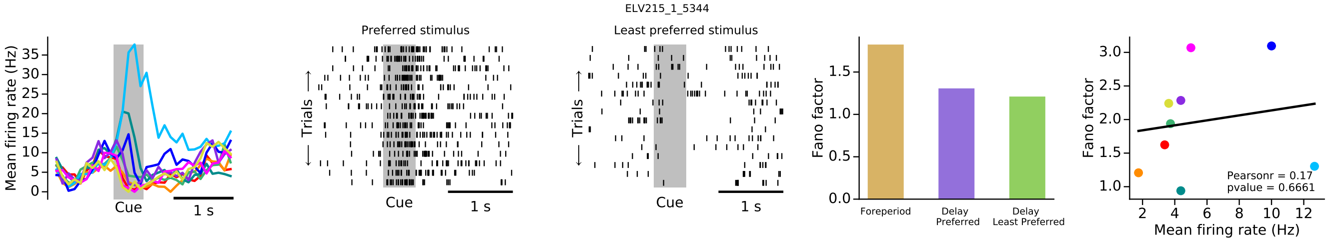
ELV215_1_5344
Preferred stimulus
Least preferred stimulus
Mean firing rate (Hz)
Trials
Cue
1 s
Fano factor
Foreperiod
Delay Preferred
Delay Least Preferred
Pearsonr = 0.17
pvalue = 0.6661
Mean firing rate (Hz)

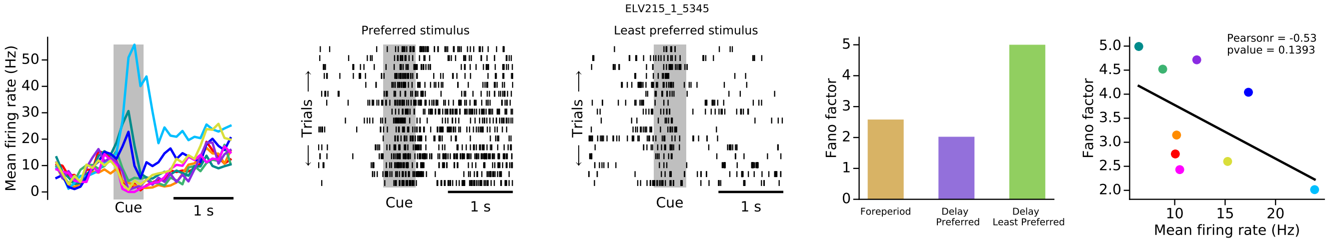
ELV215_1_5345
Preferred stimulus
Least preferred stimulus
Mean firing rate (Hz)
Trials
Cue
1 s
Fano factor
Foreperiod
Delay Preferred
Delay Least Preferred
Pearsonr = -0.53
pvalue = 0.1393
Mean firing rate (Hz)

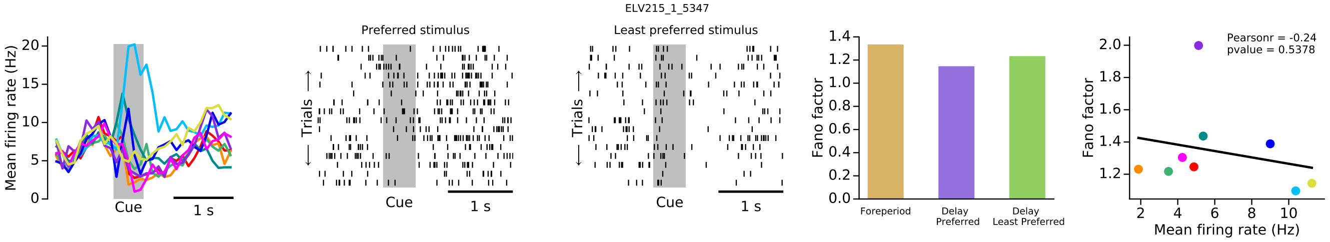
ELV215_1_5347
Preferred stimulus
Least preferred stimulus
Mean firing rate (Hz)
Trials
Cue
1 s
Fano factor
Foreperiod
Delay Preferred
Delay Least Preferred
Pearsonr = -0.24
pvalue = 0.5378
Mean firing rate (Hz)

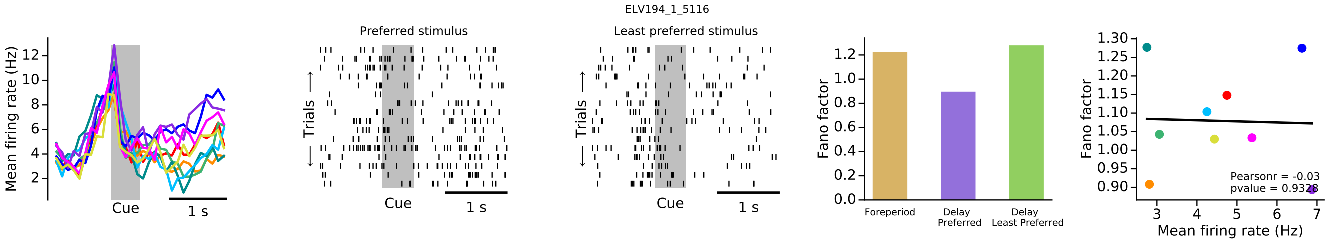
ELV194_1_5116
Preferred stimulus
Least preferred stimulus
Mean firing rate (Hz)
Trials
Cue
1 s
Fano factor
Foreperiod
Delay Preferred
Delay Least Preferred
Pearsonr = -0.03
Mean firing rate (Hz)

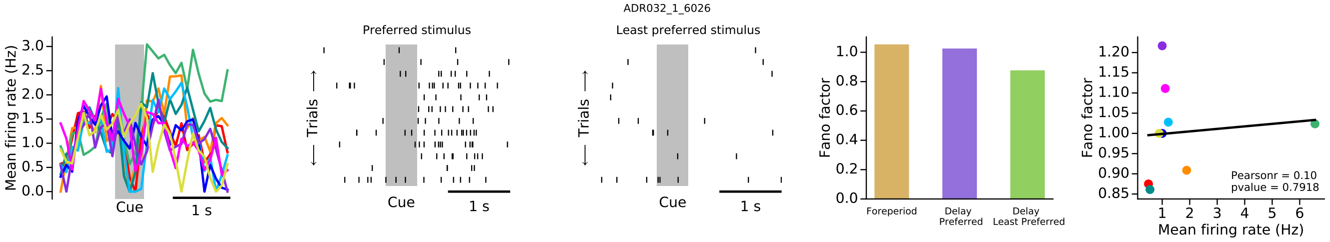
ADR032_1_6026
Mean firing rate (Hz)
Cue
1 s
Preferred stimulus
Least preferred stimulus
Trials
Fano factor
Foreperiod
Delay Preferred
Delay Least Preferred
Pearsonr = 0.10
pvalue = 0.7918
Mean firing rate (Hz)

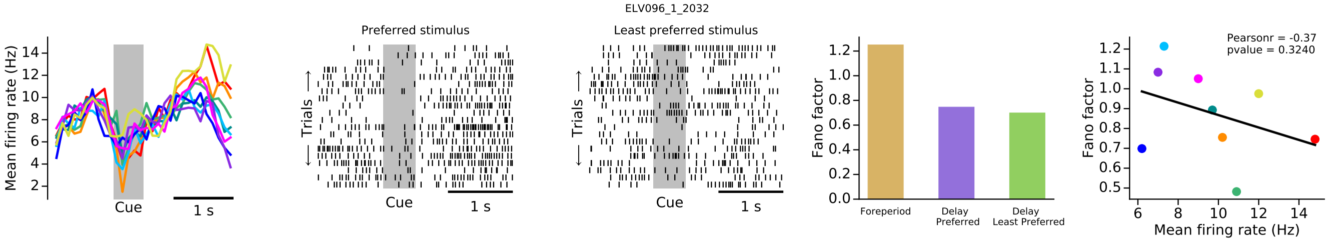
ELV096_1_2032
Mean firing rate (Hz)
Cue
1 s
Preferred stimulus
Least preferred stimulus
Trials
Fano factor
Foreperiod
Delay Preferred
Delay Least Preferred
Pearsonr = -0.37
pvalue = 0.3240
Mean firing rate (Hz)

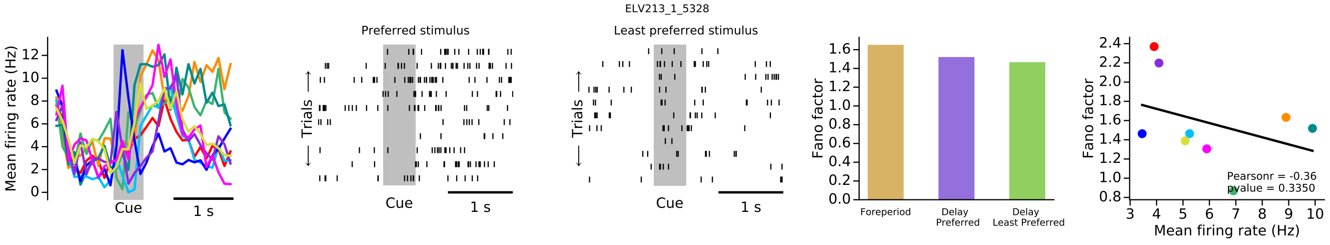
ELV213_1_5328
Mean firing rate (Hz)
Cue
1 s
Preferred stimulus
Least preferred stimulus
Trials
Fano factor
Foreperiod
Delay Preferred
Delay Least Preferred
Pearsonr = -0.36
pvalue = 0.3350
Mean firing rate (Hz)

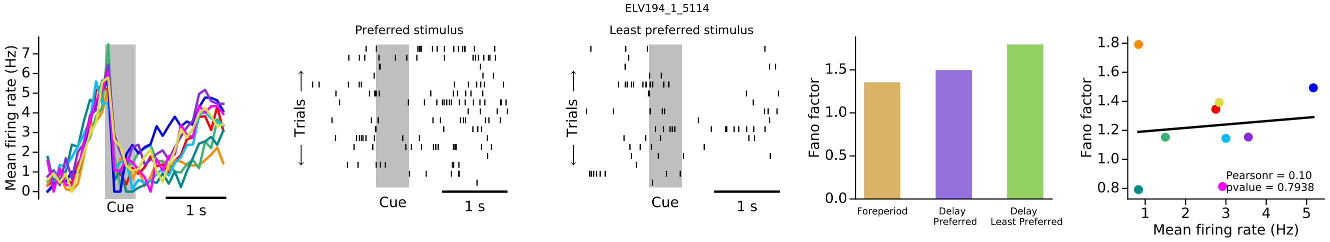
ELV194_1_5114
Mean firing rate (Hz)
Cue
1 s
Preferred stimulus
Least preferred stimulus
Trials
Fano factor
Foreperiod
Delay Preferred
Delay Least Preferred
Pearsonr = 0.10
pvalue = 0.7938
Mean firing rate (Hz)

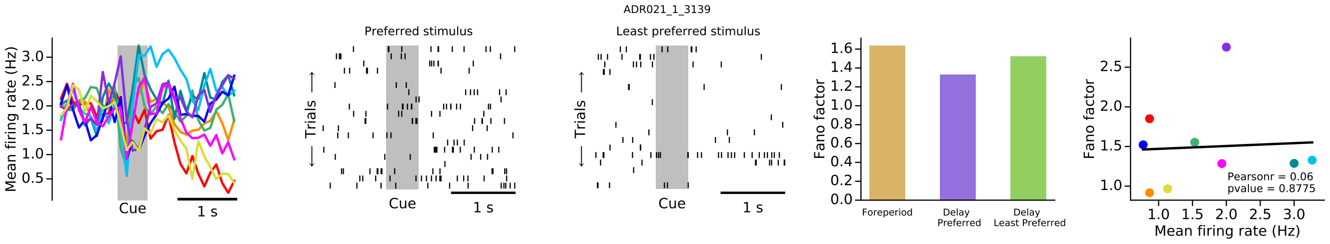
ADR021_1_3139
Mean firing rate (Hz)
Cue
1 s
Preferred stimulus
Least preferred stimulus
Trials
Fano factor
Foreperiod
Delay Preferred
Delay Least Preferred
Pearsonr = 0.06
pvalue = 0.8775
Mean firing rate (Hz)

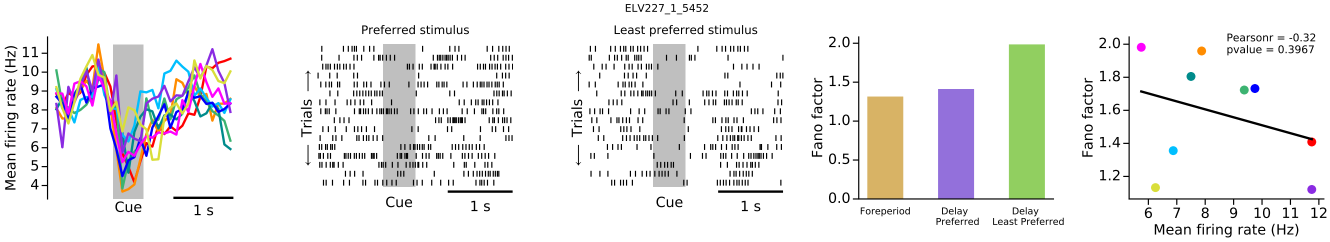
ELV227_1_5452
Mean firing rate (Hz)
Cue
1 s
Preferred stimulus
Least preferred stimulus
Trials
Fano factor
Foreperiod
Delay Preferred
Delay Least Preferred
Pearsonr = -0.32
pvalue = 0.3967
Mean firing rate (Hz)
